# Intrinsically disordered regions within transcription factors impact DNA binding through both autoinhibition and independent DNA binding

**DOI:** 10.64898/2026.11.20.768217

**Authors:** Michael G. B. Hayes, Ziwei Chen, Albert A. Lee, Stephen Plassmeyer, Scott Longwell, Alex S. Holehouse, Anshul Kundaje, Polly M. Fordyce

## Abstract

Transcription factors (TFs) recognize DNA through structured DNA-binding domains (DBDs), but the contributions of their flanking intrinsically disordered regions (IDRs) to DNA binding remain poorly understood. Here, we dissected how IDRs impact DNA binding *in vitro* by investigating three paralogous *Saccharomyces cerevisiae* TFs with highly similar DNA-binding domains but divergent IDRs (Msn2, Msn4, and Com2). Using high-throughput microfluidic binding assays, we quantified >17,000,000 TF–DNA relative binding energies between Msn2, Msn4, and Com2 variant constructs and DNA libraries comprised of systematic substitutions or genomic sequences. These measurements reveal that IDRs do not alter core ‘motif’ specificities but alter DNA binding affinities via two distinct mechanisms. First, the Msn2 and Msn4 IDRs autoinhibit their cognate DBDs, imposing a uniform free-energy penalty that depends on the combination of DNA-binding domain and IDR. Second, di-arginine pairs within the Msn2 IDR selectively bind low-complexity, homopolymeric DNA with very weak affinities only detectable under avid binding conditions. Neural-network models trained on *in vivo* Msn2 binding data (ChEC-seq) reveal similar preferences for clustered homopolymeric sequences, suggesting this weak binding impacts genomic localization in cells via intermolecular avidity. These results establish that IDRs influence TF–DNA binding through diverse mechanisms operating both independently of and in concert with DBD motif recognition.

## Introduction

Transcription factors (TFs) are the primary regulators of gene expression. Eukaryotic TFs are typically composed of structured DNA binding domains (DBDs) that bind DNA via direct recognition of short (5-10 base pair) motifs^1^ and longer intrinsically disordered regions (IDRs)^2,3^ that bind transcriptional co-activators and - repressors to alter transcriptional activity^4^. Traditionally, TF DBDs and IDRs were thought to act independently: DBDs and effector domains often remained functional when swapped between different TFs^2,5^ and many high-throughput *in vitro* studies have found minimal differences in motif preferences between full-length TFs and their DBDs alone^6,7^. As a result of this assumption and the technical difficulties associated with expressing and purifying proteins with long IDRs, most measurements profiling DNA binding specificities *in vitro* have used DBD-only constructs^6–9,9–20^.

Emerging studies have suggested that TF IDRs can play integral roles in TF-DNA binding via several distinct mechanisms. In some cases, IDRs reduce TF-DNA binding affinity through autoinhibition mediated by interactions between negatively charged D/E residues or phosphorylated S/T residues and positively charged DBD residues (as for MAX^21^, p53^22,23^, Ets1^24,25^, HMGB1^26^, and AntP^27^). This autoinhibition can speed TF target search kinetics and prevent kinetic trapping in the presence of non-specific competitive DNA binding sites, thereby *increasing* specificity^27^. In other cases, TF IDRs interact with other IDRs to form either homotypic (as for Sp1^28^) or heterotypic protein complexes (as for RelA in the presence of p53 or RPS3^29^) with distinct DNA specificities. In principle, TF IDRs could also interact with the DBD to stabilize particular conformations within the structural ensemble and alter specificity.

More recently, studies investigating the stress-responsive C2H2-zinc finger TF Msn2 from budding yeast suggested that, in cells, its large (641 amino acid) IDR was necessary and often sufficient for correct localization to target promoters^30^ via a distinct mechanism: direct, low affinity binding to DNA. Systematic mutagenesis of Msn2’s IDR *in vivo* implicated dispersed aliphatic and aromatic amino acids in this DNA recognition^31,32^.

However, IDRs in cells can simultaneously participate in all of the aforementioned interactions as well as condensate formation^33^, making it challenging to distinguish and quantify each mechanism. While *in vitro* single-molecule experiments with Msn2 recently suggested that its IDR facilitates association with DNA through independent charge-mediated IDR-DNA binding^34^, it remains unclear how widespread this binding is, whether specific sequences are preferred, and whether this IDR-DNA binding adequately explains *in vivo* observations.

Here, we exploited the same natural system (the paralogous *S. cerevisiae* C2H2 zinc finger TFs Msn2, Msn4, and Com2) to systematically examine the impacts of IDRs on TF-DNA binding *in vitro*. Using the high-throughput microfluidic binding assays BET-seq^35^ and MITOMI^36,37^, we quantified 17,493,410 TF–DNA relative binding energies (ΔΔGs) across two different DNA sequence libraries and found that IDRs alter binding in two distinct ways. First, full-length (FL) Msn2 and Msn4 constructs bound DNA with weaker absolute affinities (*K*_d_s) than their DBDs alone, consistent with autoinhibition. Chimeric constructs revealed that this autoinhibition depended on specific IDR/DBD combinations rather than simple steric occlusion. Second, the Msn2 IDR (but not the Msn4 or Com2 IDRs) independently bound DNA very weakly with a preference for homopolymeric sequences, with di-arginine pairs proximal to the DBD being both necessary and sufficient for DNA binding. Neural network models (BPNet^38^) trained on *in vivo* transcription factor binding profiles for FL and IDR-only Msn2 constructs recovered similar preferences for homopolymeric T-rich sequences, suggesting this binding is physiologically relevant in cells. Comparisons between predictions made by models trained on *in vitro* and *in vivo* data suggested that genomic localization depends substantially on both direct TF-DNA interactions and interactions between TFs and other nuclear molecules (*e.g.* protein-protein binding, competition with nucleosomes, *etc*.). Together, these results establish that IDRs can regulate TF-DNA binding via multiple distinct mechanisms independent of and in concert with DBD motif recognition.

## Results

### Msn2, Msn4, and Com2 as a model system for investigating effects of IDRs on DNA binding

Msn2, Msn4, and Com2 have highly conserved C2H2-type zinc finger DBDs with identical nucleotide-contacting residues^39^ and bind nearly identical motifs consisting of a central AGGGG sequence (**Fig. 1A**); Com2 prefers a slightly extended version of this motif with an upstream flanking AT dinucleotide^40^. These 59-62 amino acid (aa) DBDs are found at the C terminus of each TF and are preceded by sequences of variable size (641, 567, and 383 aas for Msn2, Msn4, and Com2, respectively) predicted to be disordered by both Metapredict and AlphaFold2 (**Fig. 1B**)^41–43^. These extensive IDRs are poorly conserved across Msn2, Msn4, and Com2 but share similar sequence properties, with aromatic and hydrophobic residues dispersed between abundant acidic residues throughout (**Fig. 1B, Supplementary Fig. 1**).

**Figure 1.**
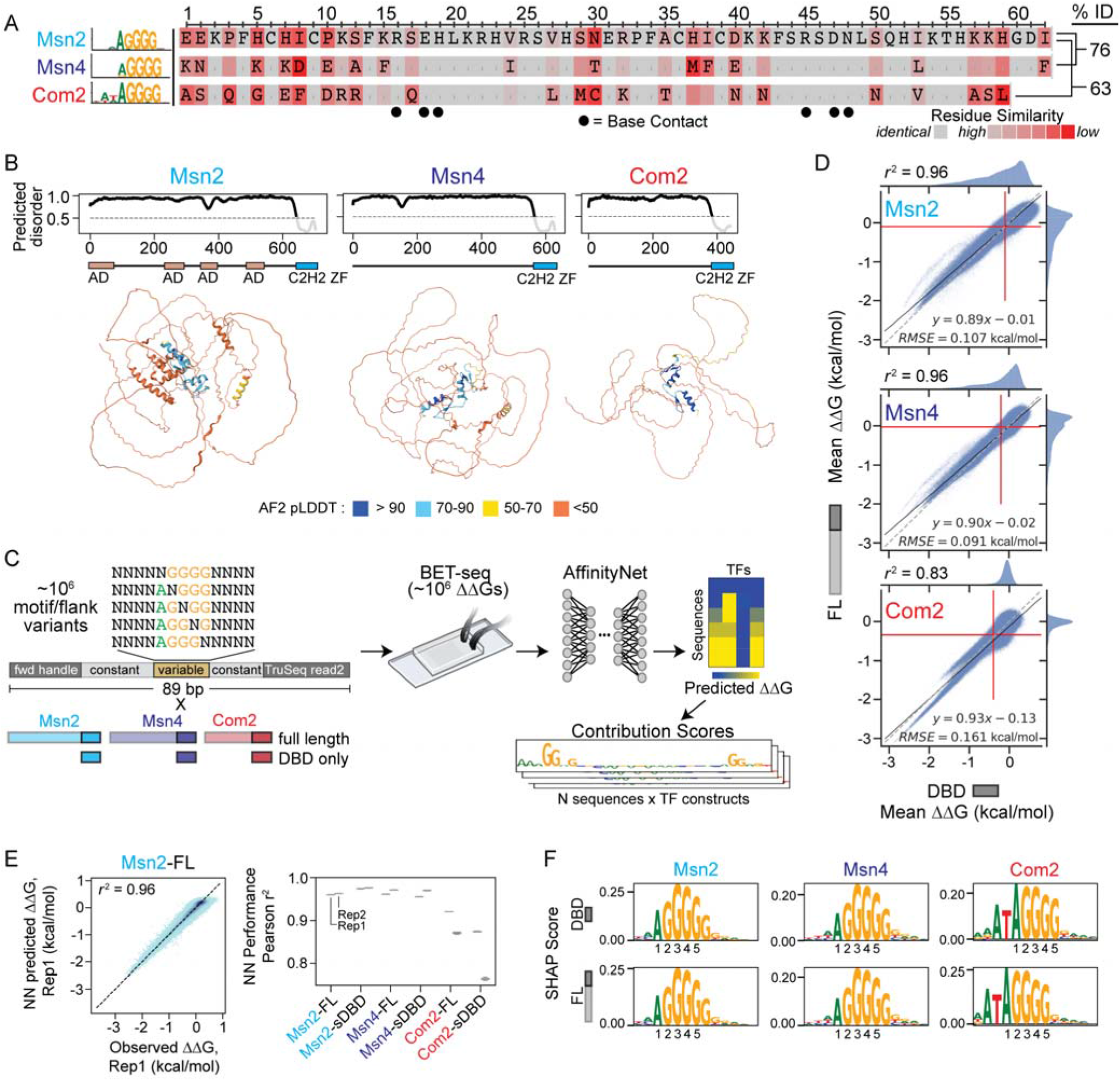
Msn2, Msn4, and Com2 IDRs have little-to-no effect on motif binding specificity. **A** Consensus motifs for Msn2, Msn4, and Com2^40^ (left) and DBD sequence alignment relative to Msn2 (right) indicating identical residues (blank), base-contacting residues (black dots), and all residues colored by amino-acid similarity. **B** Predicted disorder for Msn2, Msn4, and Com2 (top) and their AlphaFold2-predicted structures (bottom). **C** Schematic showing: (1) ‘degenerate motif’ library comprised of mutations to the AGGGG consensus motif and upstream and downstream flanking nucleotides, (2) full-length (FL) and DBD-only TF constructs, and (3) pipeline for BET-seq measurement and AffinityNet analysis. **D** Scatter plots comparing FL *vs.* DBD Msn2 (left), Msn4 (middle), and Com2 (right) relative binding energies (ΔΔGs) to the ‘degenerate motif’ library. Markers denote per-sequence ΔΔGs; dotted gray lines denote 1:1 line; solid black lines denote linear fits; red lines denote thresholds used to delineate ‘bound’ sequences (see **Methods**). Pearson *r*^2^ is reported for all sequences considered ‘bound’ by either DBD or FL constructs. **E** Representative scatter plot showing AffinityNet-predicted *vs.* BET-seq-measured ΔΔGs for FL Msn2 binding the ‘degenerate motif’ library (left) and distributions of Pearson r^2^ values between predicted and observed ΔΔGs across 5-fold test validations for each BET-seq replicate (right). Markers represent individual DNA sequences; pixel darkness indicates sequence density; box plot extents indicate ranges of Pearson *r*^2^ values. **F** Top AffinityNet-derived motif clusters for DBD (top) and FL (bottom) Msn2, Msn4, and Com2 with core consensus motif positions annotated.

To test if Msn2, Msn4, or Com2 IDRs impact DBD motif preferences, we used the BET-seq microfluidic platform^35^ to quantify relative affinities (ΔΔGs) for full-length (FL) and DBD-only constructs each binding to a ‘degenerate motif’ DNA library comprised of systematic substitutions within and around a central consensus AGGGG motif (**Fig. 1C**). Specifically, this library varied the 4 nucleotides up- and downstream of all single nucleotide variations within the motif (1,048,576 total sequences).

In BET-seq, C-terminally monomeric eGFP (meGFP)-tagged TF constructs are expressed via cell-free protein synthesis and then specifically immobilized to anti-GFP nanobody-patterned surfaces beneath pneumatically actuated ‘button’ valves within microfluidic devices **(Supplementary Fig. 2)**. This immobilization enables on-chip purification and subsequent introduction of a DNA library of interest. After interactions between introduced DNA and surface-immobilized TFs reach equilibrium, actuating ‘button’ valves mechanically traps bound DNA to allow separate elution of ‘bound’ and ‘unbound’ fractions. We then count molecules through subsequent deep sequencing of eluted pools to high read depth, making it possible to calculate the probability that a given sequence is bound, which in turn yields ΔΔGs (**Supplementary Fig. 2**).

Overall, we quantified > 6.3 million ΔΔGs for Msn4, Msn2, and Com2 FL and DBD-only constructs across 2 devices each. Measured ΔΔGs for each DNA sequence were highly reproducible across devices (Pearson *r*^2^ = 0.88-0.95, computed by considering sequences that bound with ΔΔGs > 2 standard deviations below an estimated Gaussian noise floor; **Methods**) and spanned a wide dynamic range (∼ 3.5 kcal/mol; **Supplementary Fig. 3**). Comparisons of median measured ΔΔGs between FL Msn2 and Msn4 revealed very similar binding specificities (*r*^2^ = 0.94); Com2 specificity diverged somewhat from Msn4 and Msn2 (*r*^2^ = 0.45 and 0.48, respectively) (**Supplementary Fig. 4**). STREME^44^ analysis of common and differentially bound sequences confirmed that while all paralogs tightly bound the AGGGG motif with upstream AA flanks, Com2 strongly preferred motifs with an AT upstream flank while Msn2 preferred a more degenerate tetra-G motif, consistent with prior reports^39^ (**Supplementary Fig. 4**). Together, these results confirm that BET-seq returns reliable DNA motif specificity information for each TF.

### Msn2, Msn4, and Com2 IDRs largely do not alter motif or flanking nucleotide specificities

Next, we assessed impacts of each IDR on TF specificity by comparing measured ΔΔGs for FL and DBD-only versions of each TF interacting with the ‘degenerate motif’ library (**Fig. 1D**). Binding specificity landscapes were correlated at levels similar to or exceeding experimental replicates (*r*^2^ = 0.96, 0.96, and 0.83 for Msn2, Msn4, and Com2, respectively; **Fig. 1D, Supplementary Fig. 3**), suggesting that each IDR only minimally impacts motif specificity. Consistent with this, composite position-specific affinity matrices (PSAMs) quantifying energetic impacts of single nucleotide substitutions both within and flanking the consensus AGGGG motif revealed identical motif preferences for the FL and DBD-only constructs for Msn2 and Msn4; Com2 FL and DBD-only constructs exhibited slight differences in nucleotide preferences at positions 1 and −1 (**Supplementary Figs. 5-9**). To model these data without assumptions of nucleotide independence and additivity, we constructed a convolutional neural network model designed to predict both BET-seq relative binding energies and the differences in ΔΔGs between DBD and IDR-containing constructs with the same DBD (AffinityNet). We then used DeepLIFT/DeepSHAP^45,46^ to estimate the contribution of each nucleotide in each sequence to predicted binding and clustered per-nucleotide contribution scores of bound sequences to identify distinct contributing motifs (**Methods**). AffinityNet models trained on the same data outperformed PSAMs (AffinityNet 0.97 > *r*^2^ > 0.87 *vs.* PSAM 0.78 > *r*^2^ > 0.64 across constructs; **Fig. 1E, Supplementary Figs. 10-11**) but still revealed nearly identical motif preferences between Msn2 and Msn4 constructs (**Fig. 1F, Supplementary Figs. 12-13**). For FL-Com2, AffinityNet models confirmed a preference for the GGGGG motif variant relative to Com2’s DBD, and additionally revealed a slight preference for 6xG stretches more generally (**Supplementary Figs. 14-16**).

### Msn2 and Msn4 IDRs reduce motif binding affinity in a manner consistent with autoinhibition

Results thus far establish that IDRs have little to no impact on relative motif binding specificities (ΔΔGs) but do not provide information about absolute affinities (*K*_d_s). To address this, we used the MITOMI microfluidic platform^36,37^ to quantify *K*_d_s for 8 fiducial sequences for each construct spanning a wide range of relative affinities (**Fig. 2A**). Specifically, we programmed device chambers by aligning devices to arrays of fluorescently labeled oligonucleotides printed at multiple concentrations, patterned device surfaces with surface-immobilized meGFP-tagged TF constructs, exposed solubilized printed DNA to the immobilized TFs, and then used pneumatic “button” valves to trap TF-bound DNA. After washing out unbound DNA, we imaged to quantify fractional occupancies (DNA/protein intensities) and soluble DNA concentrations (**Supplementary Fig. 17**).

**Figure 2.**
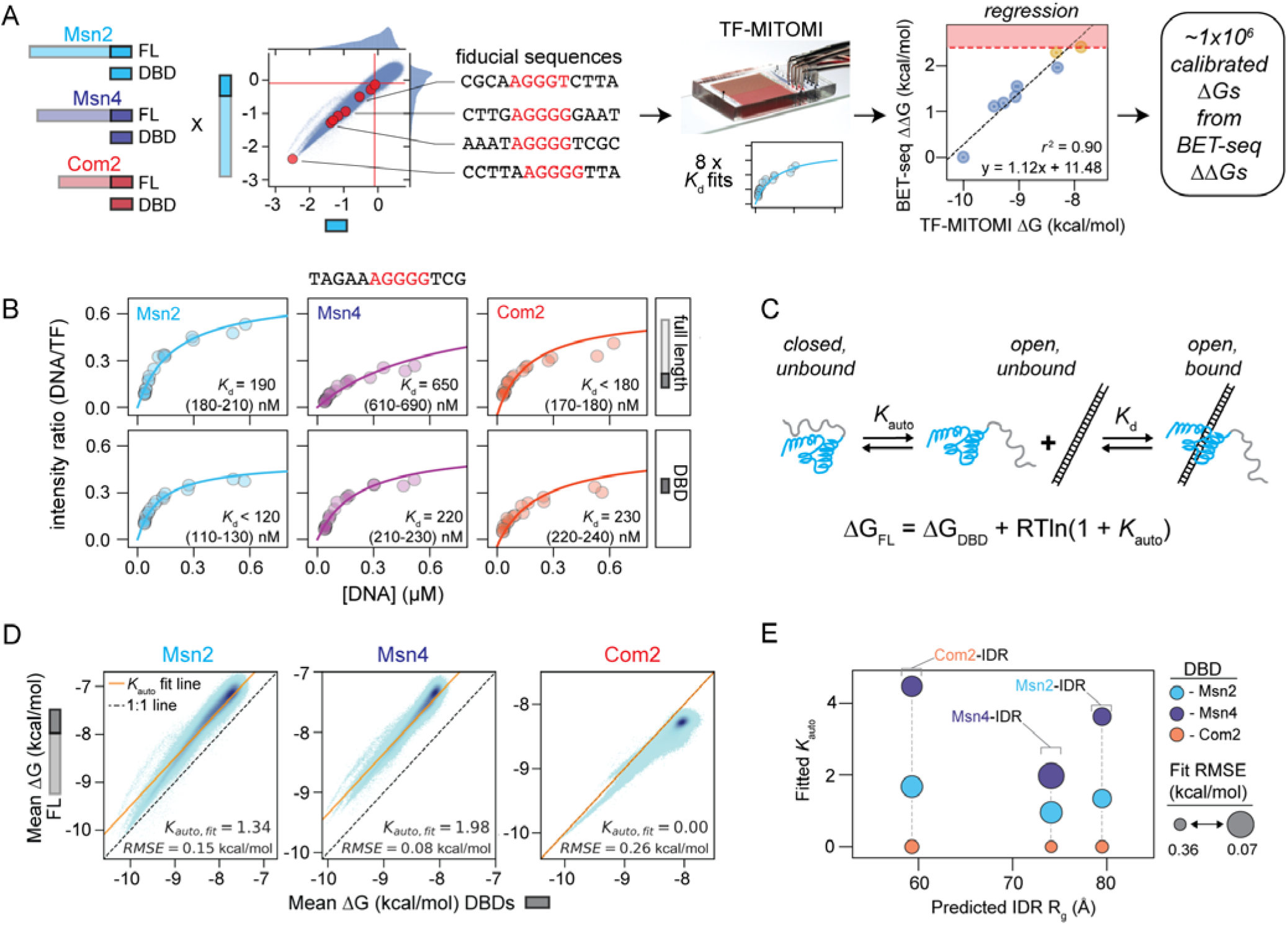
Msn2 and Msn4 IDRs autoinhibit binding to weaken affinities. **A** Schematic illustrating pipeline for calibrating BET-seq-measured relative affinities (ΔΔGs) in absolute space using MITOMI measurements of *K*_d_s for fiducial sequences; red box highlights limit of detection for weak binders. **B** Representative MITOMI concentration-dependent binding curves for FL and DBD-only constructs of Msn2, Msn4, and Com2. Markers correspond to measured intensity ratios across individual chambers; solid lines indicate fits to a Langmuir isotherm with annotated median fit *K*_d_s and interquartile ranges (all curves shown in **Supplementary File 1**). **C** Schematic of 3-state autoinhibition model and equation relating observed affinities to the DBD’s ΔG and the autoinhibition constant (*K*_auto_, defined as the ratio of ‘closed’ to ‘open’ unbound TF) (**Supplementary Note 1**). **D** 2D-histograms comparing the calibrated ‘degenerate motif’ library ΔGs for FL *vs.* DBD-only constructs of Msn2, Msn4, and Com2. Sequences were binned by mean DBD and FL ΔG; pixel darkness indicates sequence density per bin. Yellow lines indicate 3-state autoinhibition model fit lines for each DBD-FL construct pair with annotated *K*_auto_ and RMSE values; dotted black lines indicate the 1:1 line. **E** Fitted *K*_auto_ parameters for each DBD + IDR combination *vs.* the ALBATROSS-predicted radius of gyration (R_g_) for each IDR. Marker colors indicate the DBD to which each IDR was attached; marker sizes vary inversely with fit RMSE.

Fitting the resulting concentration-dependent binding curves to a single-site binding model yielded *K*_d_s that were highly reproducible across experiments (*r*^2^ > 0.98 for all constructs) and ranged from < 50 nM to > 30 µM (**Supplementary Fig. 18, 19**; all curves are available in **Supplementary File 1**). To determine upper and lower limits on resolvable affinities, we simulated fitting binding curves under the experimental conditions used here and assessed agreement between ‘ground truth’ and fitted values (**Methods**). While the specificities of DBD and FL constructs remained similar, the FL constructs of Msn2 and Msn4 consistently bound fiducial sequences more weakly, with ∼1.5-3-fold higher *K*_d_s (**Fig. 2B, Supplementary Fig. 18, 20**).

To anchor relative affinities from the larger ‘degenerate motif’ library in absolute space, we then regressed all BET-seq relative affinities (ΔΔGs) against these MITOMI-measured *K*_d_s, yielding ∼6.3 million *K*_d_s (**Fig. 2A**). BET-seq ΔΔG measurements of sequences that bound with ΔΔGs > 3 standard deviations below an estimated Gaussian noise floor (**Methods)** were linearly correlated with MITOMI-measured Κ_d_s (*r*^2^ > 0.8), with MITOMI measurements consistently detecting weaker sequence-specific binding than BET-seq (**Supplementary Fig. 21**). Calibrated ΔGs of the entire ‘degenerate motif’ DNA library confirmed that both Msn2 and Msn4 IDRs uniformly decreased binding affinities by 0.50±0.15 and 0.65±0.08 kcal/mol on average (∼2.3- and 3-fold in *K*_d_ space), respectively, while the presence of the Com2 IDR slightly increased affinities by 0.23±0.13 kcal/mol (**Supplementary Fig. 22**).

An observed decrease in affinities for FL Msn2 and Msn4 is consistent with IDR-mediated autoinhibition. The simplest model that can describe such autoinhibition is a 3-state model in which: (1) a TF can be in either a ‘closed’ (autoinhibited) or ‘open’ state, (2) only the ‘open’ state can bind DNA, and (3) an autoinhibition constant (*K*_auto_) quantifies the ratio of ‘closed’ to ‘open’ TF molecules at equilibrium (**Fig. 2C**). This model yields the following closed-form equation relating the macroscopically observed ΔG of an autoinhibited TF (*ΔG_obs_* to the microscopic ΔG of the DBD for a single DNA binding site (*ΔG_DBD_*) and the autoinhibition constant (*K*_auto_):

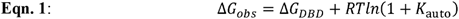

This 3-state autoinhibition model predicts that autoinhibition imposes a constant ΔG penalty that increases with increasing *K*_auto_ values (regardless of the DBD ΔG), consistent with experimental observations (**Fig. 2C, Supplementary Note 1**). Calibrated ΔGs of DBD and FL construct pairs of Msn2 and Msn4 were well-fit by this model (RMSE ≤ 0.15 kcal/mol) with fitted autoinhibition constants of 1.34 and 1.98, suggesting that ∼57% and 66% of free FL Msn2 and Msn4 molecules, respectively, were in the autoinhibited state in our assay (**Fig. 2D**); Com2 did not show signatures of autoinhibition.

### Autoinhibition by IDRs depends on DBD-IDR combination

IDR-mediated autoinhibition of DNA binding by Msn2 and Msn4 IDRs could either: (1) require specific molecular contacts between IDRs and DBDs or (2) result from simple steric blocking that limits DNA access to the DNA-binding interface. If autoinhibition requires specific intra-protein interactions, the strength of autoinhibition by a given IDR should depend on the identity of the attached DBD; conversely, if autoinhibition is due to steric blocking, the strength of autoinhibition should be constant across different DBDs and depend only on the space excluded by each IDR, which can be approximated by the radius of gyration (IDR size) and Flory scaling exponent (IDR density) parameters^47,48^.

To distinguish between these possibilities, we constructed 6 TF chimeras mixing the Msn2, Msn4, and Com2 DBDs and IDRs, quantified both their relative binding energies to the ‘degenerate motif’ library and *K*_d_s to the ‘fiducial’ sequences using BET-seq and MITOMI, respectively, and again derived calibrated ΔGs for the ‘degenerate motif’ library measurements (**Supplementary Figs. 23-26**, binding curves available in **Supplementary File 1**). All 3 IDRs autoinhibited DNA binding by Msn2 and Msn4 DBDs with observed effects well-fit by the proposed 3-state autoinhibition model (**Fig. 2E, Supplementary Fig. 27**), with several observations consistent with inhibition requiring specific IDR/DBD contacts. First, the extent of autoinhibition (fitted *K*_auto_) for each IDR varied across IDR/DBD combinations. Second, the extent of observed autoinhibition was not correlated with either IDR amino acid length or ALBATROSS^49^-predicted radius of gyration: despite no significant differences in predicted scaling exponents (ranging from 0.515-0.526, **Supplementary Fig. 27**), and therefore no differences in predicted IDR density, the smallest IDR (Com2-IDR) drove the strongest apparent autoinhibition of each DBD (*K*_auto_ = 1.67 and 4.47 for Msn2 and Msn4, respectively). Finally, chimeric constructs that paired either the Msn2 or Msn4 IDR with Com2’s DBD were not well-fit by simple autoinhibition (model fit RMSE = 0.31 and 0.37 kcal/mol, respectively); instead, these chimeric constructs showed weak binding to a population of sequences that was not bound by the DBD alone (**Supplementary Fig. 27**). Interpreting AffinityNet models trained on these data suggested that these chimeras preferred G-rich homonucleotide stretches, similar to the FL Com2 construct (**Supplementary Figs. 28, 29**). Together, these results suggest IDR-mediated autoinhibition is not explained by IDR size alone and is consistent with requiring specific DBD-IDR interactions.

### Msn2 and Msn4 IDRs substantially alter in vitro binding preferences to promoter sequences

These ‘degenerate motif’ DNA library experiments probed how differences in TF composition impacted AGGGG motif specificity but did not assess impacts on binding to actual genomic sequences. To test if IDRs alter binding to promoter sequences, we again applied BET-seq to measure relative affinities (ΔΔGs) for DBD-only and full-length Msn2, Msn4, and Com2 constructs binding a ‘promoter library’ of 213,487 sequences tiling across all 5861 *S. cerevisiae* promoters (75 bp tiles offset by 25 bp steps within the 1 kb upstream of each annotated transcriptional start site) (**Fig. 3A**). Measured ‘promoter library’ ΔΔGs were again highly reproducible, with mean replicate Pearson *r*^2^ > 0.85 and mean RMSE < 0.27 kcal/mol (**Supplementary Figs. 30-32**).

**Figure 3.**
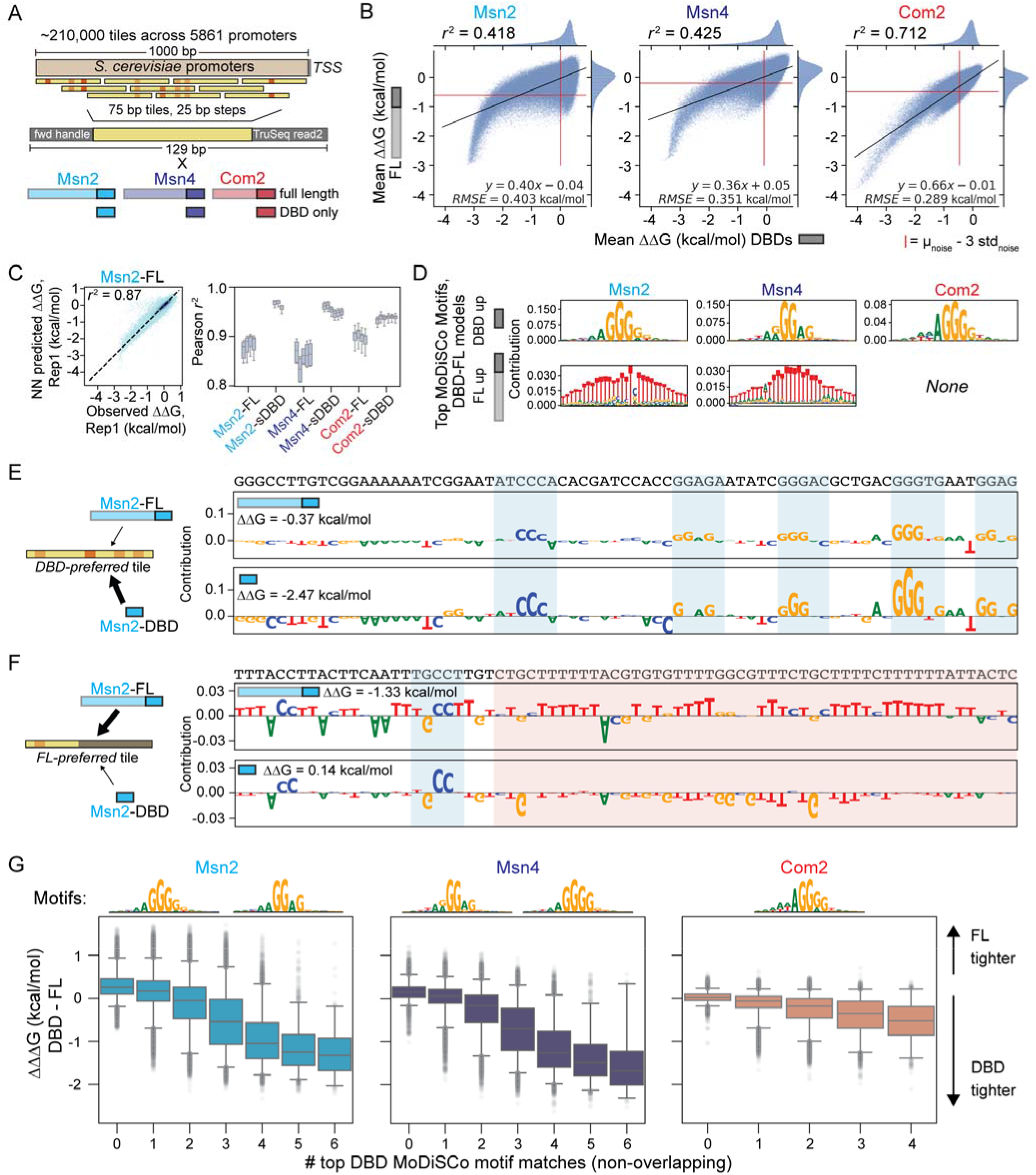
IDRs alter binding to genomic sequences *in vitro*. **A** Schematic of the ‘promoter’ library comprised of 75 bp tiles across all *S. cerevisiae* promoter sequences in 25 bp steps. **B** Scatter plots comparing measured ΔΔGs for FL *vs.* DBD-only Msn2, Msn4, and Com2 constructs binding to the ‘promoter’ library. Markers denote per-sequence ΔΔGs; black lines denote linear fits with annotated fit parameters; red lines denote thresholds used to delineate ‘bound’ sequences (see Supplementary Methods). Pearson *r*^2^ is reported for all sequences considered ‘bound’ by either DBD or FL constructs. **C** Representative scatter plot showing AffinityNet-predicted *vs.* BET-seq-measured ΔΔGs for FL Msn2 binding the ‘promoter’ library (left) and distributions of Pearson *r*^2^ values between predicted and observed ΔΔGs across 5-fold test validations of each model (right). Markers represent individual DNA sequences; pixel darkness indicates sequence density; box plot extents indicate Pearson *r*^2^ ranges. **D** Top MoDISco motifs contributing to DBD- or FL-preferred binding for Msn2, Msn4, and Com2. **E** Per-nucleotide contribution scores for FL and DBD Msn2 constructs binding an example DBD-preferred promoter tile with five separate AGGGG-variant motif hits (found by FiNeMo). **F** Per- nucleotide contribution scores for FL and DBD Msn2 constructs binding an example FL-preferred promoter tile with a homoT motif hit (found by FiNeMo). **G** Box and whisker plots showing distributions of differences in DBD and FL TF ΔΔGs (ΔΔΔGs) for the ‘promoter’ library binned by the number of AGGGG and AGGGG-variants within each promoter tile; box plot whiskers extend to the 2.5 and 95 percentiles of ΔΔΔG distributions, with outliers plotted with gray markers. Specific motif CWMs used for motif finding are shown above each plot.

In contrast to the ‘degenerate motif’ results, the presence of IDRs dramatically altered observed binding energy and specificity landscapes, especially for Msn2 and Msn4 (*r*^2^ = 0.42, 0.42, and 0.71 for Msn2, Msn4, and Com2 DBD vs. FL comparisons respectively, **Fig. 3B, Supplementary Fig. 31**). While many sequences were bound by both constructs, a substantial fraction of the library (34.4%, 19.8%, and 9.8% for Msn2, Msn4, and Com2, respectively) was bound above background levels only in the absence of the IDR; in addition, some sequences were bound only by the FL construct, particularly for Msn2 (**Fig. 3B**).

### DBD-only constructs prefer sequences with multiple motifs

To identify the sequence features driving differential DBD and FL promoter tile binding, we trained AffinityNet models on these BET-seq data, which accurately predicted held-out ΔΔGs during 5-fold cross-validation (Pearson *r*^2^ > 0.8 over 5 independent test validations; **Fig. 3C, Supplementary Fig. 33).** We then interrogated these models using a pipeline in which we: (1) calculated the predicted difference between DBD and FL constructs for each promoter tile, (2) used DeepLIFT/DeepSHAP to estimate the contribution of each nucleotide in each tile to this difference, and then (3) ran TF-MoDISco^50^ on these contribution scores to generate contribution weight matrices (CWMs) of differentially bound sequences. While all constructs strongly bound expected AGGGG motifs, the FL constructs of both Msn2 and Msn4 showed a *relative* preference for long, T-rich homonucleotide sequences (denoted as “FL-up”, **Fig. 3D**; **Supplementary Fig. 34**). Examining contribution scores across entire tiles revealed that Msn2 DBD-preferred promoter tiles were enriched for clusters of predictive AGGGG motif variants bearing 1-2 mutations (**Fig. 3E, Supplementary Fig. 35**) while Msn2 FL-preferred promoter tiles were enriched for predictive low-complexity homoT stretches (**Fig. 3F, Supplementary Fig. 36**).

As an orthogonal test of whether DBD-preferred promoter tiles generally contained multiple AGGGG motif variants, we scanned each promoter tile sequence for instances of predictive monomer motifs (using FiNeMo) and quantified the distributions of relative binding energy differences as a function of the number of motifs per tile. The magnitude of the relative DBD preference for a given promoter increased with the number of motif variants, particularly for Msn2 and Msn4 (**Fig. 3G**). Put another way, IDR-mediated autoinhibition of Msn2 and Msn4 appeared to increase for promoter tiles bearing clusters of motifs.

### DBD-driven preferences for multiple motifs reflect increased affinity from avid binding

The ‘promoter’ library is relatively long (129 bp *vs.* 89 bp for the ‘degenerate motif’ library), with many tiles containing multiple motifs in close proximity (> 50% of tiles contain > 2 Msn2 DBD-preferred motifs < 10 nm away from one another; **Fig. 4A**). As MITOMI and BET-seq experiments assessed binding of solution-phase DNA to surface-immobilized TFs spaced ∼9 nm center-to-center (**Supplementary Figs. 37-38**), a preference for binding multiple motifs could suggest that multiple surface-immobilized TFs are simultaneously binding the same DNA molecule (termed ‘avid’ binding) (**Fig. 4B, Supplementary Note 1**). In this case, two TFs can ‘bridge’ a single molecule with dramatically enhanced affinity (**Eqn. 2**):

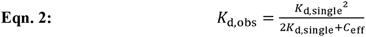

**Figure 4.**
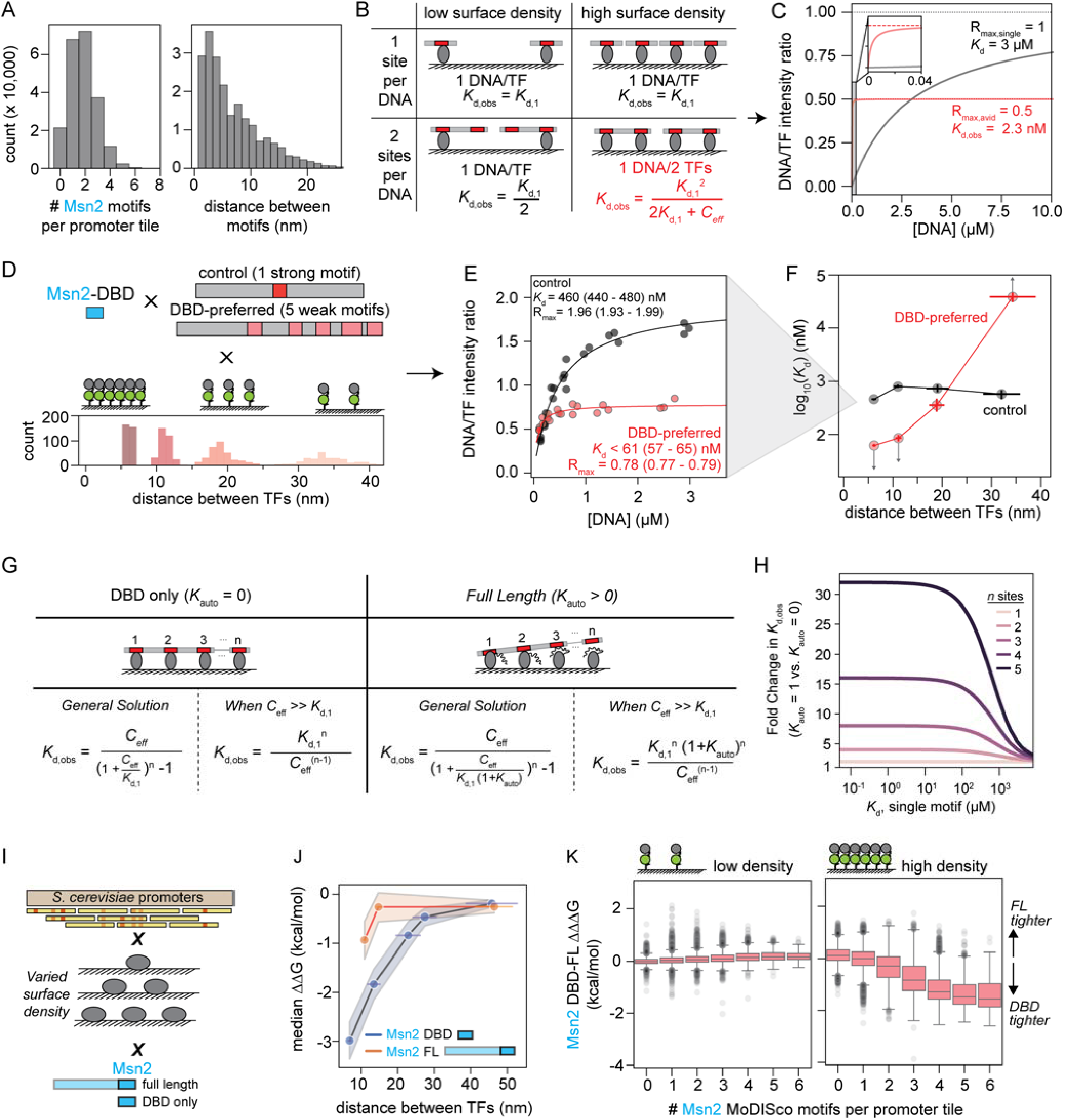
Autoinhibition by Msn2’s IDR can preferentially impact binding to sequences containing multiple binding sites. **A** Histograms showing the distributions of the number of AGGGG and AGGGG-variant motif-matches per promoter tile in the Msn2 DBD model (left) and distances between adjacent motifs (right). **B** Schematic of binding geometries and impacts on observed *K*_d_ for DNA sequences containing 1 (top) or 2 (bottom) binding motifs interacting with TFs patterned at low (left) or high (right) density (**Supplementary Note 1**). **C** Predicted concentration-dependent binding for TFs patterned at high-density interacting with DNA containing one (red) or two (blue) binding sites with a *K*_d_ of 3 µM per binding site. **D** Schematic of a MITOMI experiment measuring Msn2 DBD binding to a control sequence with one AGGGG motif and an Msn2 DBD-preferred promoter tile with 5 AGGGG-variants (from Fig. 3E) as a function of patterned TF density (top); histograms show estimated average distances between TF molecules at each density (bottom). **E** Measured concentration-dependent binding of Msn2 DBD binding to control sequence (black) and DBD-preferred promoter tile (red). Markers indicate data from single chambers; lines indicate Langmuir isotherm fits with annotated fit parameters. **F** Measured log-transformed *K*_d_s for Msn2 DBD binding the control sequence (black) or the DBD-preferred promoter tile with 5 motif variants (red) as a function of average distance between TF molecules. Markers represent *K*_d_ fits for indicated sequences and TF center-to-center distances; y errors denote interquartile ranges of *K*_d_ fit distributions; x errors denote standard deviations of TF center-to-center distances across chambers. **G** Schematic illustrating *n* adjacent immobilized TFs to a DNA molecule with *n* equivalent binding sites with (right) and without (left) autoinhibition on observed equilibrium *K*_d_ and derived equations (**Supplementary Note 1**). **H** Simulated fold difference in observed *K*_d_ between autoinhibited (*K*_auto_ = 1) and uninhibited (*K*_auto_ = 0) TF constructs to DNA with *n* identical binding sites as a function of the *K*_d_ of individual binding sites given an estimated *C*_eff_ = 3.66 mM (**Methods**); line color indicates number of sites. **I** Schematic of BET-seq measurements of Msn2 DBD and FL constructs binding the ‘promoter’ library as a function of TF surface density. **J** Median ΔΔG of Msn2 DBD (blue) and Msn2 FL (orange) binding to the top 1000 Msn2-DBD preferred promoter tiles as a function of estimated distance between TF molecules. X-axis error bars indicate standard deviations of distance distributions; y-axis error indicates standard deviation of ΔΔG distributions. **K** Distributions of differences between Msn2 DBD and FL construct ΔΔGs for promoter tiles as a function of identified contributing motifs as a function of the TF patterning density.

Here, *C*_eff_ represents the effective concentration of the second DNA binding site once the first site has been bound. For 2 sites with individual *K*_d_s of 3 µM, a ∼9 nm center-to-center distance between TFs yields an estimated *C*_eff_ of ∼4 mM and a substantially tighter observed binding affinity of 2.45 nM (**Fig. 4C, Methods**); the expected number of DNA molecules bound per surface-immobilized TF at saturating ligand concentrations should also decrease (**Fig. 4C**).

To test for ‘avid’ binding, we used MITOMI to quantify binding of Msn2’s DBD to: (1) a ‘control’ sequence containing a single AGGGG motif, or (2) the DBD-preferred promoter tile shown in **Fig. 3E** containing 5 weak motif variants, each with 2 or more nucleotides mutated relative to the AGGGG consensus (**Fig. 4D**). Consistent with ‘avid’ binding, Msn2’s DBD bound the DBD-preferred tile significantly more tightly than the ‘control’ sequence (*K*_d_ < 61 (IQR 57-65) nM vs. *K*_d_ = 462 (IQR 440-480) nM respectively; the first value is an upper limit because sites were almost fully occupied at the lowest DNA concentrations tested) and saturated at a lower fractional occupancy (maximum ratio of DNA/TF fluorescence intensities, R_max_ = 0.78 *vs*. 1.96 for the multi-site and ‘control’ sequences, respectively; **Fig. 4E**). Additional MITOMI experiments quantifying binding at different TF surface densities also revealed that, while apparent binding affinities for the ‘control’ sequence did not dramatically change as a function of surface density, apparent affinities for the multi-site sequence decreased as the average distance between TFs increased (from *K*_d_ < 61 (IQR 57-65) nM at a ∼6 nm distance to 350 (IQR 260-470) nM at a ∼19 nm distance and >39 (IQR 32-49) µM at a ∼34 nm distance; **Fig. 4F, Supplementary Fig. 39**). This dependence of high affinity binding on dense TF surface patterning confirms that ‘avid’ binding of adjacent TF molecules to single DNA molecules with multiple weak binding sites drives strong observed binding in our system. While the avid binding observed here is an artifact of the surface-immobilization geometry used in BET-seq experiments, this model may also be relevant for describing binding by complexes comprised of multiple weakly-interacting TFs and co-activators in cells (**Discussion**).

### The impacts of autoinhibition on avid binding depend on the number of binding sites

Results thus far establish that: (1) IDRs can autoinhibit DNA binding and (2) multiple surface-immobilized TFs can simultaneously bind a single DNA molecule at high densities, driving density-dependent differences in observed affinities. To understand how these distinct mechanisms alter concentration-dependent binding, we extended the autoinhibition model to predict observed binding affinities for *n* adjacent surface-immobilized TFs interacting with a DNA molecule with *n* binding sites of equal dissociation constants (*K*_d,1_) (**Fig. 4G, Supplementary Note 1**). Under the simplifying assumption that *C*_eff_ *>> K*_d,1_ (**Supplementary Fig. 40)**, this yields a closed-form solution for the observed *K*_d_ (**Eqn. 3**):

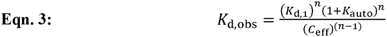

where *K*_d,obs_, represents the observed *K*_d_, *K*_d,1_ is the affinity of each individual site, *K*_auto_ is the autoinhibition constant, *C*_eff_ is the effective concentration, and *n* is the number of binding sites.

This model suggests a novel mechanism by which uniform autoinhibition could differentially impact binding to promoters containing different numbers of sites: for example, an uninhibited construct (*K*_auto_ = 0) would bind DNA containing 5 binding sites 32-fold more tightly than a modestly autoinhibited FL construct (*K*_auto_ = 1, or 50% inhibited), even for relatively weak (∼10 µM) binding sites **(Fig. 4H).** These exponentially increasing differences in *K*_d_ as a function of the number of binding sites appear as linear differences in ΔΔG space, consistent with our observations of Msn2’s DBD and FL constructs (**Fig. 3G**).

To test if autoinhibition combined with avid binding can explain the observed Msn2-DBD preferences for promoter tiles containing multiple motifs, we repeated BET-seq experiments measuring binding of Msn2 FL and DBD to the ‘promoter’ library while varying TF surface density (**Fig. 4I**). Consistent with avid binding, preferential Msn2 DBD binding to the top 1000 DBD-preferred promoter tiles decreased as the space between adjacent TFs increased (**Fig. 4J**). Further, the relative preferences of Msn2’s DBD for promoter tiles as a function of the number of motifs disappeared at low density (**Fig. 4K**).

### Msn2’s IDR can directly bind DNA in an avidity-dependent manner

Prior results established that Msn2 and Msn4 FL constructs preferentially bind sequences containing homoT stretches relative to their DBD-only counterparts, suggesting that their IDRs could independently bind DNA. To test this, we used MITOMI to measure binding of IDR-only constructs of Msn2, Msn4, and Com2 to 13 promoter tiles spanning the observed binding distributions **(Fig. 5A).** While Msn4 and Com2’s IDRs showed no observable binding to any promoter tiles, Msn2’s IDR reproducibly bound multiple sequences (replicate *r*^2^ = 0.815) with a wide range of *K*_d_s (0.8 µM to > 51.1 µM) (**Supplementary Figs. 41-42**). Sequences containing homoT motifs were among the tightest bound (**Fig. 5B, C**, **Supplementary Fig. 41**); all curves available in **Supplementary File 2**).

**Figure 5.**
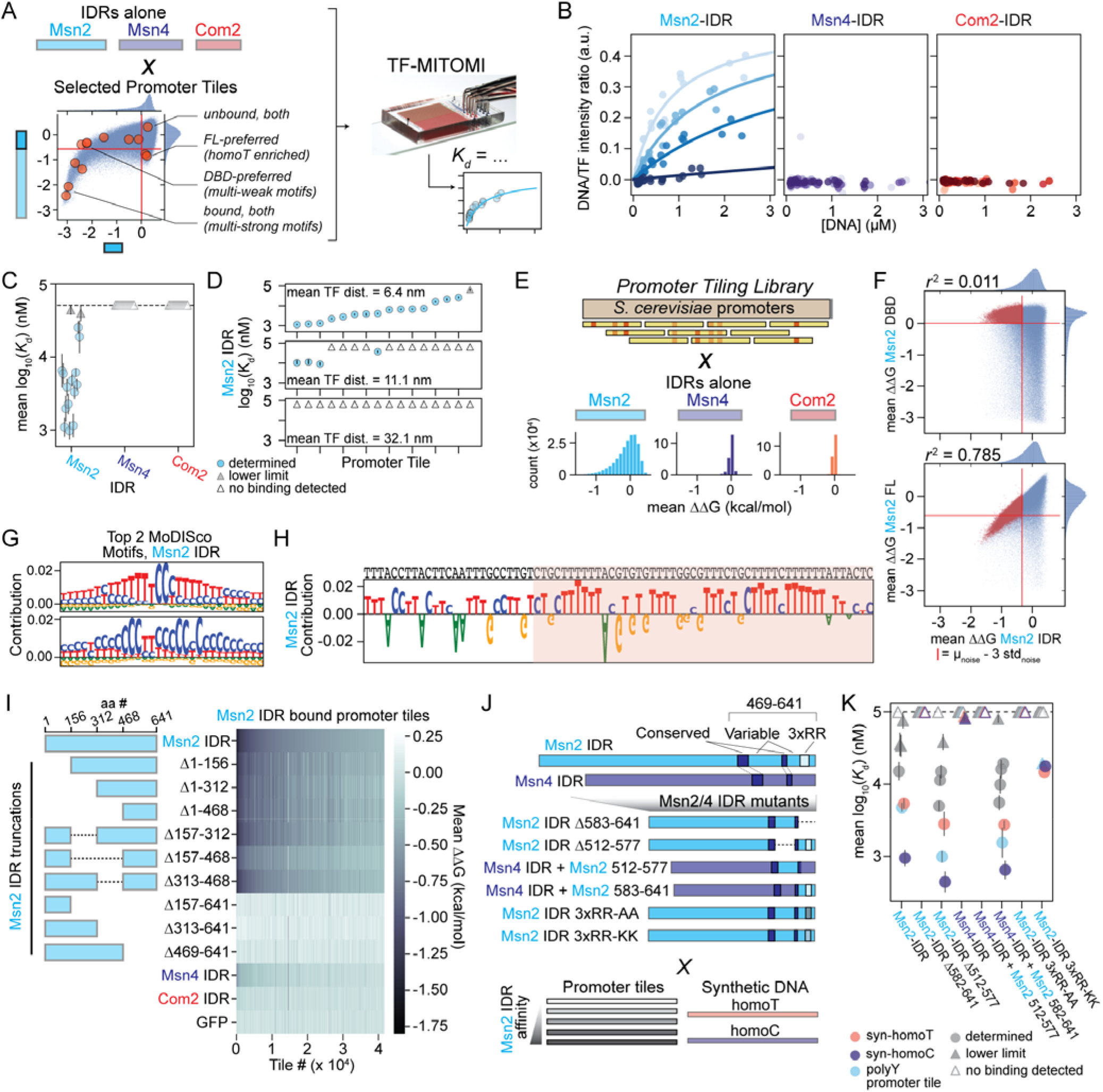
Msn2-IDR directly binds homopolymeric DNA through interactions mediated by conserved C-terminal di-arginine pairs. **A** Schematic showing TF-MITOMI measurements of Msn2, Msn4, and Com2 IDR-only constructs binding selected sequences from the ‘promoter’ library. **B** Representative concentration-dependent binding curves for Msn2, Msn4, and Com2 IDRs binding to four selected promoter tiles (sequences and individual curves shown in **Supplementary Fig. 41, Supplementary File 2**). Markers indicate fractional binding in the form of DNA fluorescence (Alexafluor 647) normalized to TF fluorescence (meGFP) at given total DNA concentrations; solid lines denote global Langmuir isotherm fits. **C** Mean log-transformed fitted *K*_d_ values for Msn2, Msn4, and Com2 IDRs across selected promoter tiles; error bars denote the difference between the mean *K*_d_ and *K*_d_ fits of two independent device replicates; blue circle markers indicate defined *K*_d_ fits, gray triangle markers indicate lower limit *K*_d_ estimates, and white triangles represent interactions for which binding was not detected. **D** Mean log-transformed fitted *K*_d_ values for Msn2 IDR binding across selected promoter tiles across three average distances between TF molecules; markers indicate the same categories as in **C**; y error represents the interquartile range of *K*_d_ fits over 400 subsamplings of the data within a single experiment (sequences and individual curves shown in **Supplementary Fig. 43**, **Supplementary File 3**). **E** Schematic of BET-seq measurements for Msn2, Msn4, and Com2 IDRs binding to the ‘promoter’ library (top) and histograms of measured mean ΔΔG distributions for each construct across two device replicates (bottom). **F** Scatter plots showing mean ΔΔGs for Msn2 DBD or FL *vs.* Msn2-IDR. Red lines indicate bound/unbound thresholds of 3 standard deviations of the noise distribution away from the mean. Red markers denote sequences preferentially bound by Msn2’s IDR relative to its DBD with annotated Pearson *r*^2^ values; blue markers denote all other sequences. **G** Top two MoDISco motif CWMs for AffinityNet models trained on Msn2 IDR ‘promoter’ library BET-seq data. **H** Per-nucleotide contribution scores predicted by AffinityNet for Msn2 IDR binding to a Msn2 FL-preferred promoter tile (from Fig. 3F); red shading highlights homoT motif match. **I** Schematic showing Msn2 IDR truncation constructs (left) and heatmap of mean BET-seq ΔΔGs (n = 2) for these truncations, Msn4’s IDR, Com2’s IDR, and an meGFP-only construct binding to ‘promoter’ library tiles bound by Msn2’s IDR. **J** Schematic showing Msn2 and Msn4 IDR constructs with mutations within non-conserved sections of the Msn2 469-641 region (top) and sequences measured by TF-MITOMI (bottom). **K** Log-transformed *K*_d_ values for Msn2 and Msn4 IDR C-terminal variable region mutants. Markers denote mean across two experimental replicates; error bars represent the absolute error between the mean log_10_(*K*_d_) and each replicate; circle markers indicate defined *K*_d_ fits, filled-in triangle markers indicate lower limit *K*_d_ estimates, and hollow triangles represent interactions for which binding was not detected (sequences and individual curves shown in **Supplementary Fig. 50-52, Supplementary File 4**).

For all binding curves, saturating occupancies (DNA/TF intensity ratios) were lower than for DBD-containing constructs (**Figs. 5B, Supplementary Fig. 41**), suggesting that multiple IDRs were interacting with single DNA molecules via avid binding. To test this, we varied the Msn2 IDR patterning density and measured binding to the same set of promoter tiles. Msn2’s IDR-DNA binding was highly sensitive to changes in patterning density, with a ∼2-fold change in TF spacing (from ∼6 to ∼11 nm) causing ∼10-fold decreases in affinity for the tightest binding sequences and a ∼5-fold change in TF spacing (at ∼32 nm) rendering binding undetectable (**Fig. 5D, Supplementary Fig. 43, Supplementary File 3**). These results confirm that detectable DNA binding requires engagement of multiple IDRs and that individual IDR-DNA binding interactions are therefore very weak.

### Msn2’s IDR binds homoT and homoC DNA sequences

To comprehensively map the sequence preferences of Msn2’s IDR and confirm that Msn4 and Com2’s IDRs do not bind DNA, we used BET-seq to measure binding of the three IDR-only constructs to the entire promoter library (**Fig. 5E**). Only Msn2’s IDR showed reproducible DNA binding over a wide dynamic range (Msn2-IDR replicate *r*^2^ > 0.71 with ΔΔGs < 2 kcal/mol); one of two Msn4-IDR replicates showed slight non-reproducible binding (ΔΔGs < 1 kcal/mol and *r*^2^ < 0.03 between replicates; **Supplementary Figs. 44, 45**). Comparisons of promoter tile binding between FL Msn2, the IDR-only construct, and the DBD-construct revealed that many sequences were similarly bound between the FL and IDR constructs but not the DBD, consistent with independent IDR binding (Msn2 FL *vs*. Msn2 IDR r^2^ = 0.786 for sequences bound by Msn2’s IDR and not its DBD) (**Fig. 5F**).

Training AffinityNet models on these Msn2 IDR promoter binding data (train vs. test *r*^2^ > 0.82, **Supplementary Fig. 46**) and using TF-MoDISco to extract CWM representations of sequences contributing to binding affinity **(Supplementary Fig. 47**) revealed a strong preference for homoT nucleotide stretches similar to those seen in FL-preferred sequences (**Figs. 5G, 3D**). These same preferences were also visible when considering contribution scores across individual promoter sequences bound by both Msn2 FL and IDR (**Figs. 5H, 3F**). The second-highest contributing motif consisted of homoC stretches interspersed with T nucleotides **(Supplementary Fig. 47**). These low complexity sequences resembled consecutive CCCCT motifs (the reverse complement of Msn2 DBD consensus motif, AGGGG), often overlapped with ‘motif’ hits of CCCCT variants (83.5%), and were rarely seen in promoter tiles lacking multiple ‘motifs’ (3588 hits with 0 or 1 ‘motif’ matches, 8924 with ≥ 2; **Supplementary Fig. 48**). Together, these results suggest that the Msn2 IDR independently binds homoT stretches while both the IDR and DBD bind these homoC stretches.

### Msn2’s IDR-DNA binding depends on positively charged di-arginine residues proximal to Msn2’s DBD

Previous studies showed that the *in vivo* effects of Msn2’s IDR on genomic localization depended on sequence features distributed across the entirety of the IDR^30–32^. To identify the residues required for *in vitro* DNA binding by Msn2’s IDR, we generated 9 Msn2 IDR deletion constructs (Δ1-156, Δ1-312, Δ1-468, Δ469-641, Δ313-641, Δ157-641, Δ157-312, Δ157-468, and Δ313-468) and quantified binding of each to the ‘promoter’ library via BET-seq (**Fig. 5I**). All constructs containing amino acids 469-641 showed equal binding and all constructs lacking these residues showed no binding, establishing that this region is both necessary and sufficient for *in vitro* DNA binding.

Msn4’s IDR does not reproducibly bind DNA but Msn2’s IDR does, suggesting that the amino acid features of Msn2 469-641 responsible for DNA binding are not conserved in Msn4. To identify the specific amino acids within the 469-641 region of Msn2’s IDR likely to bind DNA, we constructed an alignment of Msn2 and Msn4 ortholog protein sequences from *S. cerevisiae* and the related yeast species *K. lactis*, *N. glabratus*, *S. eubayanus*, *S. kudriavzevii*, and *S. mikatae* (**Supplementary Fig. 49**). Two regions within the 469-641 fragment were relatively well-conserved across Msn2 and Msn4 orthologs (485-511 and 578-582), facilitating alignment; the remaining sequence regions varied substantially (512-577 and 583-641 of *S. cer* Msn2). The 583-641 variable region contained three di-arginine (RR) pairs present only in Msn2 orthologs, which we hypothesized may mediate IDR-DNA binding since, in other DNA binding proteins, (1) arginines (and not lysines) recognize enhanced electrostatic potentials of narrow minor grooves created by A/T repeat sequences^51^ and (2) arginine-rich patches proximal to DBDs can bind RNA^52^.

To test if these residues mediate DNA binding, we generated 6 new constructs: (1) Msn2 IDR constructs with deletions that eliminated (Δ583-641) or preserved (Δ512-577) these RR pairs, (2) Msn4/Msn2 IDR chimeric constructs that did (583-641) or did not (512-577) introduce these RR pairs, and (3) Msn2 IDR constructs in which the RR pairs were mutated to AA or KK pairs, respectively (**Fig. 5J**). We then measured binding of these constructs to a small set of promoter tiles previously shown to bind Msn2’s IDR as well as synthetic homoT and homoC sequences using MITOMI (**Supplementary Fig. 50-52**). Only constructs containing intact RR pairs showed detectable binding and insertion of the RR pairs into Msn4’s IDR resulted in binding similar to Msn2’s IDR, establishing these residues as both necessary and sufficient for direct DNA binding (**Fig. 5K, Supplementary Fig. 50-52,** all curves available in **Supplementary File 4**). Strikingly, even mutation to KK severely reduced binding, suggesting that both positive charge and properties unique to arginine are essential for IDR-DNA binding. Overall, these results establish that the Msn2 IDR is capable of directly and sequence-specifically recognizing DNA under avid conditions. While individual IDR/DNA interactions are likely too weak to independently bind DNA, this property could drive either *inter*molecular binding between IDRs present within TF clusters or *intra*molecular avid binding between the DBD and IDR to drive distinct promoter localization between DBD-only and FL constructs.

### In vivo models recover IDR-associated homoT preferences

Results thus far establish that the Msn2 IDR mediates DNA binding via autoinhibition and direct (avidity-mediated) recognition of low-complexity C and T-rich sequences. To assess whether these interactions impact genomic localization in cells, we trained BPNet^38^ neural network models on two publicly available, independent *in vivo* Msn2 ChEC-seq binding datasets that included binding data for either FL, DBD-only, and IDR-only Msn2 constructs (“B”: Brodsky et al, 2020)^30^ or FL Msn2 under varied stress conditions (“M”: Mahendrawada et al, 2025)^53^ (**Fig. 6A**). While ChEC-seq datasets were generally reproducible between replicates (Pearson *r*^2^ > 0.54 and 0.98 for “B” and “M”, respectively; **Supplementary Fig. 53**), BPNet models better predicted the counts within peak regions of “M” samples compared to “B” samples (Pearson *r*^2^ = 0.45 and 0.29 > *r*^2^ > 0.19 for “M” and “B”, respectively, **Fig. 6B, Supplementary Fig. 54**).

**Figure 6.**
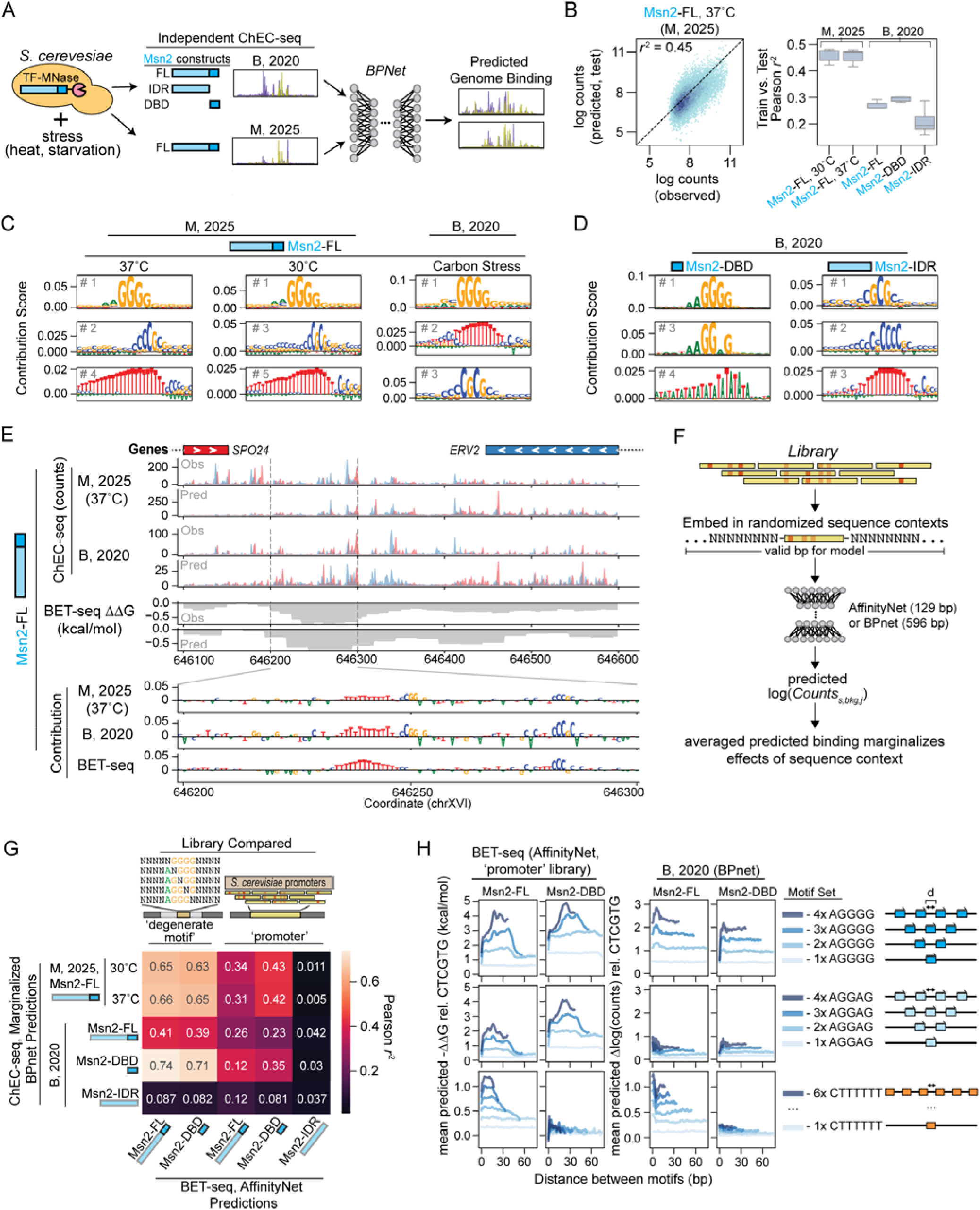
Neural network models suggest Msn2’s IDR also binds homonucleotide sequences in cells. **A** Schematic showing BPNet analysis pipeline for FL Msn2 DBD and IDR-only ChEC-seq binding data from two independent studies (Brodsky et al, 2020 and Mahendrawada et al, 2025). **B** Left: 2D histogram showing BPNet-predicted *vs.* ChEC-seq-measured log counts across peak regions; marker color denotes density of sequences per bin with annotated Pearson *r*^2^ values. Right: box and whisker plot of Pearson *r*^2^ value distributions across 5-fold test set validations for each experiment. **C** Top TF-MoDISco motifs for FL Msn2 models from each dataset. **D** Top MoDISco motifs for Msn2 DBD and IDR-only constructs. **E** Example genome browser tracks showing observed and predicted (BPNet and AffinityNet) *in vivo* and *in vitro* binding of full-length Msn2 to the ERV2 promoter region, which contains a homoT motif. Red and blue ChEC-seq traces denote + and - strand read counts, respectively. **F** Schematic of marginalization analysis for comparing *in vivo* and *in vitro* model predictions. **G** Heatmap showing Pearson *r*^2^ between AffinityNet-predicted *in vitro* BET-seq -ΔΔG values *vs*. marginalized BPNet-predicted ChEC-seq log counts for Msn2 DBD, IDR, and FL constructs binding to promoter tiles. **H** Binding predicted by *in vitro* AffinityNet (left) and *in vivo* BPNet (right) models (-ΔΔG and Δ log(counts), respectively) for Msn2 FL and DBD constructs interacting with synthetic sequences containing systematically varying numbers of consensus and homoT motifs as a function of distance between motifs.

As expected, the consensus AGGGG Msn2 binding motif was the top contributing motif for all Msn2 FL and Msn2 DBD models (**Fig. 6C, D**). Consistent with our discovery of independent Msn2 IDR binding of homoT DNA *in vitro*, models trained on Msn2 FL and IDR-only localization – but not Msn2 DBD localization – revealed preferences for homoT motifs of variable length, often capped with 2-4 C nucleotides (**Fig. 6C, D, Supplementary Fig. 55, 56**); in addition, many regions containing isolated homoT motifs were both predicted and observed to be bound by Msn2 FL and IDR-only constructs but not by Msn2’s DBD (*i.e.* the region downstream of the *ERV2* gene; **Fig. 6E, Supplementary Fig. 57**). Consistent with the existence of binding determinants in cells not recapitulated *in vitro*, the FL and IDR-only Msn2 models from both studies also predicted binding to CG dinucleotide repeats surrounded by C-rich stretches that were not bound in *in vitro* experiments (**Fig. 6C, D, Supplementary Fig. 55, 56**).

### Msn2 FL and DBD-only constructs display similar motif preferences in vitro and in cells

Differences between TF occupancies observed *in cellulo* and *in vitro* can reflect the impacts of cell-specific factors on binding (*i.e*. binding of competing factors, occlusion by nucleosomes, protein-protein binding etc.). Neural network models trained on *in vivo* data can marginalize out signal from surrounding long-range sequence contexts, thereby allowing for a fairer comparison^54^. To accomplish this, we embedded each BET-seq library sequence (89 or 129 bp) in 20 randomized sequence contexts (596 bp total) and compared averaged predicted binding for these sequences between BPNet models trained on *in vivo* data and AffinityNet models trained on *in vitro* data (**Fig. 6F**).

For the ‘degenerate motif’ library, both *in vivo* and *in vitro* models predicted highly similar binding for Msn2 FL and DBD constructs (*r*^2^ = 0.63 to 0.66 with “M” datasets, *r*^2^ = 0.39 to 0.74 with “B” datasets; **Fig. 6G, Supplementary Fig. 58**), with identical preferences for an AGGGG-like consensus motif; *in vivo* models displayed a small but consistent preference for the CGGGG motif variant with CG-rich upstream sequences relative to *in vitro* models (**Supplementary Fig. 59**). By contrast, binding predictions were poorly correlated for IDR-only Msn2 models (*r*^2^ = 0.082 to 0.087) (**Fig. 6G, Supplementary Fig. 58**).

### Models identify three classes of sequence features

For the ‘promoter’ library, predictions were substantially less well correlated, with the highest correlation observed for DBD-only Msn2 constructs (Pearson *r*^2^ = 0.23 to 0.43 and 0.26 to 0.39 for DBD and FL Msn2 respectively, **Fig. 6G, Supplementary Fig. 58**). Models trained on *in vitro* BET-seq data for Msn2’s IDR yielded predictions that were very poorly correlated with those from FL or IDR-only *in vivo* models (Pearson *r^2^* = 0.005 to 0.042, **Fig. 6G**), suggesting factors other than direct IDR-DNA binding (*e.g.*, IDR-protein or IDR-RNA interactions^52^) likely drive sequence-specific IDR-only localization in cells. Consistent with consensus motif-like sequences playing a central role *in vitro* and in cells, predicted promoter library binding for both *in vitro* and *in vivo* DBD-only and FL constructs increased substantially with increasing numbers of consensus-like motifs (**Supplementary Fig. 60**). *In vivo* models alone predicted enhanced binding of IDR-containing constructs to promoter sequences containing multiple CG-rich motifs, suggesting these motifs likely impact binding via cell-specific factors (*i.e.* IDR-driven oligomerization with other DNA-binding proteins; **Supplementary Fig. 60**). Finally, *in vitro* and *in vivo* FL Msn2 models each predicted slightly enhanced binding for sequences containing ‘homoT’ stretches (**Supplementary Fig. 60**), consistent with direct weak binding of ‘homoT’ stretches by the Msn2 IDR *in vitro*.

### Predicted binding to synthetic libraries suggests intermolecular avidity in cells

In *vitro* measurements revealed 2 types of *inter*molecular avidity for densely patterned TF constructs: (1) enhanced binding to clusters of AGGGG-like motifs by DBD-containing constructs, and (2) detectable binding of homonucleotide stretches by IDR-only constructs. If either type of *inter-*molecular avidity also plays a role in cells, binding predicted by *in vivo* models should be enhanced by the presence of multiple motifs in close proximity. To test this, we used the AffinityNet (*in vitro*) and BPNet (*in vivo*) models to predict binding to libraries of synthetic DNA sequences containing different numbers of sequence features at various equidistant spacings (**Fig. 6H**, **Supplementary Fig. 61**). Consistent with observed avid binding *in vitro*, *in vitro* models predicted increased binding of both FL and DBD-only Msn2 to closely clustered consensus-like motifs (**Fig. 6H**, **Supplementary Fig. 61**). While *in vivo* models also predicted increased binding to multiple AGGGG-like motifs, this binding did not depend strongly on motif spacing, consistent with independent, non-avid binding. Both *in vitro* and *in vivo* models predicted increased binding to multiple homoT stretches that was enhanced when these stretches were closely spaced (<30 bp, or 10 nm). These predictions, in combination with our observation that IDR-DNA binding requires *inter*molecular avidity *in vitro*, suggest the presence of avidity-driven IDR-DNA binding *in vivo*.

## Discussion

TF IDRs have been proposed to affect DNA binding through multiple distinct mechanisms, but the relative importance of these mechanisms for Msn2, Msn4, and Com2 genome localization has remained unclear. Here, we establish that Msn2 and Msn4 IDRs (but not the Com2 IDR) autoinhibit DNA binding by their cognate DBDs. In addition, we find that Msn2’s IDR (but not the Msn4 or Com2 IDRs) binds low-complexity homopolymeric T and C DNA sequence-specifically, yet very weakly. This study confirms prior reports that Msn2’s IDR influences genomic localization via direct binding of DNA^34^ but further establishes that this binding is sequence-specific, maps the sequence-specificity landscape, and establishes that DBD-proximal di-arginine repeats are both necessary and sufficient for direct DNA binding. While this IDR binding is likely too weak to be physiologically relevant in isolation, intermolecular avidity drives altered localization *in vitro* and may influence localization *in cells*.

Our observations agree with some previously published observations but not others. Consistent with prior reports that Msn2 constructs lacking the IDR localized to different promoters in cells, we observed different binding energy landscapes for Msn2 FL and DBD-only constructs interacting with promoter-derived sequences. However, prior systematic truncations of or sequential deletions of aliphatic and aromatic residues within the IDR in cells (residues 1-567) progressively decreased differences in genomic localization, while mutating positively charged residues had no effect^30,31^, leading the authors to propose that hydrophobic residues distributed throughout Msn2’s IDR drive IDR-DNA binding. By contrast, we find that 3 pairs of arginine residues unique to Msn2 orthologs just N-terminal to the DBD (residues 617-631) are both necessary and sufficient for direct binding of Msn2’s IDR to DNA *in vitro* (**Fig. 5I-K**). These residues are just outside of the region mutated in the cell-based screen, potentially explaining why this screen saw no impact of mutating positively charged residues. Models trained on *in vivo* datasets also revealed preferences for sequences not bound *in vitro*, consistent with cellular localization via protein-protein interactions and potentially explaining observed effects of truncations.

Prior *in vitro* single-molecule optical tweezers experiments suggested that Msn2’s IDR promotes association with DNA and one-dimensional scanning via weak, charge-mediated interactions^34^. While these observations are broadly consistent with our finding that di-arginine repeats within Msn2 IDR bind DNA, this study suggested that Msn2 IDR/DNA binding is relatively strong (∼40 nM *K*_d_), at odds with our observations of weak binding detectable only under avid binding conditions. We propose the strong binding observed in these optical tweezers experiments likely reflects avid binding by larger complexes of homotypic IDR assemblies, supported by observations of multimeric complexes in electrophoretic mobility shift assays and DNA unzipping traces that show multiple Msn2 unbinding events along a single molecule^34^. However, the immobilization of TFs in our assays precludes observation of sliding during target search, complicating direct comparisons.

The observation that deleting (or even conservatively mutating) the three conserved DBD-proximal di-arginine pairs in Msn2’s IDR ablates DNA binding suggests that this DNA recognition requires properties unique to arginine. Arginine residues within DBDs bind more strongly than lysines to narrow minor grooves of DNA associated with homoT-rich sequences^51^. Though this potentially explains Msn2’s IDR binding of homoT-rich sequences, it does not explain observed binding to homoC- and polypyrimidine-rich sequences, suggesting that disordered arginine pairs may confer specificity through mechanisms distinct from those described in DBDs. The cluster of DBD-proximal arginines within Msn2’s IDR appears similar to DBD-proximal arginine-rich patches that mediate RNA binding by TFs^52,55,56^ and recent live-cell single-molecule imaging studies of SP1 found that DBD-adjacent basic residues in its IDR are required for proper chromatin association^28^, suggesting these sequences may be involved in nucleic acid binding more generally.

The direct measurements of absolute binding affinity presented here also provide novel evidence that Msn2 and Msn4 IDRs autoinhibit binding, likely via specific IDR-DBD interactions, with different impacts on binding affinity to promoter sequences containing isolated or clustered motifs. Even modest autoinhibition per monomer (*e.g. K*_auto_ = 1, corresponding to 50% inhibition) can yield a compounding (1 + *K*_auto_)^n^-fold reduction in observed affinity when bound with avidity to n sites (**Fig. 4H**). Thus, autoinhibition need not simply uniformly reduce affinity; instead, it can act as a specificity filter that preferentially suppresses avid binding to clustered sites while leaving strong single-motif interactions relatively intact. Further, subtle differences in autoinhibition resulting from post-translational modification of IDR sequences could substantially alter the binding landscape across promoter sequences.

While solution-phase biochemistry is considered the ‘gold standard’ benchmark for quantifying biomolecular interactions, measuring binding to surface-immobilized molecules displayed at carefully titrated densities can facilitate measurement of extremely weak binding undetectable in solution-phase assays and dissect how multivalency and avid binding nonlinearly tune apparent affinities. While surface-immobilization is inherent to our experimental platform, we note that analogous conditions may arise in cells. Activation domains within TF IDRs interact with coactivators and other TF IDRs, concentrating TFs *in vivo* at particular genomic loci in local hubs and phase-separated condensates^28,33,57–59^. Recent measurements of direct interactions between over 200 human activation domains and coactivators such as P300 revealed extensive multivalency in which TF activation domains can bind multiple domains within a single coactivator molecule^60^. Coactivators could therefore act as a scaffold, bridging TF molecules and creating the high local TF densities required for avid binding of DBDs to clustered motifs and IDRs to homopolymeric DNA without requiring formation of phase-separated condensates. The widespread prevalence of homotypic clusters of TF binding sites within eukaryotic promoters and enhancers^53,61^ suggests a functional role for avidity; the exponential sensitivity of avid binding to autoinhibition further suggests a powerful mechanism for tuning occupancy at these clustered elements.

These interactions *could* occur in cells – but do they? To address this question, we explicitly compared binding predictions from neural network models trained on *in vitro* and *in vivo* Msn2 binding. Many aspects of binding observed *in vitro* were also observed in cells: (1) DBD-dependent strong binding to consensus and consensus-like motifs, (2) IDR-dependent binding to T-rich DNA stretches, and (3) distance- and multiplicity-dependent enhancement of binding to clusters of these elements. Given the avidity-dependence of IDR-homonucleotide binding we observe *in vitro*, we hypothesize that these latter two similarities may suggest the presence of *inter*molecular avid binding in cells driven by IDR-dependent protein-protein interactions. However, more experiments directly examining this possibility are needed. Genomic localization in cells also exhibited IDR-dependent binding to CG-dinucleotide rich sequences not detected *in vitro*, suggesting impacts of either interactions in *trans* (*i.e.* interactions between IDRs and other nucleic acid-binding proteins) or interactions dependent on post-translational modification of Msn2’s IDR.

Overall, predictions of binding to promoter sequences across *in vitro* (BET-seq) and *in vivo* (ChEC-seq) models were less well correlated than previously characterized human or yeast TFs (Pearson *r*^2^ = 0.23 to 0.43 and 0.26 to 0.39 for DBD and FL Msn2 compared to *r*^2^ = 0.44 (MAX), 0.58-0.59 (Myc), 0.58 (GABPA), 0.40 (E2F1), 0.67 (Pho4), and 0.66 (Cbf1))^54^. Greater differences between binding *in vitro* and in cells for Msn2 may reflect a larger role for mechanisms absent in our *in vitro* binding assays, such as competition with TFs and nucleosomes, TF-RNA binding, post-translational modifications that may alter IDR-DNA binding, and IDR-mediated interactions with other TFs, cofactors, and transcriptional machinery that concentrate interacting proteins around genomic loci^33,57,58^. Alternatively, these differences could simply reflect the fact that methods that permeabilize cells to allow nuclear access and activity of antibody- or TF-conjugated nucleases (*e.g.* CUT&RUN^62^ or ChEC-seq) may rapidly deplete nuclei of proteins weakly associated with chromatin, thereby preferentially emphasizing DBD-dependent binding^28^.

While this study is limited to the TFs Msn2, Msn4, and Com2, DBD-proximal arginine-rich motifs (ARMs) linked to TF-RNA binding were found in 80% of human TFs across diverse TF structural families^52^. These ARMs are similar to the cluster of di-arginine pairs we identify as necessary and sufficient for IDR-DNA binding, suggesting that direct IDR-DNA binding through similar mechanisms may be relevant for other TFs and organisms not examined here. More broadly, models integrating multivalent IDR-DNA binding will be important for understanding TF-chromatin interactions. Overall, our results challenge the traditional view that TF DBDs and IDRs function independently and highlight the importance of considering full-length TFs and multivalent binding when interpreting DNA binding specificity and genomic occupancy.

## Methods

### Sequence alignment and conservation analysis

Msn2, Msn4, and Com2 amino acid sequences were aligned and visualized using the NCBI COBALT tool^63^ (**Fig. 1A**). Full length Msn2 and Msn4 ortholog amino acid sequences were aligned using MUSCLE, visualized using Jalview^64^, and colored by Blosum62 score (**Supplementary Fig. 49**). DNA binding domain annotations used can be found in **Supplementary File 5**.

### TF expression plasmid cloning

Genomic DNA was extracted from *S. cerevisiae* (BY4741) using standard Lithium Acetate precipitation. Full length Msn2, Msn4, and Com2 ORFs were amplified with homologous overhangs from genomic DNA through PCR with hot-start Q5 (New England Biolabs, M0494) and cleaned up using Zymo Clean and Concentrator-5 (CC-5) columns (Zymo D4004). Vector backbone containing an N-terminal 3xFLAG tag and C-terminal meGFP tag (full vector sequence available in **Supplementary File 5**) was amplified by PCR, treated with 1 µL of DpnI (New England Biolabs, R0176L) to remove intact template plasmid, and cleaned up as above. If undesired fragments were present, desired PCR fragments were further purified by gel extraction using the QiaQuick Gel Extraction kit (Qiagen, 28704) with manufacturer recommended protocols after running the PCR products on a 1% agarose, 1x TAE (Tris-Acetate-EDTA) gel stained with 1x GelGreen (Biotium, 41005). Purified ORFs were assembled into amplified expression vector by combining 0.03 pmol of insert and vector backbone with an equal volume of 2x NEBuilder HiFi DNA assembly mix (New England Biolabs, E2621) and incubating for 1 hour at 50°C.

Because the basic RGRK motif in the disordered 15 amino acid region upstream of the C2H2 zinc fingers of Com2 impacts DBD binding preferences and is partly responsible for its paralog-specific binding^39^, all DBD-only constructs tested in this study included the 15-18 amino acids upstream of the annotated zinc finger domains (annotations found in **Supplementary File 5**). IDR-only constructs also contained these 15-18 amino acid regions upstream of the zinc-fingers.

DBD-only, IDR-only, and Msn2 IDR-truncation expression vectors were constructed by amplifying subsections of the full-length (or IDR-only) plasmids, clean up with Zymo CC-5 columns, and simultaneous phosphorylation of 5’ ends, ligation of blunt ends, and digestion of template plasmid using KLD (Kinase-Ligase-DpnI) mix (NEB, M0554) with manufacturer recommended protocols. Chimeric TF expression vectors were constructed by PCR amplification of the two fragments to be combined and assembly as above.

All constructs in this study were transformed into NEB-5alpha chemically competent *E. coli* (New England Biolabs, C2987) using standard procedures and plated on LB-carbenicillin (100 µg/mL) plates. Carbenicillin-resistant clones were picked, grown for 16 hours in LB supplemented with 100 µg/mL carbenicillin at 37°C with 250 rpm shaking, and miniprepped using the QiaPrep spin miniprep kit (Qiagen, 27106) according to manufacturer instructions. Plasmid sequences were confirmed using either Sanger sequencing (ELIM Bio) or whole plasmid sequencing (Plasmidsaurus).

### Transcription factor production and purification

To ensure that we measured the correct full-length constructs, we expressed each construct with an N-terminal 3X-FLAG tag and a C-terminal meGFP tag using PURExpress (New England Biolabs, E6800L) *in vitro* transcription/translation (IVTT) mix with 200 ng of plasmid per 25µL of IVTT mix, allowing expression to proceed for 8 hours at 30°C. We then performed an initial purification by immunoprecipitation of the N-terminal 3xFLAG tag with magnetic anti-FLAG agarose beads (Thermo Scientific, A36797) (**Supplementary Figs. 62, 63**). Briefly, for each TF sample, 10 µL of beads were washed two times in 100 µL of 1x WB buffer (20 mM KH_2_PO_4_, 200 mM KCl, 10µM ZnCl_2_, 0.05% (v/v) Tween-20) and resuspended in 125 µL 1x WB buffer. 125 µL of washed beads were added directly to IVTT mixtures and TF was allowed to bind to beads for 1 hour covered with foil, at room temperature, and with end-over-end rotation to ensure beads remained in solution. TF-bound beads were collected using a 96-side magnetic plate (Invitrogen, 12331D), supernatant removed, and TF-bound beads washed by adding 175 µL of 1x WB buffer and rotating tubes end-over-end for 10 minutes. Beads were collected, supernatant removed, and TF eluted by incubating beads in 20 µL of 1.5 mg/mL 3xFLAG peptide (Thermo Scientific, A36805) in 1x WB buffer for 20 minutes with end-over-end rotation. Supernatant with eluted TF was collected and beads were eluted one additional time. To remove any aggregated protein, eluted TF was centrifuged at 20,000 x g at 4°C for 10 minutes and 30 µL of protein remaining in solution was collected. To ensure proteins were loaded onto microfluidic devices at consistent concentrations, meGFP fluorescence of each TF sample was measured using the blue excitation and green emission channels in a fluorometer (DeNovix). Using 30µL of 1x WB buffer as a blank, TF concentration (rfu/µL) was roughly determined by subtracting blank fluorescence and dividing the background subtracted fluorescence by 30 and all TF samples were normalized to the concentration of the most dilute TF sample in 1x WB buffer. TF samples were then flown onto devices patterned with biotinylated anti-eGFP VHH (Proteintech, gtb), thus immunoprecipitating the TFs using the C-terminal meGFP tag.

### Degenerate DNA library production and characterization

To ensure equal coverage of each single nucleotide variation of the AGGGG consensus motif, we designed the ‘degenerate motif’ library as 5 separate single-stranded oligonucleotides with 4 degenerate ‘N’ positions up- and downstream of NGGGG, AHGGG, AGHGG, AGGHG, or AGGGH (where H denotes A, C, or T nucleotides) binding sites embedded within constant flanking sequences, with the 5’ end starting with the Illumina TruSeq Read 2 sequence and the 3’ end ending with a primer landing site for primer extension and addition of UMIs during library prep (full sequences for all MITOMI experiments in this study available in **Supplementary File 6**). Each oligonucleotide was ordered from Integrated DNA technologies as custom DNA oligos with degenerate positions receiving hand-mixed nucleotides at equimolar ratios and standard desalting purification.

We duplexed each library oligonucleotide by primer extension by mixing 9 µL of 100 µM oligo with 9 µL of common 3’ annealing primer, 9 µL of 10 mM dNTPs (Thermo Scientific, R0191), 3 µL of 10x NEBuffer 2 (B7002S) and annealing by heating the mixtures in a thermocycler to 94°C for 3 minutes and ramping down to 37°C at 1°C per 50 seconds. To each reaction, we then added 1.5 µL of Klenow fragment, exo – (New England Biolabs, M0212L), 1.5 µL of 10x NEBuffer 2, and 12 µL of UltraPure water (Invitrogen, 10977015), and extended the annealed primers by incubating at 37°C for 1 hour, inactivated the polymerase at 72°C for 20 minutes, and ramped down to 10°C at a rate of −1.5°C per 60 seconds. To remove remaining unextended oligonucleotides, we added 2 µL of Exonuclease I (New England Biolabs, M0293L) to each reaction, incubated at 37°C for 40 minutes, heat inactivated the enzyme for 15 minutes at 80°C, and ramped to 10°C at −0.1°C per second. We then buffer exchanged each reaction and removed unincorporated nucleotides using Amicon Ultra10 kDa molecular weight cut off (MWCO) filter spin columns (Millipore Sigma, UFC501024). 47 µL of each reaction was mixed with 400 µL of TloE (10 mM Tris pH 8.0, 0.2 mM EDTA), added to a MWCO column, and concentrated to ∼40 µL by centrifugation for 10 minutes at 10,000 x g. We repeated this filtering step 3 times, collected the retained solution, and brought the DNA samples to a volume of 60 µL with TloE. Extended oligonucleotide concentrations were then quantified by measuring the absorbance at 260 nm with a NanoDrop spectrophotometer (Thermo Scientific, ND-ONE-W). We checked products for correct size (90 base pairs) and purity by running 10-fold dilutions of extension product on a 2.5% agarose buffered with 1x Tris-Acetate-EDTA (TAE) and stained with 1x GelGreen and comparing to a 50 base pair DNA ladder (New England Biolabs, N0556S). Since the first library oligo (NNNN-NGGGG-NNNN) contained a greater number of unique sequences (4^9^) compared to the other four oligos with H degenerate positions (4^8^ x 3), we mixed the extension products to achieve equimolar ratios of each unique sequence at a total concentration of 2 µM in TloE.

### Microfluidic device fabrication and operation

Flow and control molds were fabricated as described previously^35,37^ and all design files are available on the Fordyce Lab website (http://www.fordycelab.com/microfluidic-design-files). We fabricated two-layer MITOMI devices from PS1800 4- and 8-inlet (PS4 and PS8 respectively) molds using polydimethylsiloxane (PDMS) polymer (RS Hughes, RTV615) in the Stanford Microfluidics Foundry as described^65,66^. Two-layer BET-seq devices were fabricated from Spline 6-block molds (**Supplementary Fig. 64**) as above.

Microfluidic devices were controlled by a custom pneumatic manifold^67^ and imaged with a fully automated microscope (Nikon Ti2) and custom software (https://github.com/FordyceLab/RunPack)^68,69^.

Initial setup of control lines of devices and loading of reagents onto devices were performed as previously described^66^. Once control lines were pressurized and tested, device surfaces were passivated by dead-end filling with 5 mg/mL ultrapure bovine serum albumin (BSA, Invitrogen, AM2616) with ‘button’ valves (and ‘neck’ valves if running MITOMI devices) pressurized and closed and then flowing the BSA through each flow inlet. Sandwich valves were actuated while devices were filled with BSA to ensure passivation of surfaces around button edges. Devices were then washed with 1x phosphate-buffered saline (PBS, diluted from 10x stock, Corning 46-013-CM), flowing again through each flow inlet, for 10 minutes.

### Microscopy Instrumentation

Device imaging was performed using a Nikon Ti-2 Microscope equipped with motorized XY-stage (Applied Scientific Instruments, MS-2000 XY stage), LED light source (Lumencor SOLA SE Light Engine), and CMOS camera (Andor Zyla 4.2). The Nikon Ti-2 core itself was equipped with an automated Z-axis, turret aperture, and filter turret, containing pre-arranged filter cubes for EGFP (Semrock, GFP-4050B), Cy5 (Semrock, Cy5-5070A). The microscope stage, filter, and camera were controlled using Micro-Manager^70^; all images were acquired with a 4x objective lens (CFI Plan Apo 4X NA 0.20, Nikon) and 2×2 binning.

### BET-seq relative affinity measurements for ‘degenerate motif’ DNA library

Spline 6-block devices were placed on epoxy- or amino-silane treated 25×75 mm glass slides (ArrayIt, SME2M48BC or Santa Cruz Biotechnology, sc-363560) cleaned by wiping with isopropanol and bonded to slides by baking for 4 hours at 80°C on a programmable hot plate. Initial set up and passivation of devices was performed as above. Device surfaces under button valves were patterned sequentially with biotinylated BSA (bBSA; 2 mg/mL, ThermoFisher Pierce, 29130), NeutrAvidin (1 mg/mL, Thermo Scientific, 31000), and anti-eGFP VHH (16 µg/mL, Proteintech, gtb) as previously described^66^ with the following modifications: anti-eGFP VHH was diluted to 16 µg/mL in 1x PBS with 280 µg/mL bBSA and reagents were flown onto the devices at constant pressure of 1.8 – 2.1 psi. After surface patterning, 20 µL of meGFP-tagged TF preparations were flown onto devices through block-specific inlets (6 TF constructs per device) at ∼1.8 psi with button valves open to allow for TF pull-down. To assist with anti-eGFP VHH saturation, TFs were flowed enough to visibly load the entire device, flow was stopped, and TFs allowed to bind button surfaces without flow for ∼10 minutes. This flow-stop-bind cycle was repeated until button surfaces were saturated and did not continue accumulating meGFP fluorescence. Button valves were then closed and excess TF was washed out of devices with 1x PBS, and devices were imaged in the GFP channel at 200 or 500 ms to quantify TF concentrations and confirm purification and completion of device washing.

We prepared 15 µL of 1 µM ‘degenerate motif’ library for introduction to the device by mixing 7.5 µL of 2 µM library in TloE with 7.5 µL of 2x binding buffer (300 mM NaCl, 20 mM Tris-HCl pH 7.5, 2 mM DTT, 0.02 mM ZnCl_2_, 0.1% (v/v) Tween-20, 0.5 mM Spermidine, 2.1 mM Magnesium Acetate, 53 mM Potassium Acetate, 0.5% ultrapure BSA, and 6 mg/mL Trehalose) with DTT, Spermidine, and BSA added fresh. Library was then flowed onto the device through a common inlet at 2.5 psi flow pressure until about ¼ inch of library is left in the reagent tubing, at which point we closed outlet valves to stop flow and opened button valves to fully fill the device. Inlet valves were then closed and the binding reaction was allowed to come to equilibrium for 90 minutes.

After binding reactions were complete, we closed button valves to trap the bound fractions, connected 100 µL gel-loading pipet tips (USA Scientific, 1022-0810) to each of the 6 block-specific outlets, and eluted the unbound fractions by flowing elution buffer (20 mM Tris-HCl pH 8.0, 0.2 mM EDTA, 0.05% (v/v) Tween 20, 0.25% BSA) through block-specific inlets into the pipet tips for 5 minutes at a flow pressure 2.5 psi. Unbound fractions were then transferred to 0.5 mL DNA LoBind tubes (Eppendorf, 022431005) and stored at −20°C. Devices were then washed by flowing elution buffer for 10 minutes at a flow pressure of 3.5 psi.

To elute bound fractions from button surfaces, we first flowed on ∼60 µL of digestion buffer with proteinase K (1 mg/ml Proteinase K (New England Biolabs, P8107S), 1x NEBuffer 2, 0.05% Tween 20) through a common inlet at 3 psi for ∼10 minutes (or until ∼1 inch of liquid remains in the reagent tubing) with button valves closed. We then closed outlet valves, opened button valves, dead-end filled the digestion reaction mixture into the device for 5 seconds, closed inlet valves, and allowed the digestion of immobilized TF to proceed for 20 minutes, checking the progress of TF digestion by imaging devices in the GFP channel at 500 ms. We again connected 100 µL gel-loading pipet tips into the block specific outlets and flowed elution buffer into the tips for 5 minutes at a flow pressure of 3.5 psi. Eluted bound fractions were transferred to 0.5 mL DNA LoBind tubes and stored at −20°C.

### BET-seq library preparation and sequencing

5 µL of bound fractions, 1 µL of 1:50 dilutions of unbound fractions, and 3 µL of 1 nM dilutions of input libraries were taken for library preparation. Samples were annealed to a primer (oMH_160, consisting of Illumina TruSeq Read 1, 26 degenerate nucleotide positions that act as unique molecular identifiers (UMIs), and a region complementary to the constant primer landing sites within all library sequences) by mixing samples with 100 nM UMI oligo and 50 mM NaCl, with the reaction being brought to 7 µl total with TT (10 mM Tris-HCl pH 8.0, 0.05% Tween-20). Reactions were incubated in a thermocycler at 95°C for 5 minutes to inactivate any residual Proteinase K and then the temperature was ramped down to 37°C at a rate of 1°C per 30 seconds.

To extend the annealed primer, we preheated 7 µL of NEBNext Ultra II Q5 mastermix per reaction to 98°C for 50 seconds to activate the polymerase, added 7 µL of activated mastermix to each sample, incubated them for 1 hour at 55°C, and then slowly ramped to 10°C at a rate of 0.2 °C per second. To remove excess UMI primer, we added 1 µL of Exonuclease I (New England Biolabs, M0293L), incubated reactions at 37°C for 45 minutes, heat inactivated the nuclease at 80°C for 20 minutes, and slowly ramped to 10°C at a rate of 0.2°C per second. We then amplified the libraries and attached sequencing indexes and Illumina P5 and P7 adapters by adding to each reaction 8 µL of NEBNext Ultra II Q5 mastermix, 0.5 µM of forward (oMH_185) and reverse primers (oMH_161-184 or oMH_242-265, sequences available at Zenodo^71^), and water to 30 µL total and using the following thermocycling conditions: 98°C 30 seconds, 12x [98°C 5 seconds, 63°C 10 seconds, 72°C 20 seconds], 72°C 1 minute, 4°C. Amplified library was then precipitated onto magnetic beads by adding 24 µl of 20% PEG-8000/2.5 M NaCl and 2 µl carboxylated speedbeads (Cytiva, 24152105050250) for 8.5% PEG-8000 final and incubating at room temperature for 10 minutes. Beads were collected on a 96-side magnetic plate (Invitrogen, 12331D), supernatant was removed, and beads were washed two times with 150 µL of 80% DNase-free ethanol by resuspending beads in the 80% ethanol, collecting beads, and removing the wash solution. After the final wash was removed, beads were collected on the magnetic plate and allowed to dry for 10-15 minutes (or until beads appeared cracked). Precipitated library was eluted from the beads in 20 µL of TT (10 mM Tris-HCl pH 8.0, 0.05% Tween-20), library concentrations were measured using the high-sensitivity dsDNA Qubit detection kit (Thermo-Fisher, Q32854) with the Qubit Flex Fluorometer (Thermo-Fisher, Q33327), and library size and cleanliness were confirmed by running libraries on 2% agarose gels buffered with 1x TAE and stained with 1x GelGreen.

Libraries were then pooled at equimolar ratios at final concentrations of 4-10 nM in TT, and sequenced single-end (130 cycles, 8 cycles index) with the Chan-Zuckerberg Biohub sequencing core on an Illumina NextSeq 2000, NovaSeq S4, or NovaSeq X with 14-25% PhiX spike-in to offset the lack of library diversity in constant regions.

### BET-seq sequencing data processing for ‘degenerate motif’ library

Raw base calls were converted to fastq format and demultiplexed using Bcl2Fastq2. For all subsequent steps, we used GNU Parallel^72^ to parallelize jobs submitted to the Stanford Sherlock computing cluster. UMIs were then extracted from reads using UMI-tools^73^ extract with the following pattern setting: (26)N(104)X. Next, we used cutadapt^74^ to trim constant sequences from ‘degenerate motif’ library reads and remove any reads that were not the correct length (13 bp) post-trimming with the following parameters: cutadapt -j 5 -m 20 --nextseq-trim=20 -a ^GTCATACCGCCGGACAAGACTCAGATACATT… AATACTGATGAAGATCGGAAGAGCACACGTCTGAACTCCAGTCAC --discard-untrimmed --minimum-length 13 --maximum-length 13. We then used fastp^75^ to filter reads on read quality, requiring a minimum of Q23 for all base calls, with the following settings: fastp --cut_right --cut_right_window_size 1 – cut_right_mean_quality 23 --length_required 13 --length_limit 13 -A -Q -w 5 -z 4. We then aligned the trimmed and filtered reads to an index containing all possible sequences in the ‘degenerate motif’ to convert the files to SAM format and confirm integrity of the reads using Bowtie2^76^ using default settings. Aligned reads were then converted to sorted BAM files using Samtools^77^, after which we used UMI-tools dedup to deduplicate reads using the “unique” method. The deduplicated BAM file was then converted back to SAM format, and the number of sequences matching one of the possible 4^10^ sequences were counted using a custom python script. Reads that did not match a designed library sequence (0.2% of all unique reads for the ‘degenerate motif’ library) were not used for subsequent analysis. Designed library sequences that were unobserved (i.e. had no reads matching them) in all samples were removed from further analysis.

We then computed probabilities of observing each sequence in the bound and input libraries by normalizing the read counts to 30 million reads, adding a pseudo-count of 4 to each count, and then taking the ratio of each count to the total read counts (including pseudocounts) for each sequence using the following equations:

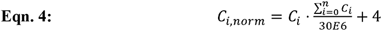

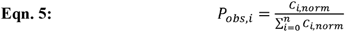

where *C_i_* is the number of unique reads for the *i*^th^ sequence, *C_i,norm_*, is the normalized read count for the *i*^th^ sequence, and *P_obs,i_*, is the probability of observing the *i*^th^ sequence. We then calculated relative binding energies of each sequence as previously described^35^ using the following equation:

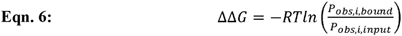

where *R* is the ideal gas constant (1.987 E-3 kcal/mol/K), T is the temperature in Kelvin (298 K), *P_obs,i,bound_* is the probability of observing the *i*^th^ sequence in the bound fraction, and *P_obs,i,input_*, is the mean probability of observing the *i*^th^ sequence across input fraction replicates.

### BET-seq data centering and normalization for the ‘degenerate motif’ library

Because BET-seq ΔΔGs are not centered on the ΔΔG value of any particular sequence, we first centered ΔΔG measurements on a single sequence (AGTTCGGGGAACG for the ‘degenerate motif’ library) selected by virtue of having minimal differences between the median ΔΔG values for all protein constructs and replicates. To account for variation between experiments, we then normalized ΔΔG measurements for a given protein across all experimental replicates to a reference replicate. We designated the replicate with the highest ΔΔG dynamic range as the ‘reference’ experiment and normalized replicates by performing a linear regression against the reference experiment using sequences with ΔΔG < −0.8 in the reference experiment and transforming ΔΔG values using the fit slope and intercept parameters. These normalized ΔΔG values were then used in replicate-to-replicate comparison plots.

We then calculated the mean normalized ΔΔGs between experimental replicates and recentered these values on the mean normalized ΔΔG of a sequence (CTTCCGGGGCCGG for the ‘degenerate motif’ library) selected again for having minimal differences with median mean normalized ΔΔGs of all protein constructs. These recentered mean normalized ΔΔGs are those shown in plots comparing relative binding energies of different protein constructs and all other analyses use these values unless otherwise stated.

### BET-seq data analysis

To distinguish between the signal and ‘noise’ of BET-seq ΔΔG measurements, we devised a method to determine how far a given ΔΔG is from the experimental ‘noise’ distribution as follows. First, we found the empirical ‘center’ (µ_noise_) of ΔΔGs distributions by splitting the values across 50 bins of equal size, selecting the ΔΔG bin with the highest count, and taking the mean of the upper and lower bin edge values. Since true binding events form a characteristic tail on the left side of the distribution (i.e. the more negative ΔΔG values), we constructed a ‘noise’ distribution by taking the ΔΔG values greater than the ‘center’ ΔΔG and reflecting these across that center ΔΔG. We then fit a normal distribution to this ‘noise’ distribution, extracting a standard deviation (std_noise_) parameter (**Supplementary Fig. 65**). We then used µ_noise_-3(std_noise_) as a rough threshold to determine whether sequences were either ‘bound’ or ‘unbound’ in a given experiment.

### Motif finding with STREME

STREME^44^ was used to perform set-based motif finding using dinucleotide shuffled input sequences as background sequences, a minimum motif length of 5 bp, and otherwise default parameters.

### Extraction of energetic penalties of single nucleotide variants of AGGGG consensus motif

To determine energetic penalties of each single nucleotide variant of the AGGGG motif, we first calculated, for each construct, pairwise differences in ΔΔG between sequences with a single nucleotide variant of the core motif (eg. TGGGG) and those with the same flanking nucleotides but surrounding a consensus AGGGG motif, summarized by the following equation.

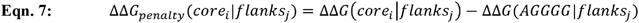

Reported ΔΔG penalties are the medians of the resultant ΔΔG penalty distributions for each core motif variant across all flanking nucleotide combinations excluding those that contain or create alternative AGGGG motifs (**Supplementary Fig. 5**).

### Linear mononucleotide modeling of motif-flanking nucleotide energetic contributions

To fit linear mononucleotide models of AGGGG-flanking nucleotides, we first filtered the ‘degenerate motif’ library for sequences with the AGGGG consensus motif and recentered ΔΔGs of each replicate of each construct on the median ΔΔG value of these filtered sequences. Motif flanking sequences 4 bp up- and downstream of the AGGGG core were then one-hot encoded. We then used the least squares method (scipy) to optimize a model with additive weights for each nucleotide in each flanking nucleotide position, bounding weight values at −4 and 4 kcal/mol.

Model-predicted ΔΔGs were calculated by taking the dot product of the one-hot encoded flanking sequence matrix and the model weight matrix (i.e. the sum of the weights of the nucleotides present in each sequence).

### AffinityNet model training on BET-seq data

We trained AffinityNet, a Convolutional Neural Network (CNN), on the BET-seq ‘degenerate motif’ and ‘promoter’ libraries. AffinityNet takes one-hot encoded sequences and simultaneously predicts binding affinities for a single Transcription Factor (TF) across multiple constructs with a given DNA binding domain. The model architecture comprises eight sequential convolutional layers, each with 256 filters and a kernel size of 8, with Rectified Linear Unit (ReLU) activation applied between layers. The output from the convolutional blocks was average-pooled across the sequences and then passed through a dense layer to predict TF-DNA affinity. We utilized Mean Squared Error (MSE) as the primary loss function between predicted and measured activities. To encourage the model to capture differences between FL, DBD, and chimera constructs, we introduced an additional differential loss term defined as the mean squared error (MSE) between the predicted and experimentally measured differences for construct pairs (FL vs. DBD and chimera vs. DBD). Models were trained for a maximum of 100 epochs using an initial learning rate of 1×10−4. To mitigate overfitting, we employed early stopping with a patience of 8 epochs. Additionally, a learning rate scheduler was implemented to halve the learning rate following 4 consecutive epochs of stagnant validation loss. All models were trained using five-fold cross-validation. For the promoter library, data were partitioned by chromosome, while the degenerate library was partitioned via random assignment. We trained independent models for each TF and library combination, resulting in six distinct training configurations. Implementing a five-fold cross-validation strategy for each setup resulted in a total of 30 trained model instances. In addition to the main ‘promoter’ and ‘degenerate motif’ library models, we trained dedicated AffinityNet models on the IDR-only construct binding data with the ‘promoter’ library to characterize their sequence-dependent binding. These models were trained independently of the primary ‘degenerate motif’ and ‘promoter’ libraries to avoid cross-task feature leakage, ensuring that the learned representations reflected only the IDR constructs. To account for experimental uncertainty, the model is trained to predict the activity of individual replicates during training. During inference and interpretation, we produce a finalized prediction by averaging across all replicates and folds.

### Interpreting AffinityNet using DeepSHAP/DeepLift

To characterize the features learned by AffinityNet, we employed the DeepSHAP/DeepLift algorithm (https://github.com/kundajelab/shap) to generate base-resolution sequence contribution scores. DeepLift is an interpretation method that utilizes backpropagation to quantify the impact of individual nucleotides by decomposing the difference in model output between the input sequence and reference sequences. We interpreted our models using their respective training data: models trained on the promoter library were interpreted using promoter library sequences, and those trained on the degenerate library were interpreted using degenerate library sequences. Interpretations were performed on the predicted binding affinities for each TF construct, averaged across replicates, using 20 dinucleotide-shuffled versions of the input sequence as reference sequences. To further identify the features driving differences in binding affinity between constructs, we performed differential interpretation by applying DeepLift directly to the predicted differences in binding affinity between constructs. All interpretations were averaged across folds for downstream analysis.

### Motif discovery on the ‘degenerate motif’ library AffinityNet

The ‘degenerate motif’ library contained a massive number of sequences (1,048,576), with all positions fixed outside the central 13 bp. To accommodate this, we implemented a custom algorithm to extract learned patterns directly from the models trained on the degenerate library.

First, to identify sequences with binding events within the degenerate library, we modeled the background noise as a normal distribution. We first estimated the mean by binning the data into 1,000 intervals and identifying the most frequent bin as the center. Because true binding events formed a distinct tail on the left side of the distribution, we excluded these events from our noise estimate by fitting a Gaussian model exclusively to the right half of the data and reflecting it across the noise mean, as described above. Finally, we isolated the bound sequences by selecting all data points with a binding energy more than four standard deviations (4σ) below the modeled mean.

Next, we extracted the central 13-bp variable regions from the interpretation scores and calculated sequence similarity using cosine similarity. To construct a sparse k-nearest neighbors (k-NN) graph, we processed the data in batches of 4096, retaining only the top 1000 edges per batch that exceeded a similarity threshold of 0.7. We then applied the Leiden algorithm to cluster the sequences with an initial resolution parameter of 0.5. To avoid over-fragmentation, if the algorithm generated more than 200 clusters, we re-clustered the sequences by reducing the resolution by 0.1 until fewer than 200 clusters were obtained.

Finally, to account for variable motif positions within the sequences, we aligned the motifs generated in the previous step by allowing shifts of up to ±3 bp. We then clustered these aligned motifs using the Leiden algorithm at a resolution of 1.0, applying a strict cosine similarity threshold of 0.98.

### Analysis of motif-flanking nucleotide SHAP scores in ‘degenerate motif’ library models

SHAP scores for the ‘degenerate motif’ library were first averaged over the replicate models, and then averaged for each nucleotide and position across sequences sharing the same core motif variant.

### Preparation of DNA for MITOMI

We designed all DNA sequences for MITOMI binding assays with a 3’ region complementary to an Alexa Fluor-647 (AF647) dye-conjugated primer (**Supplementary File 6**). We ordered all DNA sequences as single-stranded oligonucleotides from Integrated DNA Technologies (IDT) with standard desalting purification and shipped in ‘LabReady’ formulation (100 µM in IDTE buffer, pH 8.0). We then duplexed these single-stranded DNA sequences by (1) annealing the universal AF647-labeled primer to the 3’ region of the oligonucleotide and (2) extension using the primer as a template using Klenow fragment, exo-, polymerase. Both steps (1) and (2) were performed as previously described^68^.

After the Klenow extension, we removed any un-extended dye-conjugated primer from the reactions using Amicon Ultra 30 kDa MWCO filter spin concentrator columns (Millipore Sigma, UFC503024). We added 60 µL of duplexed DNA to the filter spin columns, added 400 µL of TloE (10 mM Tris pH 8.0, 0.2 mM EDTA), mixed by pipetting, and concentrated the extended DNA to ∼40 µL by centrifugation for 10 minutes at 10,000 x g. We repeated this filtering step 3 times, collected the retained solution, and brought the DNA samples to a volume of 61 µL.

We then checked each DNA sequence for proper extension by running them on Novex 10% polyacrylamide TBE-Urea denaturing gels (Thermo Scientific, EC68752BOX) as previously described^66^. A sample gel can be found in **Supplementary Fig. 66**.

### DNA library printing

For each 60 µL sample of extended DNA, we added 15 µL of 5x print buffer to 60 µL of extended DNA for 1x print buffer final (1x SSC, 1% BSA, 12 mg/mL Trehalose) and prepared six 1.92-fold serial dilutions of the extended DNA in 1x print buffer. Serial dilutions were then transferred to a 384-well plate using a 96-well Liquidator pipette (Rainin, 17014207) and arrayed on epoxy- or amino-silane treated 25×75 mm glass slides (ArrayIt, SME2M48BC or Santa Cruz Biotechnology, sc-363560) using a SciFlex arrayer S3 (Scienion AG) with a PDC70 (type 1 coating) nozzle as previously described^66^. To ensure replicable and confident measurements, we included at least three spot replicates in randomized positions of each sequence-concentration combination. To prevent and allow for downstream filtering out of contamination between chambers we added a “blank” row of 1x print buffer without DNA in between every 2^nd^ or 3^rd^ row. While printing, we covered the arrayer windows with black vinyl cloth panels to prevent light from bleaching printed DNA spots.

After printing and drying arrays at least overnight, we manually aligned MITOMI devices to the DNA array such that each chamber contained one oligonucleotide spot or one empty buffer spot. If DNA spots were not visible by eye, slides were baked for 1 hour at 50°C to further dry the spots before device alignment. Devices were then bonded to the glass slide by baking on a hot plate at 80°C for approximately 4 hours.

### MITOMI affinity measurements for ‘fiducial’ sequences from ‘degenerate motif’ DNA library

Surfaces beneath button valves were functionalized and meGFP-tagged TFs immobilized in the same manner as in the BET-seq measurements of the ‘degenerate motif’ library. Surface patterning steps were done on day 1 of the experiment and the devices allowed to sit overnight to allow osmosis of gaseous water into fluorescently labelled DNA spots to assist with spot solubilization. We then flowed 2x MITOMI binding buffer (225 mM NaCl, 20 mM Tris-HCl pH 7.5, 2 mM DTT, 20 µM ZnCl_2_, 0.1% (v/v) Tween-20, 0.1% BSA, 0.5 mM spermidine, 2.1 mM magnesium acetate, 53 mM potassium acetate – with DTT, BSA, and spermidine added fresh) onto the device for 10 minutes. To solubilize spots, we then closed outlet valves, opened button valves and allowed for the full chamber volume to dead-end fill with binding buffer for ∼10 seconds, closed inlets and sandwich valves to isolate chambers, and then opened neck valves. We toggled button valves 3-5 times to assist with mixing, left button valves closed, and allowed spots to solubilize and mix with binding buffer for 20 minutes. We then opened button valves and allowed binding reactions to come to equilibrium for 90 minutes.

‘Pre-wash’ images of devices were then taken in the Cy5 channel (50-125 ms exposures) to quantify the total DNA concentrations. Equilibrium bound TF-DNA complexes were then mechanically trapped by closing button valves, neck valves were closed, and unbound DNA was washed away with 1x PBS, after which ‘post-wash’ images were taken of devices in both Cy5 (750-1500 ms exposures) and GFP (200-500 ms exposures) channels to quantify the bound DNA and total TF under each button. Microfluidic devices were imaged with a gridded acquisition, with adjacent images overlapping by 10% to facilitate image stitching (see below for more details).

### MITOMI and BET-seq image processing and quality control

Raw image rasters of fluorescence images covering the full device area were flat-field corrected using manual flat-field images for ‘pre-wash’ Cy5 images as previously described^78^ or an implementation of BaSiCpy^79^ for ‘post-wash’ Cy5 and GFP images and stitched into single full-device images using ImageStitcher, an in-house publicly-available software package (https://github.com/FordyceLab/ImageStitcher).

These stitched images were processed using ProcessingPack, an in-house publicly-available software package (https://github.com/FordyceLab/ProcessingPack) as previously described^66,68^. Briefly, to process ‘pre-wash’ images, the software stamps the stitched image of the device with a 32 x 56 or 24 x 64 array (for PS1800 or Spline 6-block devices respectively), finds the centers of chambers within rough boundaries demarked by the array using a Hough-transform, and quantifies median fluorescence intensities. To process ‘post-wash’ images, circle-finding algorithms identified fluorescent spots under chamber “button” valves and per-spot metrics were mapped to Scienion print array position. Downstream analysis of buttons primarily utilized background-subtracted summed intensities of GFP and Cy5 per-spot fluorescence. ‘Pre-wash’ Cy5 images and ‘post-wash’ GFP images can then be converted to effective DNA/TF concentrations per chamber respectively using standard curves (described below).

Before analyzing data, we manually examined images of each device and culled data from device chambers and/or columns based on the following criteria: (1) incomplete spot solubilization, (2) evidence of DNA contamination between chambers in the form of either AF647 fluorescence in ‘blank’ chambers or ‘pre-wash’ chamber Cy5 fluorescence intensities that are outliers relative to the distribution of intensities of replicate spots, (3) protein aggregates within the chamber, (4) chambers with visible debris and/or device defects that otherwise did not impair device operation.

### DNA fluorescence standard curve

To convert measured AF647 fluorescence intensities to DNA concentrations, we measured the fluorescence intensities of a series of known concentrations of AF647-labelled DNA within our devices as previously described with modifications^66^. Briefly, we prepared AF647-labelled dsDNA as above, and made a series of two-fold dilutions in 1x binding buffer (150 mM NaCl, 10 mM Tris-HCl pH 7.5, 1 mM DTT, 0.01 mM ZnCl_2_, 0.05% (v/v) Tween-20, 0.25 mM Spermidine, 1.05 mM Magnesium Acetate, 26.5 mM Potassium Acetate, 0.25% ultrapure BSA, and 3 mg/mL Trehalose) targeting the following concentrations: 4 µM, 2 µM, 1 µM, 500 nM, 250 nM, 125 nM, and 62.5 nM. We then quantified DNA concentrations by measuring A260 using a NanoDrop spectrophotometer.

To measure fluorescence intensities of these DNA concentrations, we used PS8 MITOMI devices as above, passivating devices with 5 mg/mL BSA as in a standard MITOMI experiment. We then flowed each DNA concentration onto the device through block-specific inlets for 10 minutes with button valves open, and imaged the device in the Cy5 channel with a 100 ms exposure. Using the same image processing steps as above, we quantified median fluorescence intensities within each chamber associated with each known DNA concentration. Using all chambers, we then fit a line to the relationship between the DNA concentrations and the median chamber fluorescence intensities, constraining the y-intercept (*b*) to be greater than or equal to 0.

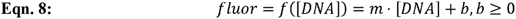

To convert fluorescence intensities of any exposure time to DNA concentrations we used the following formula:

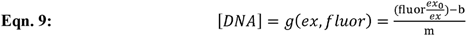

where *fluor* is the median chamber Cy5 intensity, *ex*_0_ is the exposure of standard curve images, *ex* is the exposure of the ‘pre-wash’ images, and *b* and *m* are the fitted coefficients from the equation above.

### Determination of equilibrium binding constants

To determine equilibrium binding constants from MITOMI data, we first converted ‘pre-wash’ median Cy5 fluorescence intensities into DNA concentrations using the parameters fit from a standard curve above. Then, as a proxy for fractional binding, we quantified the DNA bound by surface immobilized TF as the ratio (*R*) of summed, background subtracted ‘post-wash’ DNA fluorescence (AF647) to summed, background subtracted ‘post-wash’ TF fluorescence (GFP). Finally, we fit a Langmuir isotherm to these data:

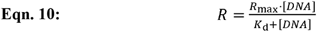

where *R*_max_ is the fit saturation point of the binding curve. In MITOMI experiments measuring binding equilibrium constants for ‘fiducial’ sequences from the ‘degenerate motif’ library, *R*_max_ values were fit globally^37,66^ across all sequences tested for binding with a given TF while *K*_d_ values were fit for each oligo individually to allow for accurate estimation of *K*_d_s of weak binders that did not saturate within the DNA concentrations tested. In MITOMI experiments measuring binding of promoter tiles containing multiple potential binding sites, for which global *R*_max_ fits did not yield a good fit to the data (such as in Fig. 4D, E), we locally fitted *R*_max_ values for each oligo given that we expect avidity to alter the stoichiometric ratios of TF to DNA.

To minimize the influence of outliers on estimation of *R*_max_ and *K*_d_ fits, we randomly selected 70 percent of the data without replacement for each sequence tested, fit *R*_max_ and *K*_d_ parameters to the subset of data, and then repeated this process 400 times to create a distribution of fit parameters. For individual binding curves, the *R*_max_ and *K*_d_ values and errors reported represent the medians and interquartile ranges of the fit distributions, excluding failed fits, which default either to the initial *K*_d_ guess or maximum *K*_d_ bound allowed.

We used three conditions to determine whether there was detectable binding for a given TF-DNA sequence pair. (1) We required that bound DNA signal roughly be increasing by requiring that the median “post-wash” summed background AF647 fluorescence (BgSub AF647) within button valve areas of the chambers containing the top 50% of [*DNA*] concentrations must be greater than the median “post-wash” BgSub AF647 over button areas of chambers containing no DNA for a given experiment. (2) We required that the distribution of “post-wash” BgSub AF647 over button areas for a given TF-DNA sequence pair (the ‘test’ distribution) was significantly different from the distribution of “post-wash” BgSub AF647 over button areas of chambers with no DNA (the ‘null’ distribution). To test this, we calculated the observed Wasserstein Distance between the ‘test’ and ‘null’ distributions, took 10,000 random samples of *N* values from the ‘null’ distribution with replacement, where *N* is equal to the number of measurements in the ‘test’ distribution, calculated the Wasserstein Distance between the whole ‘null’ distribution and each random sample, and then calculated a p-value by calculating the probability that the Wasserstein Distance between the full ‘null’ distribution and the random samples of the ‘null’ distribution were greater than the Wasserstein Distance between the ‘null’ and ‘test’ distributions. (3) Finally, we required that the median *K*_d_ fit for a given interaction be below the maximum *K*_d_ value allowed during the fitting, or 100 µM. If any of these conditions were not met, we reported a *K*_d_ fit value as “binding not determined”.

To minimize over-interpretation of *K*_d_ fits to very strong or weak binders, we report *K*_d_ values below or above thresholds at which fit values are expected to deviate > 1.4-fold from “true” *K*_d_s as upper and lower limits, respectively, based on thresholds set through binding simulations, as described below.

### Determination of equilibrium binding constant upper and lower limit thresholds

To determine lower and upper threshold *K*_d_ values under and over which, respectively, to call MITOMI fit *K*_d_ values as limits, we needed to estimate *K*_d_ ranges within which our *K*_d_ fits accurately represent the “true” *K*_d_s of the measured interactions given our experimental conditions (**Supplementary Fig. 67**). To accomplish this, we simulated a 2-state binding model in which the two states are comprised of free TF and DNA and a DNA-bound state (B) whose concentrations are described by the following equations:

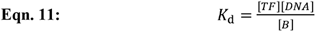

The total concentrations of TF and DNA are also described thus:

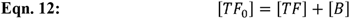

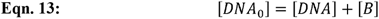

We then solved for [*B*] in terms of [*TF*_0_], [*DNA*_0_], and *K_d_* using the Python package Sympy, and simulated binding curves for synthetic DNA sequences over a range of binding strengths from 6 nM to 250 µM as listed below in units of nM:

“True” = [6, 12.5, 25, 50, 100, 250, 500, 1000, 1500, 2000, 2500, 3000, 3500, 4000, 4500, 5000, 6000, 7000, 8000, 9000, 10000, 12500, 15000, 17500, 20000, 25000, 30000, 35000, 40000, 45000, 50000, 55000, 65000, 70000, 75000, 80000, 85000, 90000, 95000,100000,125000, 150000, 175000, 200000, 250000]

To mimic experimental conditions, binding curves were sampled along two sets of log-spaced [*DNA_0_*] concentrations seven concentrations from 32.5 to 1000 nM or 62.5 nM to 3000 nM, each concentration of which was represented with three replicates and to each of which jitter was added by multiplying by a random factor normally distributed with a mean of 1 and standard deviation of 0.15. Bound TF concentrations were calculated for each [*DNA*_0_] and *k_d_* using [*TF*_0_] = 50 nM. Experimental noise was simulated in two ways: by (1) multiplying the true fraction bound by a random log normal distributed factor (mean = 0, standard deviation = 0.2), and (2) adding a random normally distributed value (mean = 0.015, standard deviation = 0.0156) to the fraction bound. Bound fractions were simulated across 10 different random seeds.

These two noise factors were parameterized as follows. First, to estimate the spread in fractional binding between measurements done with roughly equal [*DNA_0_*] concentrations (multiplicative noise), we fitted log normal distributions to the measured “post-wash” AF647/GFP ratios for each TF-oligo-dilution combination across all MITOMI experiments in which there was measurable binding and for which there were at least 3 chambers with replicate DNA dilutions for a given TF-oligo combination post-chamber culling. The standard deviation of the multiplicative log normal noise was determined by taking the average standard deviation across all fits (**Supplementary Fig. 68**). Second, to estimate the background noise in imaging (additive noise), the standard deviation of the additive noise was determined by fitting normal distributions to the measured “post-wash” AF647/GFP ratios of “blank” chambers without printed DNA spots included within each MITOMI experiment. The average standard deviation of these fit normal distributions was then used as the standard deviation of the additive normal noise (**Supplementary Fig. 68**).

We then fit Langmuir isotherms (**Eqn. 10**) to the resultant simulated binding curves by fitting a “global” *R*_max_ as described above for MITOMI experiments, with the following modifications. For each set of [*DNA_0_*] values and each of the 10 random seeds we performed separate fits over 60 samplings of 70% of datapoints from each binding curve. To mimic experiments exhibiting avid binding, we also separately fit Langmuir isotherms using “local” *R*_max_ fitting, in which the saturating AF647/GFP ratios are fit individually per binding interaction.

To determine thresholds beyond which fit *k*_d_ values should be considered either upper or lower limits, we compared the median fit *k*_d_ values across samplings to the “true” *k*_d_ values from which their binding curves were derived and set a conservative threshold of 1.4-fold difference (∼0.2 kcal/mol) between “true” and fit *k*_d_, expressing these *k*_d_ thresholds in terms of the fit *k*_d_ values (**Supplementary Fig. 67**). Using this method, the following lower and upper *k*_d_ thresholds were determined: 0.19 and 15.0 µM for global *R*_max_ fitting with 1 µM max [*DNA_0_*], 0.10 and 51.1 µM for global *R*_max_ fitting with 3 µM max [*DNA*_0_], and 0.13 and 18.9 µM for local *R*_max_ fitting with 3 µM max [*DNA_0_*].

### Calibration of ‘degenerate motif’ library ΔΔGs with ‘fiducial’ ΔG measurements

First, we converted MITOMI-derived *K*_d_s from 2 independent device replicates to ΔGs using the following equation:

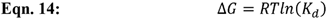

where *R* is the ideal gas constant (1.987 E-3 kcal/mol/K) and *T* is the temperature in degrees Kelvin (298°K). We noticed that the MITOMI measurements often had higher dynamic range for weakly bound sequences found within and near the ‘noise’ distribution of the BET-seq ΔΔGs, and broke the linear relationship between MITOMI ΔG and BET-seq ΔΔG. Thus, for each set of constructs with a given DBD (for example, Msn2-DBD, Msn2-FL, Msn2-DBD-Msn4-IDR, Msn2-DBD-Com2-IDR), we filtered out sequences with mean ΔΔG values within 3 standard deviations of the ‘noise’ distribution mean of the BET-seq ΔΔGs for any of the constructs within the group (shown in orange in **Fig. 2A**, **Supplementary Fig. 21**). Using the remaining sequences, we performed a linear regression of the BET-seq ΔΔGs against the MITOMI ΔGs with the following model:

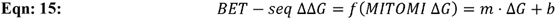

where *m* and *b* are the fit slope and y-intercept parameters respectively. We then used these parameters and the inverse of the **Eqn. 15** to calibrate the entire BET-seq ΔΔG datasets, transforming them into ΔG space for each construct.

### Autoinhibition constant fitting to calibrated ‘degenerate motif’ library ΔGs

To fit autoinhibition constants (*K*_auto_) to calibrated ΔGs, we began with the following equation related the observed *k*_d_ to the values of the *k*_d_ of a single binding site and *K*_auto_ (derived in **Supplementary Note 1**):

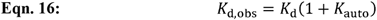

We then plugged this into our formula for ΔG (**Eqn. 14**) to get the following:

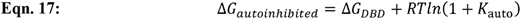

We fit *K*_auto_ values to the ‘degenerate motif’ library calibrated ΔGs of each DBD-IDR fusion by first randomly sampling 33% of sequences bound by either the DBD or DBD-IDR fusion (using a threshold of µ_noise_-3(std_noise_) to determine ‘bound’ sequences as described above) using the ΔGs of the DBD-only construct as the Δ*G_DBD_* variable in **Eqn. 17** and then fitting by minimizing residuals. We repeated this process 200 times to build a distribution of *K*_auto_ fits for each DBD-IDR fusion construct and estimate error in the fits. All reported *K*_auto_ fits are means of these distributions.

### Prediction of IDR radii of gyration

We predicted dimensions of Msn2, Msn4, and Com2 IDRs using ALBATROSS-colab^49^ using default settings.

### ‘Promoter’ library construction and characterization

We designed the ‘promoter’ BET-seq library by taking the 1000 base pairs (bp) upstream of all annotated ORFs in the R64 assembly of the S288C *S. cerevisiae* strain and tiling across them in 75 bp windows in 25 bp steps. To the 5’ and 3’ ends of these tiles we added a constant primer landing site and part of the Illumina TruSeq read 2 sequence respectively, yielding 213,487 unique sequences 117 bp long that we ordered as a SurePrint High Fidelity Oligo Pool (Agilent, G7278A) at 10 pmol scale (ordered library available at Zenodo^71^). We then PCR amplified the library using NEBNext Ultra II Q5 mastermix (New England Biolabs, M0544X) in twelve 100 µL reactions for 10 cycles with 7.8 ng of library as template, 1 µM of each primer (oMH158/159), 100 µM addition dNTPs, and a melting temperature of 65.5°C. Reactions were then split in two 50 µL aliquots and cleaned up using polyethylene glycol (PEG)-NaCl precipitation onto magnetic beads. To each aliquot, we added 2 µL of carboxylated speedbeads (Cytiva, 24152105050250) and 78 µL of a 20% PEG-8000/2.5 M NaCl mixture to achieve 12% PEG-8000 final. After mixing and incubating at room temperature for 10 minutes, beads were collected on a 96-side magnetic plate (Invitrogen, 12331D), supernatant was removed, and beads were washed two times with 150 µL of 80% DNase-free ethanol by resuspending beads in 80% ethanol, collecting beads, and removing the wash solution. After the final wash was removed, beads were collected on the magnetic plate and allowed to dry for 10-15 minutes (or until beads appeared cracked). Precipitated DNA was eluted from the beads in 5 µL of TT (10 mM Tris-HCl pH 8.0, 0.05% Tween-20) and elutions were pooled together. We quantified PCR yields by measuring A260 with a NanoDrop spectrophotometer and verified the size of the library (129 bp) by running the purified product alongside a 1 kb ladder (New England Biolabs, N0552L) on a 2% agarose gel buffered with 1x TAE and stained with 1x GelGreen. Library was diluted to 2 µM in TloE and stored at −20°C.

### BET-seq relative affinity measurements for ‘promoter’ DNA library

BET-seq measurements for the ‘promoter’ library were performed in the same way as for the ‘degenerate motif’ library with the following modifications. For device surface patterning of all experiments other than those varying TF density, we used a mixture of 100 µg/mL anti-eGFP VHH and 100 µg/mL bBSA in 1x PBS. In experiments varying TF-density, we used a constant 280 µg/mL of bBSA mixed with anti-eGFP VHH concentrations of 100, 16, 3, 2, 1, 0.75, or 0.66 µg/mL, flowed these varied mixtures into the device from block-specific inlets, and modified the patterning script in order to allow for multiple densities to be tested on the same device.

### BET-seq sequencing data processing for ‘promoter’ DNA library

Sequencing data from ‘promoter’ library measurements were processed in the same way as for the ‘degenerate motif’ library with the following modifications. We used cutadapt^74^ to trim adapters and remove constant sequences with the following parameters: cutadapt -j 5 -m 50 -q 18,18 --nextseq-trim=20 -a ^ AGATCGGAAGAGCACACGTCTGAACTCCAGTCAC… GTCATACCGCCGGACAAGAC. In case of sequencing errors in constant regions, we output untrimmed reads, removed 20 bp and 9 bp from the 5’ and 3’ ends respectively, and added these reads back to the trimmed reads. We then filtered these trimmed reads to ensure >90% of base calls with quality scores > 20, further trim low quality base calls from the 5’ and 3’ ends of reads, and remove reads smaller than 50 bp with the following settings: fastp -q 20 -u 10 --cut_front -- cut_front_window_size 1 --cut_front_mean_quality 20 --cut_tail --cut_tail_window_size 1 -- cut_tail_mean_quality 20 --length_required 50 --length_limit 80 -A. Trimmed and filtered reads were aligned to the sacCer3 (R64) reference genome using Bowtie2. Due to trimming of low-quality base calls on variable region ends, unique reads were then matched to designed promoter tiles using a python implementation of the ahocorasick algorithm (https://github.com/WojciechMula/pyahocorasick) to search for read matches within each promoter tile (as substrings within each 75 bp promoter tile string). Reads that were not either exact matches or substrings of a promoter tile (∼14% of all reads) were not used for subsequent analysis, since most of these reads were for sequences that either had low coverage (<10 reads per sequence) or were not observed in both the bound and unbound fractions, leaving only ∼1-2% of non-tile matching reads that could have been salvaged for further analysis (**Supplementary Fig. 69**).

Probabilities of observing each promoter tile and relative binding energies were calculated in the same manner as for the ‘degenerate motif’ library.

### BET-seq data centering and normalization for the ‘promoter’ library

BET-seq ΔΔGs for the ‘promoter’ library were centered and normalized in the same manner as for the ‘degenerate motif’ library with the following modifications. ΔΔGs were first centered on a single promoter tile sequence (TGCAATTACGTAATTTTAGCCGCTGAGAAGCGGATGGTAATGAGACAAGTTGATATCAAACAGATACATATTTAA) selected by virtue of having minimal differences between the median ΔΔG values for all protein constructs and replicates. ΔΔGs were normalized replicates by performing a linear regression against the ‘reference’ experiment using sequences with ΔΔG < µ_noise_-2(std_noise_) in both the reference experiment and lower dynamic range experiments and transforming ΔΔG values using the fit slope and intercept parameters. These normalized ΔΔG values were then used in replicate-to-replicate comparison plots.

We then calculated the mean normalized ΔΔGs between experimental replicates and recentered these values on the mean normalized ΔΔG of a sequence (TAATCCTTGGATTCTTAAAAGGTTATTAAAGTTCCGCACAAAGAACGCTTGGAAATCGCATTCATCAAAGAACAA) selected again for having minimal differences with median mean normalized ΔΔGs of all protein constructs. These recentered mean normalized ΔΔGs are those shown in plots comparing relative binding energies of different protein constructs and all other analyses use these values unless otherwise stated.

### Motif discovery on ‘promoter’ library AffinityNet models

We applied TF-MoDISco to interpretation scores to identify and categorize the motifs driving binding and differential binding within the promoter library. The TF-MoDISco algorithm extracted high-importance sequence windows as seqlets, and organized them into clusters representing distinct patterns. For this analysis, we utilized a window size of 128 and a maximum of 50,000 seqlets to run TF-MoDISco.

### Motif finding using FiNeMo for ‘promoter’ library AffinityNet models

To find contributing motif occurrences within ‘promoter’ library sequences for AffinityNet models, we used Fi-NeMo (https://github.com/kundajelab/Fi-NeMo) to call motifs on the DeepSHAP contribution scores generated previously with the global-lambda parameter set to 0.6. To find contributing consensus AGGGG and variants thereof, we searched for the top two Msn2 and Msn4 DBD TF-MoDISco CWMs and the single top Com2 DBD CWM (**Supplementary Fig. 34**) using their respective DBD models (**Fig. 3G, Supplementary Fig. 35**). To find contributing homoT motifs, we searched for the #9 Msn2-FL TF-MoDISco CWM (**Supplementary Fig. 34**) using the Msn2-FL model (**Fig. 3F, Supplementary Fig. 36**). When searching for two motifs, we filtered for non-overlapping motif instances when reporting total motif counts.

### eGFP fluorescence standard curve

PS1800 8-inlet devices were set up and patterned using 100 µg/mL anti-eGFP VHH and 100 µg/mL bBSA in 1x PBS as described above. We prepared 1 nM of eGFP protein (Chromotek, EGFP) in 1x PBS, 0.5% BSA, 0.05% Tween-20 and flowed three separate preparations of this onto the device from block-specific inlets for 10 minutes with button valves closed and then stopped flow and incubated for 15 minutes to ensure device walls were sufficiently blocked. To fully bind the equivalent of one chamber volume of 1 nM of eGFP onto a functionalized button surface in each chamber we (1) flowed on 1nM eGFP onto the device for 2 minutes from block-specific inlets with buttons closed, (2) stopped flow, closed sandwich valves, and opened button valves, (3) allowed eGFP to bind button surfaces for 30 minutes, (4) imaged the in the GFP channel (500 and 1000 ms exposures) to ensure no eGFP remained in solution, (5) closed buttons and washed out device with 1x PBS for 2 minutes, (6) stopped flow and imaged the device in the GFP channel (500 and 1000 ms exposures) to quantify bound eGFP. We then repeated these steps 35 times to quantify the successive binding of one chamber’s volume of 1 nM eGFP.

Device image rasters were stitched together as described above and processed by finding button locations using the 35^th^ binding step images as a reference and quantifying the summed eGFP fluorescence in both the button and the button annulus for each binding step in those button locations. To quantify the relationship between eGFP fluorescence and concentration, we first calculated the eGFP signal-to-noise for each button at each binding step by dividing the summed button fluorescence (signal) by the summed annulus fluorescence (noise). We used the signal-to-noise ratio as a metric since it is functionally equivalent to background-subtracted summed fluorescence intensities, but has the advantage that it is robust to differences in exposure times and overall excitation efficiency across microscopes. Assuming all eGFP molecules in solution were stably bound to the button surface after 30 minutes of binding in each step (due to the picomolar affinity of anti-eGFP VHH for eGFP), the fluorescence of each step represented binding of an additional chamber volume of 1nM eGFP to the button. We then performed linear regression of the eGFP signal-to-noise against the cumulative bound eGFP concentration at each step for each chamber, fitting to the following form:

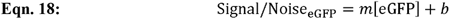

where *m* is the eGFP signal-to-noise ratio per nM of eGFP and *b* is the baseline eGFP signal-to-noise ratio with no eGFP bound, which is set to a value of 1. The mean slope values across all buttons were then taken for further calculations (**Supplementary Fig. 37**).

### Determination of TF surface density

To estimate the average distance between meGFP-tagged TF molecules immobilized on a device button surface, we first calculated the equivalent chamber concentration of eGFP using a rearrangement of **Eqn. 18:**

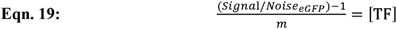

where *Signa*l/*Noise*_eGFP_ is calculated as above per button, *m* is the fitted slope from the eGFP fluorescence standard curve, and [TF] is the concentration of meGFP-tagged TF in nM. To calculate the density of TFs in terms of molecules per µm^2^, we used the following formula:

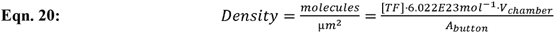

where [TF] is the concentration of TFs in units of mols per liter and *V_chamber_* and *A_button_*, the chamber volume and button area respectively, are defined by the following equations and known values for chamber and button geometry:

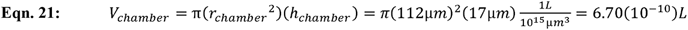

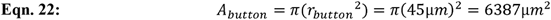

where *r* represents the radius of either the chamber or button and *h_chamber_* represents height of the cylindrical chamber. The value of 45 µm for the button radius here is a representative example, but is determined empirically on an experiment-by-experiment basis. We then plugged these values of chamber volume, button area, and TF concentration into **Eqn. 20** to calculate TF density for each button. We then convert TF density to average distance between TF molecules using the following equation (**Supplementary Fig. 37**):

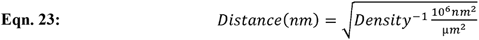

All reported average distances between TF molecules are the mean values of buttons with which the reported measurement was made. Reported errors are the standard deviations of those distances across buttons.

### Estimation of effective concentrations for avid binding models

Prediction of equilibrium dissociation constants under avid binding conditions requires estimation of the effective concentration (*C*_eff_) of a DNA binding site relative to nearby, immobilized TFs once another site on the same DNA molecule has been bound (**Figs. 4B, 4F**). To estimate the *C*_eff_ we used the following formula:

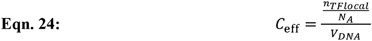

where *V_DNA_* is the volume accessible to a second DNA binding site once the DNA molecule has been bound by a single immobilized TF, *n_TFlocal_* is the number of adjacent surface-immobilized TFs within this volume, and *N_A_* is Avogadro’s number (6.022 x 10^23^ mol^-1^).

We estimated *V_DNA_* as a hemisphere whose radius is the sum of (1) the distance between the two binding sites in question and (2) the length of the flexible linker between the immobilized meGFP and the TF. Since the distance between the two binding sites depends on the sequence, for simplicity, we calculated the average distance between all AGGGG and AGGGG variant FiNeMo motif hit pairs called in the Msn2 DBD ‘promoter’ library AffinityNet model (8.74 nm, using a 0.34 nm per bp as a conversion factor). Once bound, the DNA can rotate and sway the length of the linker between meGFP and the TF, which, including the disordered residues at the end of Msn2 and the beginning of meGFP (GDIGGGSGGGGSGM), we predict to be 2.17 nm (end-to-end distance) using ALBATROSS-colab^49^. Thus, we estimate the radius of the hemisphere to be 10.91 nm, and calculate *V_DNA_* as follows:

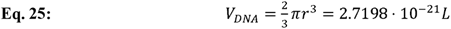

We then estimated *n_TFlocal_* as the average number of TF molecules expected within the hemisphere with a radius of 10.91 nm when drawn on a lattice. While the distance between TFs depends on the experiment, for library with DBD and FL (9.43 nm) as the distance between TFs in our lattice, which results in *n_TFlocal =_* 6. simplicity we used the average distance between TF molecules for all BET-seq measurements of the ‘promoter’ Inserting these values into **Eqn. 24** results in C*_eff_* = 3.66 mM.

### Simulation of avid binding with autoinhibition

Expected observed equilibrium dissociation constants for arbitrary numbers, *n*, of adjacent, immobilized TFs with autoinhibition binding to DNA sequences with *n* binding sites with identical *K*_d_ values were calculated using the general solution shown in **Fig. 4G**, derived in **Supplementary Note 1**, with varied values of *n* (1 - 5), *K*_d_ (0.05 – 1000 µM), *K*_auto_ = 0 or 1, and *C*_eff_ = 3.66 mM.

### ChEC-seq Data Processing

Raw ChEC-seq reads were downloaded using fastq-dump from the NCBI BioProject database (PRJNA573518 for Brodsky et al, 2020 and PRJNA993170 for Mahendrawada et al, 2025, accession numbers available in **Supplementary File 7**). Reads were trimmed of adapter sequences using cutadapt, removing the Illumina TruSeq Read2 (AGATCGGAAGAGCACACGTCTGAACTCCAGTCAC) from 3’ ends of single-end reads and TruSeq Read1 (AGATCGGAAGAGCGTCGTGTAGGGAAAGAGTGT) from the 3’ ends of read 2 of paired-end sequenced samples. Trimmed reads were aligned to the sacCer3 (R64) reference genome using Bowtie2 with the settings “-I 10 -X 700 --local --very-sensitive-local --no-unal” for single-end samples and with the additional settings “--no-mixed --no-discordant” for paired-end samples.

Initial sequence alignments in BAM format were pooled using samtools merge when replicates are available. The 5’ positions of the reads were extracted and separated by strand using bedtools genomecov −5 -bg -strand +/-. Finally, these strand-specific bedGraph files were converted into BigWig format using bedGraphToBigWig. Peaks were called on the merged BAM files using MACS3^80^ with model building disabled and parameters set to a shift of 0, an extension size of 20 bp, and summit calling enabled (--shift 0 --extsize 20 --nomodel --call- summits). Free MNase data was used as the control during peak calling for the data from Mahendrawada et al. 2025. No control was used for the data obtained from Brodsky et al. 2020. Outlier peaks were identified and removed using a threshold of 1.2 times the 99th percentile of the counts across all peaks.

### ChEC-seq model training

We utilized the bpnet-refactor repo (https://github.com/kundajelab/bpnet-refactor/tree/main) to train BPNet-style models on the ChEC-seq data. To account for the smaller genome size and fewer peak regions, we utilized an input window of 596 bp and an output window of 250 bp. The model processes one-hot encoded DNA sequences to simultaneously predict two values: total read counts within the 250 bp regions and the corresponding base-resolution probability profiles. The model was optimized using MSE as the loss function for the count head and Multinomial Negative Log-Likelihood (MNLL) Loss as the loss function for the profile head. The model architecture consists of an initial convolutional layer (64 filters, kernel size = 21) followed by a stack of six dilated convolutional layers (64 filters each, kernel size = 3). We used reverse complement augmentation and random jittering of up to 150bp during training. To mitigate overfitting, dropout (p = 0.15) was applied between convolutional layers, and an early stopping criterion with a patience of 30 epochs was implemented. All models were trained using five-fold cross-validation splitting across chromosomes. For the Brodsky et al. 2020 dataset, we excluded chromosome M for training and evaluation as the absence of experimental controls resulted in an excessive number of peaks on mitochondria.

### ChEC-seq model interpretation

To extract learned motifs, we first applied DeepLIFT to both the count and profile heads on peak regions. We then averaged these interpretations over five folds and used TF-MoDISco to cluster the interpretations into patterns for each head.

### Motif finding using FiNeMo for AffinityNet (‘promoter’ library) and ChEC-seq BPNet models

To find contributing motif occurrences within ‘promoter’ library sequences for ‘promoter’ library AffinityNet and BPNet models, we again used FiNeMo (https://github.com/kundajelab/Fi-NeMo with the global-lambda parameter set to 0.6. Specifically, we searched for the following CWMs using each BPNet model: AGGGG’ - #1 CWM from the Msn2-DBD ‘promoter’ library AffinityNet model **(Supplementary Fig. 34**); ‘AGGAG’ - #2 CWM from the Msn2-DBD ‘promoter’ library AffinityNet model (**Supplementary Fig. 34**); ‘CGCG’ - #1 CWM from the B, 2020 Msn2-IDR BPNet model (**Fig. 6D**); ‘homoT’ - #9 CWM from the Msn2-FL ‘promoter’ library AffinityNet model (**Supplementary Fig. 34**), the #5 CWM from the M, 2025 (30°C) Msn2-FL BPNet model, and the #2 CWM from the B, 2020 Msn2-FL BPNet model (**Fig. 6D**).

### AffinityNet and ChEC-seq model simulations

We first constructed 200 randomized background sequences 596 bp in length by dinucleotide shuffling the *S. cerevisiae* genomic region chrIV:1464250-1464846 (a non-peak region) and filtering out any shuffles containing the following sequences: GGGG, CCCC, AGGAG, CTCCT, AAAAAAA, and TTTTTTT. Motif sets (shown in **Fig. 6H** and **Supplementary Fig. 61**) were then embedded in the center of each background sequence and spaced equidistantly from 0 to 75 bp maximum between motifs, with a maximum sequence length of 129 bp including the motifs (the valid sequence length for AffinityNet ‘promoter’ library models). The central 129 bp of each background sequence with embedded motifs were then used as inputs for Affinity model prediction, while the whole 596 bp sequences were used as input for BPNet-style ChEC-seq models. Reported predictions were averaged first across all 5-folds for each model, then averaged across the 200 shuffled background sequences, and finally normalized to the mean predicted binding of the background sequences with an embedded E-box motif (CACGTG).

## Acknowledgements

This work was funded by an NSF CAREER Award (2142336) and an NIH Pioneer Award (DP1CA290563) to P.M.F., and an NSF graduate research fellowship (DGE-2146755) awarded to M.G.B.H. P.M.F. is a Chan Zuckerberg Biohub San Francisco Investigator. The authors thank Julia Zeitlinger, Jesse Engreitz, Connor Horton, Peter Suzuki, Renee Hastings, Craig Markin, Lucas Melo, and the members of the Fordyce lab for their helpful discussions.

## Author contributions

M.G.B.H. measured and analyzed transcription factor-DNA affinities in MITOMI and BET-seq microfluidic assays. Z.C. developed, trained, and tested NN models on BET-seq and genomic binding data. A.A.L. and M.G.B.H derived biochemical models of binding with autoinhibition and avidity. S.P. analyzed conservation of Msn2 and Msn4 orthologs and designed mutant constructs. S.L. designed microfluidic devices used in BET-seq experiments. M.G.B.H. fabricated microfluidic molds and devices. M.G.B.H. and P.M.F. conceived of the study. M.G.B.H. and P.M.F. wrote the manuscript with input from all authors. P.M.F., A.K., and A.S.H. supervised the project.

## Declaration of generative AI and AI-assisted technologies in the manuscript preparation process

During the preparation of this work, the authors used Claude to collect some initial references and help with initial drafting of some sections of the manuscript and used Gemini to assist with generating code for data visualization. After using these tools, the authors reviewed and edited the content as needed and take full responsibility for the content of the published article.

## Data availability

BET-seq library sequences are available at Zenodo^71^. Processed MITOMI and BET-seq data will be available at Zenodo. Raw BET-seq sequencing data will be available at the NCBI BioProject database. Due to the large size of imaging data, raw microscopy images are available upon request.

## List of Supplementary Files

Supplementary File 1: MITOMI concentration-dependent binding curves used to calibrate ‘degenerate motif’ library affinities for FL, DBD-only, and chimeric constructs of Msn2, Msn4, and Com2.

Supplementary File 2: MITOMI concentration-dependent binding curves for IDR-only constructs binding to selected ‘promoter’ library tiles.

Supplementary File 3: MITOMI concentration-dependent binding curves for Msn2-IDR binding to selected ‘promoter’ library tiles across varied surface patterning densities.

Supplementary File 4: MITOMI concentration-dependent binding curves for Msn2 and Msn4 mutants IDR constructs investigating impacts of di-arginine repeats on binding.

Supplementary File 5: Protein domain annotations, plasmids, and plasmid sequences

Supplementary File 6: Oligos use in MITOMI experiments

Supplementary File 7: ChEC-seq data accession numbers

Supplementary Note 1: Derivation of biochemical models of autoinhibition and avid binding

## Supplementary Figures

**Supplementary Figure 1.**
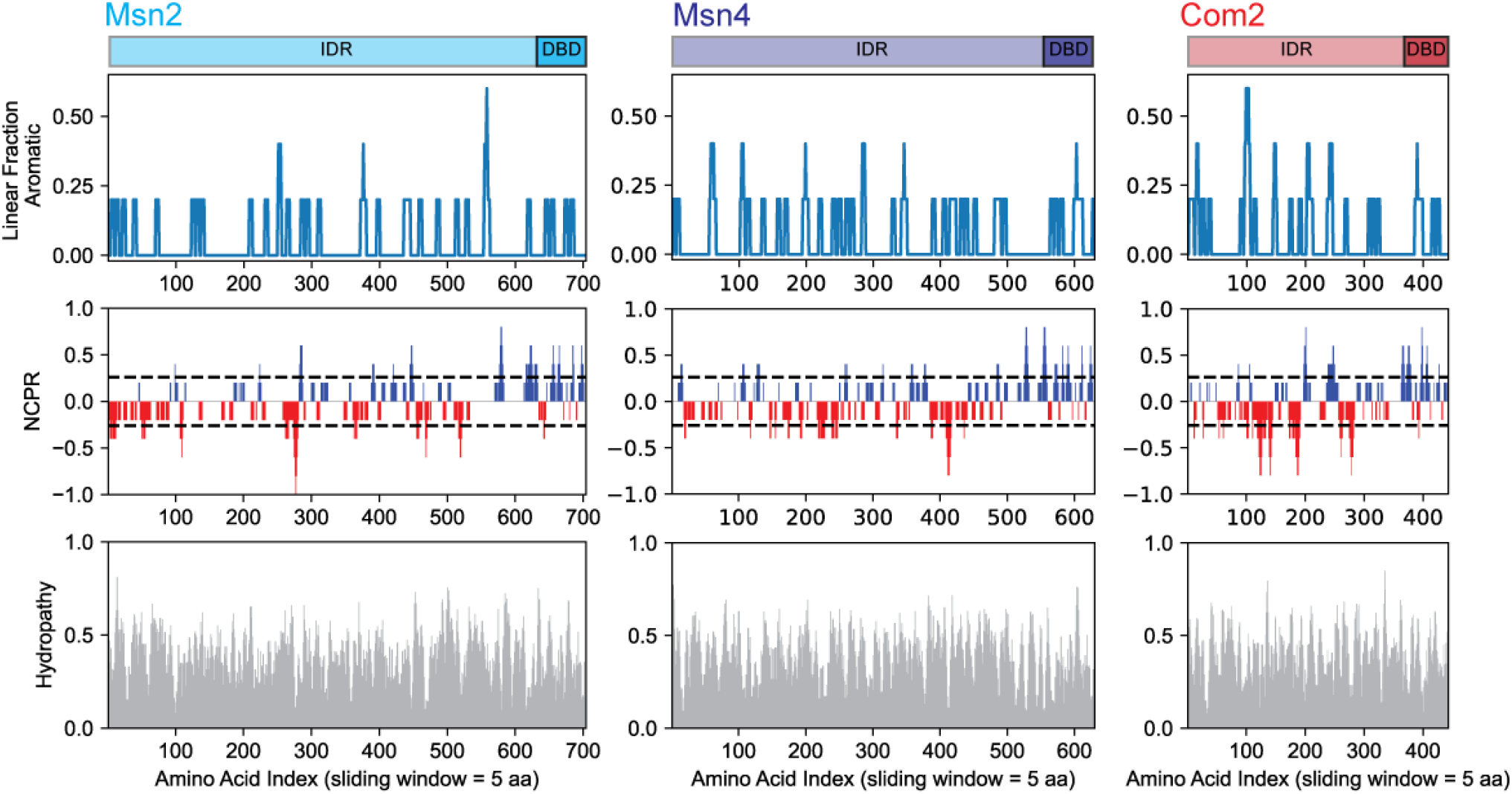
Amino acid properties of Msn2, Msn4, and Com2. Linear fraction aromatic (top), net charge per residue (NCPR, middle), and hydropathy (bottom) as a function of residue position within Msn2 (left), Msn4 (middle), and Com2 (right). We calculated all properties using a 5-amino-acid sliding window. For NCPR, dashed lines indicate values of +0.25 and −0.25, respectively; red and blue coloring indicates regions of net negative and net positive charge, respectively.

**Supplementary Figure 2.**
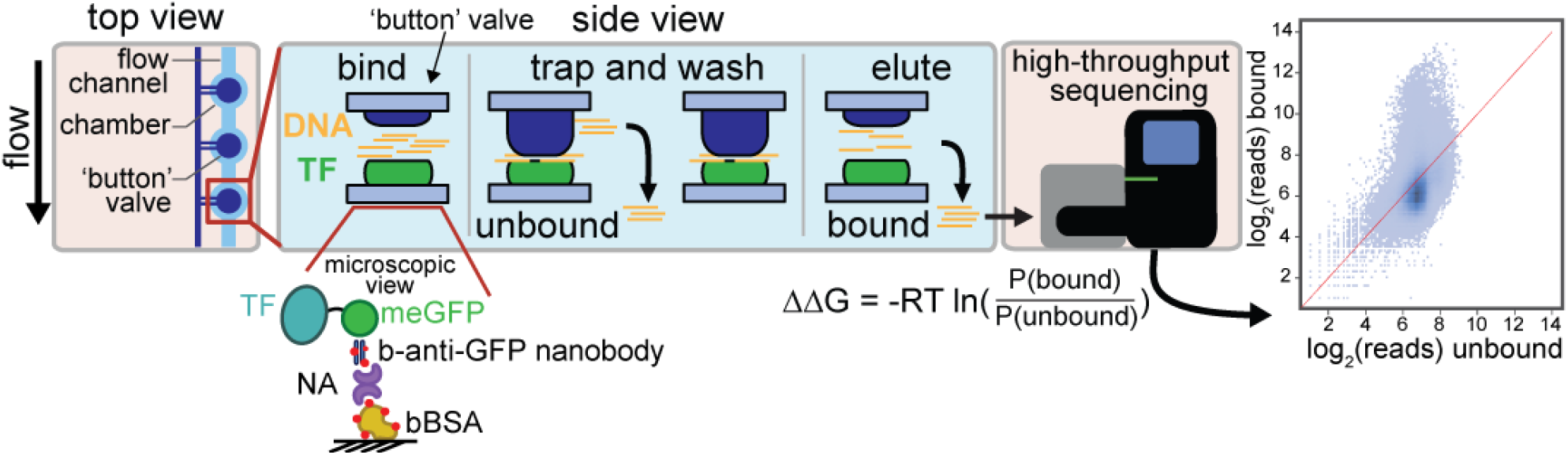
Schematic illustrating BET-seq experimental pipeline and returned data. Individual chambers along columns of flow paths within a microfluidic device each have a pneumatic “button” valve. Under each button valve, we pattern surfaces with biotinylated BSA, neutravidin, and biotinylated anti-GFP nanobody, which in turn makes it possible to recruit and immobilize meGFP-tagged TFs (see **Methods** for details). After surface patterning, we introduce a DNA library onto the chip and allow the library to bind to the surface-immobilized TF. After incubating to allow binding interactions to come to equilibrium, closing the “button” valve physically traps the bound fraction, making it possible to elute the unbound fraction from the device. Next, we wash and open the button valve to elute the bound fraction. Counting the number of molecules corresponding to each sequence in each fraction via Illumina sequencing makes it possible to compute a probability that each sequence is bound and extract relative binding energies.

**Supplementary Figure 3.**
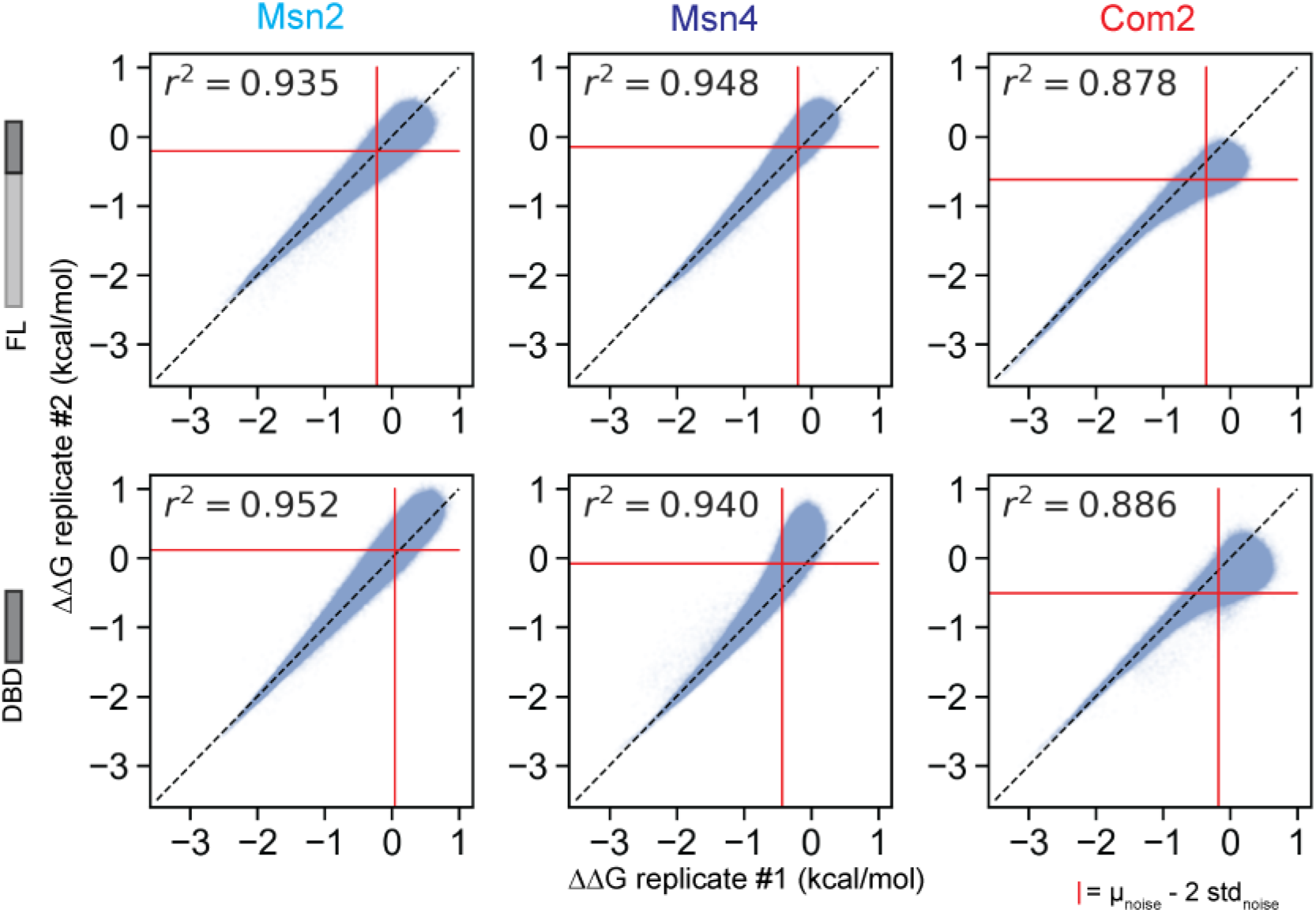
Scatter plots comparing measured ‘degenerate motif’ library ΔΔG values for each FL and DBD-only construct across 2 technical replicates. Blue markers denote measured ΔΔGs for each sequence (computed relative to CTTC-CGGGG-CCGG). Dotted black lines indicate 1:1 lines; red lines indicate thresholds of 2 standard deviations from noise mean values (calculated by fitting a Gaussian curve to an inferred ‘noise’ distribution of each experiment obtained by taking the ΔΔG values greater than the empirical center and their reflection across that center value – see **Methods**). Pearson r^2^ values shown were calculated from ΔΔG values for sequences lying outside of 2 standard deviations from the noise of either replicate.

**Supplementary Figure 4.**
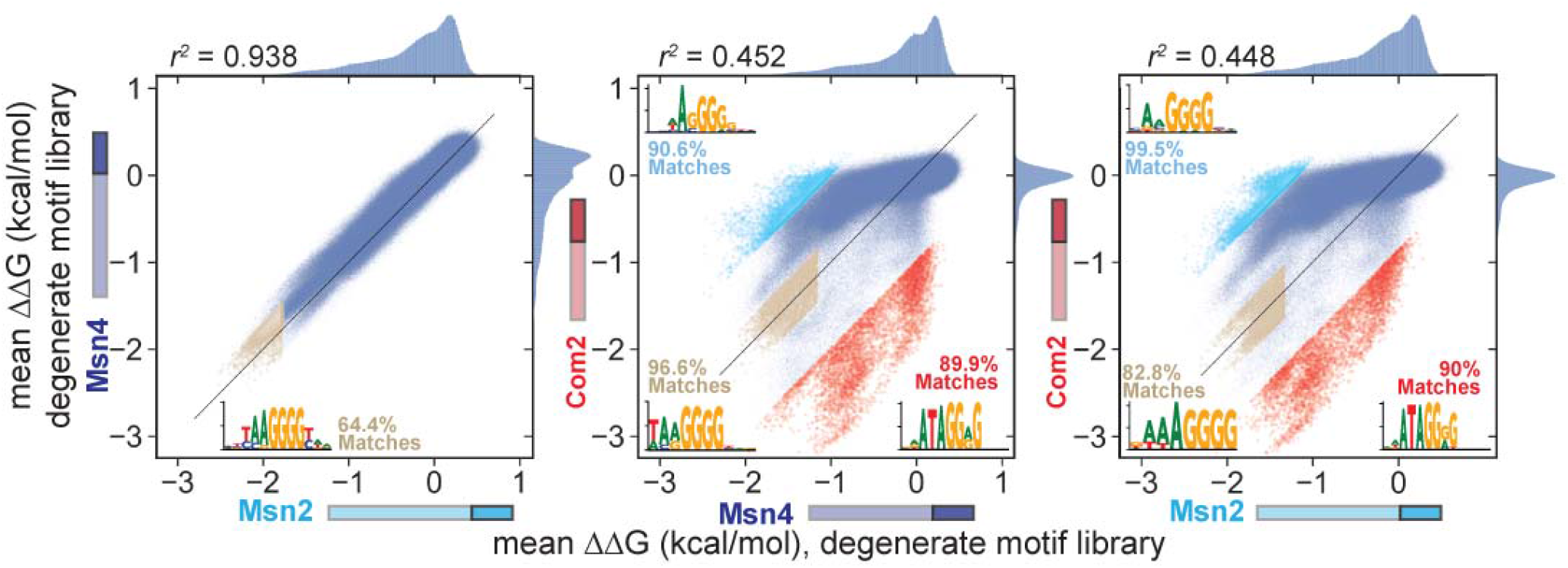
Scatter plots comparing measured ‘motif’ library ΔΔG values for FL Msn4, Msn2, and Com2. Scatter plots of measured ΔΔGs for FL Msn4 *vs.* Msn2 (left), Com2 *vs.* Msn4 (middle), and Com2 *vs.* Msn2 (right). Across all plots, each marker represents the ΔΔG value for a particular sequence (computed relative to CTTC-CGGGG-CCGG). In all plots, tan points represent the top 5000 sequences bound similarly between the compared TFs (with a maximal difference in ΔΔG of 0.3 kcal/mol); light blue and red points represent the top 5000 sequences preferentially bound by the TF on the y- or x-axis, respectively; and all other sequences are colored in blue. We then analyzed each sequence set using STREME to call *de novo* motifs with the percent of these sequence sets that match the top motif call indicated. For all 3 comparisons, we found either TA-AGGGG or AA-AGGGG motifs were highly enriched in the top similarly bound sequences. Msn2 and Msn4 showed no strong specificity differences. Com2 consistently preferred sequences enriched with an AGGGG or AGGAG motif with a strong preference for an upstream AT flank relative to Msn2 and Msn4, while Msn2 and Msn4 preferred sequences enriched for degenerate versions of the AGGGG motif, with variation at either the first or second positions.

**Supplementary Figure 5.**
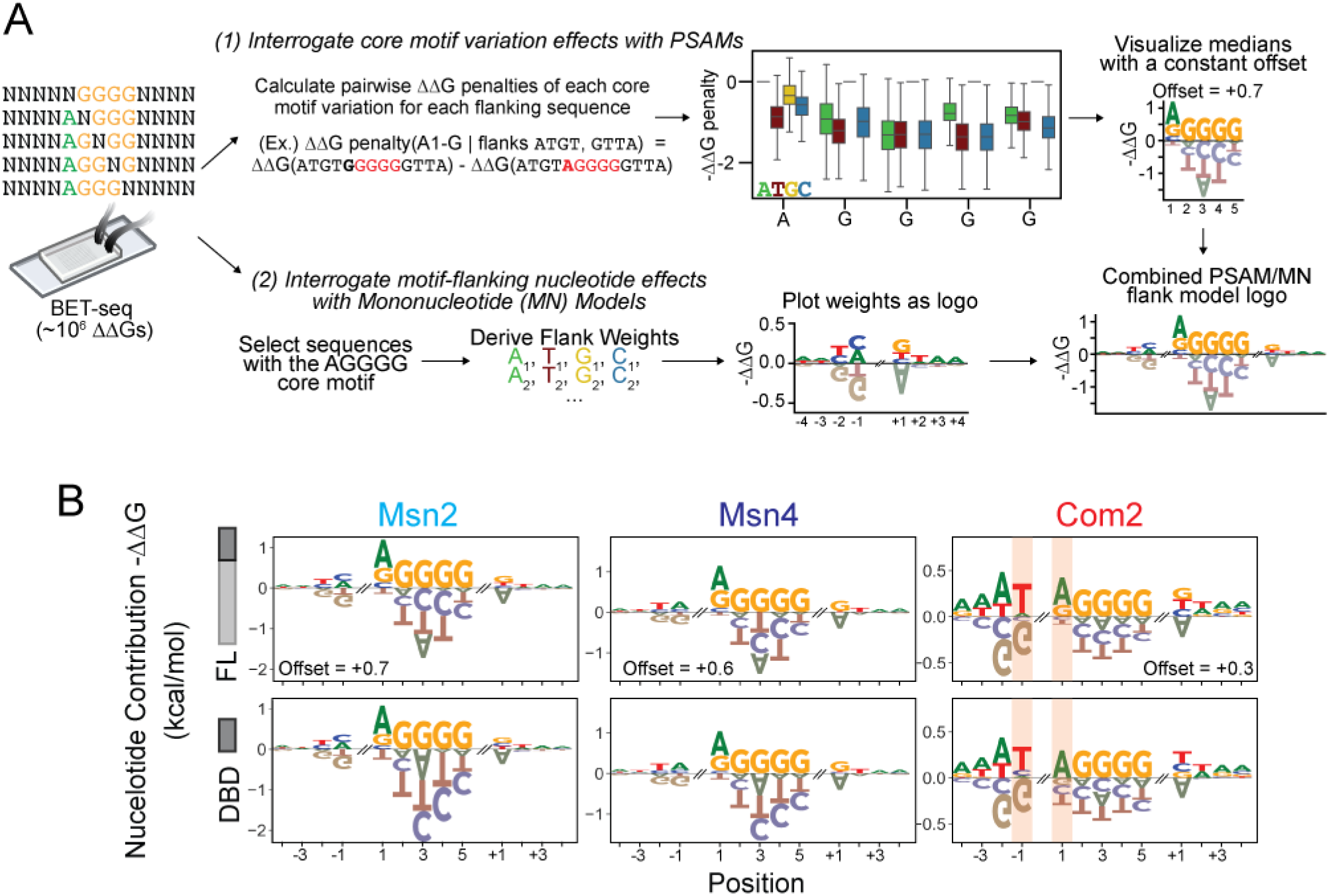
Analysis of ‘degenerate motif’ library BET-seq data using Position Specific Affinity Matrix (PSAM) and Linear Mononucleotide (MN) models. (**A**) Analysis pipeline used to construct composite PSAMs. First, we calculated pairwise ΔΔG penalties for each core motif variant relative to the consensus AGGGG motif for each unique flanking nucleotide combination in the library. We used the median ΔΔG penalty for each single nucleotide core variant and added a constant offset to all median penalties to create a logo representation of the core motif nucleotide preferences. Next, we constructed mononucleotide models of the flanks by selecting only sequences containing a consensus ‘AGGGG core’ and training a linear model with weights for each nucleotide at each flanking position. These weights represent energetic contributions of each flanking nucleotide and can be plotted as a logo. Finally, we visualized both PSAMs and MN flanking-nucleotide models together by plotting combined logos for each DBD and FL construct. (**B**) Returned logos for FL (top) and DBD-only (bottom) constructs for Msn2 (left), Msn4 (middle), and Com2 (right). Pink bars highlight positions exhibiting slightly different sequence preferences between Com2 FL and DBD-only constructs (DBD preference for C at the −1 flank and FL preference for G at core motif position 1).

**Supplementary Figure 6.**
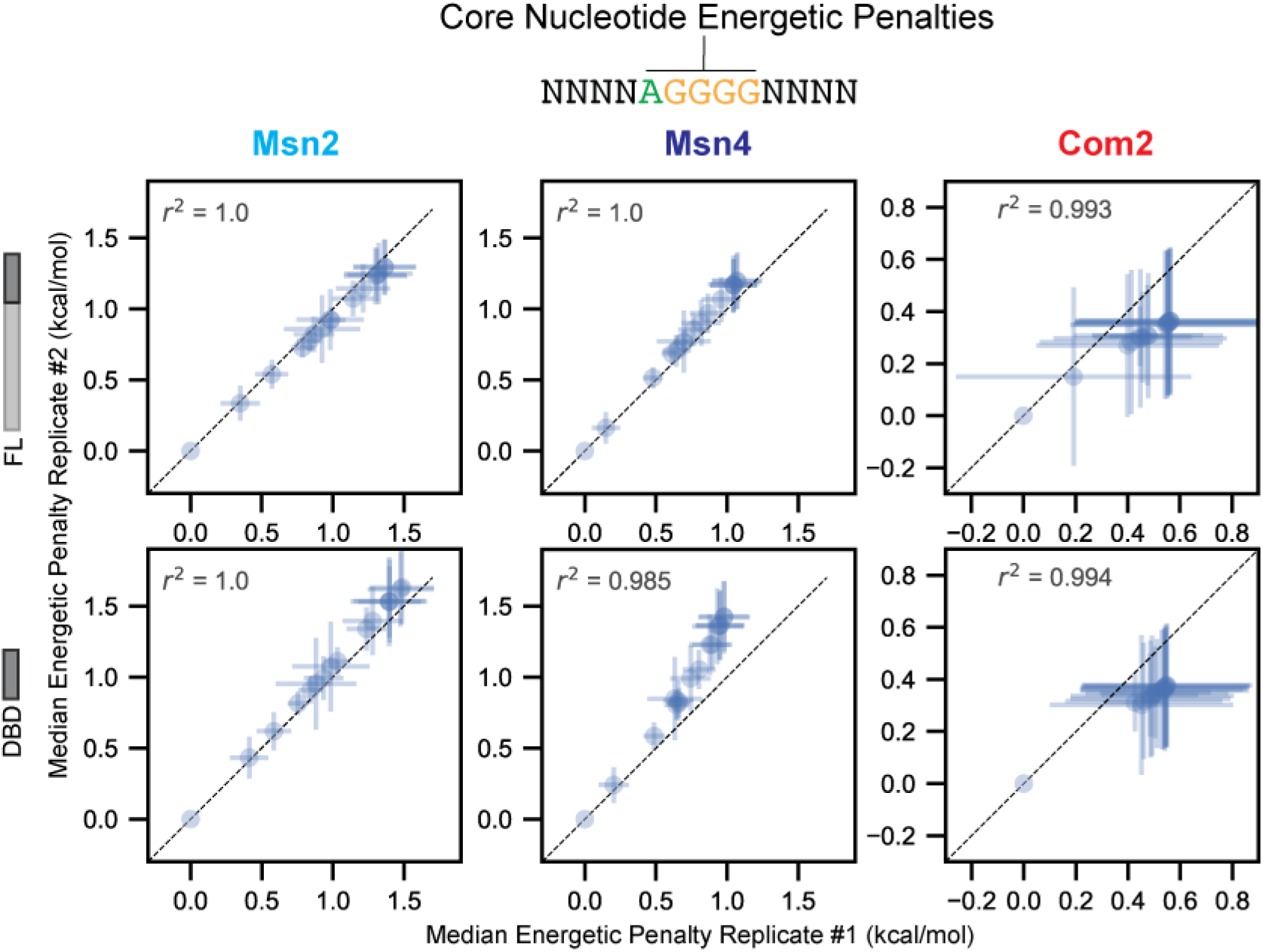
Scatter plots comparing PSAM median energetic penalties of core motif single nucleotide substitutions across two experimental replicates. Each point represents the median energetic penalty of a single-nucleotide substitution variant of the AGGGG consensus motif relative to AGGGG (as in **Supplementary Fig. 5**) for FL (top row) and DBD-only (bottom row) constructs of Msn2 (left column), Msn4 (middle column), and Com2 (right column). Points at 0 represent the consensus AGGGG motif. Dotted black lines represent 1:1 lines. Distribution medians were calculated for all sequences with a given core motif excluding those that contained alternative AGGGG motifs outside of the central five nucleotides. Error bars represent the variance of the energetic penalty distribution for a given substitution.

**Supplementary Figure 7.**
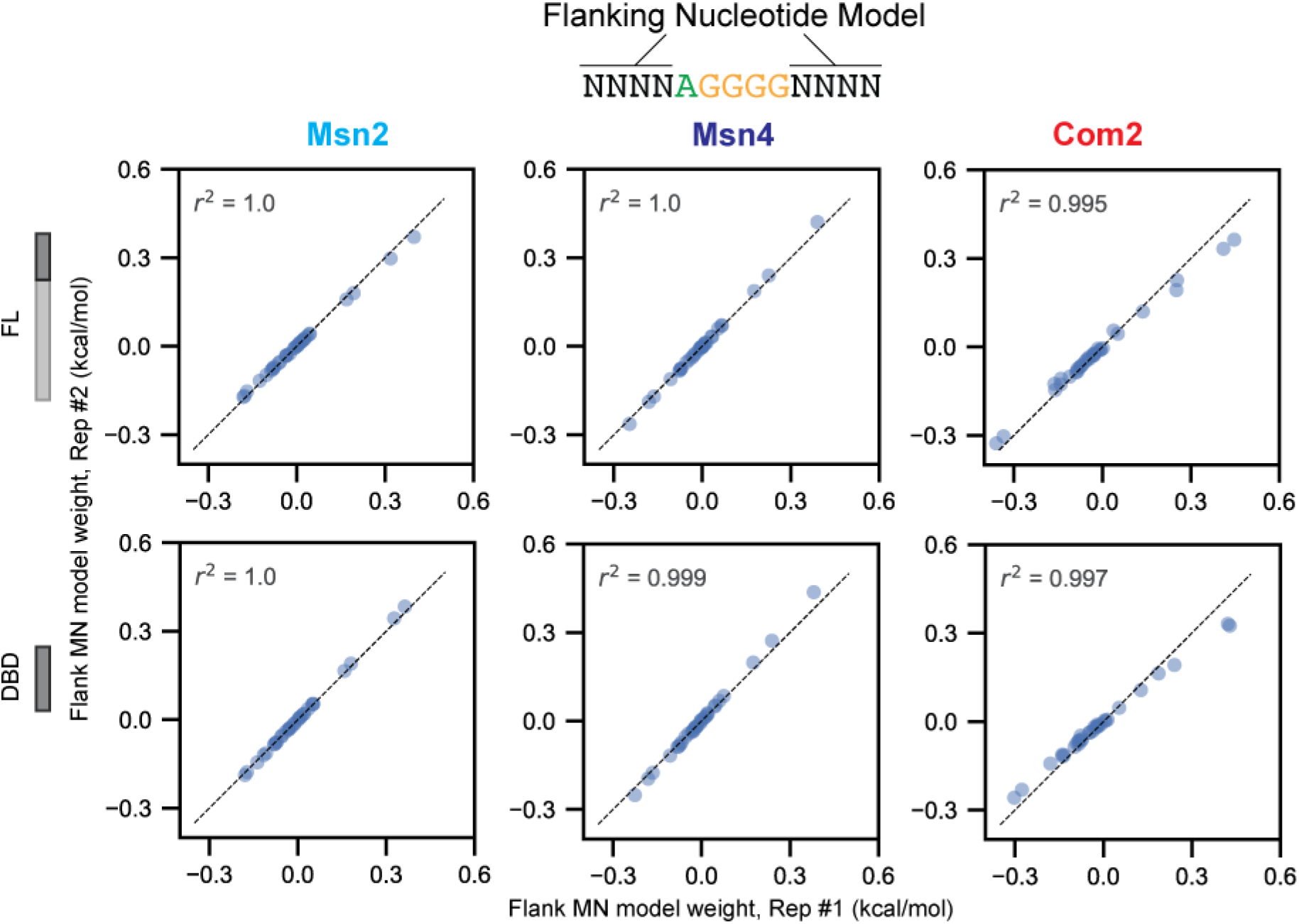
Scatter plots comparing mononucleotide model weights of motif-flanking nucleotides 4 bp up- and downstream of an AGGGG motif across two experimental replicates. Each point represents the model weight of a nucleotide at a position 4 base pairs up- or downstream of the core AGGGG motif for FL (top row) and DBD-only (bottom row) constructs of Msn2 (left column), Msn4 (middle column), and Com2 (right column). Dotted black lines represent 1:1 lines.

**Supplementary Figure 8.**
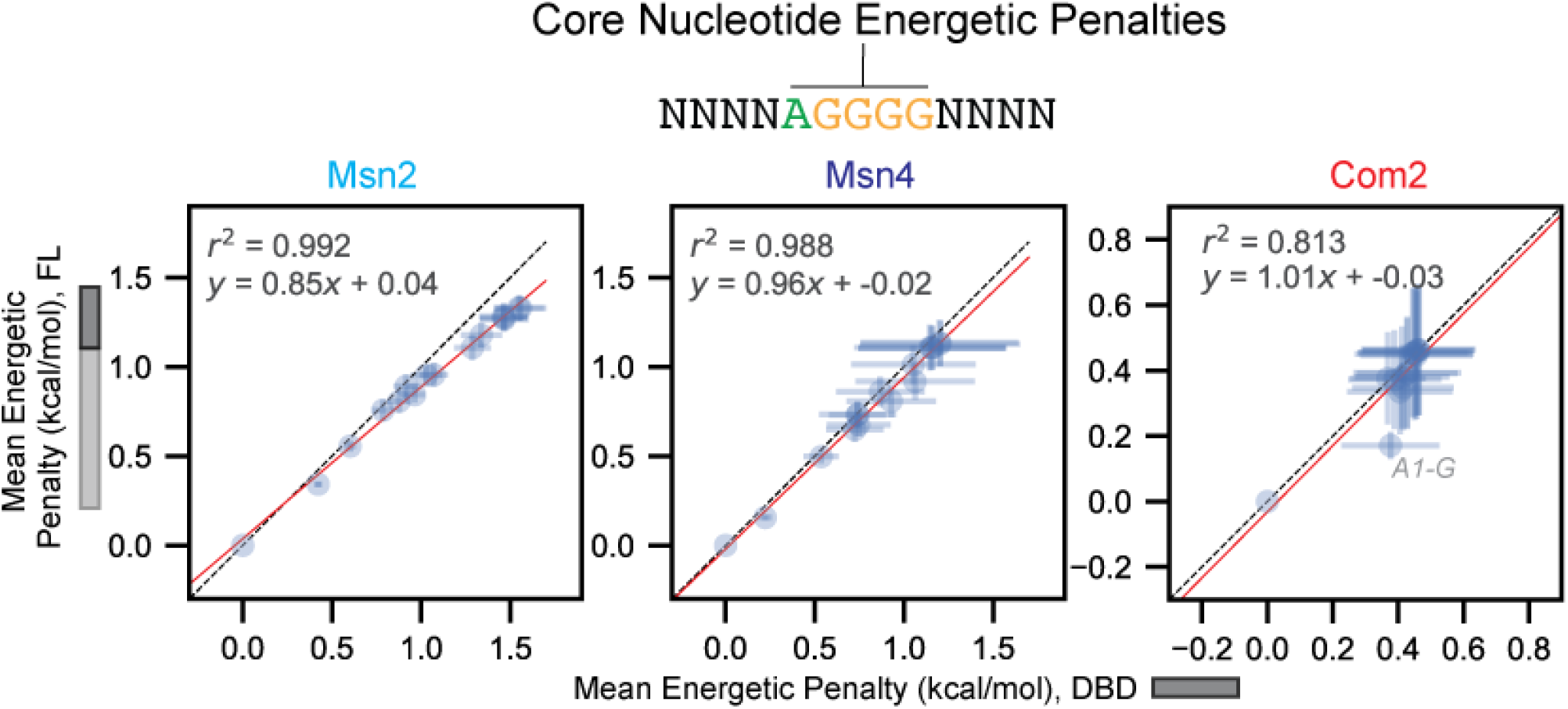
Scatter plots comparing mean energetic penalties for each core motif single nucleotide substitution between FL and DBD-only constructs. Blue markers represent mean energetic penalties of a single-nucleotide substitution variant of the AGGGG consensus motif relative to AGGGG for DBD (x-axis) and FL (y-axis) constructs of Msn2 (left), Msn4 (middle), and Com2 (right); error bars represent the standard deviation between the two replicates. Black dashed line indicates 1:1 line; red line indicates linear regression with the annotated fit parameters.

**Supplementary Figure 9.**
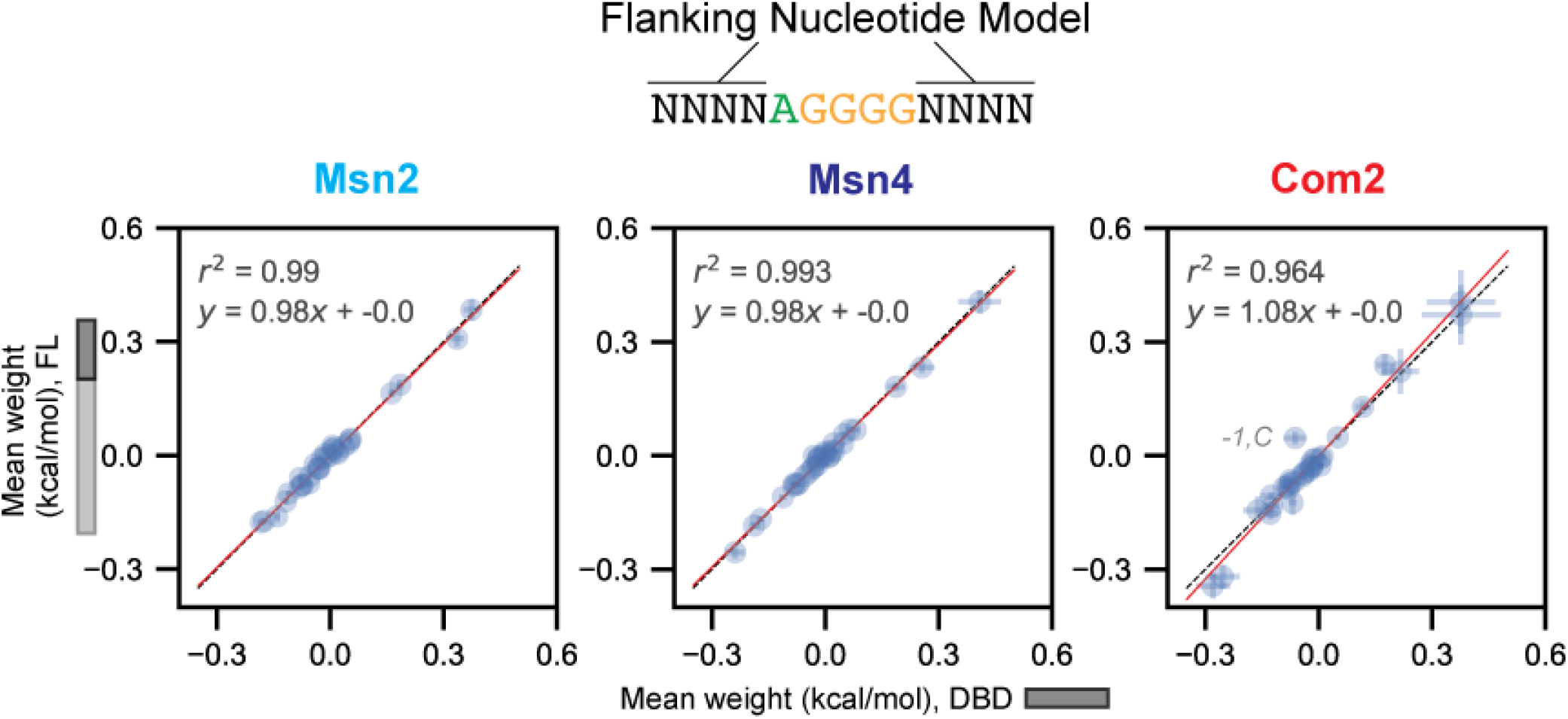
Scatter plots comparing mononucleotide flank model weights between FL and DBD-only constructs. Blue markers indicate mean flanking nucleotide weights for each nucleotide at each position 4 bp up- and downstream of an AGGGG motif for DBD vs FL constructs of Msn2 (left), Msn4 (middle), and Com2 (right). Error bars represent the standard deviation of the replicate weights. Black dashed line indicates 1:1 line; red line indicates linear regression with the annotated fit parameters.

**Supplementary Figure 10.**
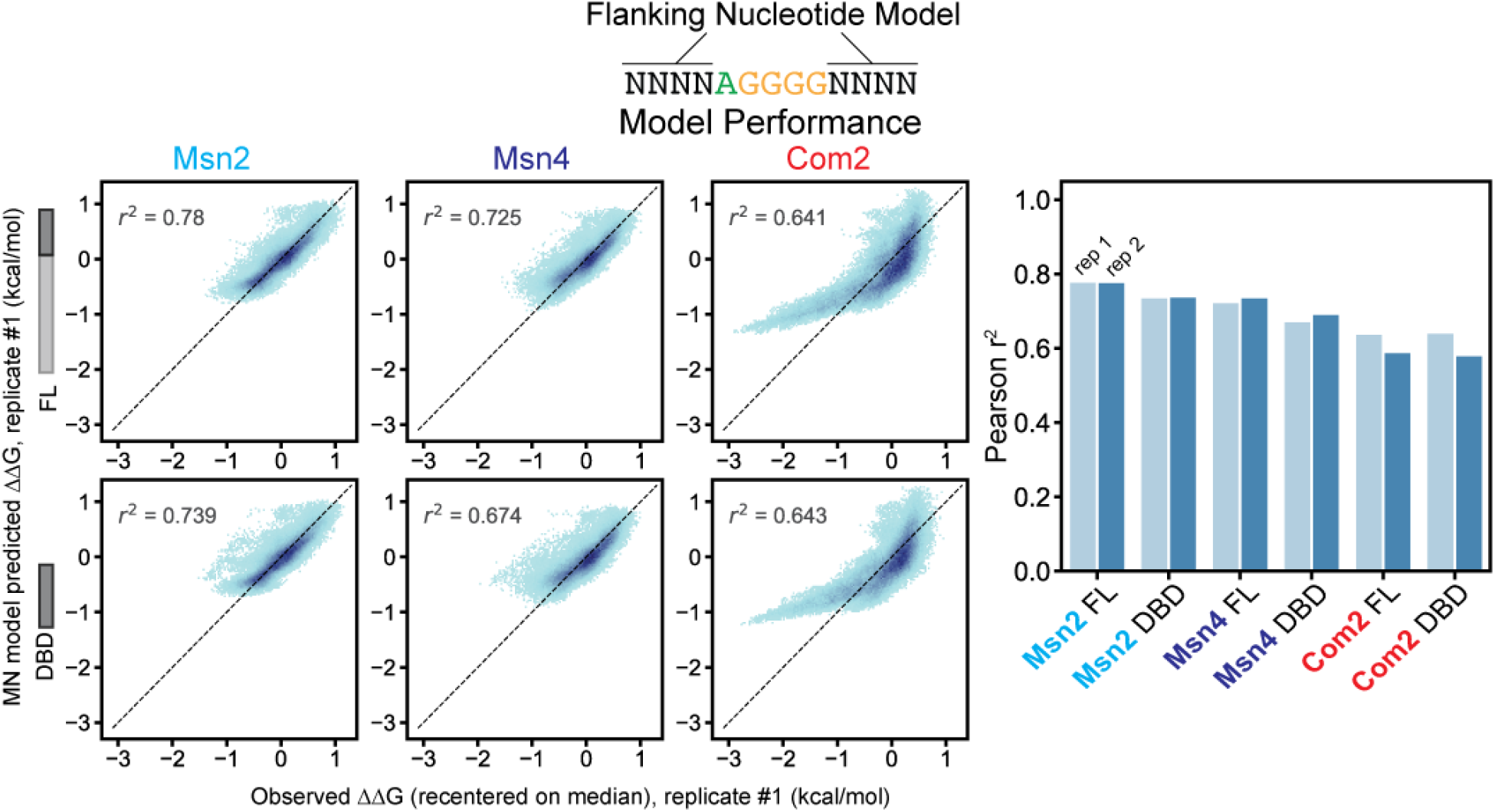
Comparisons between flanking nucleotide mononucleotide model (with AGGGG core) binding predictions and measured binding affinities. Left: 2D histograms comparing mononucleotide model-predicted ΔΔGs and observed ΔΔGs for FL (top row) and DBD-only (bottom row) constructs of Msn2 (left), Msn4 (middle), and Com2 (right) for experimental replicate #1. Sequences in the ‘degenerate motif’ library with AGGGG in the central five nucleotides were binned by observed and predicted ΔΔG and pixel darkness indicates density of sequences per bin. Black dashed line indicates 1:1 line; annotated Pearson r^2^ values are computed for all points. Right: Pearson r^2^ values for both experimental replicates of each construct.

**Supplementary Figure 11.**
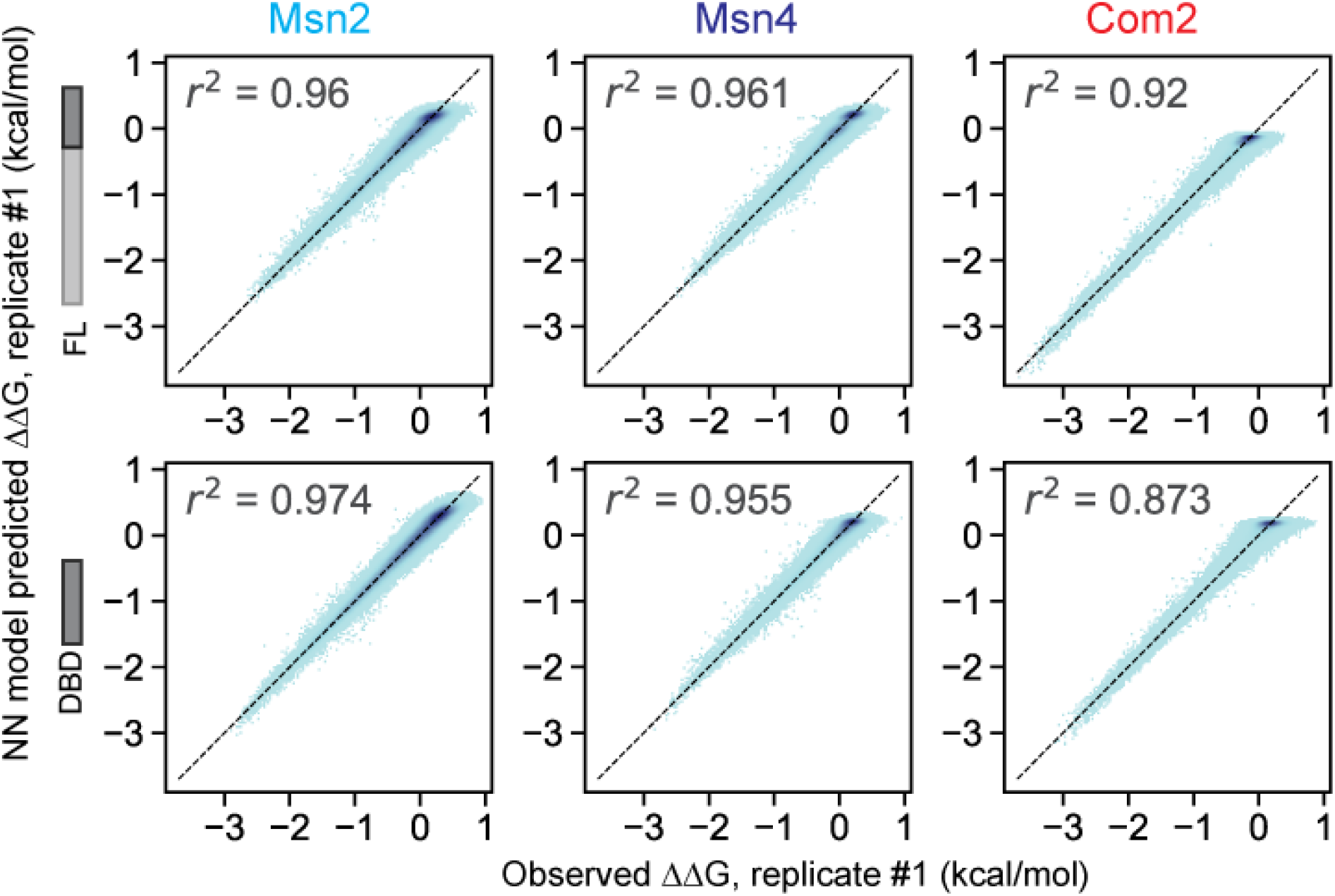
Neural network model performance predicting ‘degenerate motif’ library ΔΔG measurements. 2D histograms comparing AffinityNet model-predicted ΔΔGs versus observed ΔΔGs for replicate #1 of each FL (top row) and DBD-only (bottom row) constructs of Msn2 (left), Msn4 (middle), and Com2 (right). Sequences were binned by observed and predicted ΔΔG values and pixel darkness indicates density of sequences. Black dashed line indicates 1:1 line, annotated Pearson *r*^2^ values are computed for all points (Pearson *r*^2^ > 0.87 for all constructs).

**Supplementary Figure 12.**
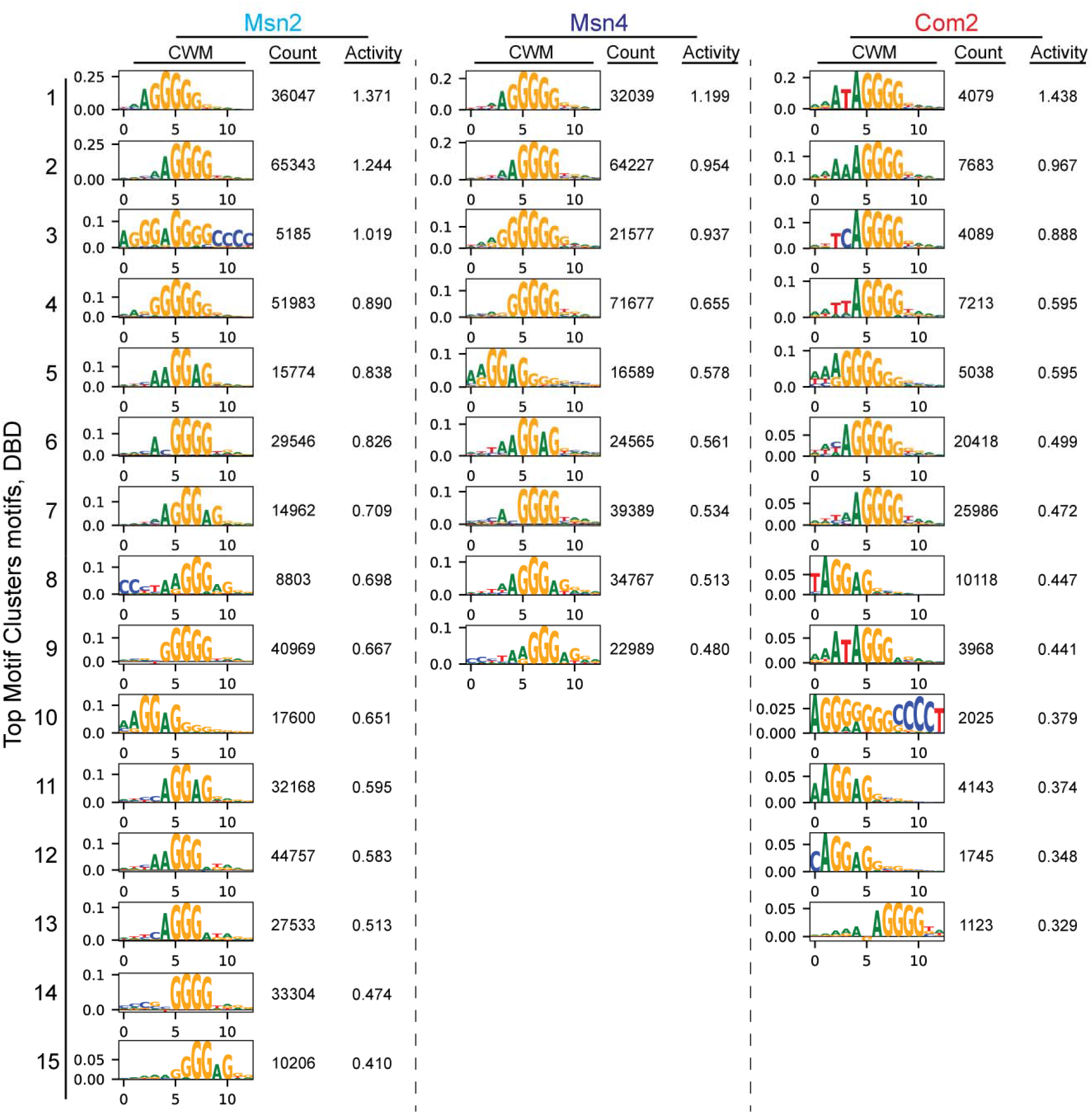
Top motif clusters derived from degenerate motif library AffinityNet models for each DBD-only TF construct ordered by decreasing activity. Count column values represent the number of library sequences assigned to a given motif cluster. Activity column values represent the summed contribution scores across the nucleotides of aggregated contribution-weight-matrices for a given motif cluster. Only motif clusters with summed predicted activity ≥ 0.2 are shown.

**Supplementary Figure 13.**
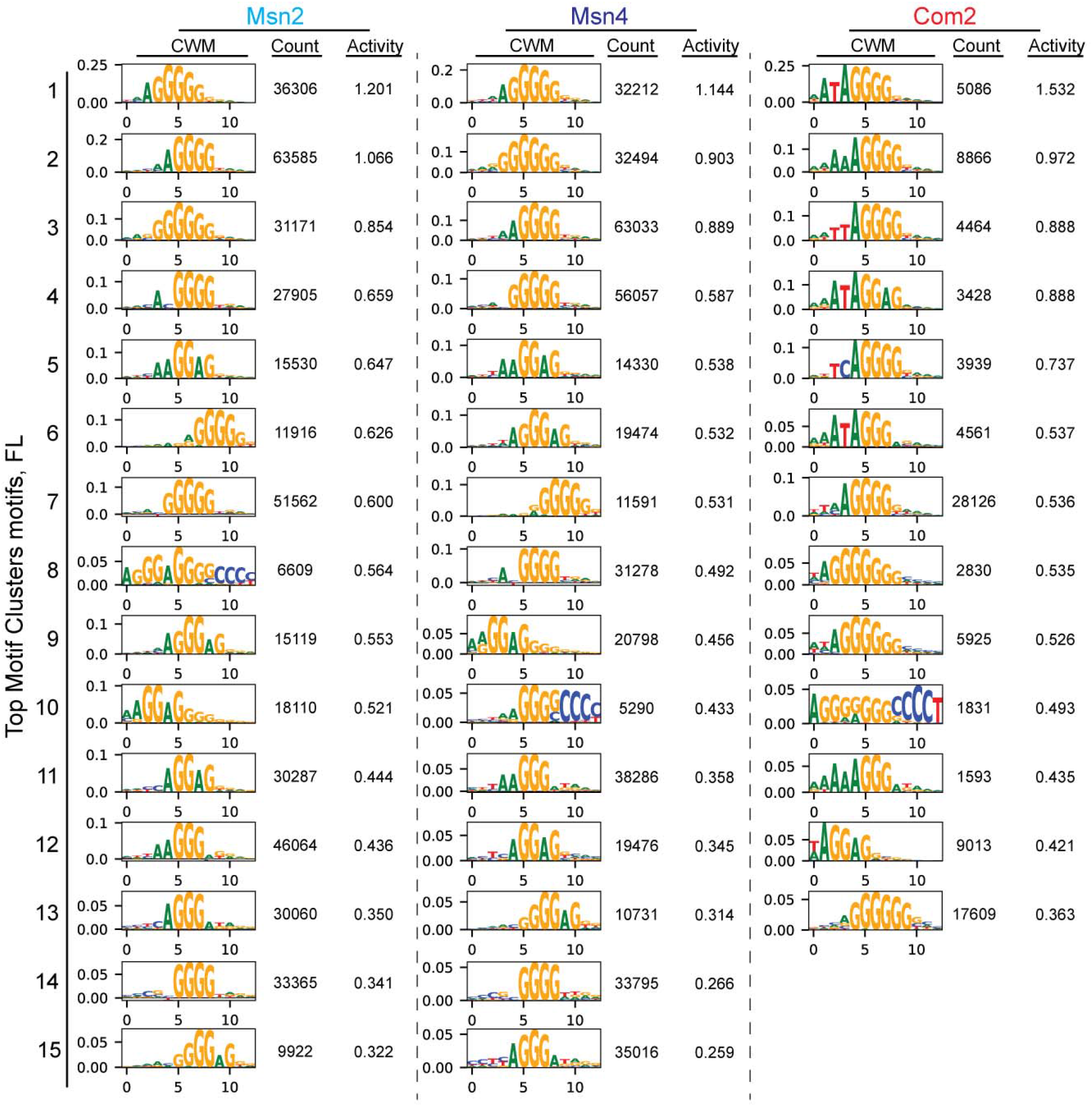
Top motif clusters derived from degenerate motif library AffinityNet models for each full-length TF construct ordered by decreasing activity. Count column values represent the number of library sequences assigned to a given motif cluster. Activity column values represent the summed contribution scores across the nucleotides of aggregated contribution-weight-matrices for a given motif cluster. Only motif clusters with summed predicted activity ≥ 0.2 are shown.

**Supplementary Figure 14.**
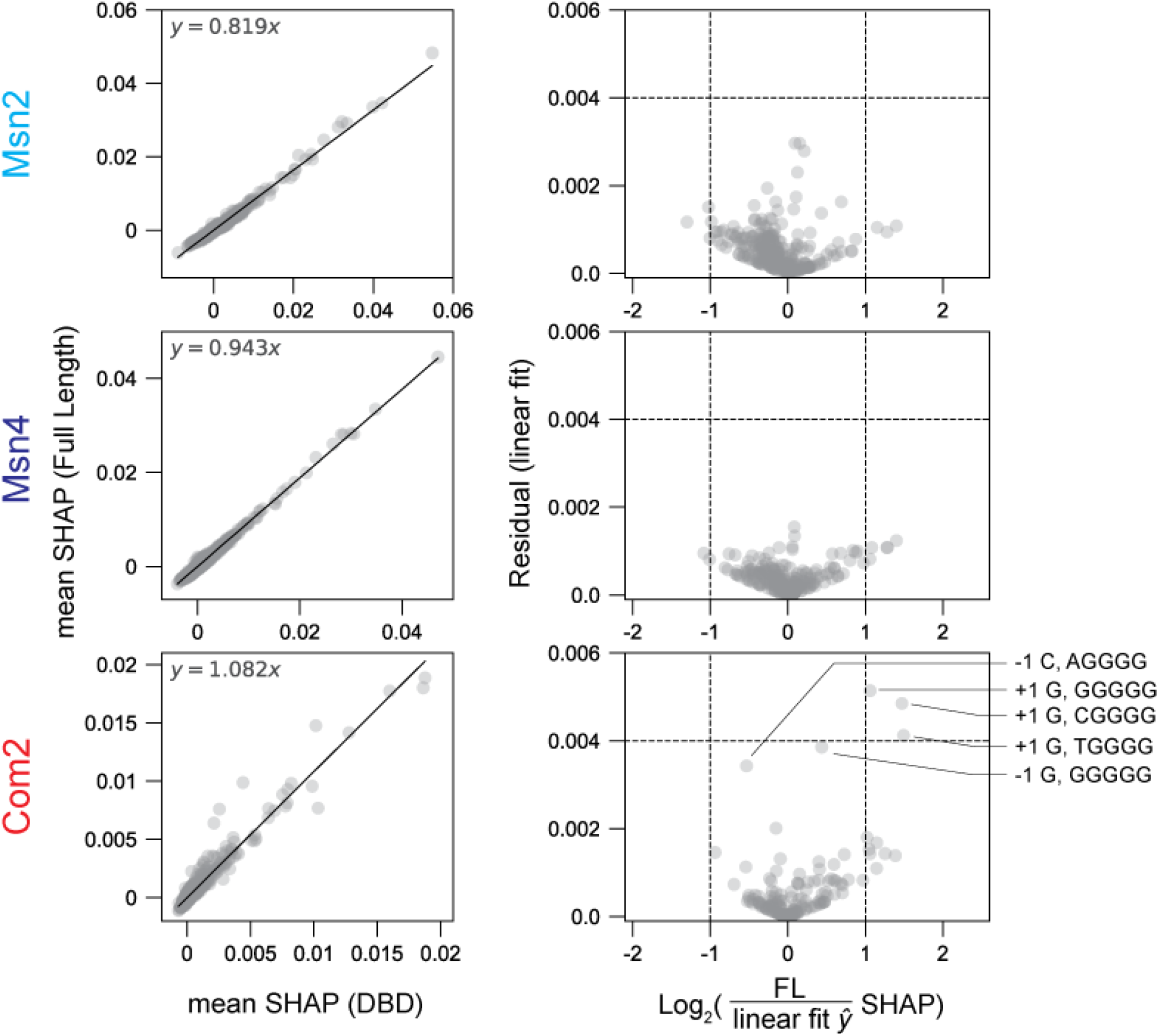
Analysis of motif-flanking nucleotide contributions to binding for the consensus AGGGG motif and all single nucleotide variants of the consensus motif using NN SHAP scores. Left: Scatter plots showing mean DBD versus FL construct mean SHAP scores of nucleotides in each motif flanking position 4 bp up- and downstream of an AGGGG motif or each single nucleotide variant of the AGGGG motif for Msn2 (top), Msn4 (middle), and Com2 (bottom). Black lines indicate linear fits with the annotated fit parameters. Right: Residual from linear fits versus log_2_-transformed ratio of the FL to linear fit-predicted SHAP scores for all possible flanking nucleotide-core motif combinations. To account for noise within low magnitude SHAP scores and limit the error of the FL/fit prediction SHAP ratio, all SHAP scores below 0.0008 were brought to a value of 0.0008 for calculation of ratios. Annotations identify specific sequence combinations with high residuals and fold changes between measured and predicted SHAP scores. Dotted lines indicate thresholds for fold change ≥ 2 and residual ≥ 0.004.

**Supplementary Figure 15.**
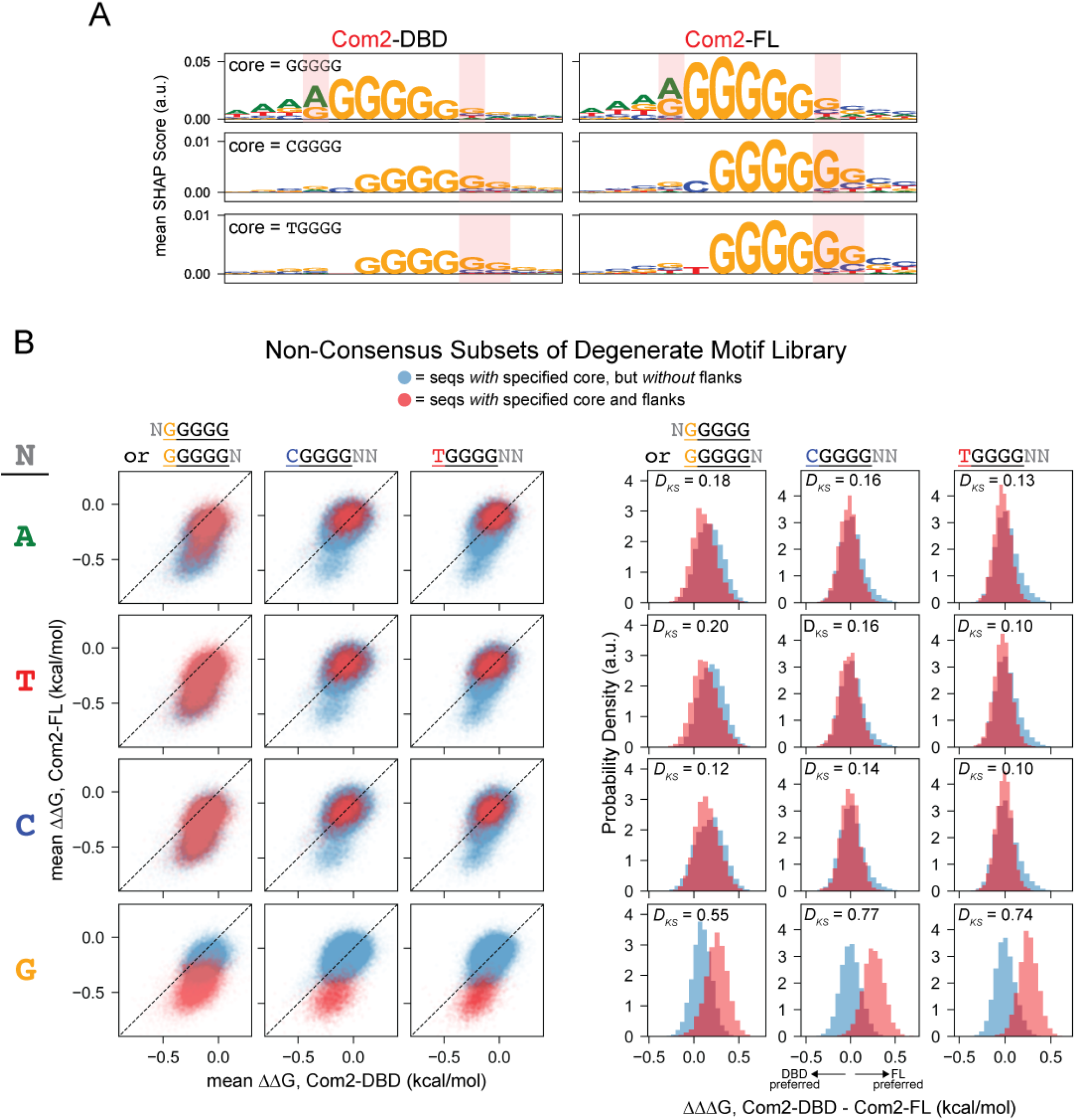
Com2-FL exhibits subtle preferences for 6xG motifs relative to Com2-DBD. (**A**) Logo plots of mean SHAP scores for core and flanking nucleotides 4 bp up- and downstream of either GGGGG, CGGGG, or TGGGG core motif variants; red boxes highlight flank positions (identified in **Supplementary Fig. 14**) that differ between DBD and FL constructs. (**B**) Left: Scatter plots comparing BET-seq-derived mean ΔΔGs for Com2-FL *versus* Com2-DBD for subsets of the degenerate motif library. Red points indicate sequences with both the specified core and flanking nucleotides (columns and rows respectively); light blue points indicate sequences with the specified core nucleotides but not the specified flanks. Right: Histograms of the differences in ΔΔGs between Com2-DBD and Com2-FL (ΔΔΔG) for the same sets of core and flanking nucleotides (same color-coding). Annotations denote Kolmogorov-Smirnov test statistics (*D_KS_*) for each comparison of cores with and without specified flanking nucleotides (*p* < 1E-30 for all test statistics).

**Supplementary Figure 16.**
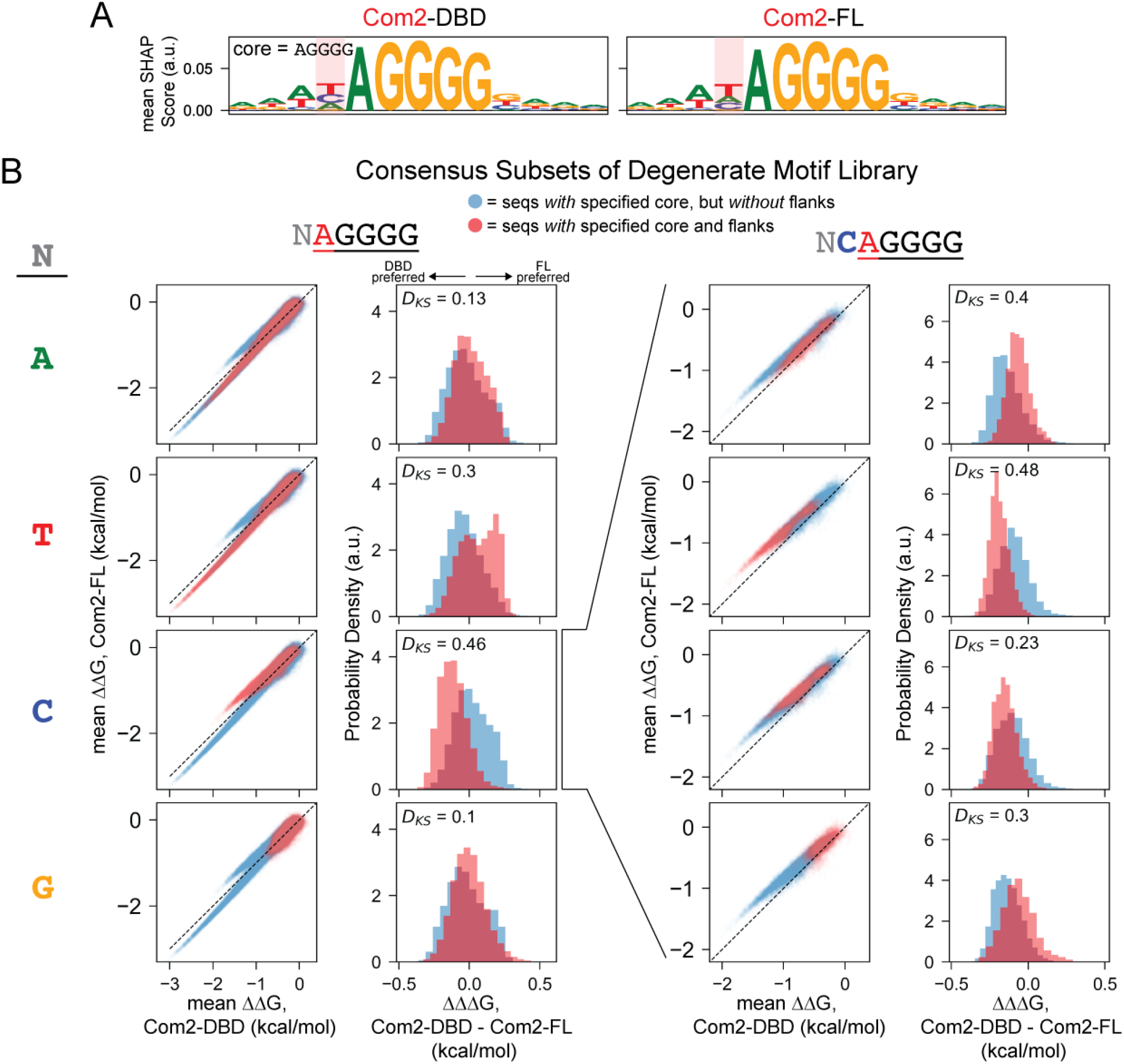
Com2-DBD exhibits subtle preferences for the TCAGGGG motif relative to Com2-FL. (**A**) Logo plots of mean SHAP scores of core and flanking nucleotides 4 bp up- and downstream of the AGGGG core motif; red boxes highlight positions that differ between DBD and FL constructs. (**B**) Scatter plots comparing BET-seq-derived mean ΔΔGs for Com2-FL versus Com2-DBD for subsets of the degenerate motif library; red points indicate sequences with both the specified core and flanking nucleotides (columns and rows, respectively) and light blue points indicate sequences with the specified core nucleotides but not the specified flanks. Right: Histograms of the differences in ΔΔGs between Com2-DBD and Com2-FL (ΔΔΔG) for the same sets of core and flanking nucleotides (same color-coding). Annotations denote Kolmogorov-Smirnov test statistics (*D_KS_*) for each comparison of cores with and without specified flanking nucleotides (*p* < 1E-30 for all test statistics).

**Supplementary Figure 17.**
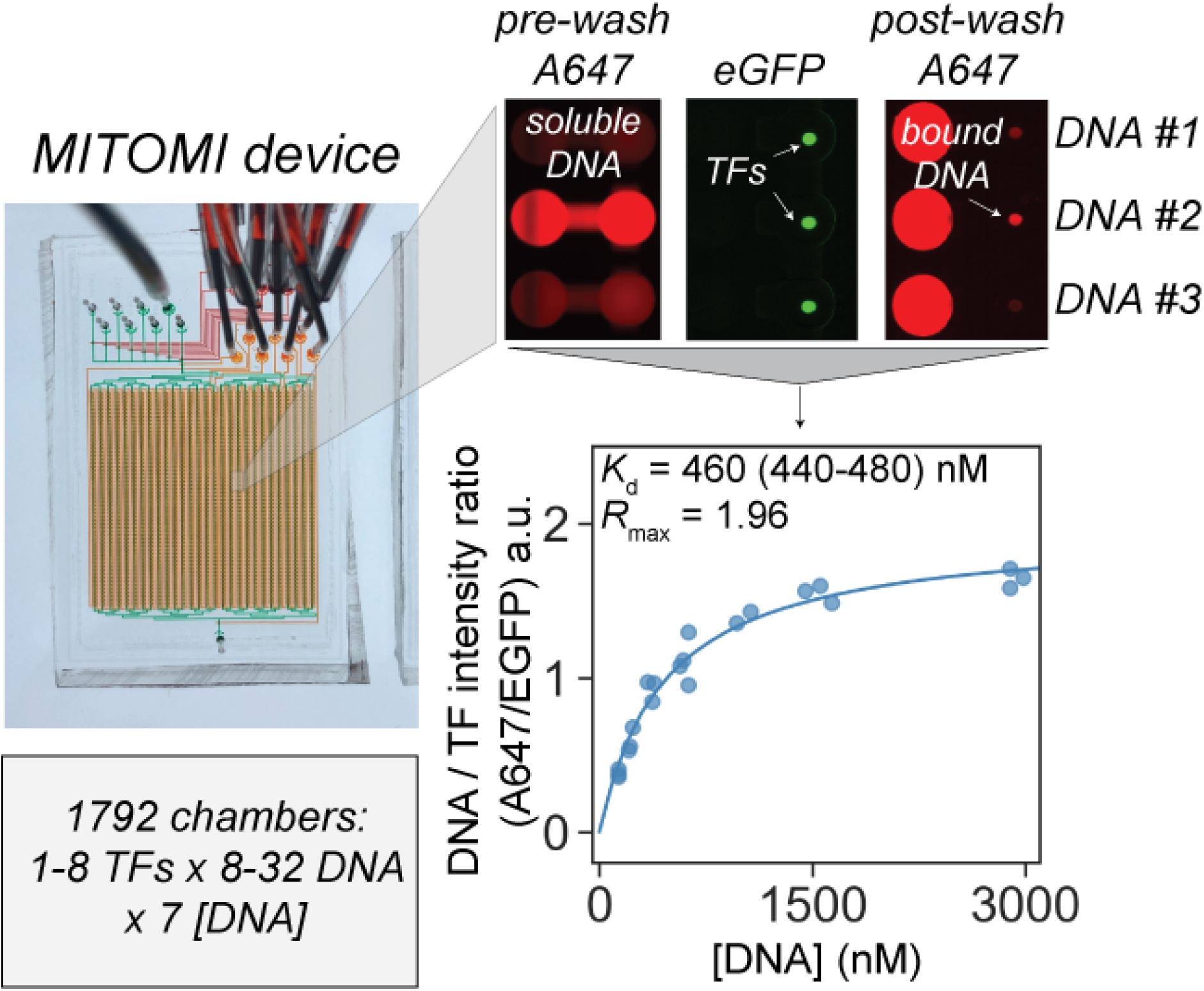
Schematic of MITOMI binding assay and representative images, binding curves, and fits to quantify *K_d_*. Left: Representative image of a MITOMI device with flow channels filled with green dye and control valves filled with red dye. Top Right: Zoomed-in representative fluorescence images in either the Cy5 (red) or GFP (green) channels showing solubilized Alexafluor647-(A647)-labelled DNA in chambers exposed to immobilized TFs (‘pre-wash’, left), immobilized meGFP-tagged TFs under button valves (middle), and TF-bound A647-labelled DNA after closing button valves to trap bound DNA and washing away unbound DNA (‘post-wash’, right). Bottom right: Representative measured ‘post-wash’ ratios of bound DNA to TF fluorescence intensities as a function of ‘pre-wash’ solubilized DNA concentrations. Blue markers indicate the median pixel intensity ratio for the circular region subtended by surface-immobilized GFP-tagged TFs within a given chamber; line indicates fit to a Langmuir isotherm with the returned annotated maximum occupancy and affinity values (with interquartile range of *K*_d_ fits annotated) (see **Methods**).

**Supplementary Figure 18.**
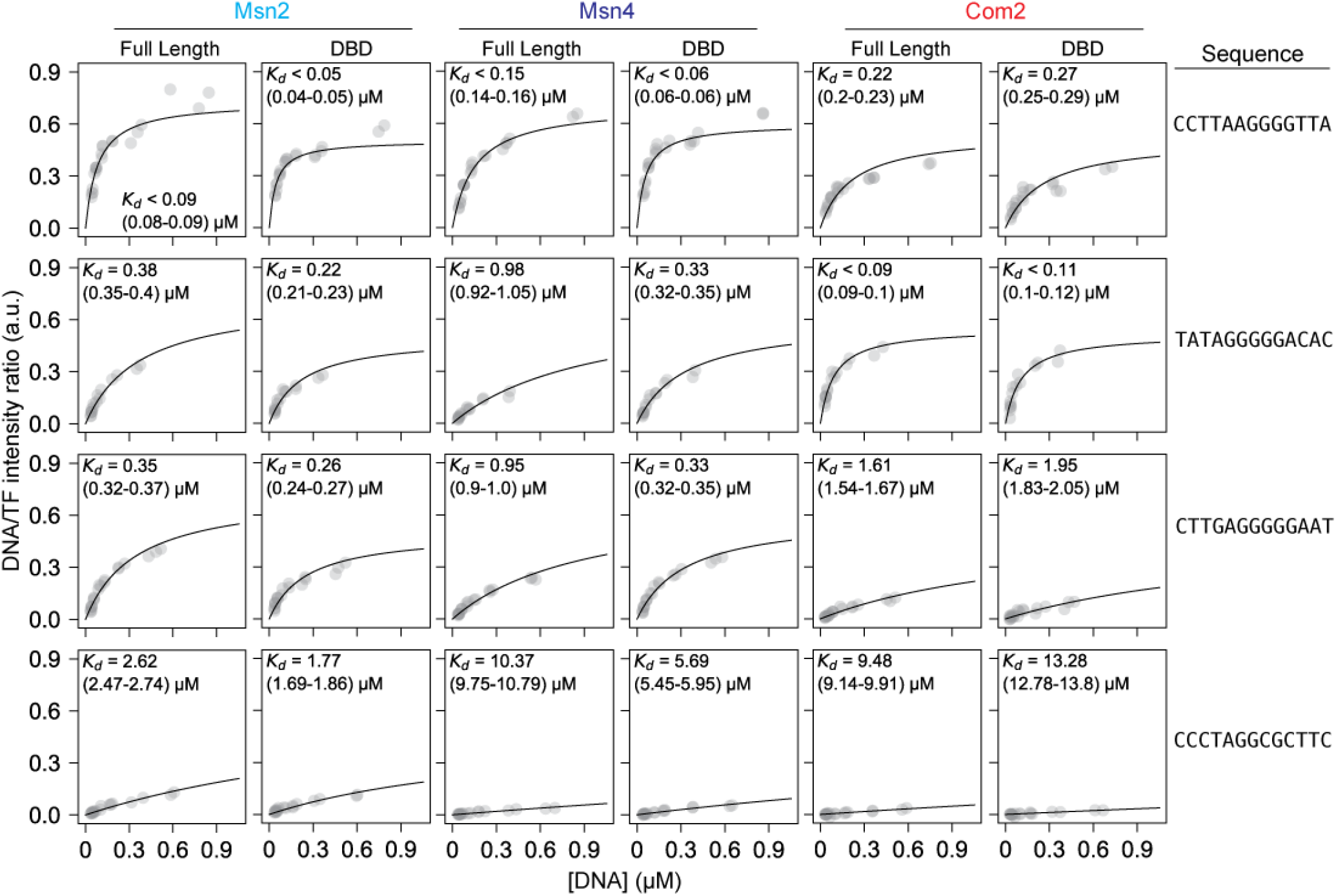
Representative TF-MITOMI concentration-dependent binding curves for FL and DBD constructs binding fiducial sequences from the ‘degenerate motif’ library. Measured fluorescence intensity ratios (Cy5/GFP or DNA/TF) as a function of soluble DNA concentration available for binding for FL and DBD-only Msn2, Msn4, and Com2 constructs interacting with representative fiducial sequences from the ‘degenerate motif’ library. Gray markers indicate per-chamber ratios of median background-subtracted fluorescence intensities; black lines indicate global Langmuir isotherm fits. Reported *K*_d_ values denote the median fit *K*_d_ for each sequence over 400 subsamplings of the data; reported errors denote interquartile ranges of *K*_d_ fit distributions (see **Methods**).

**Supplementary Figure 19.**
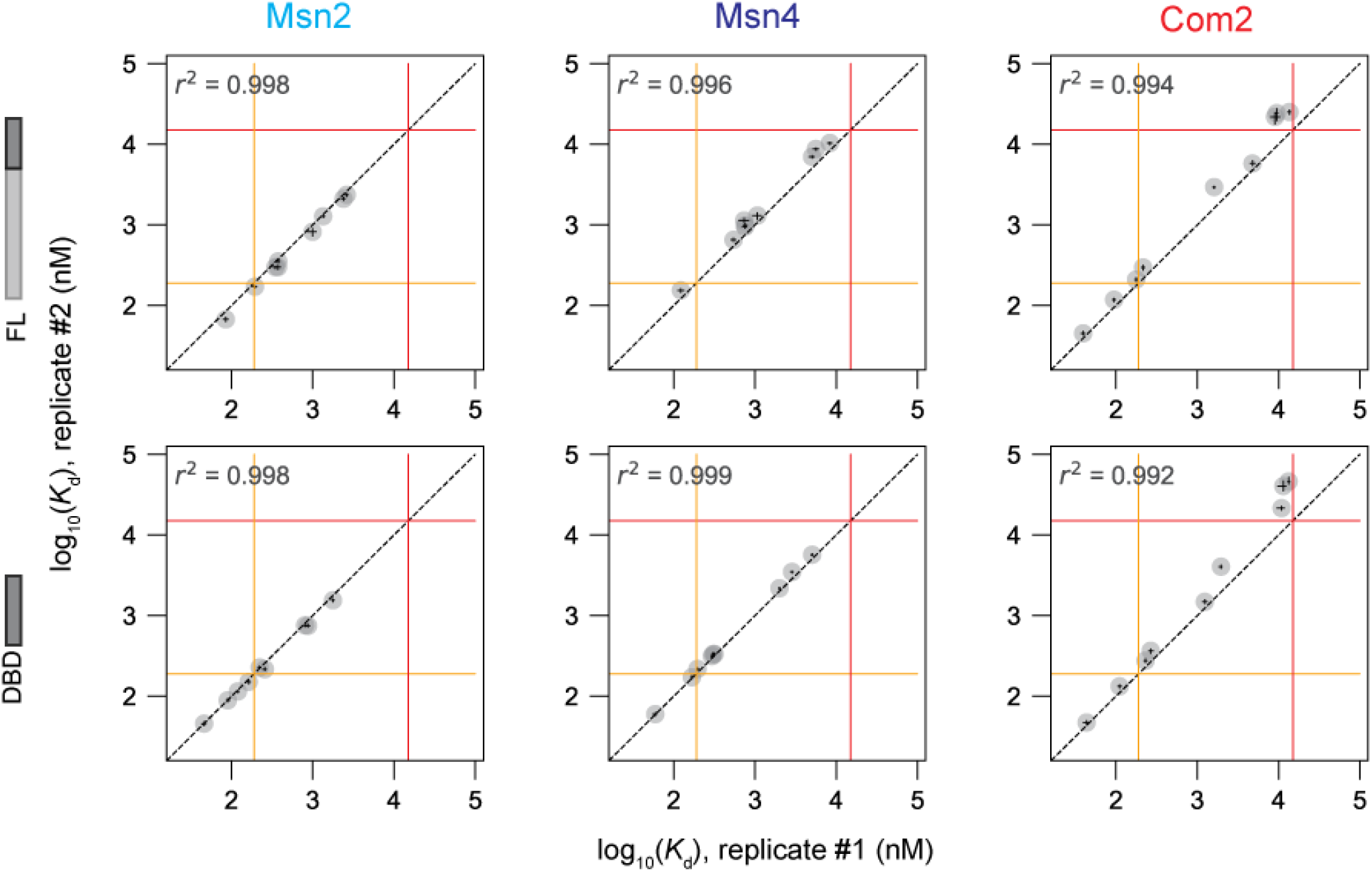
Scatter plots comparing TF-MITOMI fitted *K*_d_ values for fiducial sequences from ‘degenerate motif’ library across experimental replicates. Scatter plots showing log_10_-transformed fitted *K*_d_ values from TF-MITOMI measurements of fiducial sequences from ‘degenerate motif’ library for experimental replicate #2 versus experimental replicate #1. Each marker indicates the mean value for a given experiment; error bars represent interquartile ranges of fit *K*_d_ distributions for each sequence over 400 subsamplings and fits to the data (**Methods**). Black dashed line indicates 1:1 line; solid orange and red lines indicate thresholds (190 and 15000 nM), below and above which fit *K*_d_s are classified as upper and lower limits, respectively, based on binding simulations (**Methods**).

**Supplementary Figure 20.**
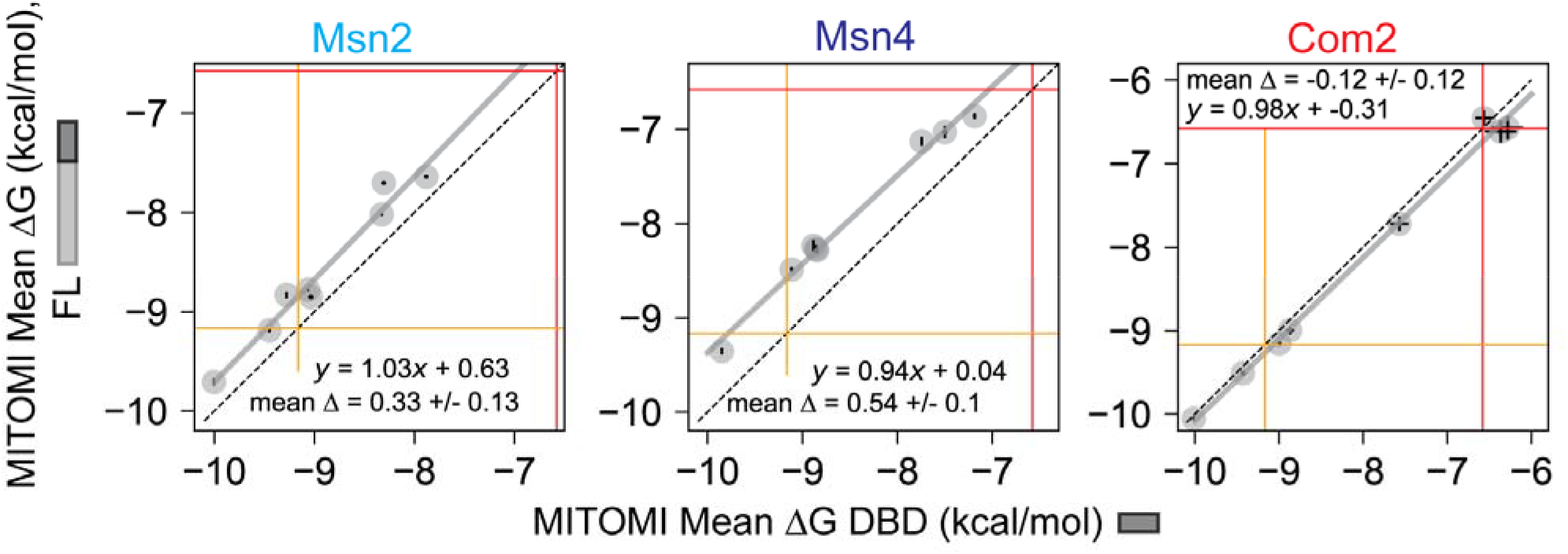
Scatter plots comparing TF-MITOMI fitted ΔG values for fiducial sequences from ‘degenerate motif’ library between FL versus DBD-only constructs. Markers represent mean ΔGs of DBD and FL constructs for each fiducial sequence; error bars indicate absolute error in fit ΔGs between the two replicate measurements. Dotted black lines indicate the 1:1 line; solid gray lines indicate linear regression fit line with annotated fit parameters; solid orange and red lines indicate lower and upper threshold *K*_d_ values of 190 and 15000 nM, below and above which mean fit values are considered as upper and lower limits, respectively.

**Supplementary Figure 21.**
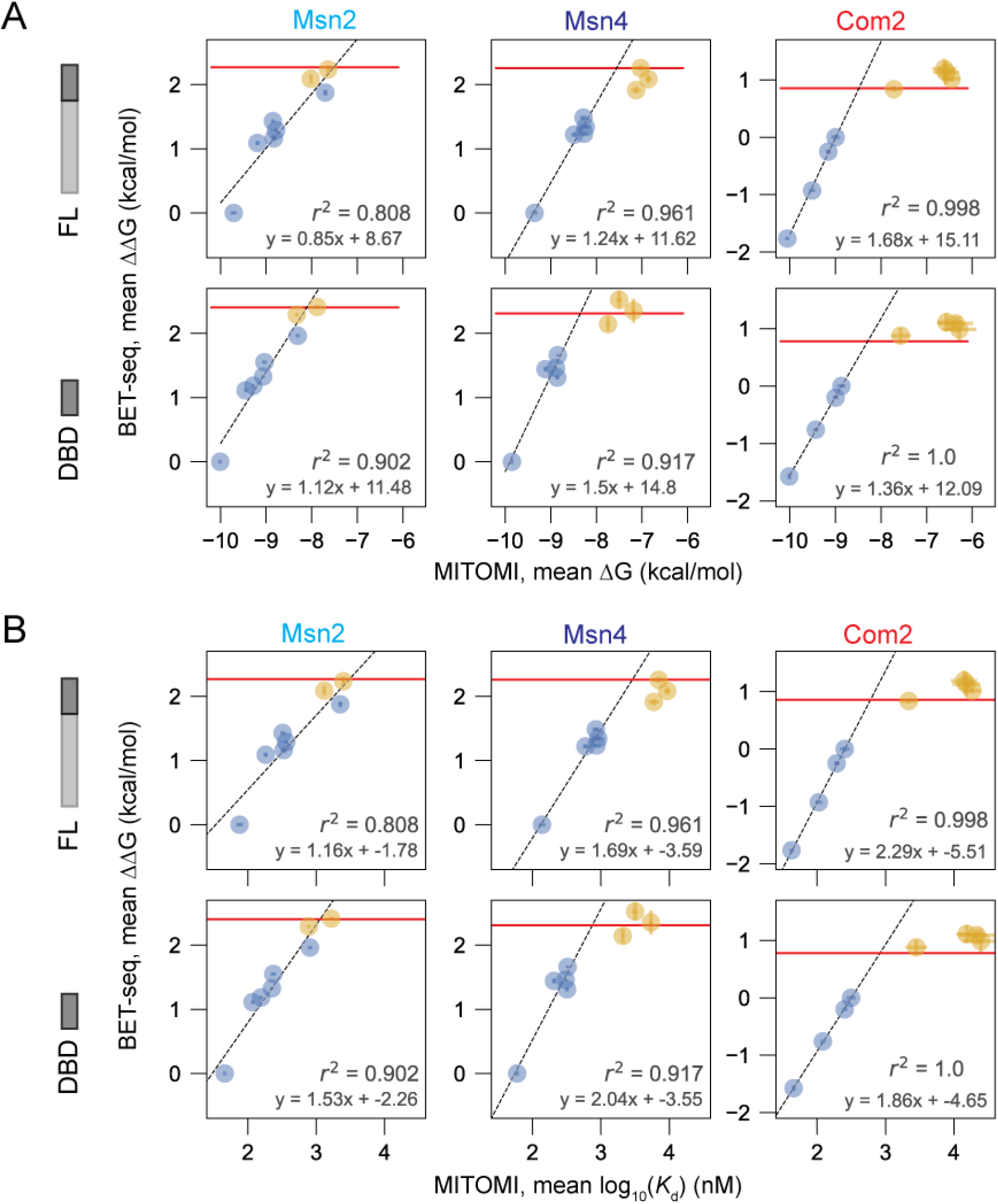
Calibration of BET-seq ΔΔGs using MITOMI-measured ΔGs and *K*_d_s of fiducial sequences from the ‘degenerate motif’ library. Markers represent the mean MITOMI ΔGs (**A**) or log_10_-transformed *K*_d_s (**B**) and mean BET-seq ΔΔGs for each fiducial sequence (centered on the sequence CCTTAAGGGGTTA); error bars indicate the differences between each replicate measurement and the mean of the two replicates. Horizontal red lines denote a noise threshold of 3 standard deviations away from the mean of the fitted noise distribution of the BET-seq data. Markers shown in blue were below the threshold and included in the regression; markers shown in orange were at or above the threshold and not included in the regression. Black dashed line indicates linear regression fit with the annotated fit parameters.

**Supplementary Figure 22.**
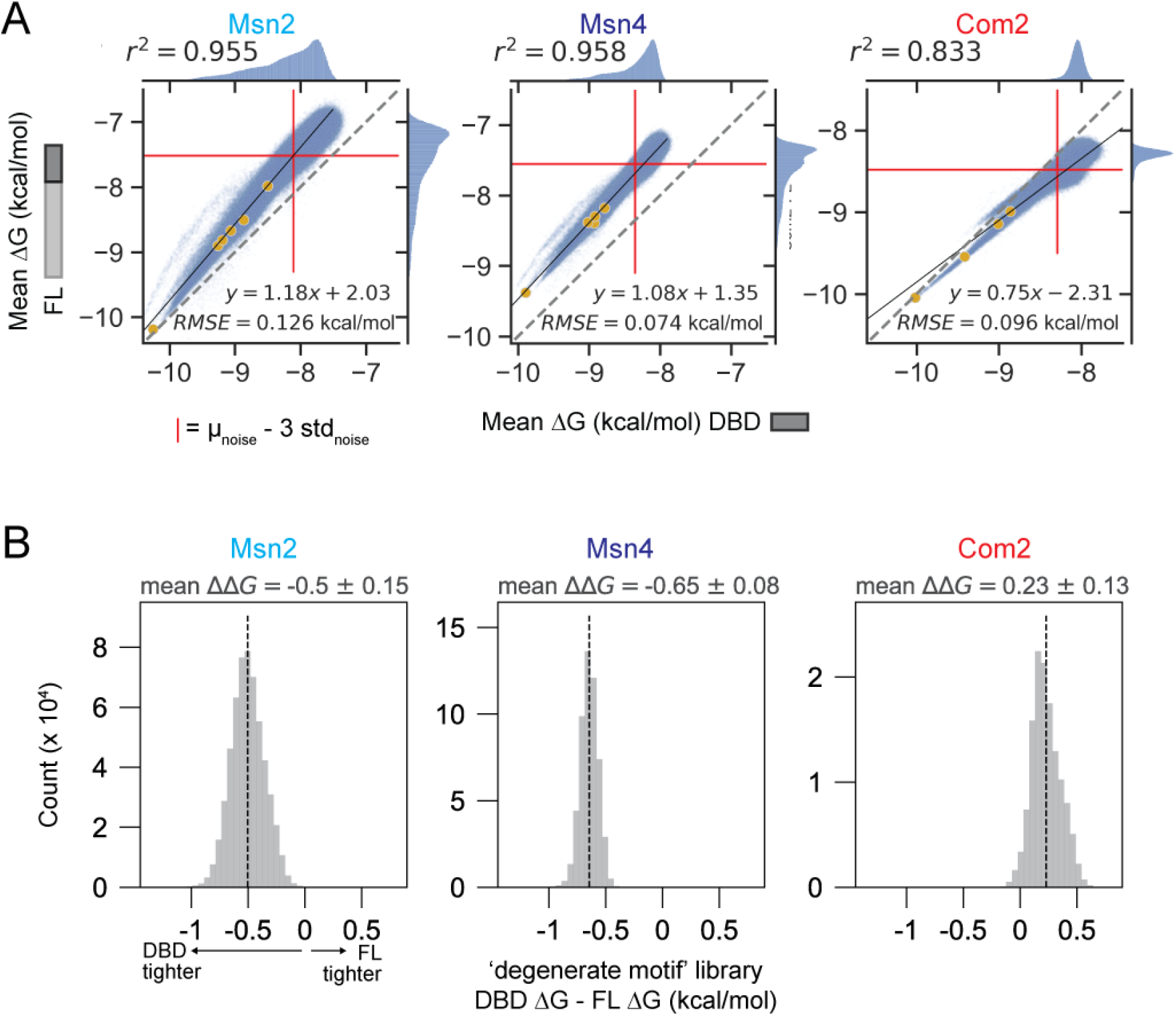
Comparison of DBD and FL calibrated ΔGs of binding to the ‘degenerate motif’ library. (**A**) Scatter plots comparing calibrated ΔGs of DBD and FL constructs for Msn2 (left), Msn4 (center), and Com2 (right). Blue markers indicate mean ΔGs of individual ‘degenerate motif’ library sequences; yellow markers indicate mean ΔGs of fiducial sequences used in the regression. Dotted lines indicate 1:1 lines. Red lines indicate noise thresholds of the mean of the noise fit minus 3 standard deviations of the noise. Pearson r^2^ values are reported for ΔΔG measurements outside of the noise of either replicate. (**B**) Histograms showing distributions of differences in ‘degenerate motif’ library calibrated mean ΔGs between DBD and FL constructs of Msn2 (left), Msn4 (center), and Com2 (right). Mean differences (ΔΔG) are reported for each pair and are indicated by vertical black dotted lines.

**Supplementary Figure 23.**
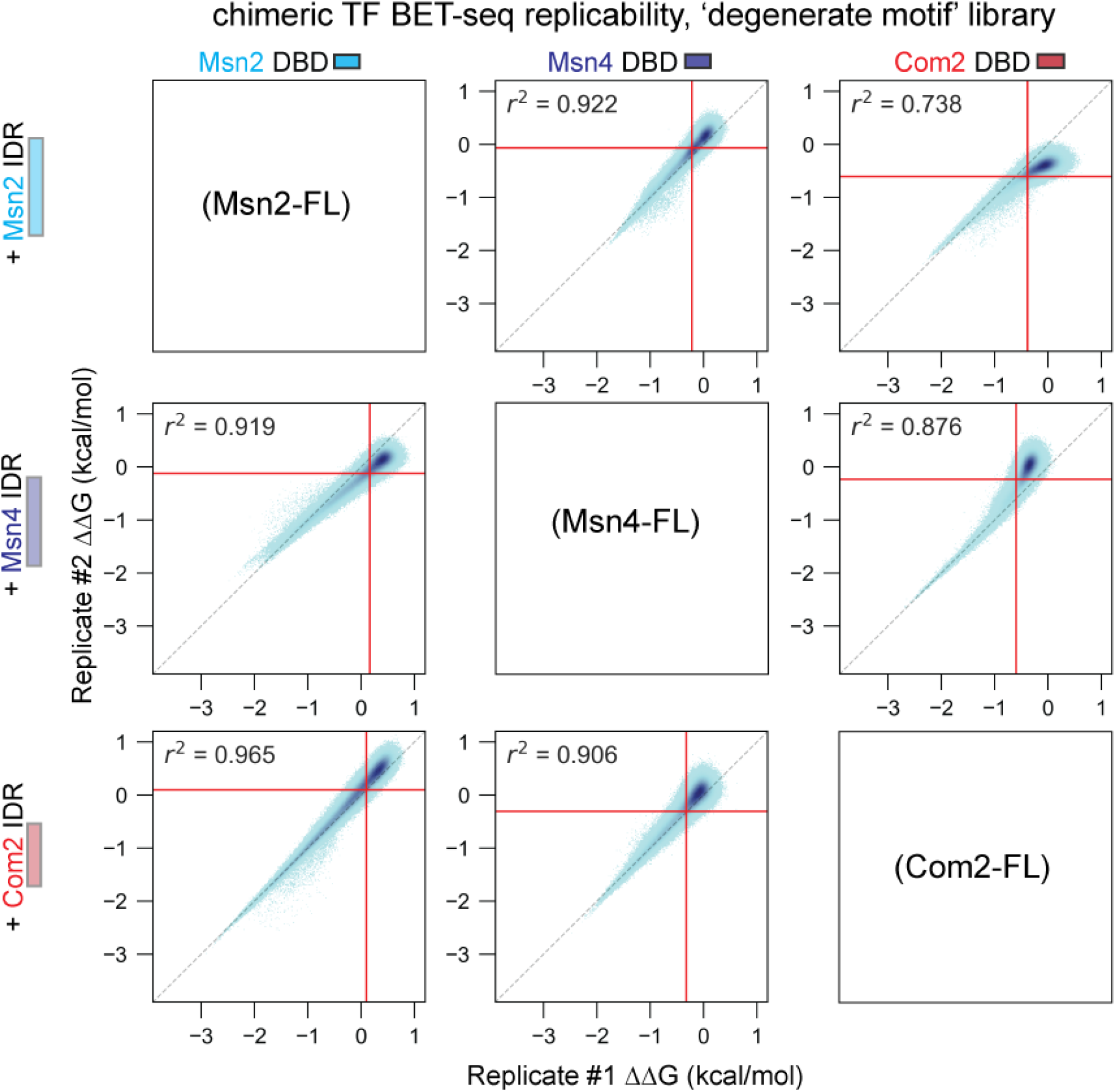
2D histograms comparing BET-seq ΔΔG measurements for each chimeric TF construct binding to the ‘degenerate motif’ library across 2 technical replicates. Columns and rows indicate which DBD is attached to which IDR, respectively. Sequences were binned by replicate 1 and 2 ΔΔG values (computed relative to CTTCCGGGGCCGG) and pixel darkness indicates density of sequences per bin. Dotted lines indicate 1:1 lines. Red lines indicate noise thresholds of the mean of the noise fit minus 2 standard deviations of the noise. Pearson *r*^2^ values are reported for ΔΔG measurements below the noise threshold of either replicate.

**Supplementary Figure 24.**
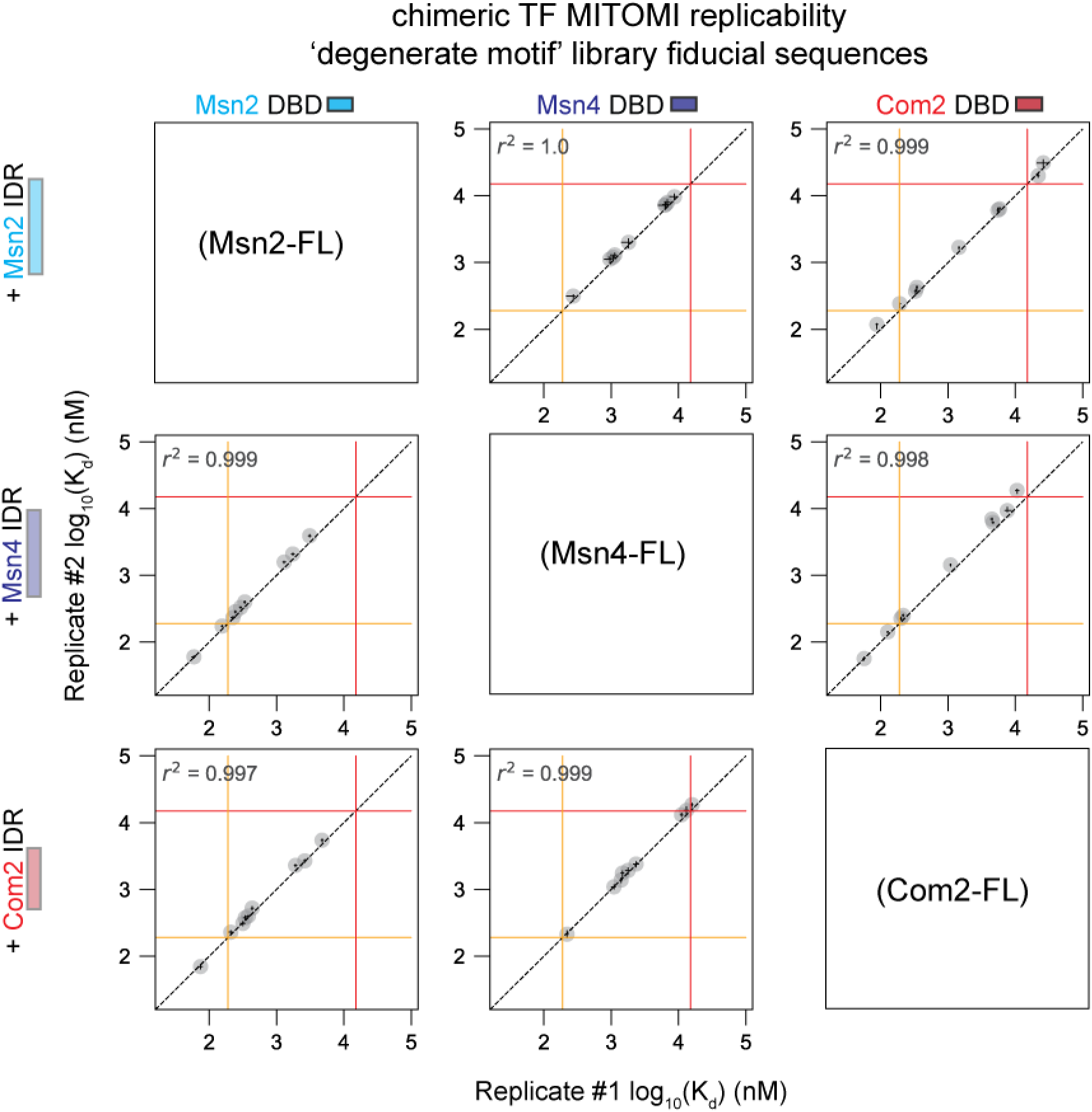
Scatter plots comparing MITOMI-measured *K*_d_s for chimeric TF constructs interacting with ‘degenerate motif’ library fiducial sequences across 2 experimental replicates. Row and column identities specify the DBD and IDR comprising each construct. Each marker indicates the log_10_-transformed mean fitted *K*_d_ value for a given sequence within a given experiment replicate. Error bars represent the propagated log-transformed interquartile range of the distribution of fitted *K*_d_s for a given sample over 400 subsamplings of the binding data. Black dotted lines indicate 1:1 lines; solid orange and red lines indicate thresholds (190 and 15000 nM), below and above which fit *K*_d_s are classified as upper and lower limits, respectively, based on binding simulations (**Methods**). All curves are available in **Supplementary File 1**.

**Supplementary Figure 25.**
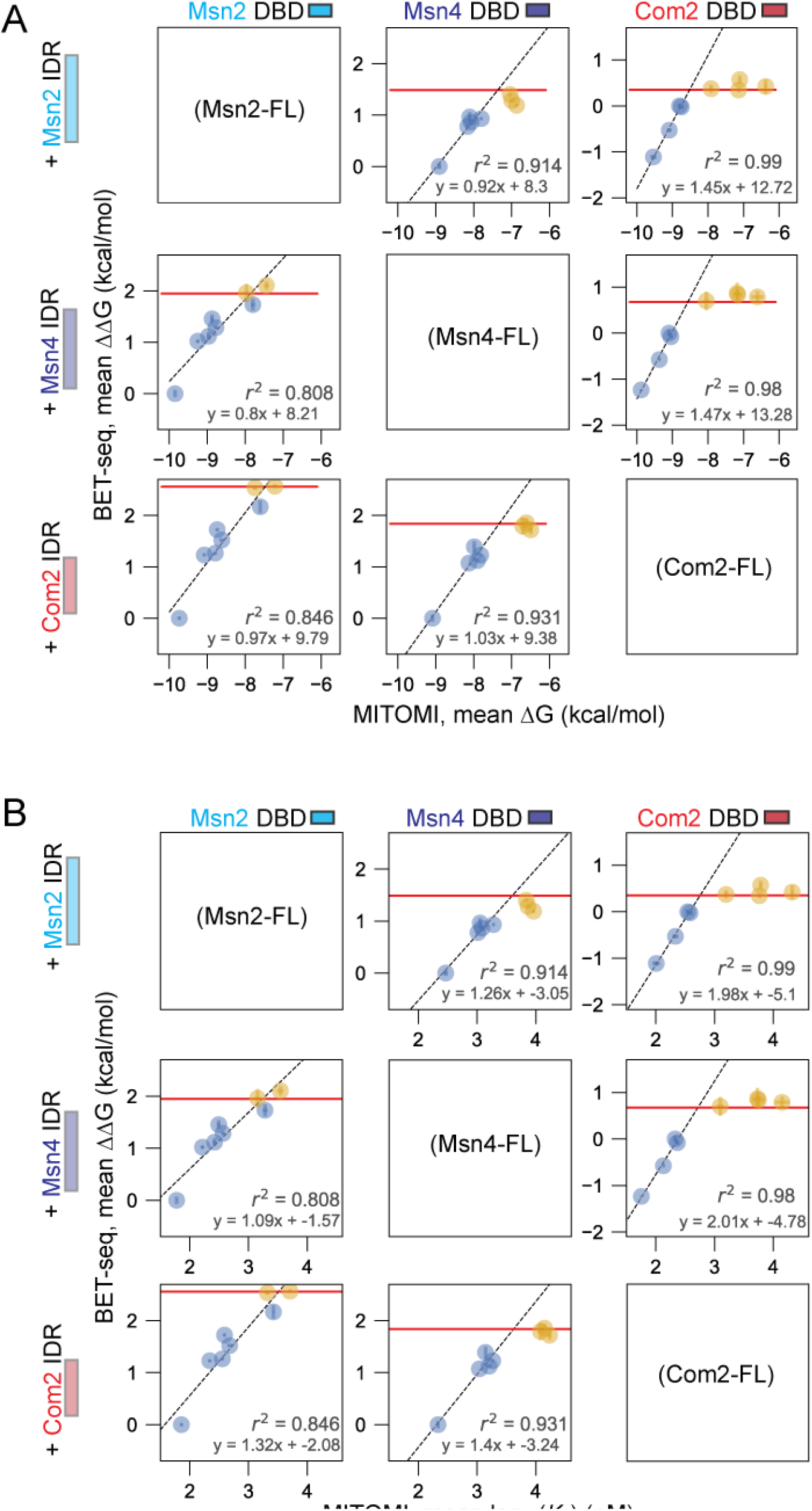
Calibration of BET-seq ΔΔGs for chimeric constructs using MITOMI-measured *K*_d_s of fiducial sequences from the ‘degenerate motif’ library. Columns and rows indicate which DBD is attached to which IDR, respectively. Markers represent the mean MITOMI ΔGs(**A**) or log_10_-transformed *K*_d_s (**B**) and mean BET-seq ΔΔGs for each fiducial sequence (centered on the sequence CCTTAAGGGGTTA); error bars indicate the differences between each replicate measurement and the mean of the two replicates. Horizontal red lines denote a noise threshold of 3 standard deviations away from the mean of the fitted noise distribution of the BET-seq data. Markers shown in blue were below the threshold and included in the regression; markers shown in orange were at or above the threshold and not included in the regression. Black dashed line indicates linear regression fit with the annotated fit parameters.

**Supplementary Figure 26.**
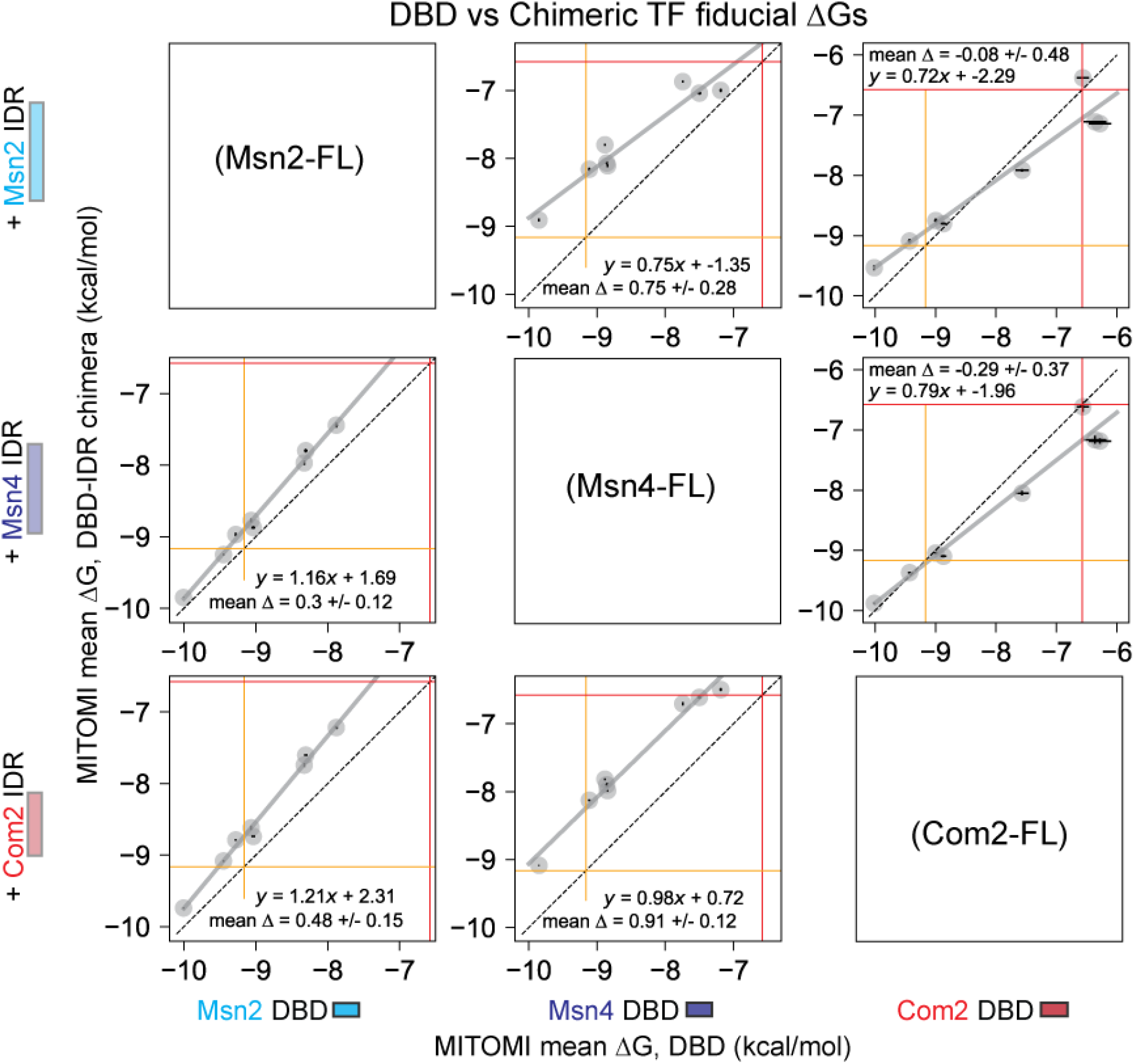
Scatter plots comparing TF-MITOMI fitted ΔG values for fiducial sequences from ‘degenerate motif’ library between chimeric and DBD-only constructs. Columns and rows indicate which DBD is attached to which IDR, respectively. Markers indicate mean ΔGs for each fiducial sequence; error bars indicate absolute error in fit ΔGs between two replicate measurements. Dotted black lines indicate the 1:1 line; solid gray lines indicate the linear regression fit line with annotated fit parameters; solid orange and red lines indicate thresholds (190 and 15000 nM), below and above which fit *K*_d_s are classified as upper and lower limits, respectively, based on binding simulations (**Methods**).

**Supplementary Figure 27.**
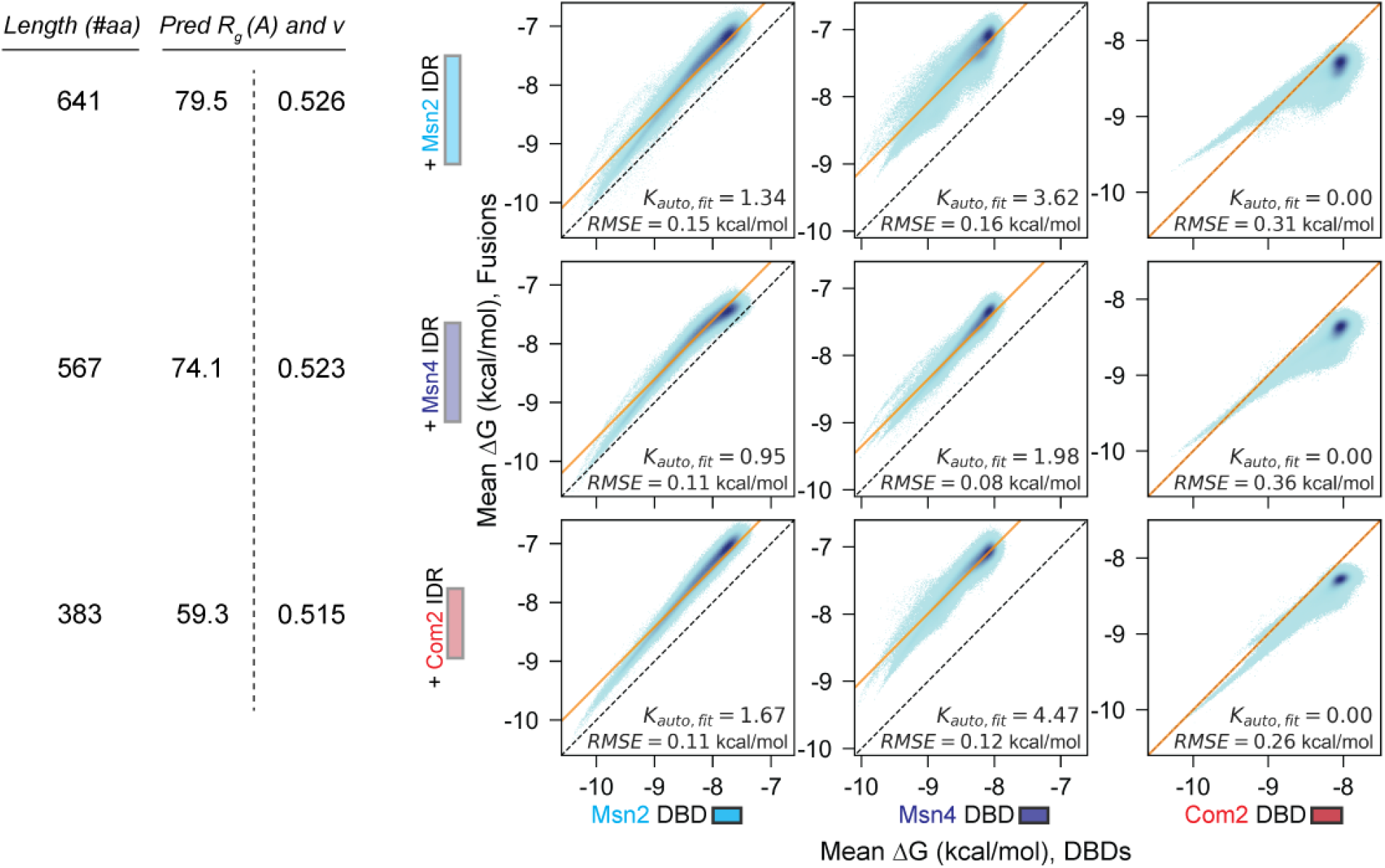
2D histograms comparing calibrated ‘degenerate motif’ library ΔGs of chimeric and DBD-only constructs with *K*_auto_ fits. Columns and rows indicate which DBD is attached to which IDR, respectively. Library sequences were binned by DBD and IDR-fusion calibrated ΔGs and pixel darkness indicates density of sequences. Yellow lines indicate 3-state autoinhibition model fit lines for each DBD-Fusion construct pair with the fit *K*_auto_ and root-mean-squared-error (RMSE) of the fit reported. Dotted black lines indicate the 1:1 line. IDR lengths and ALBATROSS predicted radii of gyration (R_g_) and Flory scaling exponents (ν) are shown for each row.

**Supplementary Figure 28.**
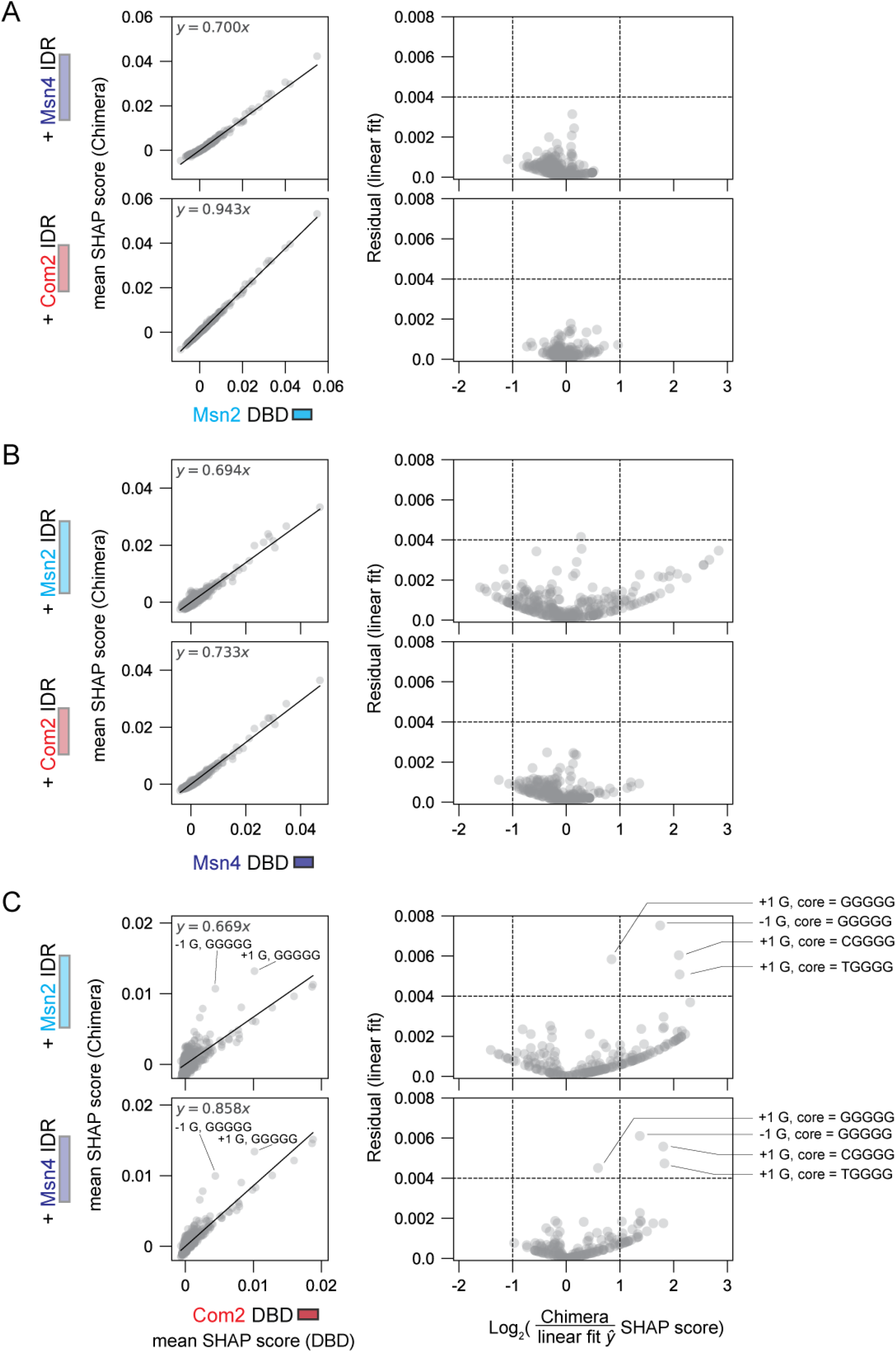
Comparison of flank nucleotide contributions within each core motif variant in DBD and chimeric TF constructs. Left: Mean SHAP scores for chimeric TF constructs versus DBD-only construct for each nucleotide in each motif-flanking position 4 bp up- and downstream of either an AGGGG motif or a single nucleotide variant of the AGGGG motif. Black lines indicate linear fits with the annotated fit parameters. Right: Residual from the linear fit versus the log_2_-transformed ratio of the chimeric TF DFIM to linear fit-predicted SHAP scores. To account for noise within low magnitude SHAP scores, all SHAP scores below 0.0008 were brought to a value of 0.0008 for this analysis. Dotted lines indicate 2-fold changes and residuals of 0.004. Annotations highlight flanking nucleotide/motif pairs of interest. Plots are shown for chimeric constructs in comparison with Msn2 DBD (**A**), Msn4 DBD (**B**), and Com2 DBD (**C**).

**Supplementary Figure 29.**
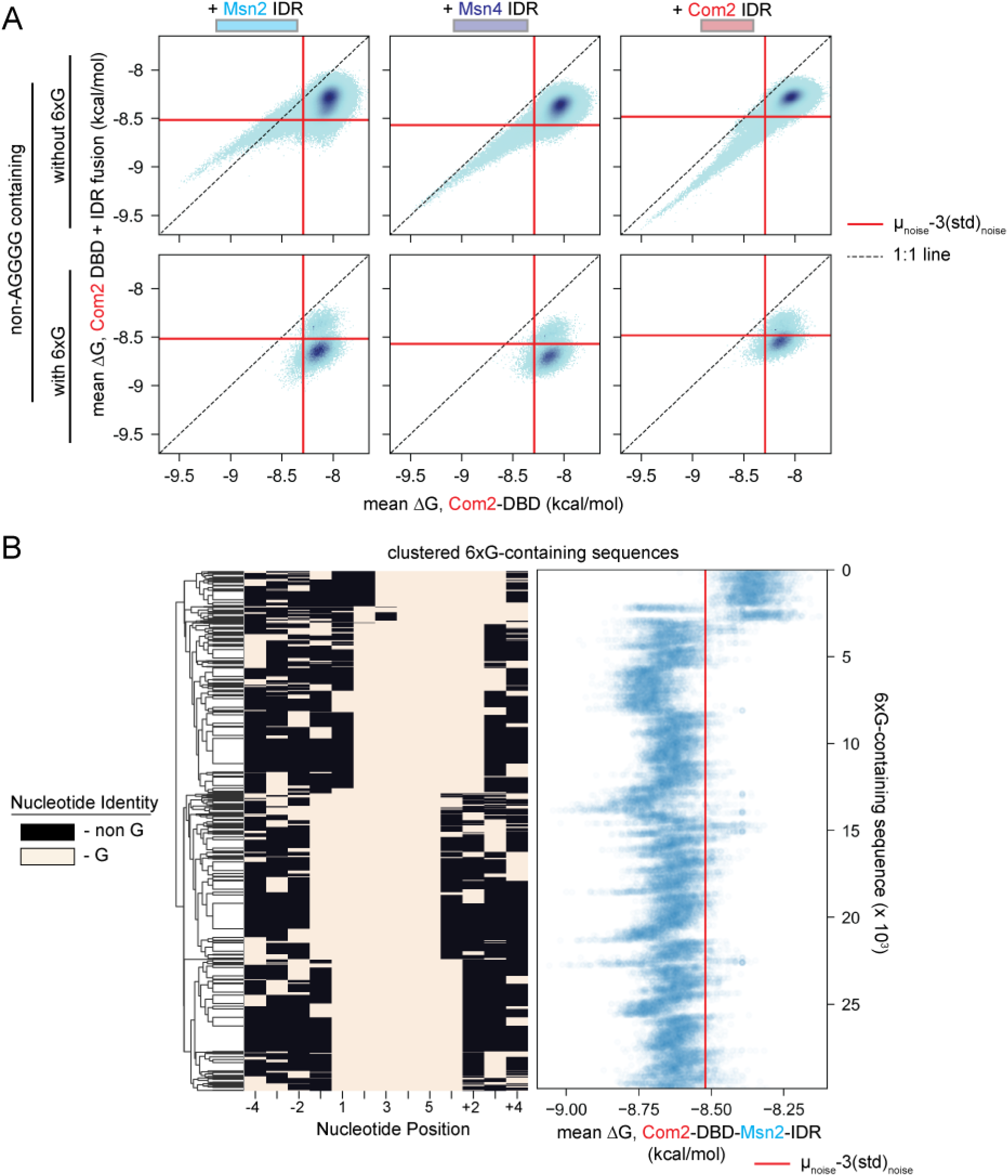
Comparison of 6xG-motif binding in the ‘degenerate motif’ library between Com2-DBD and its associated chimeric TFs. (**A**) 2D histograms comparing mean calibrated ΔGs for the Com2 DBD fused to the Msn2 (left), Msn4 (middle), or Com2 (right) IDR versus the Com2 DBD alone for sequences lacking AGGGG sequences and either without (top) or with 6xG-motifs (bottom). Library sequences were binned by DBD and IDR-fusion calibrated ΔG and pixel darkness indicates density of sequences. Dotted lines indicate 1:1 lines; red solid lines indicate thresholds of 3 standard deviations above the mean of the noise distribution used to delineate ‘bound’ from ‘unbound’ sequences. (**B**) (Left) Binarized heatmap showing positions of G *vs* non-G nucleotides within the ‘degenerate motif’ library sequences containing 6xG stretches but no AGGGG after clustering based on the positions of G nucleotides. (Right) Measured mean ΔG of the Com2-DBD-Msn2-IDR chimera for each sequence depicted in the heatmap. Red solid line indicates the same threshold used to delineate ‘bound’ from ‘unbound’ sequences as in (**A**).

**Supplementary Figure 30.**
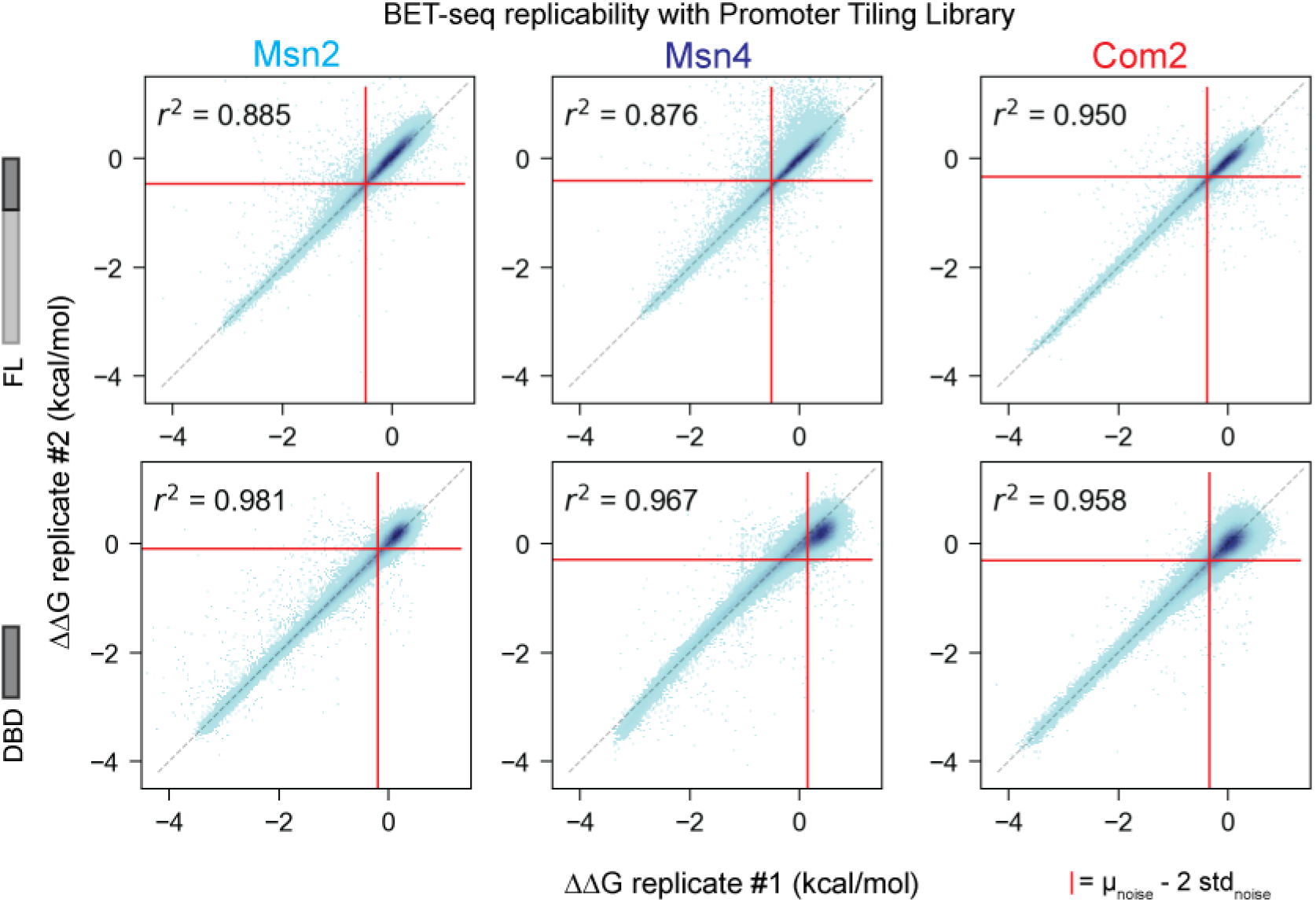
2D histograms comparing measured ‘promoter tile’ library ΔΔG values for each FL and DBD-only construct across 2 technical replicates. Sequences were binned by replicate 1 and 2 ΔΔG (computed relative to TGCAATTACGTAATTTTAGCCGCTGAGAAGCGGATGGTAATGAGACAAGTTGATATCAAA CAGATACATATTTAA) and pixel darkness indicates density of sequences. Dotted black lines indicate 1:1 lines; red lines indicate thresholds of 2 standard deviations from noise mean values (calculated by fitting a Gaussian curve to a synthetic ‘noise’ distribution of each experiment constructed by taking the ΔΔG values greater than the empirical center and their reflection across that center value – see **Methods**). Pearson *r*^2^ values shown were calculated from ΔΔG values for sequences lying outside of 2 standard deviations from the noise of either replicate.

**Supplementary Figure 31.**
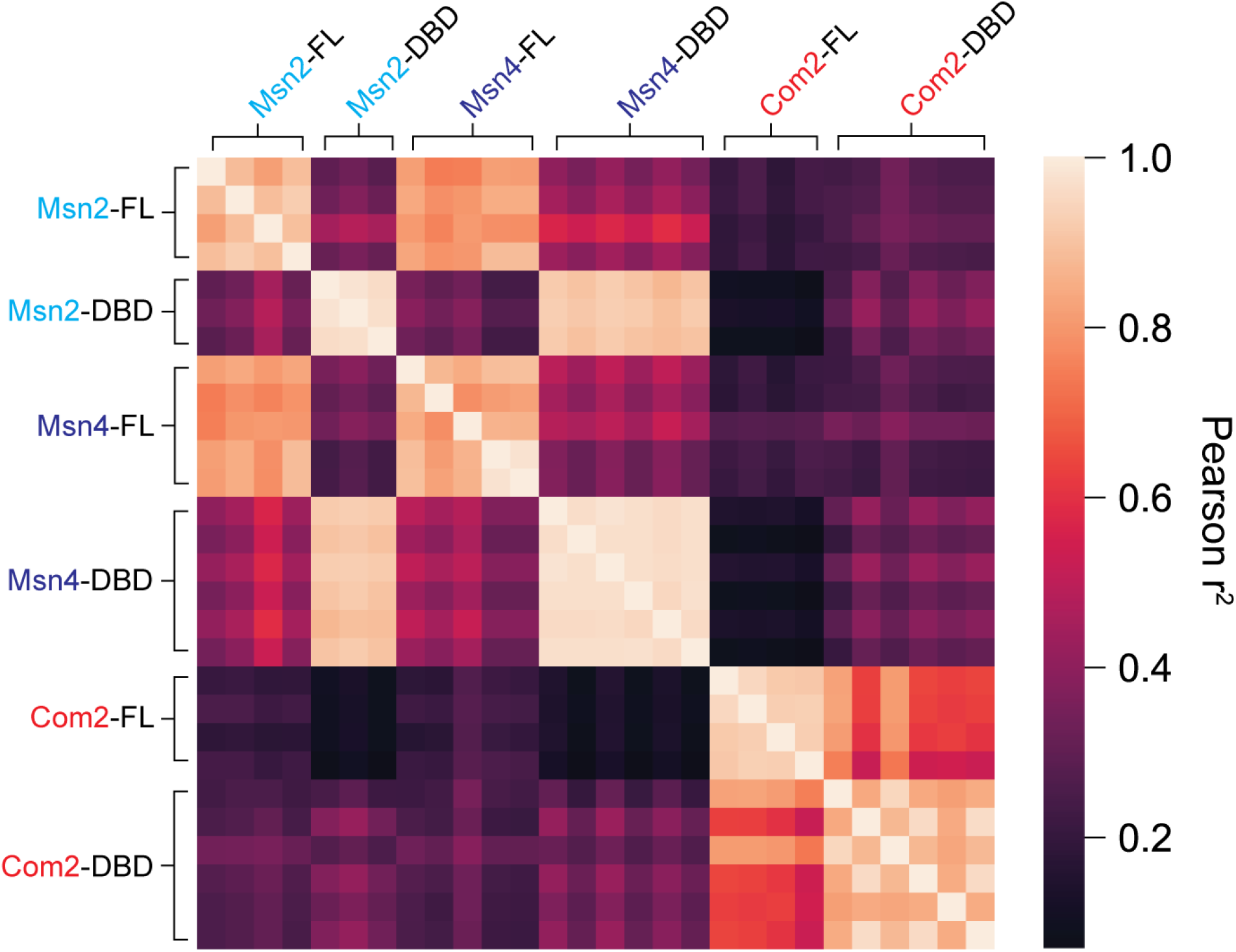
Heatmap of pairwise Pearson *r*^2^ values for ΔΔGs for promoter tiles between all constructs and replicates. Pearson r^2^ values are reported for ΔΔG measurements outside of the noise of either replicate (defined by using a threshold of 2 standard deviations above the mean of the noise distribution, see **Methods**).

**Supplementary Figure 32.**
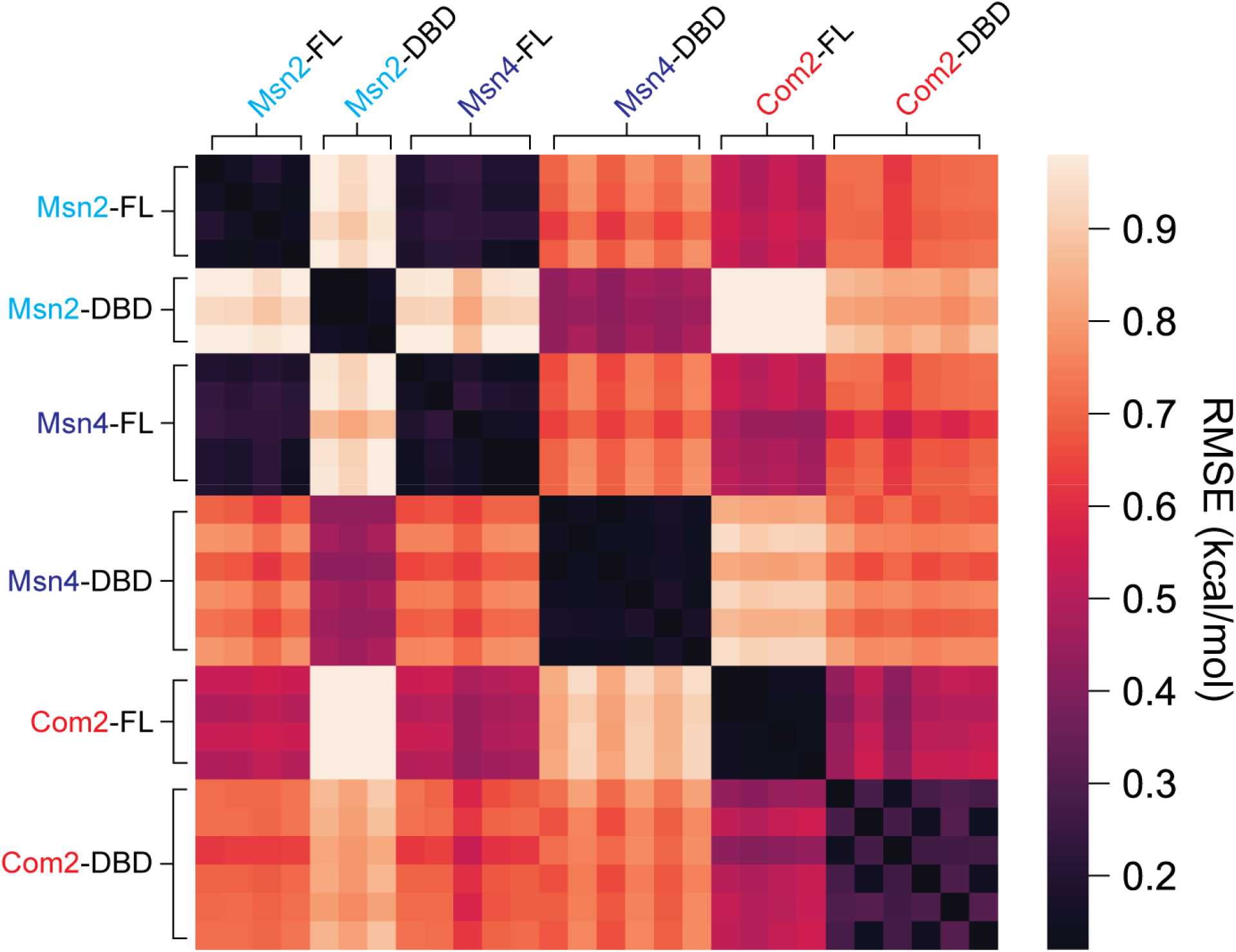
Heatmap of pairwise RMSE for ΔΔGs for promoter tiles between all constructs and replicates. RMSEs are reported for ΔΔG measurements outside of the noise of either replicate (defined by using a threshold of 2 standard deviations above the mean of the noise distribution, see **Methods**).

**Supplementary Figure 33.**
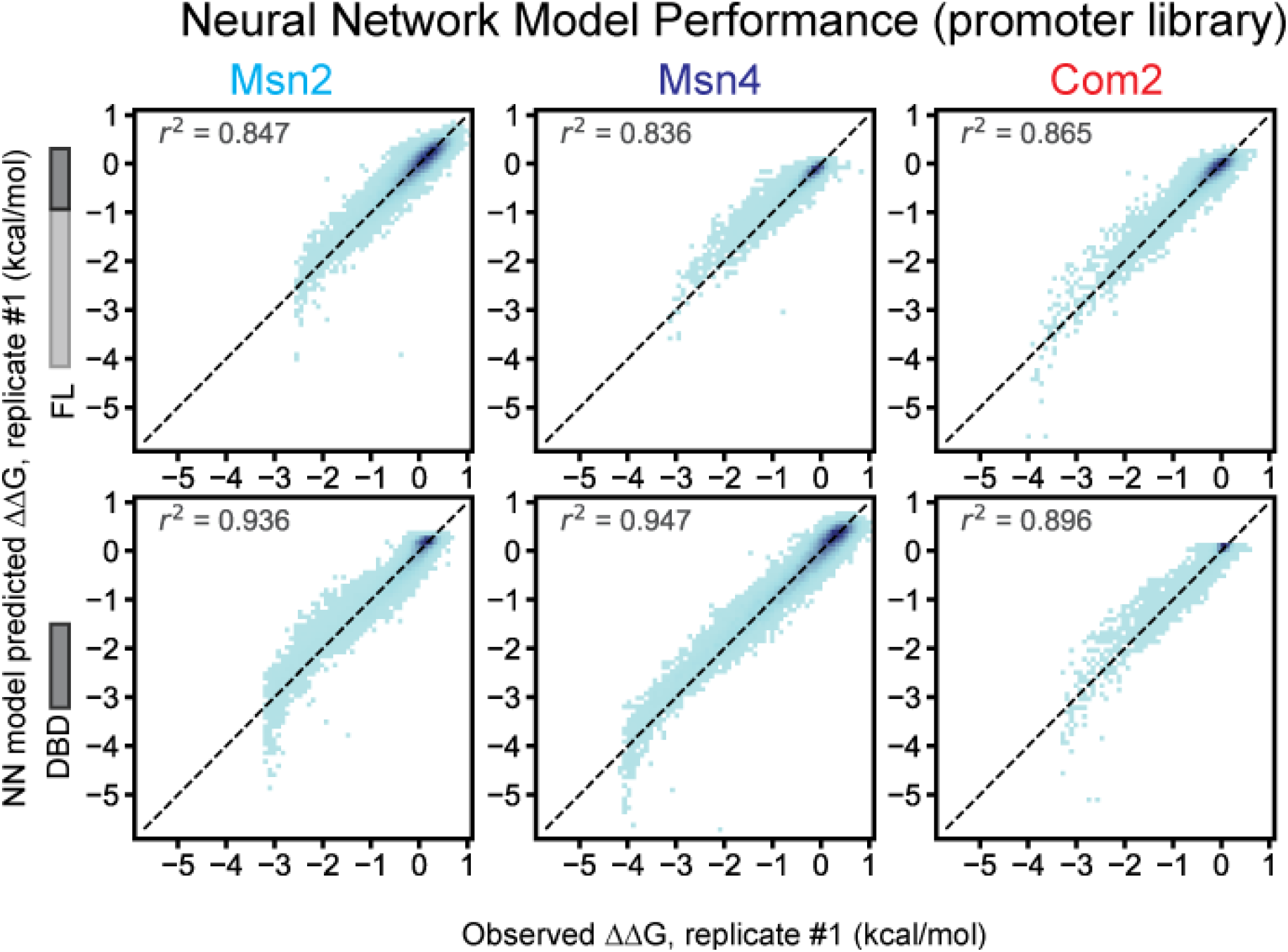
Neural network model performance predicting ‘promoter’ library ΔΔG measurements. 2D histograms comparing AffinityNet model-predicted ΔΔGs versus observed ΔΔGs for replicate #1 of each FL (top row) and DBD-only (bottom row) constructs of Msn2 (left), Msn4 (middle), and Com2 (right). ‘Promoter’ library sequences were binned by observed and predicted ΔΔGs and pixel darkness indicates density of sequences per bin. Black dashed line indicates 1:1 line, annotated Pearson *r*^2^ values are computed for all points (Pearson *r*^2^ > 0.84 for all constructs).

**Supplementary Figure 34.**
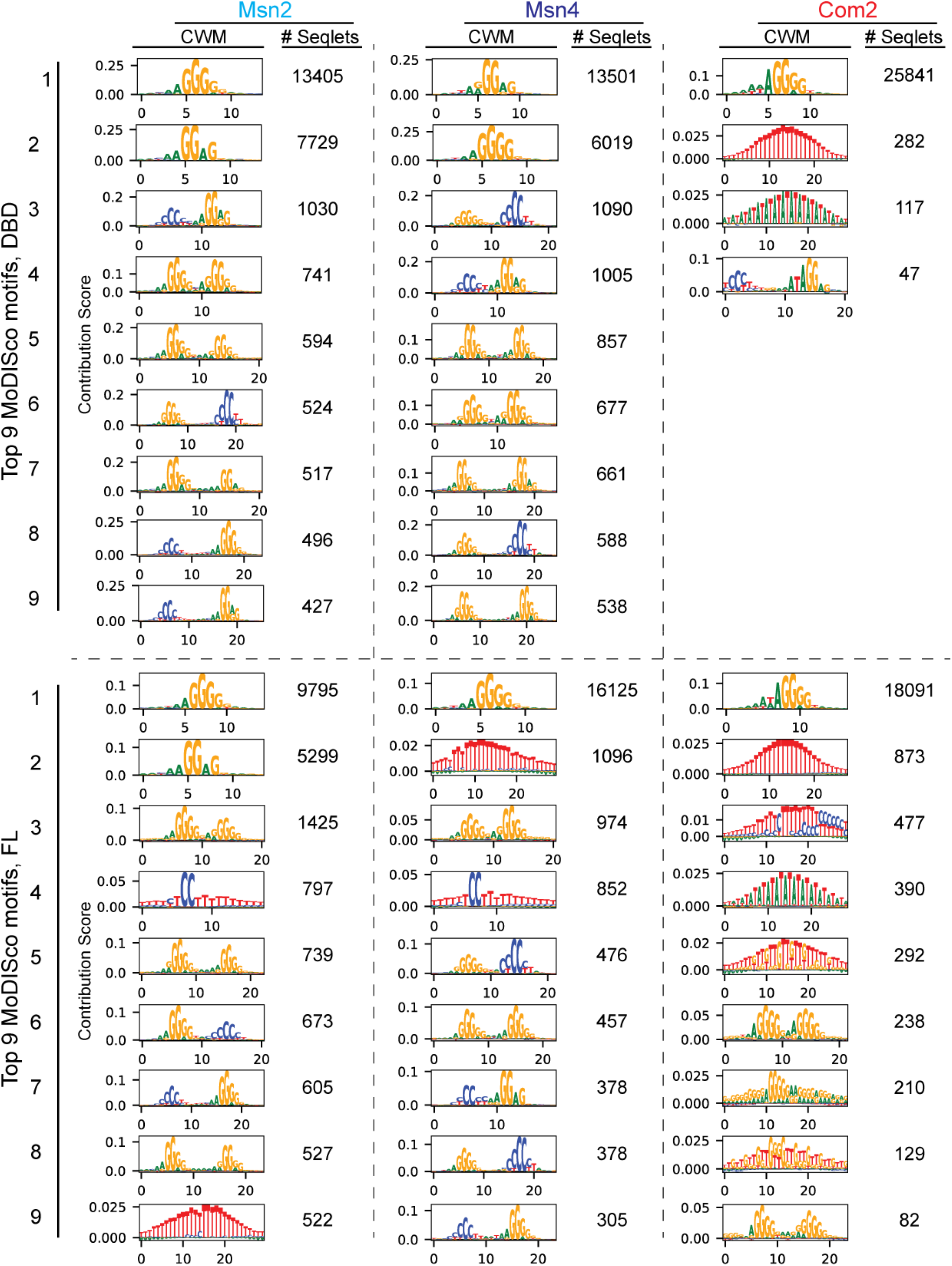
Top 9 MoDISco CWM motifs found in models of ‘promoter’ library binding of FL and DBD-only Msn2, Msn4, and Com2 constructs. Count column values represent the number of seqlets assigned to a given motif cluster.

**Supplementary Figure 35.**
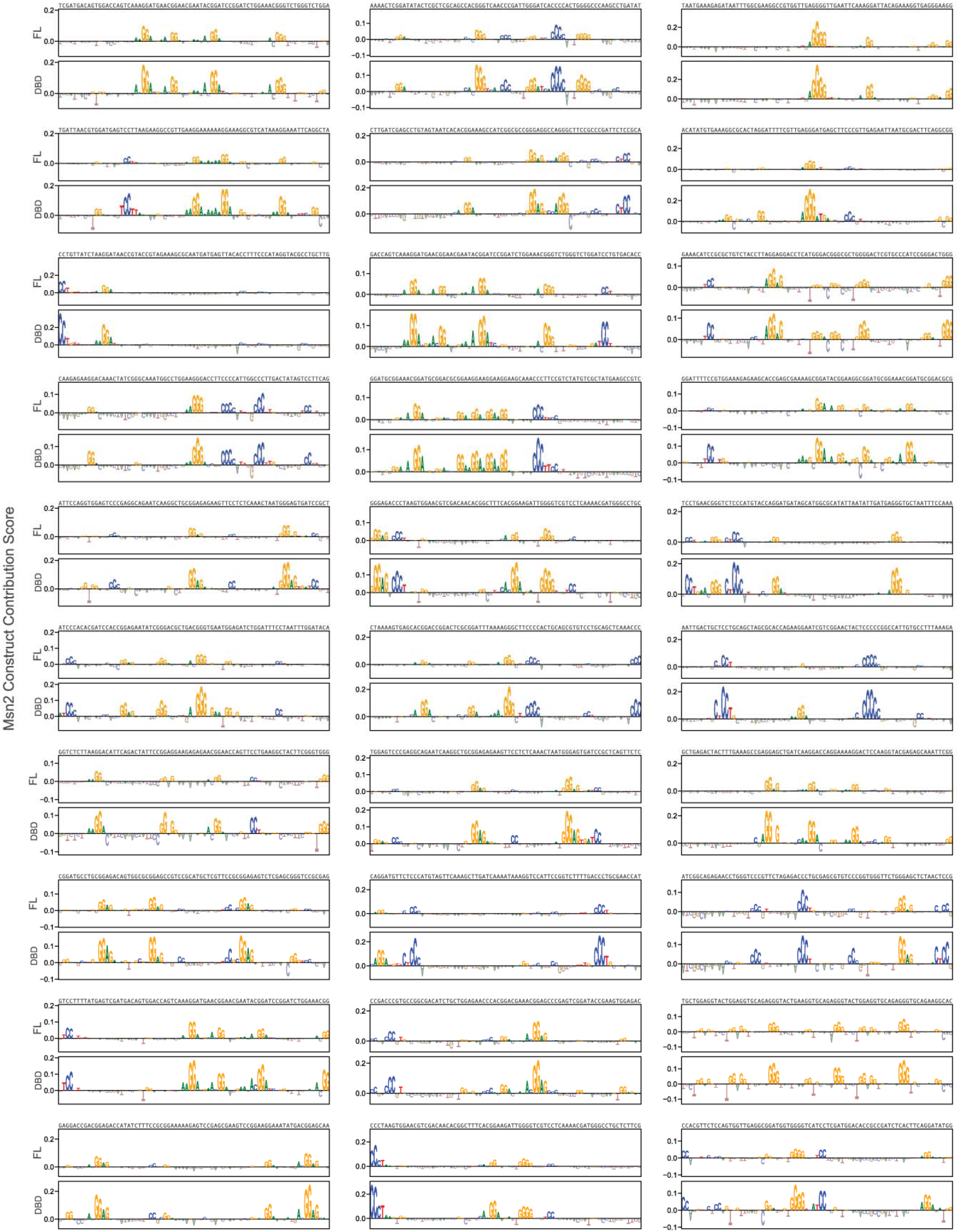
Contribution scores of top 30 ‘promoter’ library sequences preferred by Msn2’s DBD relative to FL Msn2. FL and DBD contribution scores are plotted together for each promoter tile, with each promoter tile’s sequence printed above plotted contribution scores.

**Supplementary Figure 36.**
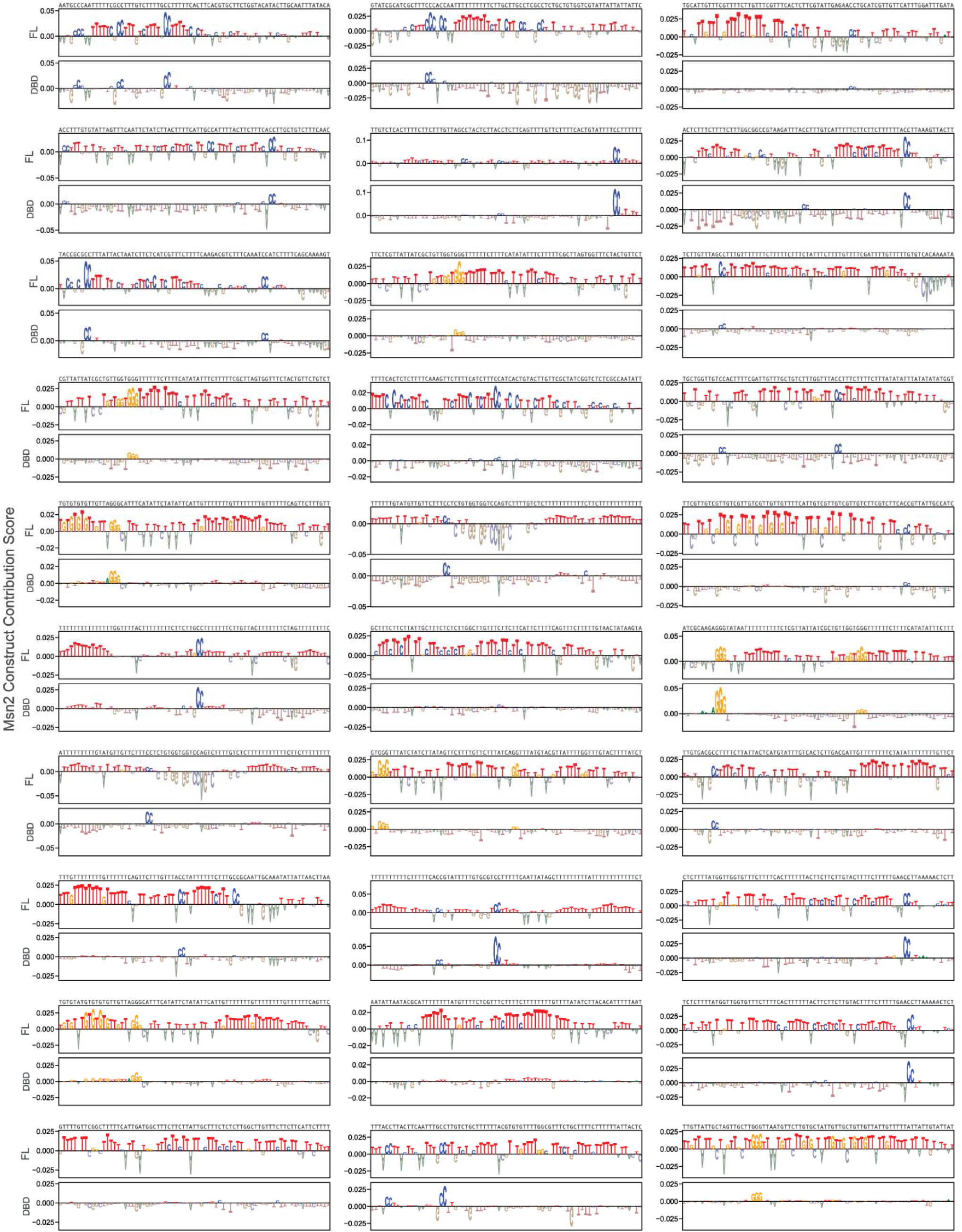
Contribution scores of top 30 ‘promoter’ library sequences preferred by Msn2 FL relative to Msn2 DBD. FL and DBD contribution scores are plotted together for each promoter tile, with each promoter tile’s sequence printed above plotted contribution scores.

**Supplementary Figure 37.**
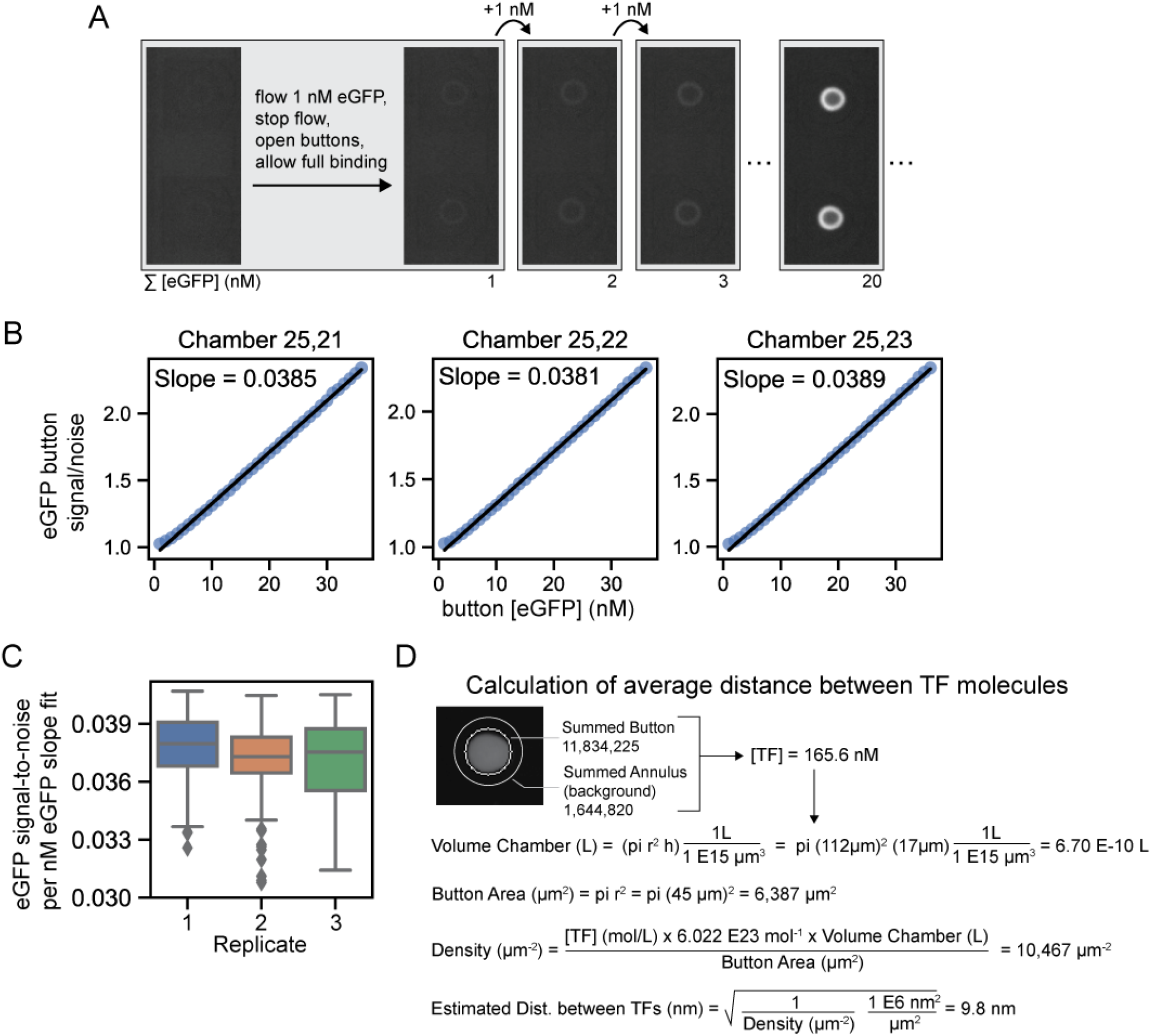
eGFP standard curve and estimation of average distances between protein molecules. (**A**) Schematic of calibration curve acquisition showing iterative introduction of 1 nM eGFP into each reaction chamber, opening of the button valves to expose anti-eGFP-patterned device surfaces, and binding of eGFP until saturation Each titration repeated this process 35 times. (**B**) Plots show linear fits of concentrations of eGFP bound to the button surface to the GFP channel signal-to-noise ratio (the ratio of fluorescence in the button area and the “annulus” area which represents the background fluorescence over the same total area). (**C**) Box plot showing the distribution of slopes fitted across all chambers of the device for three replicates. Boxes denote distribution median and 1st and 3rd quartiles; whiskers denote the maximum and minimum values within 1.5 interquartile ranges and outliers are plotted with individual markers. (**D**) Example calculation of average distance between TF molecules for a given button using known chamber volumes, button areas, and TF concentration (calculated using the fitted slope from the eGFP standard curve).

**Supplementary Figure 38.**
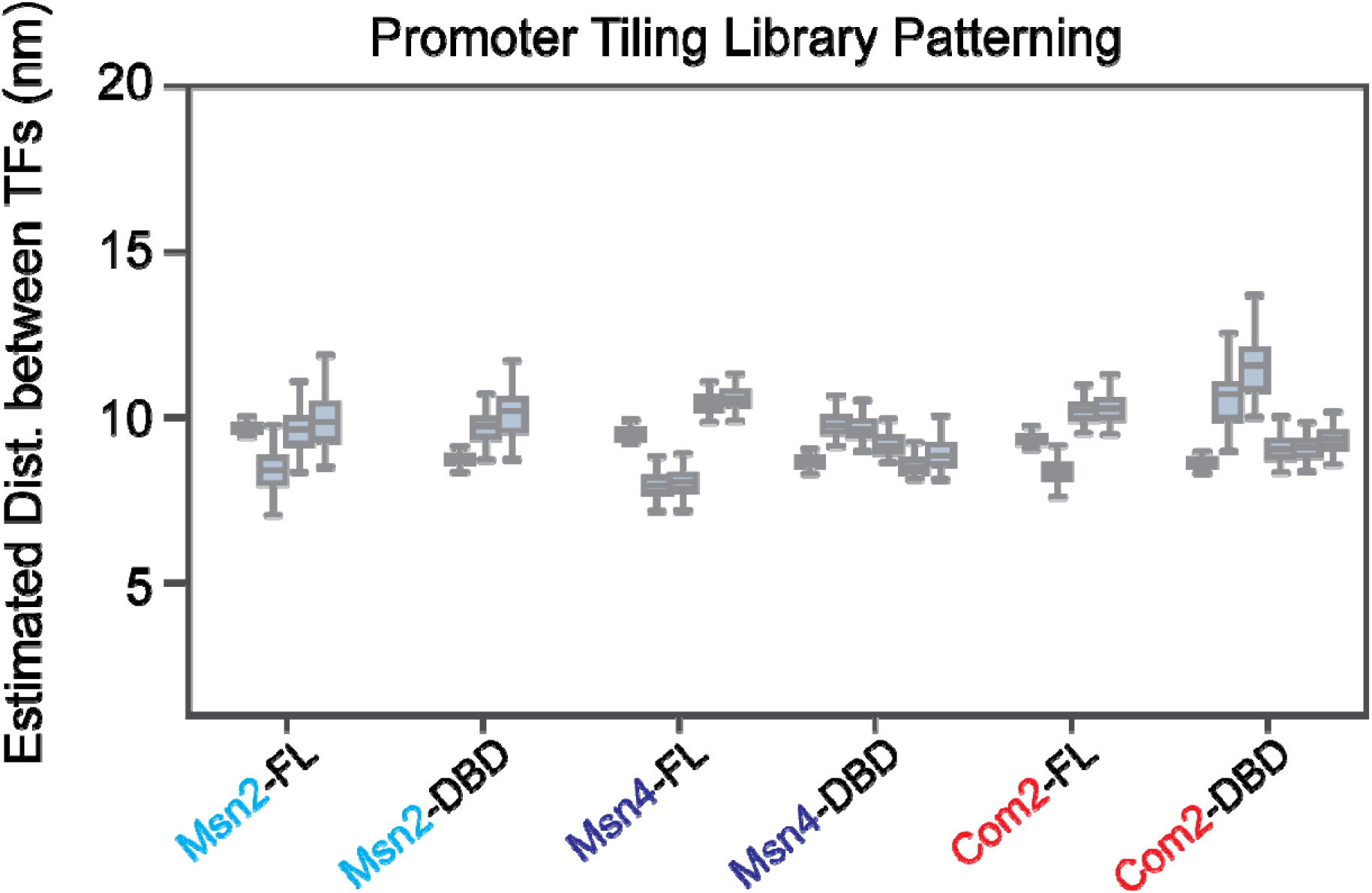
Distributions of estimated distances between adjacent TF molecule for FL and DBD constructs of Msn2, Msn4, and Com2 in BET-seq experiments measuring binding to the ‘promoter library’. Each box plot shows the distribution of estimated distances between adjacent TF molecules (calculated from background-subtracted eGFP fluorescence intensities of ‘button’ areas within the 256 chambers of a device, as in Supplementary Fig. 37); each box represents one BET-seq experiment. Boxes denote distribution median and 1st and 3rd quartiles; whiskers denote the maximum and minimum values within 1.5 interquartile ranges.

**Supplementary Figure 39.**
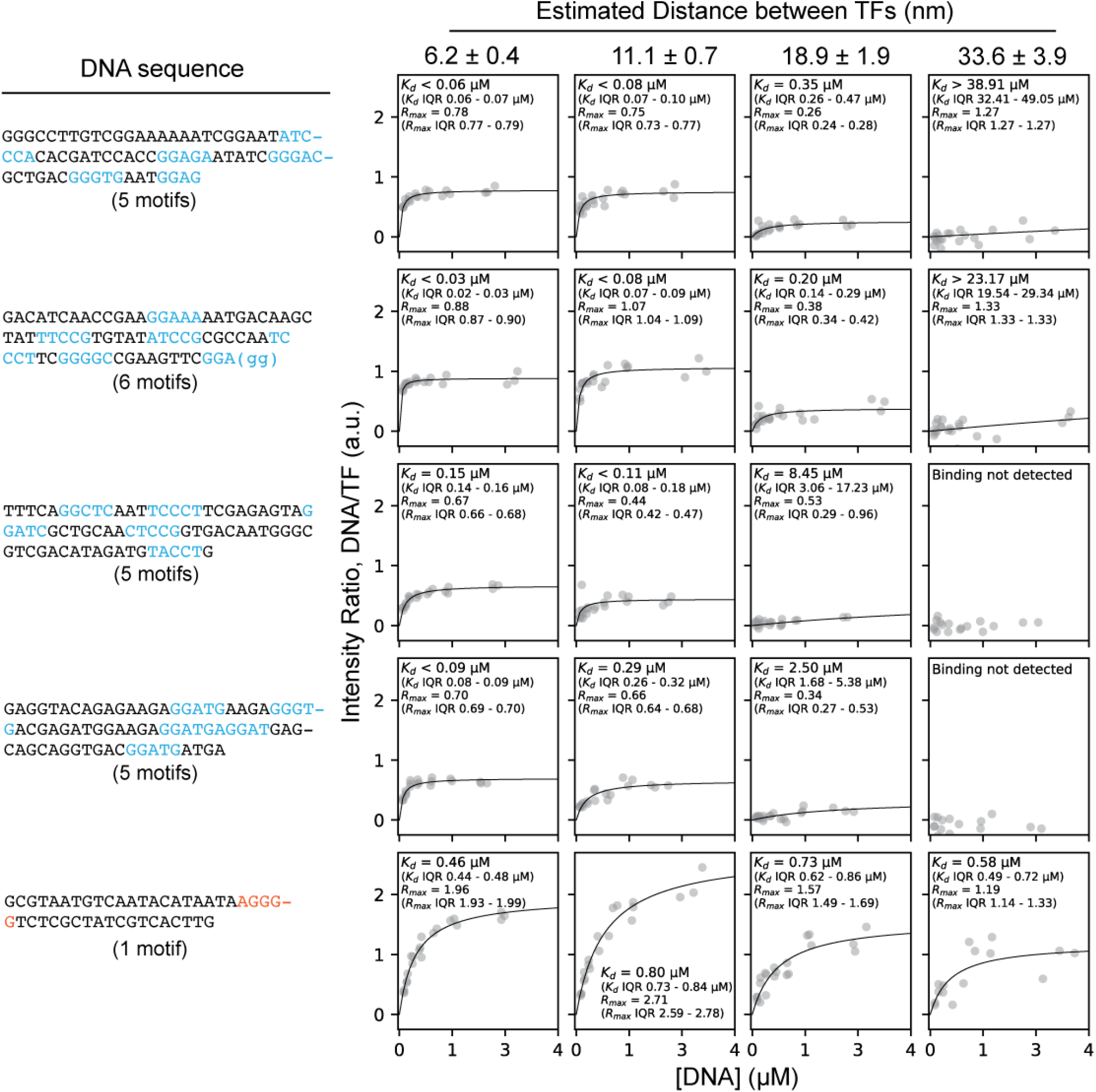
Example binding curves for the Msn2-DBD interacting with 5 different DNA sequences across different surface-immobilized TF densities. Each row shows a different DNA sequence with AGGGG motif variants (identified using FiNeMo) highlighted in blue (left) and measured intensity ratios (DNA/TF) as a function of soluble DNA concentration with associated Langmuir isotherm fits (right). Reported *K*_d_ and *R*_max_ values represent median fits for a given sequence across 400 subsamplings and fits to the data, and interquartile ranges of the fit distributions are reported for each (see **Methods**). The first and last sequences shown are the same as those shown in Figure 4D-E.

**Supplementary Figure 40.**
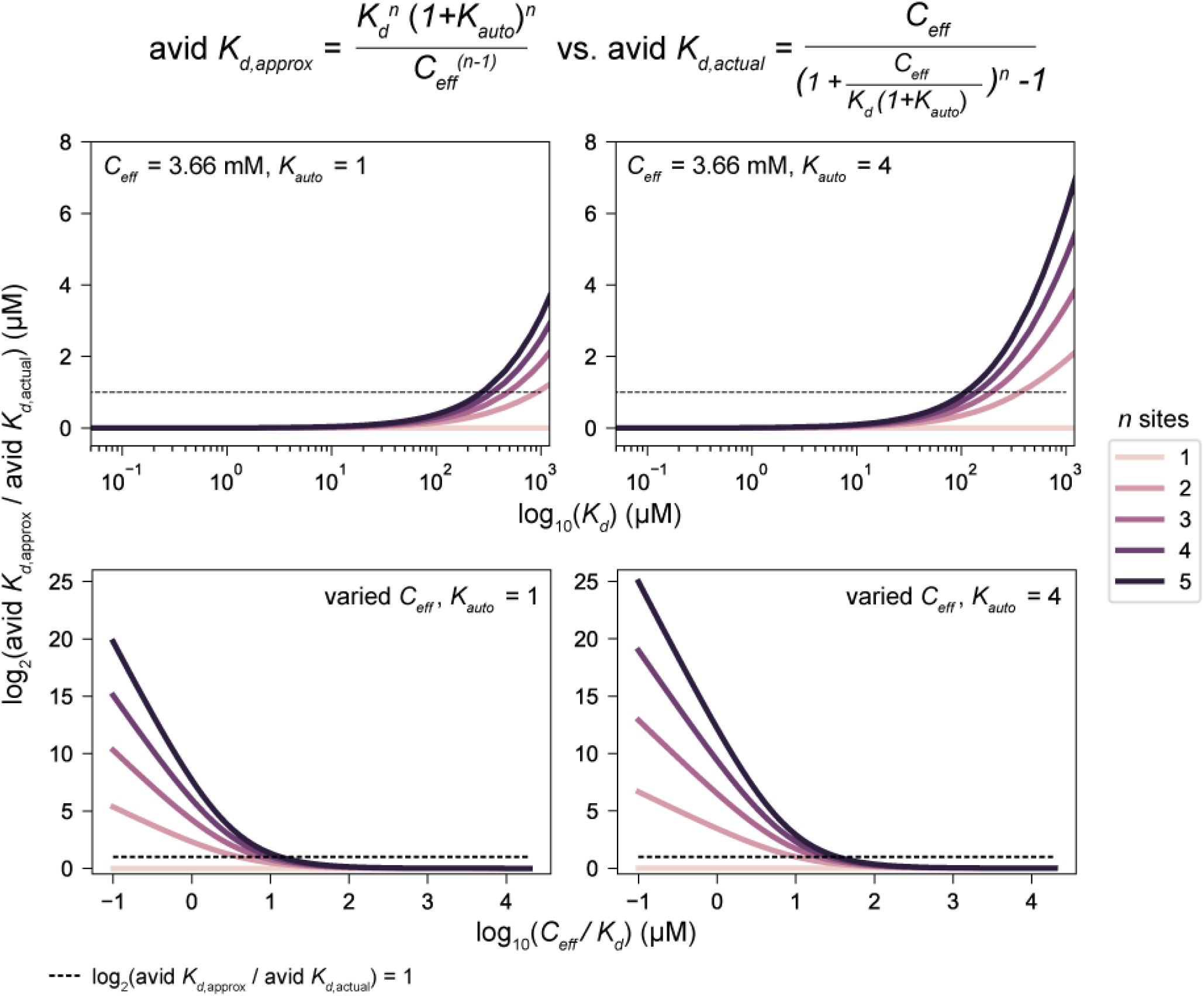
Analysis of when the simplifying assumption, *C*_eff_ >> *K*_d_, is valid for estimation of avid *K*_d_ with *n* binding sites and autoinhibition. To estimate the observed dissociation constant of *n* immobilized TFs to *n* identical binding sites on a DNA molecule under conditions of autoinhibition, we make the assumption *C*_eff_ >> *K*_d_ (where *C*_eff_ is the effective concentration of additional binding sites once the first of *n* binding sites is bound and *K*_d_ is the equilibrium dissociation constant describing binding of a TF to a single binding site). To clarify the regimes in which this assumption holds, we calculated the ratio of the approximated avid *K*_d_ (valid under *C*_eff_ >> *K*_d_) to the general solution for avid *K*_d_, here denoted by “actual”, (derived in **Supplementary Note 1**) across varied *C*_eff_*, K*_d_, and *K*_auto_ values. Here, subplots show calculated log_2_-transformed ratios as a function of log_10_(*K*_d_) (top row) or the log_10_-transformed ratio of *C*_eff_ to *K*_d_ (bottom row) for weaker (*K*_auto_ = 1, left column) and stronger (*K*_auto_ = 4, right column) autoinhibition. Line colors indicate the number of binding sites and TFs; dashed lines indicate a 2-fold difference between avid *K*_d,approx_ and avid *K*_d,actual_. For *K*_auto_ = 1, the fold change between avid *K*_d,approx_ and avid *K*_d,actual_ is less than 2 when *C*_eff_ is ∼5 and ∼14-fold greater than the single site *K*_d_ when binding to 2 or 5 sites, respectively. For *K*_auto_ = 4, the fold change between avid *K*_d,approx_ and avid *K*_d,actual_ is less than 2 when *C*_eff_ is ∼11 and ∼40-fold greater than the single site *K*_d_ when binding to 2 or 5 sites, respectively.

**Supplementary Figure 41.**
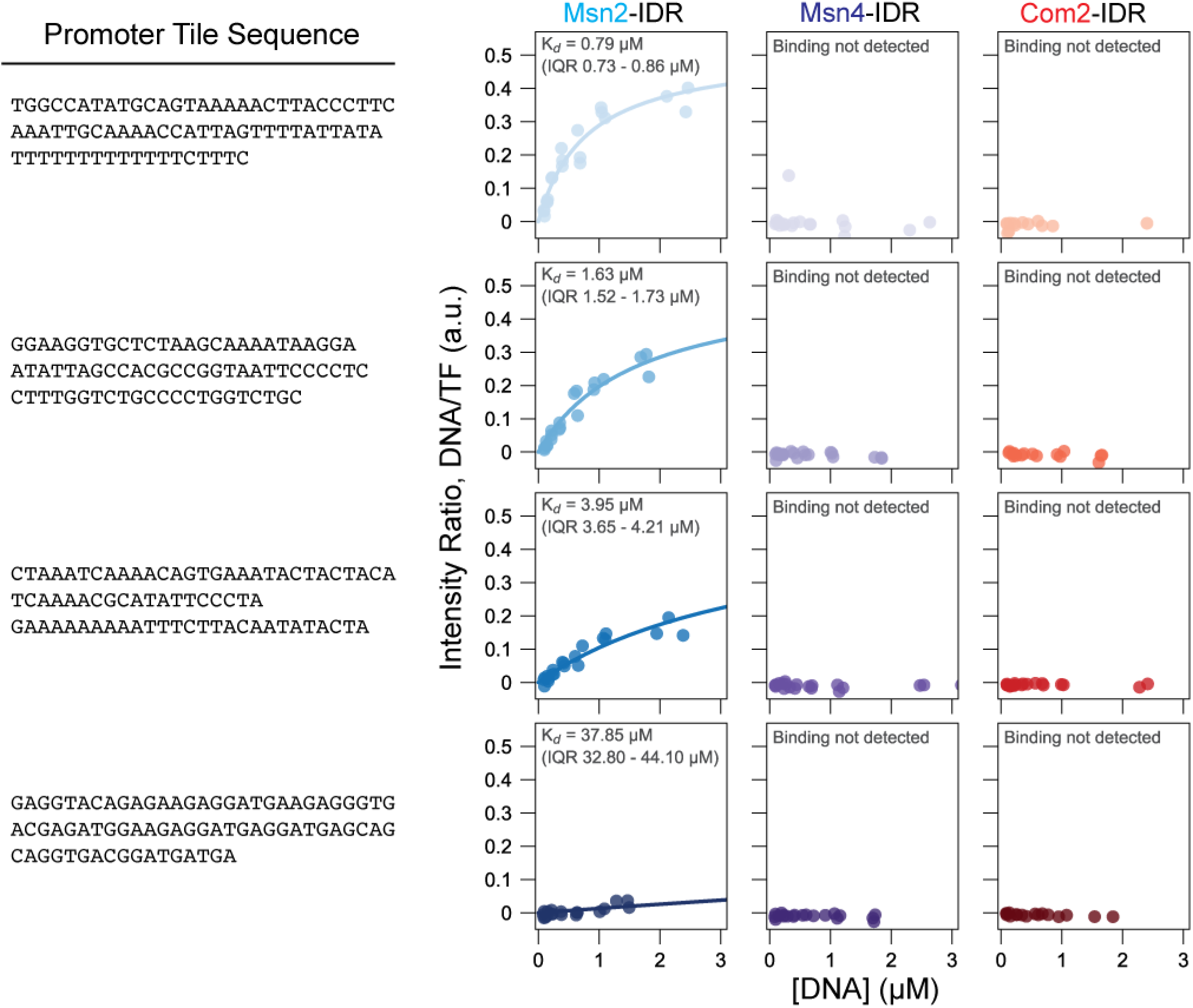
Representative binding curves of Msn2, Msn4, and Com2 IDRs to four promoter tiles (the same as those shown in Figure 5B) with fitted *K_d_*s. Rows indicate the promoter tile measured, with its sequence shown (left), and columns indicate the IDR-only construct measured. Markers indicate per-chamber ratios of summed background-subtracted DNA (Alexafluor647) to TF (meGFP) fluorescence intensities; lines indicate Langmuir isotherm fits with the annotated median fitted *K*_d_ across 400 subsamplings of the data per sequence; interquartile ranges of the fitted *K*_d_ distributions across subsamplings are annotated. Interactions for which no binding was detected are annotated (see **Methods**).

**Supplementary Figure 42.**
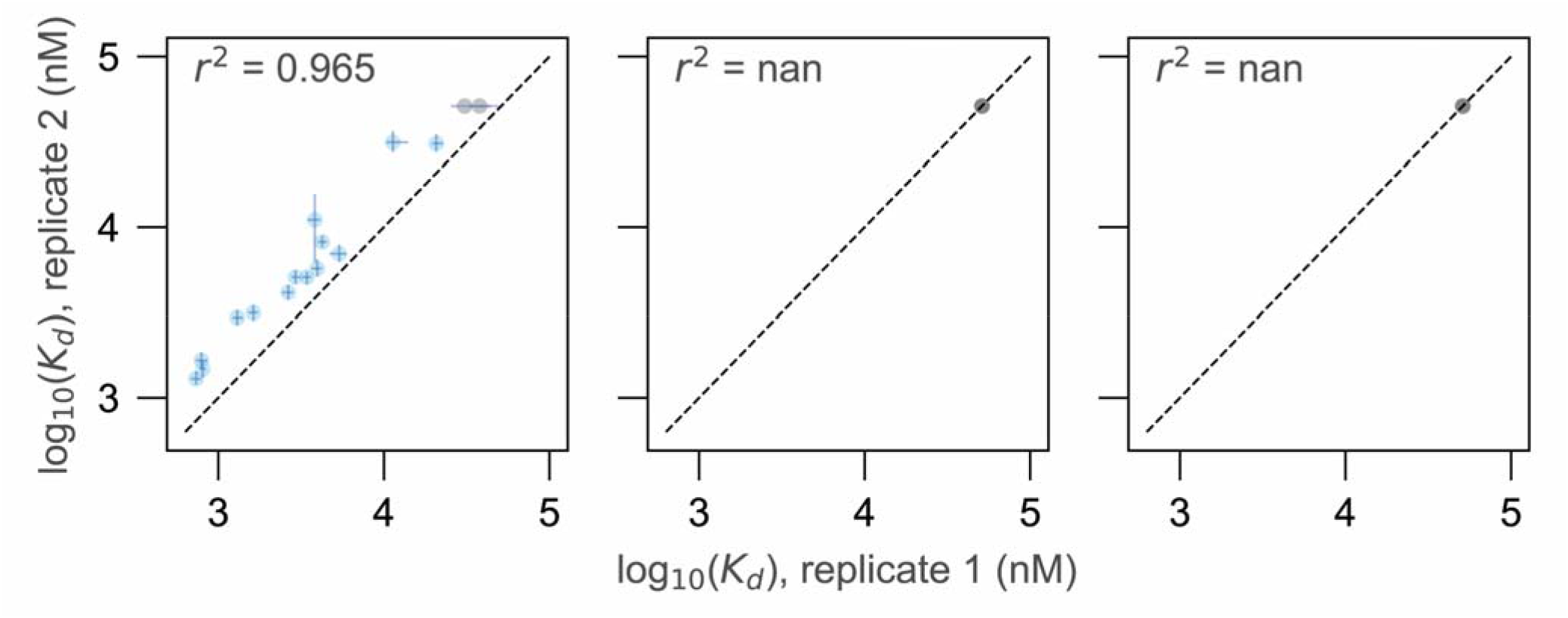
Replicate scatter plots of log_10_(*K*_d_) of Msn2, Msn4, and Com2 IDR binding to selected promoter tiles as measured by MITOMI. Dotted lines indicate 1:1 lines. Each marker indicates the median fitted *K*_d_ for a given promoter tile; error bars indicate the interquartile distance of the fitted *K*_d_ distributions across 400 subsamplings of the data. Blue points represent sequences whose fit *K*_d_s were within detection limits, while gray points represent TF-DNA interactions which were not detected in at least one replicate and the values of which were arbitrarily set to 51.1 µM (the value above which *K*_d_s are called as lower limits for this experiment, **Methods**) for the sake of visualization.

**Supplementary Figure 43.**
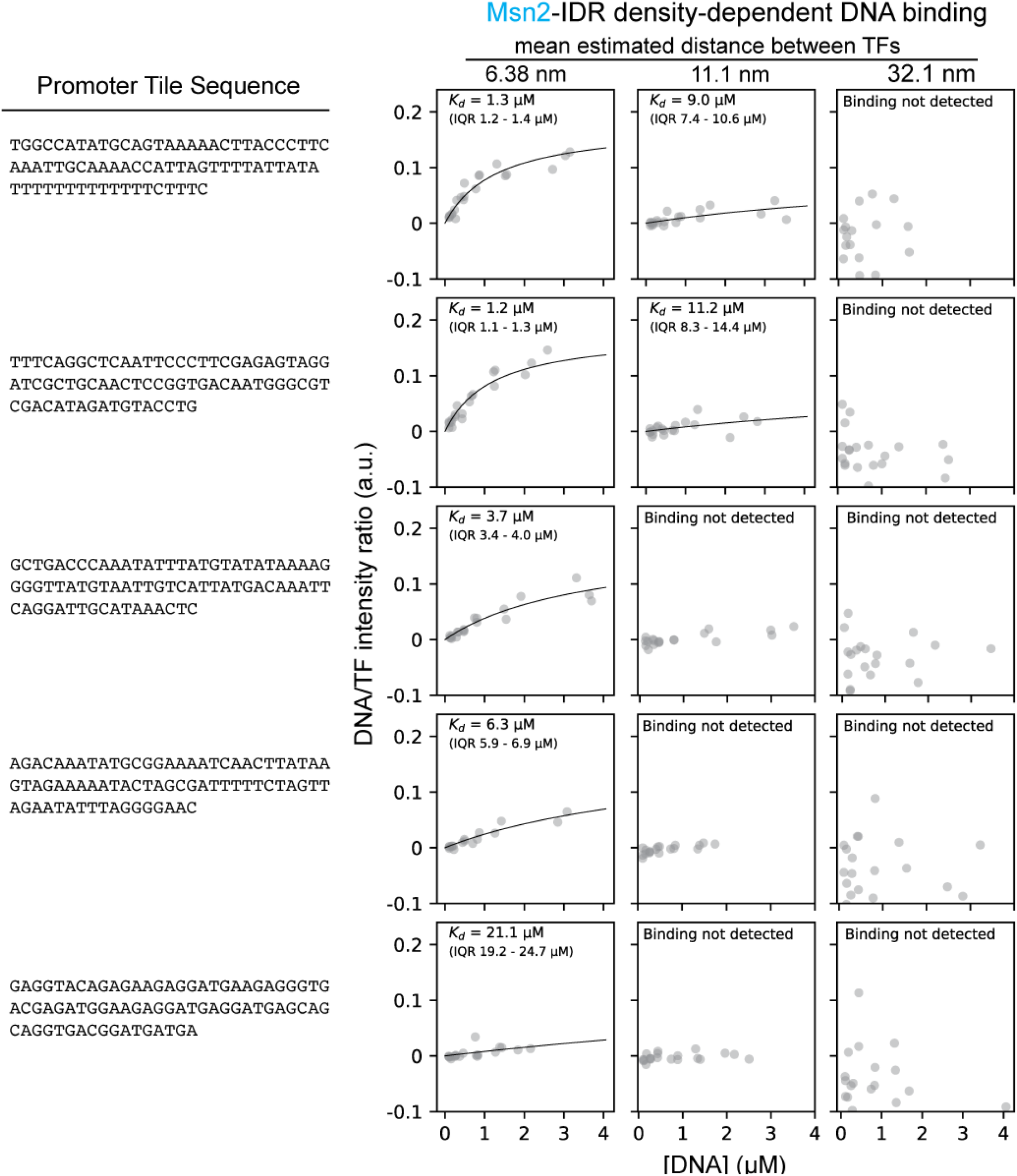
Representative binding curves of Msn2’s IDR to example promoter tiles across a range of patterning densities. Rows indicate the promoter tile measured, with its sequence shown (left), and columns indicate the density at which binding was measured with the average estimated distance between TF molecules denoted. Markers indicate per-chamber ratios of summed background-subtracted DNA (Alexafluor647) to TF (meGFP) fluorescence intensities; lines indicate Langmuir isotherm fits with the annotated median fitted *K*_d_ across 400 subsamplings of the data per sequence; interquartile ranges of the fitted *K*_d_ distributions across subsamplings are annotated for detected interactions (see **Methods**).

**Supplementary Figure 44.**
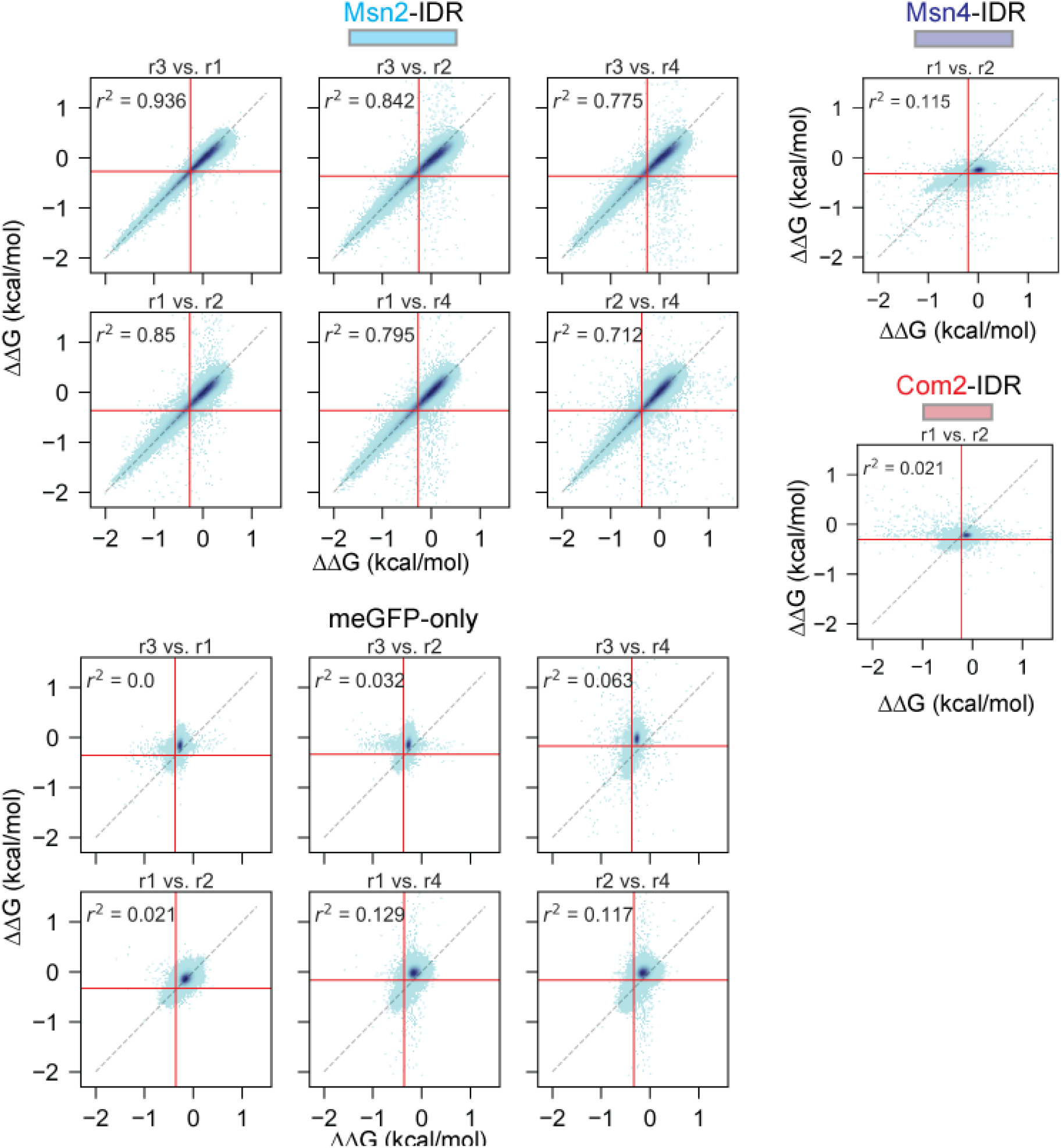
2D histograms comparing measured ‘promoter tile’ library ΔΔG values for each IDR-only construct across 2 technical replicates. Sequences were binned by the ΔΔGs of the replicates indicated (computed relative to TGCAATTACGTAATTTTAGCCGCTGAGAAGCGGATGGTAATGAGACAAGTTGATATCAAACAGATACATATTTAA) with pixel color indicating density of sequences per bin. Dotted black lines indicate 1:1 lines; red lines indicate thresholds of 2 standard deviations from noise mean values (calculated by fitting a Gaussian curve to a synthetic ‘noise’ distribution of each experiment constructed by taking the ΔΔG values greater than the empirical center and their reflection across that center value –Methods). Pearson *r*^2^ values shown were calculated from ΔΔG values for sequences lying outside of 2 standard deviations from the noise of either replicate.

**Supplementary Figure 45.**
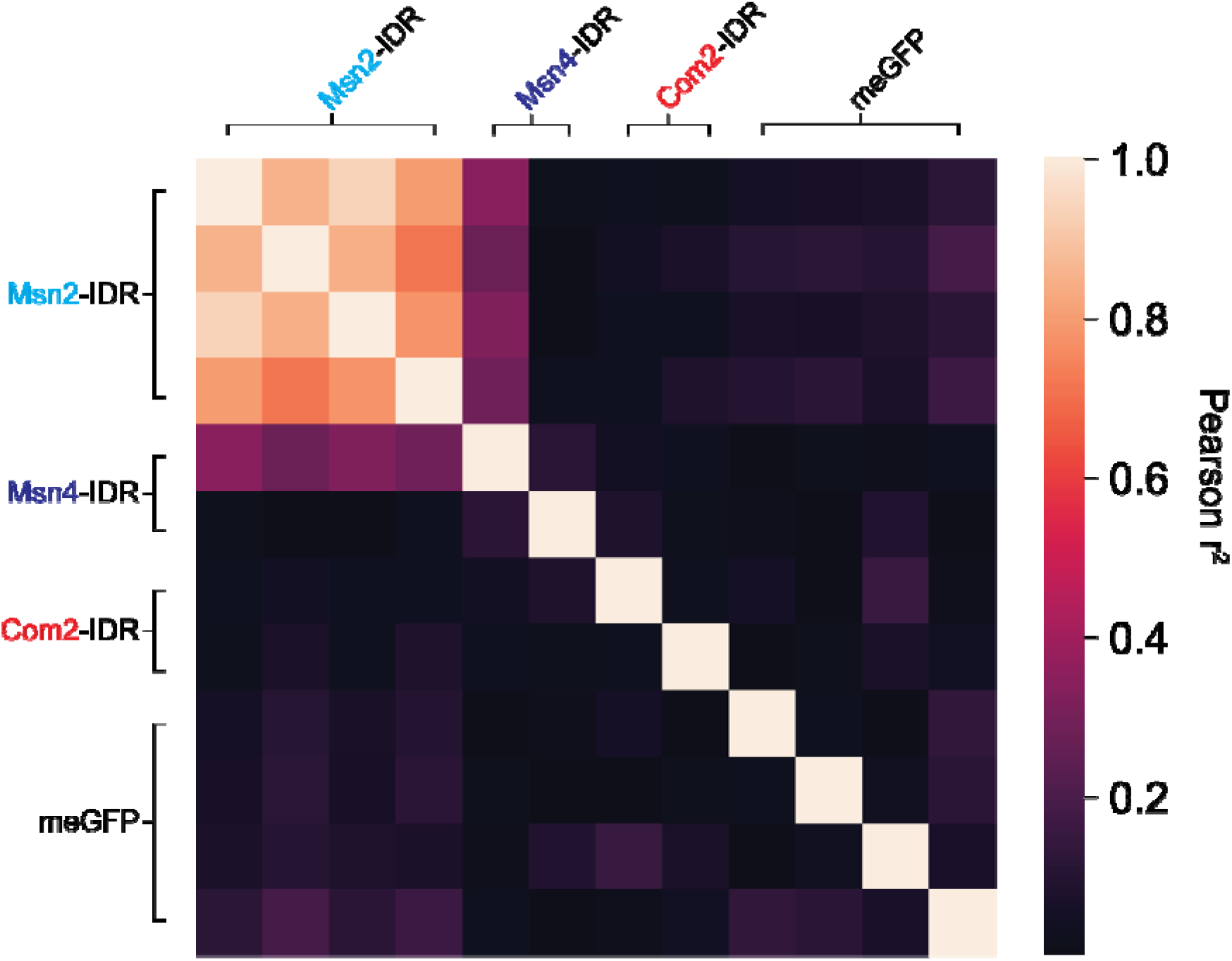
Heatmap of pairwise Pearson r^2^ values for ΔΔGs for promoter tiles between all replicates of IDR-only experiments. Pearson r^2^ values are reported for ΔΔG measurements outside of the noise of either replicate (defined by using a threshold of 2 standard deviations above the mean of the noise distribution, see **Methods**).

**Supplementary Figure 46.**
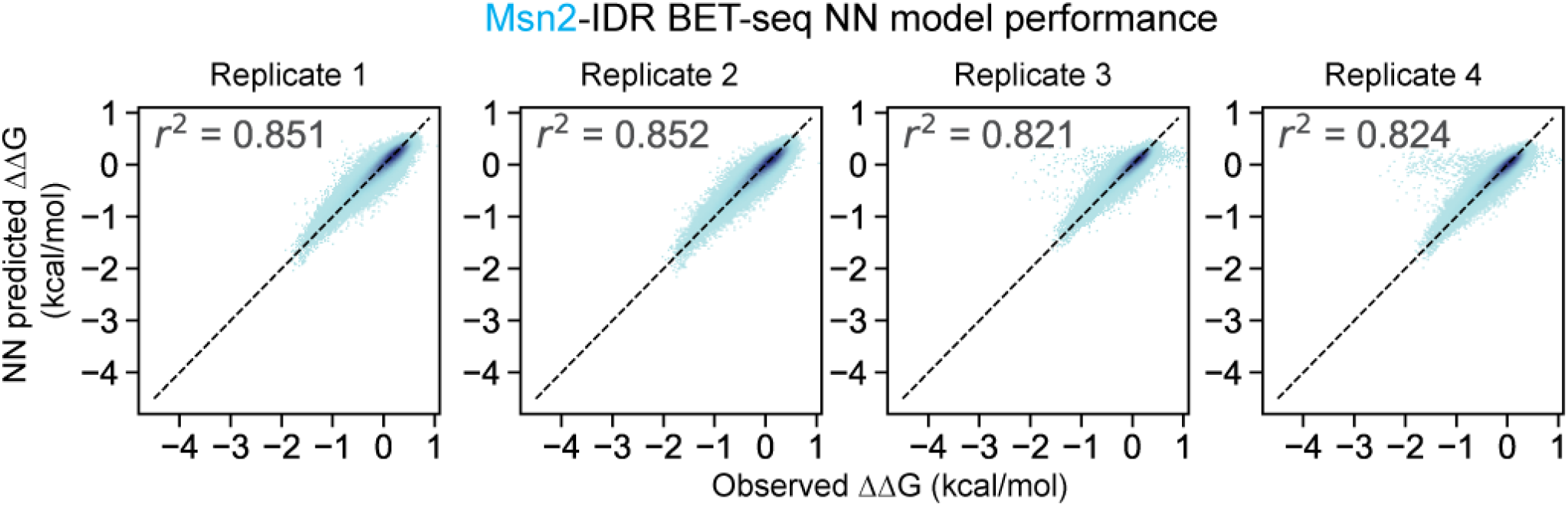
2D histograms showing neural network model-predicted ΔΔGs versus observed ΔΔGs for each Msn2-IDR BET-seq replicate measurement of the ‘promoter’ library. Library sequences were binned by observed and predicted ΔΔG and pixel color indicates density of sequences per bin. For each replicate, test-set predictions from all 5-folds are combined here. Dotted black lines indicate 1:1 line.

**Supplementary Figure 47.**
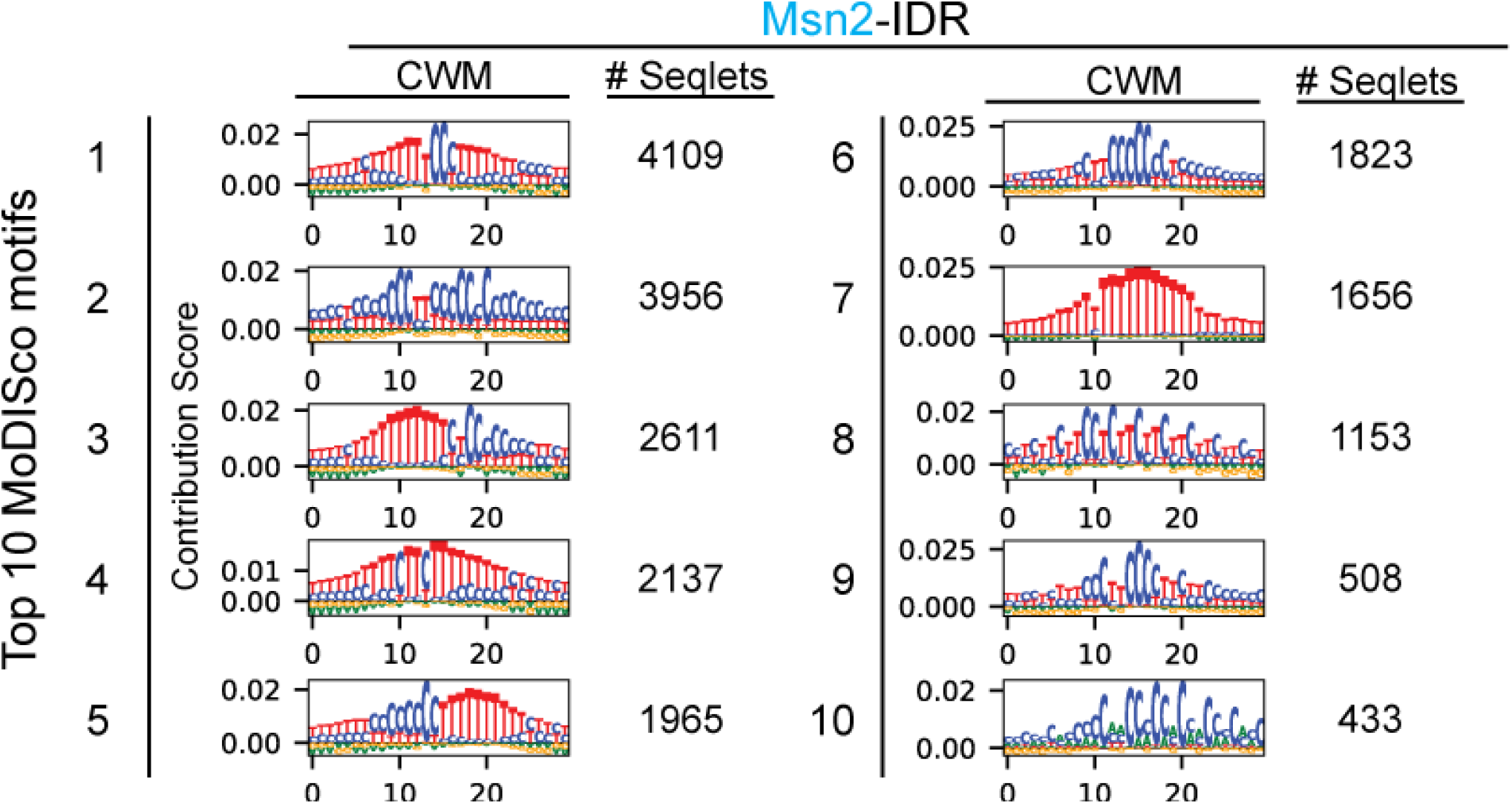
Top 10 MoDISco CWM motifs found in models of ‘promoter’ library binding by Msn2-IDR. Count column values represent the number of seqlets assigned to a given motif cluster.

**Supplementary Figure 48.**
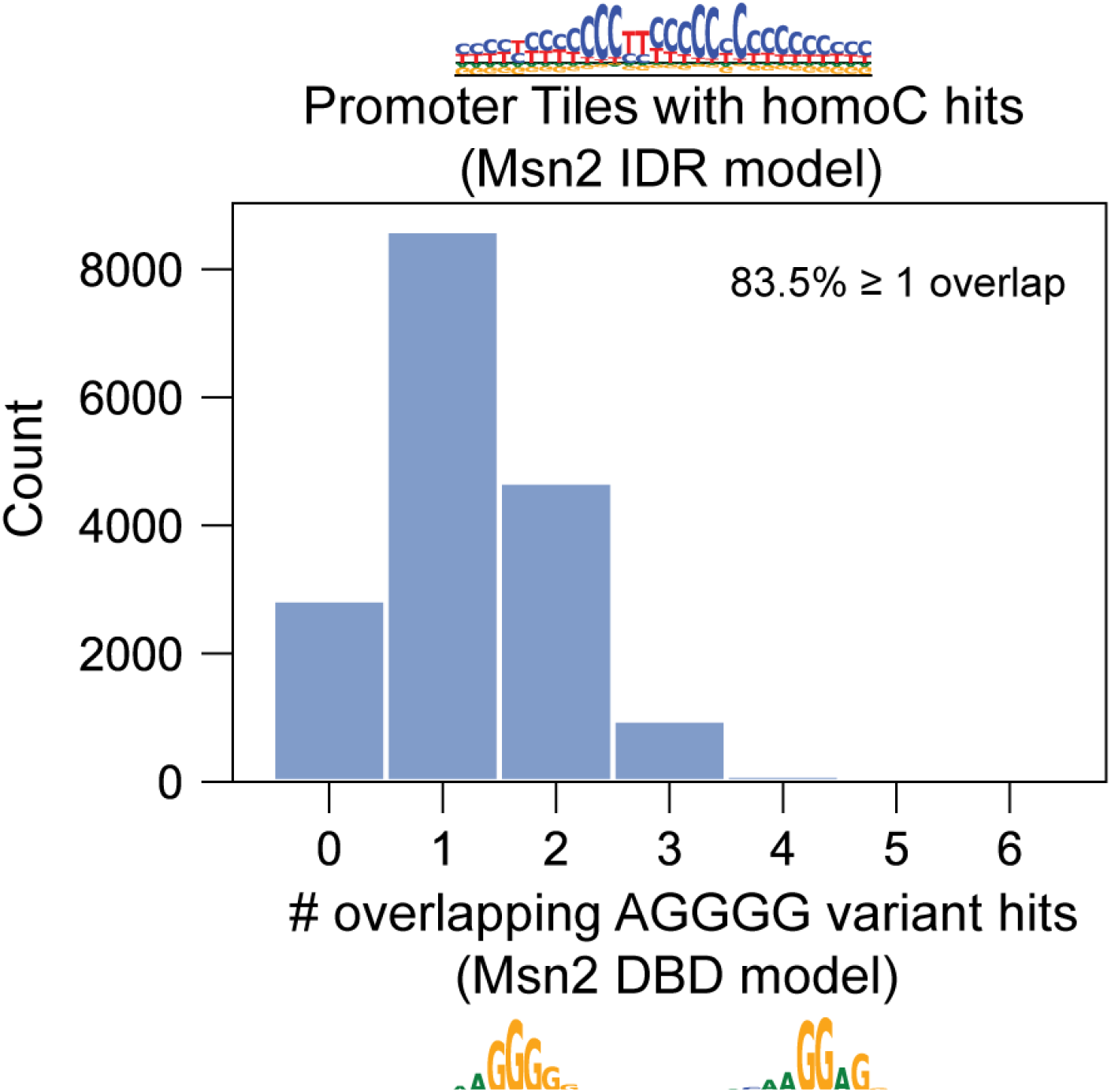
Overlap of AGGGG motif variants with Msn2-IDR homoC motif. Histogram of the number of AGGGG motif variant hits from the Msn2-DBD AffinityNet model that overlapped with FiNeMo homoC matches in the Msn2-IDR AffinityNet model.

**Supplementary Figure 49.**
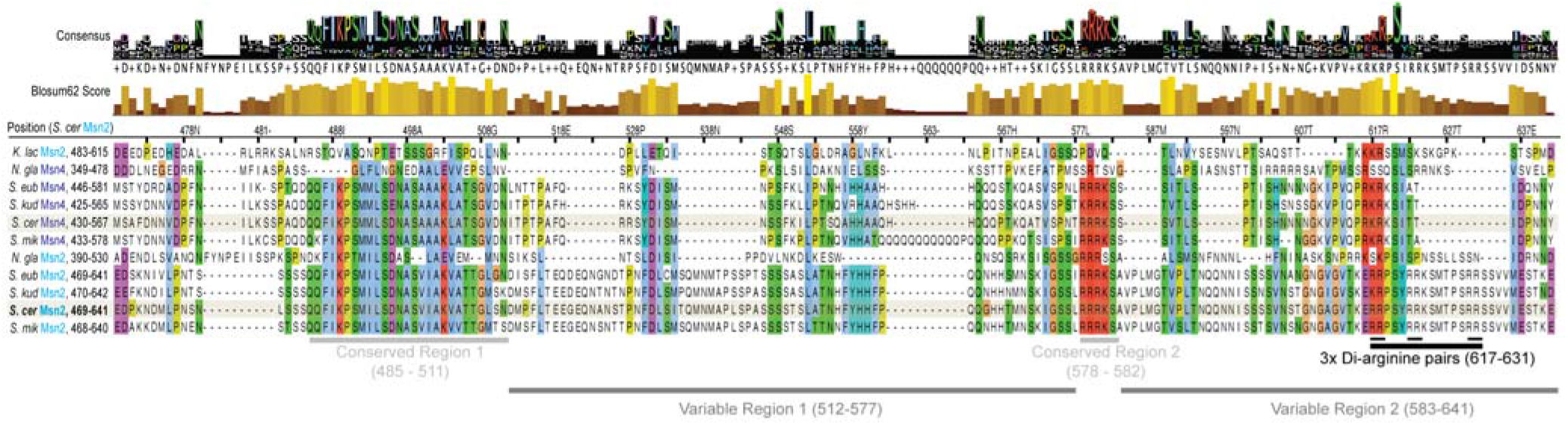
MUSCLE alignment of Msn2 and Msn4 ortholog protein sequences from *K. lactis*, *N. glabratus*, *S. eubayanus*, *S. kudriavzevii*, *S. mikatae*, and *S. cerevisiae* centered on *S. cerevisiae* Msn2’s 469-641 IDR region. Top rows indicate consensus residues (with heights proportional to the percentage of orthologs with a given residue) and Blosum62 conservation scores for each position. Highlighted sequences indicate *S. cerevisiae* Msn2 and Msn4; all residues are colored using the Clustal coloring scheme. Underlines indicate two regions that differ in conservation between Msn2 and Msn4 orthologs and the three di-arginine (RR) pairs that appear in Msn2 orthologs but not in Msn4 orthologs.

**Supplementary Figure 50.**
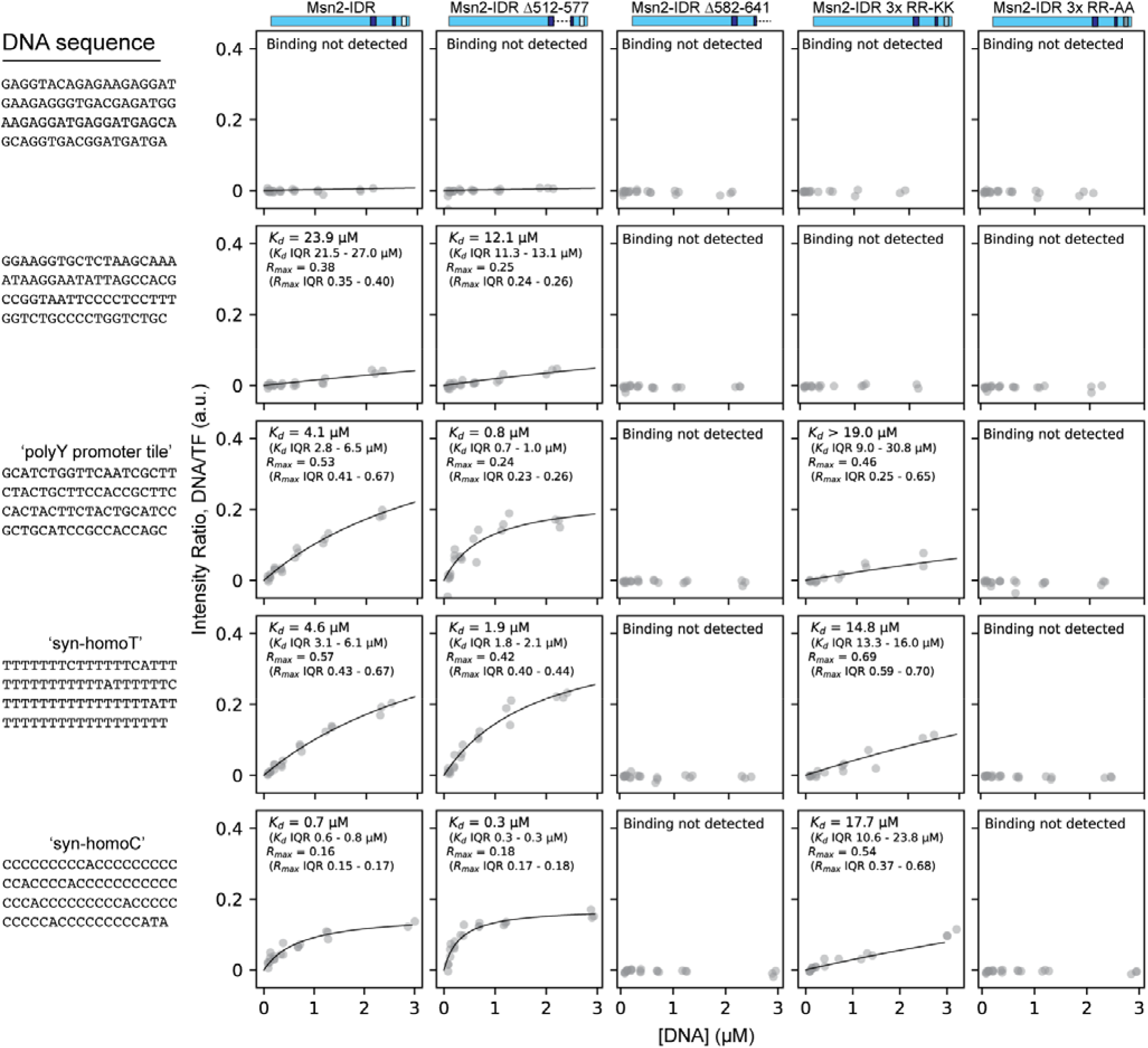
Representative MITOMI binding curves of Msn2’s IDR and derived mutants tested in Figure 5J-K. Gray points represent fractional binding in the form of DNA fluorescence (Alexafluor 647) normalized to TF fluorescence (meGFP) of indicated constructs to indicated DNA sequences. Black lines represent binding isotherms fitted to each dataset with the fitted parameters of *K*_d_ and R_max_ (the saturating fractional occupancy) shown for each curve with interquartile ranges of fits across 400 subsamplings. Columns represent binding for the Msn2-IDR mutant constructs shown (left to right: Msn2-IDR, Msn2-IDR Δ512-577, Msn2-IDR Δ582-641, Msn2-IDR 3xRR-KK, and Msn2-IDR 3xRR-AA). Rows represent binding to the indicated DNA sequences. A ‘global’ R_max_ fitting algorithm was used to fit R_max_ values to the first two sequences shown since they do not show saturation, and a ‘local’ fitting algorithm was used to fit R_max_ values to the last three sequences shown since, for the constructs that bound DNA tightly, they clearly showed saturation but often at visually distinct saturation points (**Methods**).

**Supplementary Figure 51.**
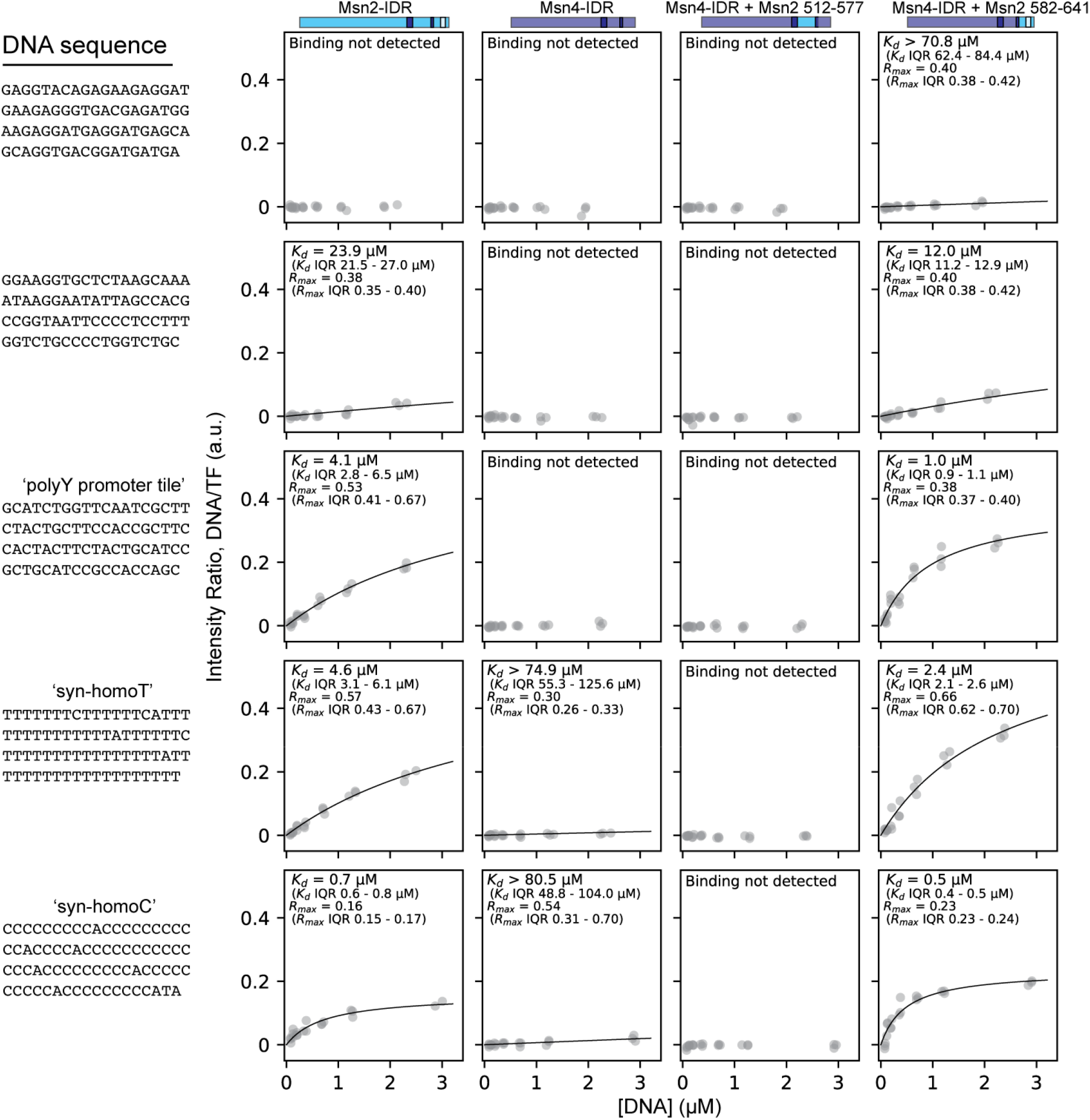
Representative MITOMI binding curves of Msn2’s IDR and chimeric Msn4-IDR mutants tested in Figure 5J-K. Gray points represent fractional binding in the form of DNA fluorescence (Alexafluor 647) normalized to TF fluorescence (meGFP) of indicated constructs to indicated DNA sequences. Black lines represent binding isotherms fitted to each dataset with the fitted parameters of *K*_d_ and R_max_ (the saturating fractional occupancy) shown for each curve with interquartile ranges of fits across 400 subsamplings. Columns represent binding for the constructs shown (left to right: Msn2-IDR, Msn4-IDR, Msn4-IDR + Msn2 512-577, Msn4-IDR + Msn2 582-641). Rows represent binding to the indicated DNA sequences. A ‘global’ R_max_ fitting algorithm was used to fit R_max_ values to the first two sequences shown since they do not show saturation, and a ‘local’ fitting algorithm was used to fit R_max_ values to the last three sequences shown since, for the constructs that bound DNA tightly, they clearly showed saturation but often at visually distinct saturation (**Methods**)

**Supplementary Figure 52.**
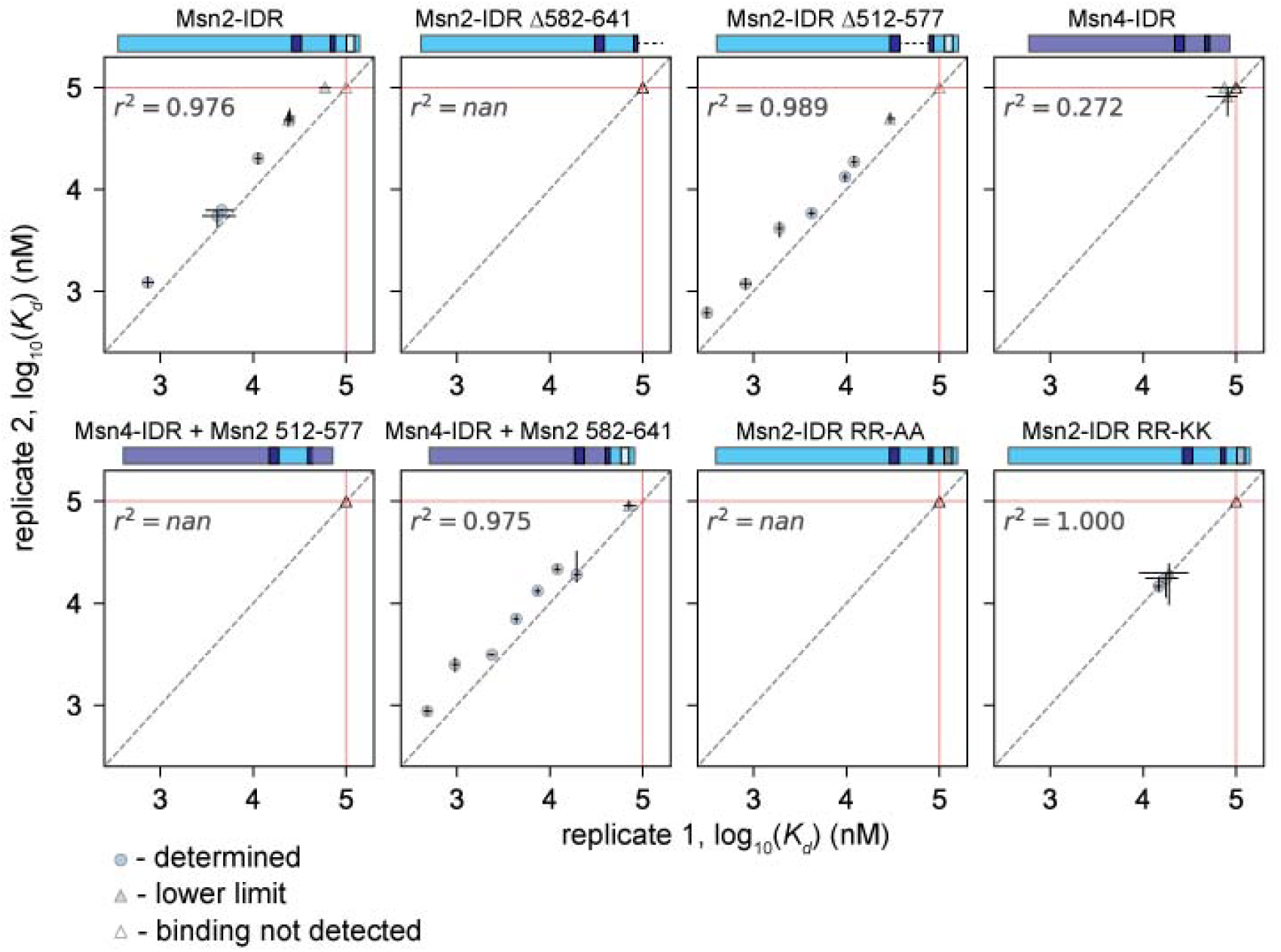
Scatter plots comparing fitted *K*_d_s between replicates of MITOMI measurements of Msn2 and Msn4 IDR mutants. Each subplot compares 2 technical replicates for the indicated construct interacting with selected promoter tiles and synthetic homoT and homoC sequences (summarized in Figure 5J-K). Each marker indicates the log-transformed median fitted *K*_d_ value across 400 subsamplings of the data per sequence in a given experiment; blue circles indicate determined *K*_d_ values, gray triangles indicate interactions for which *K*_d_ values represent lower limits in at least one replicate, and white triangles indicate interactions for which binding was not detected (with *K*_d_ values set to 100 µM for the sake of visualization) error bars indicate log-transformed interquartile ranges of the *K*_d_ fits across subsamplings. Black dotted lines indicate 1:1 lines; red dotted lines indicate linear regression fits with the annotated correlation coefficients. Solid red lines indicate upper *K*_d_ fitting limit.

**Supplementary Figure 53.**
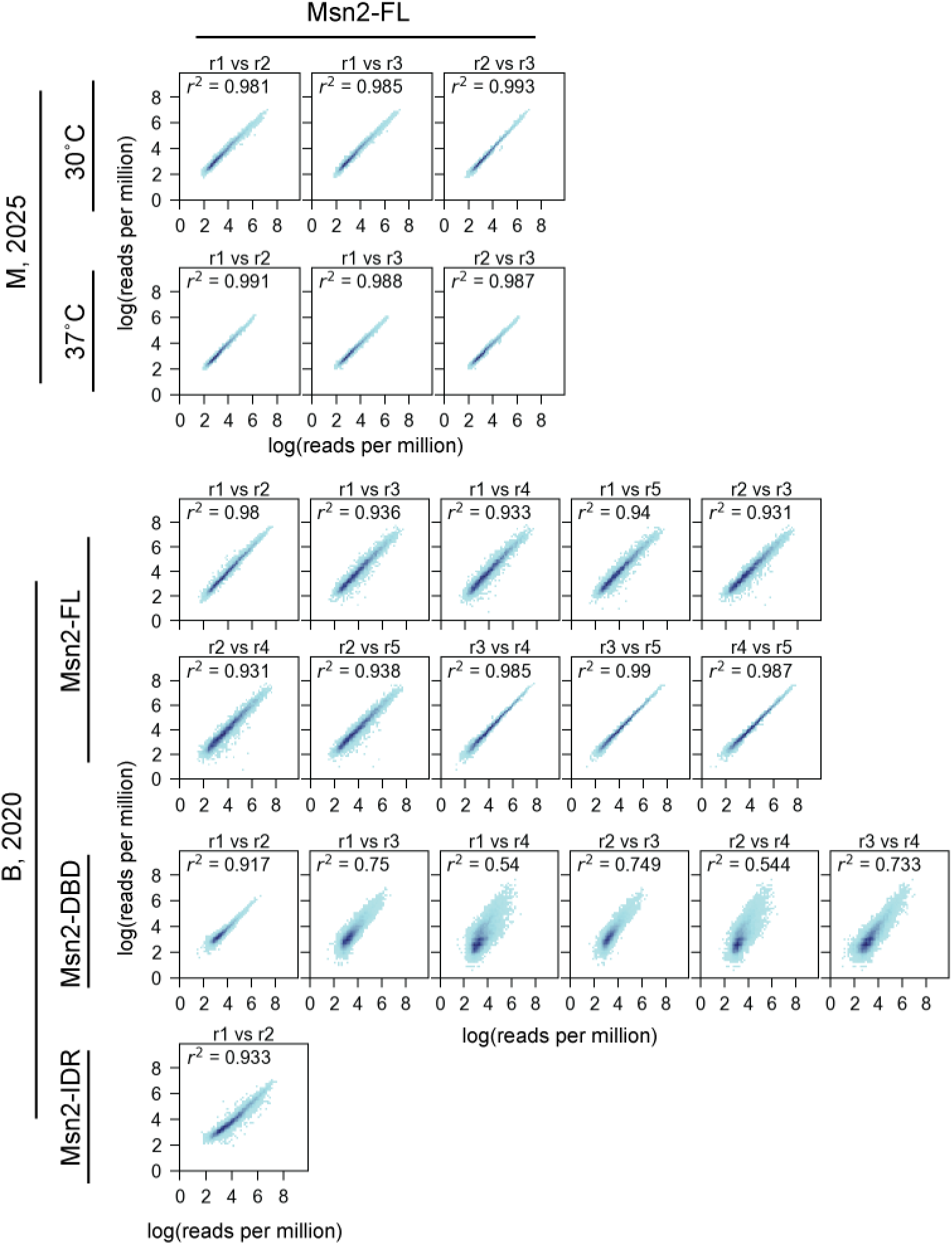
2D histograms of ChEC-seq normalized unstranded read counts within peak regions between experimental replicates for both Brodsky et al., 2020 (B, 2020) and Mahendrawada et al., 2025 (M, 2025) datasets. Peak regions were binned by normalized log-transformed read counts for each replicate comparison shown; pixel color indicates density of peaks per bin. Specific replicate pairs plotted are indicated per plot and Pearson r^2^ is reported per replicate pair. Rows indicate the construct and dataset.

**Supplementary Figure 54.**
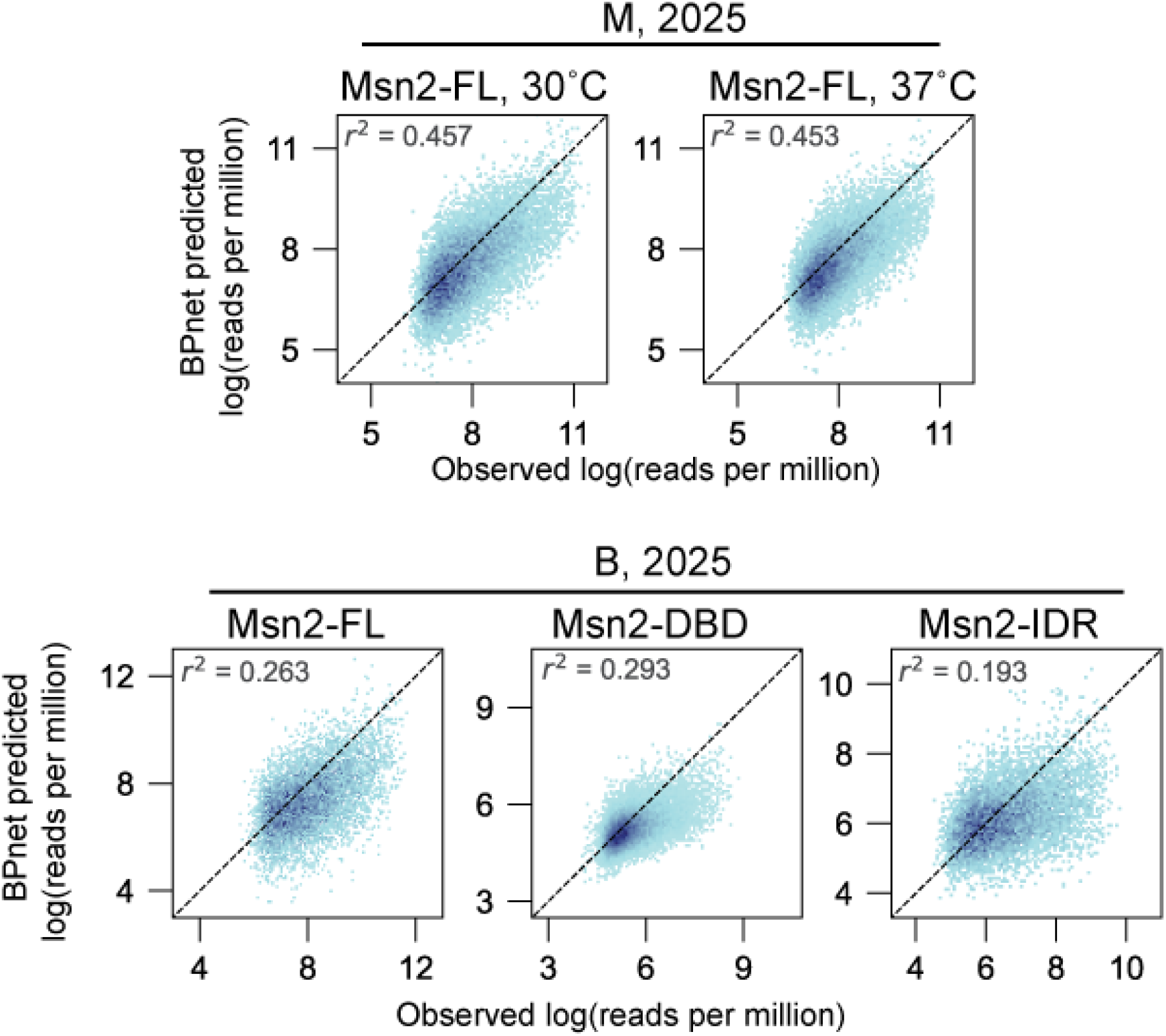
BPNet model performance for ChEC-seq datasets over peak regions. 2D histograms of observed vs. BPNet-predicted log-transformed read counts within peak regions for ChEC-seq binding data of Msn2 constructs from Mahendrawada et al., 2025 (M, 2025) and Brodsky et al., 2020 (B, 2020) datasets. Peak regions were binned by observed and predicted log-transformed read counts, and pixel color indicates density of peaks per bin. Pearson r^2^ is reported per construct. Dotted black lines indicate 1:1 lines.

**Supplementary Figure 55.**
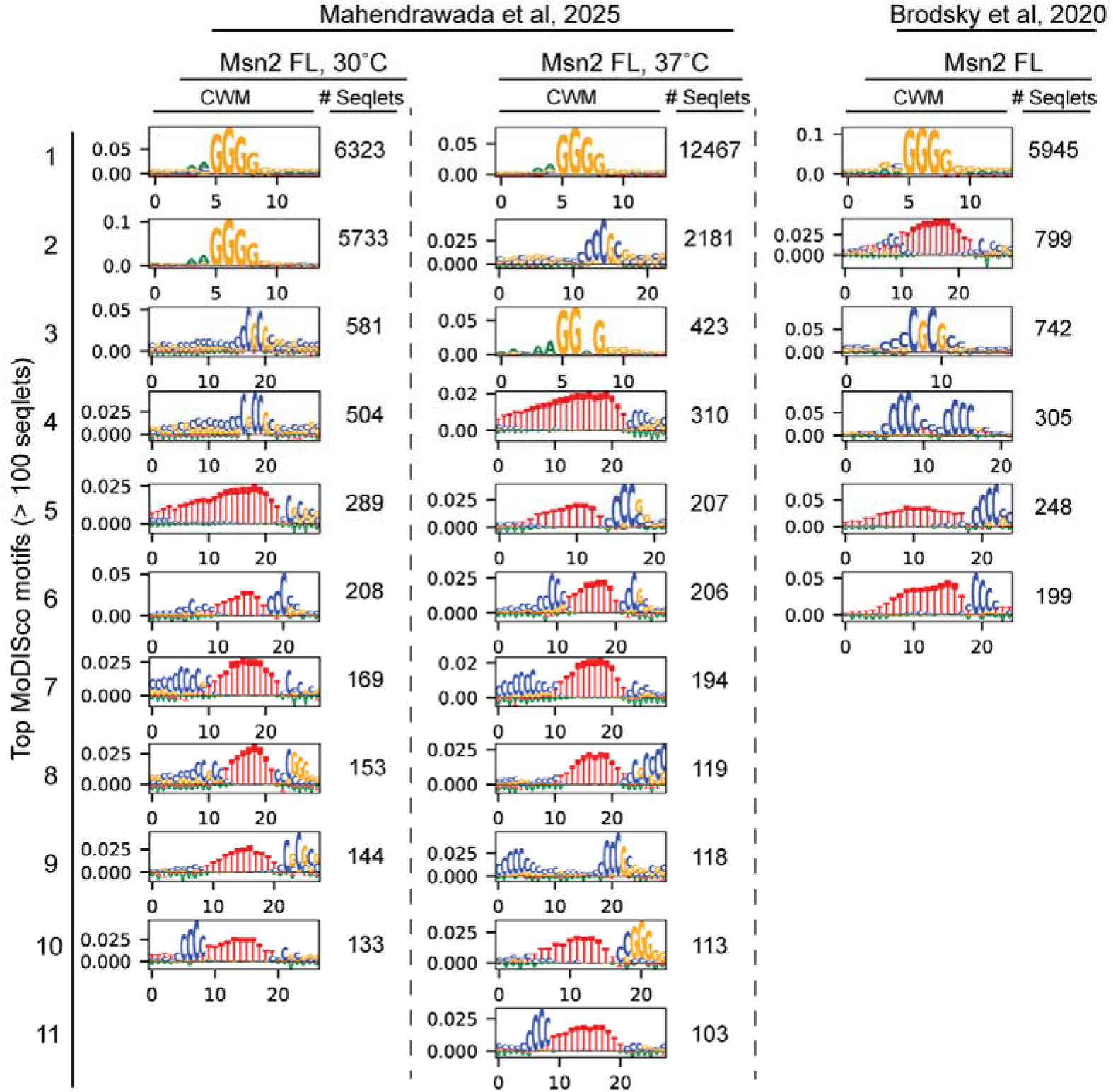
MoDISco CWMs discovered through BPNet count heads for Msn2-FL experiments from both Mahendrawada et al., 2025 and Brodsky et al., 2020 datasets. All CWMs with greater than 100 seqlets are reported.

**Supplementary Figure 56.**
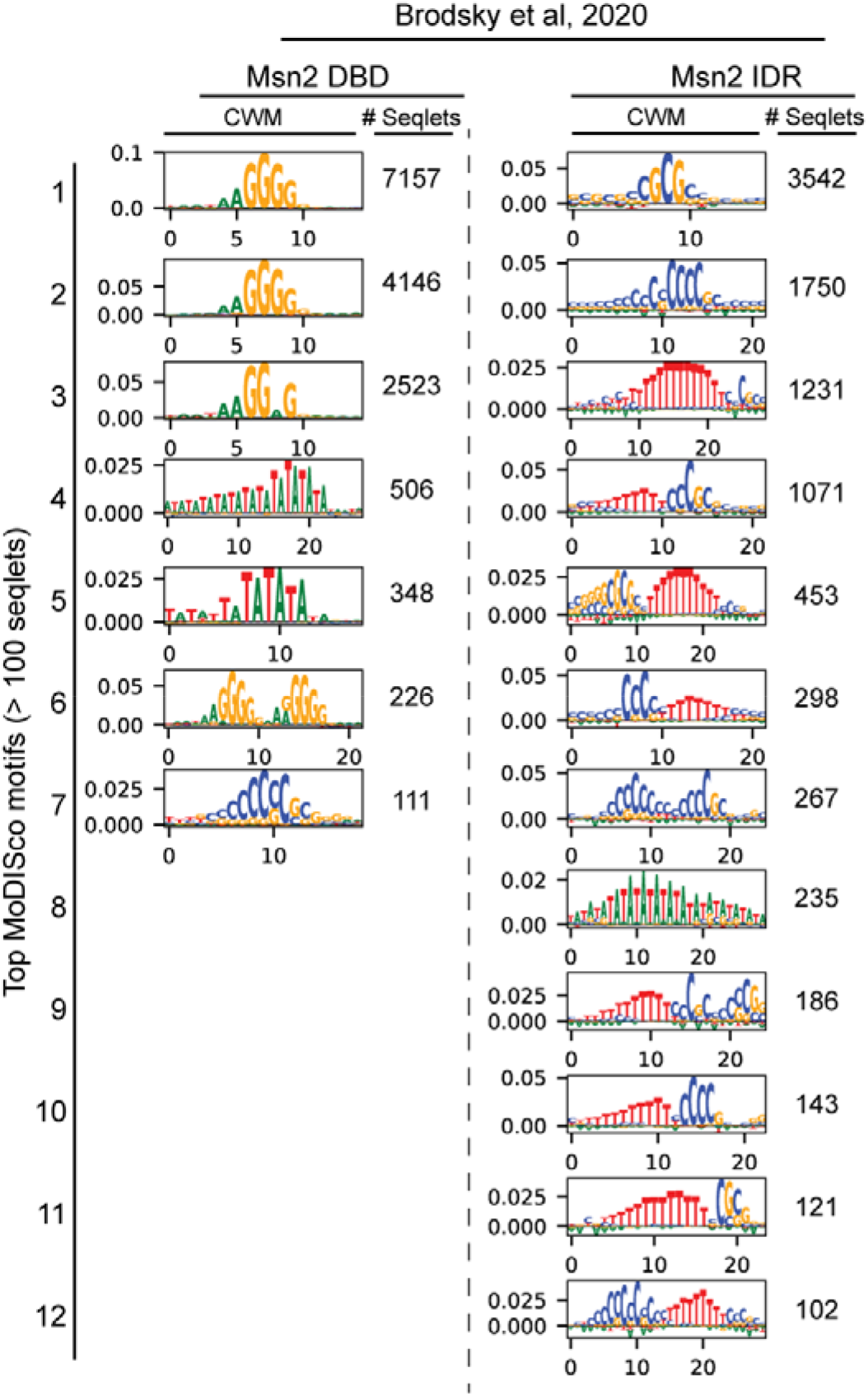
MoDISco CWMs discovered through BPNet count heads for Msn2-DBD and IDR experiments from the Brodsky et al., 2020 dataset. All CWMs with greater than 100 seqlets are reported.

**Supplementary Figure 57.**
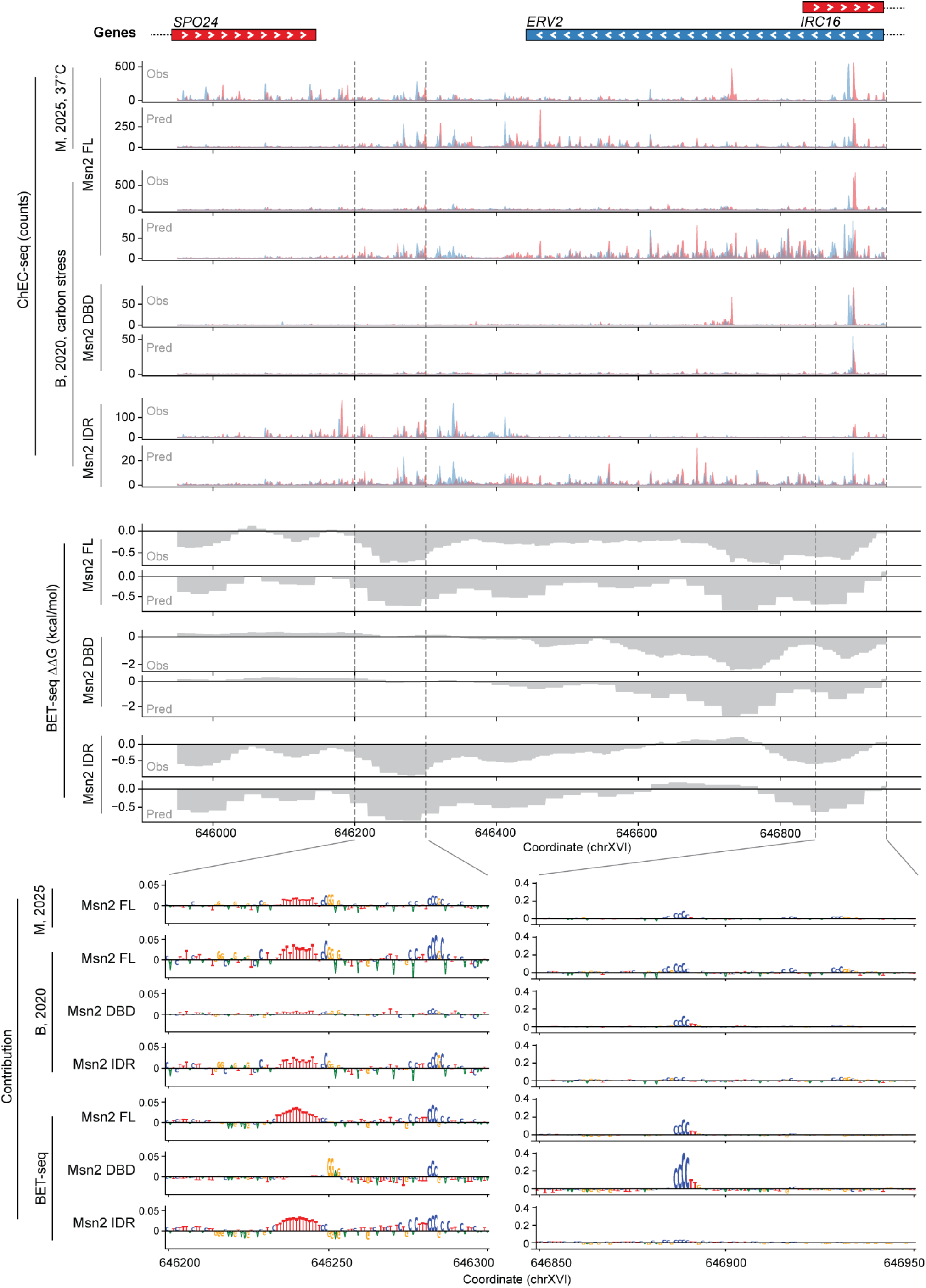
Binding of FL, DBD-only, and IDR-only Msn2 constructs *in vivo* and *in vitro* to the ERV2 locus. (Top) Genome browser-style plots of observed and neural network-predicted ChEC-seq counts and BET-seq ΔΔGs over the *ERV2* region (chrXVI:645,950-646,950) for the following datasets and constructs: ChEC-seq - M, 2025 (37°C) Msn2-FL and B, 2020 Msn2-FL, Msn2-DBD, and Msn2-IDR; BET-seq – Msn2-FL, Msn2-DBD, and Msn2-IDR. The plus and minus strand counts are colored as red and blue, respectively. (Bottom) Zoomed in plots of BPNet and AffinityNet per-nucleotide contribution scores (for ChEC-seq and BET-seq samples respectively).

**Supplementary Figure 58.**
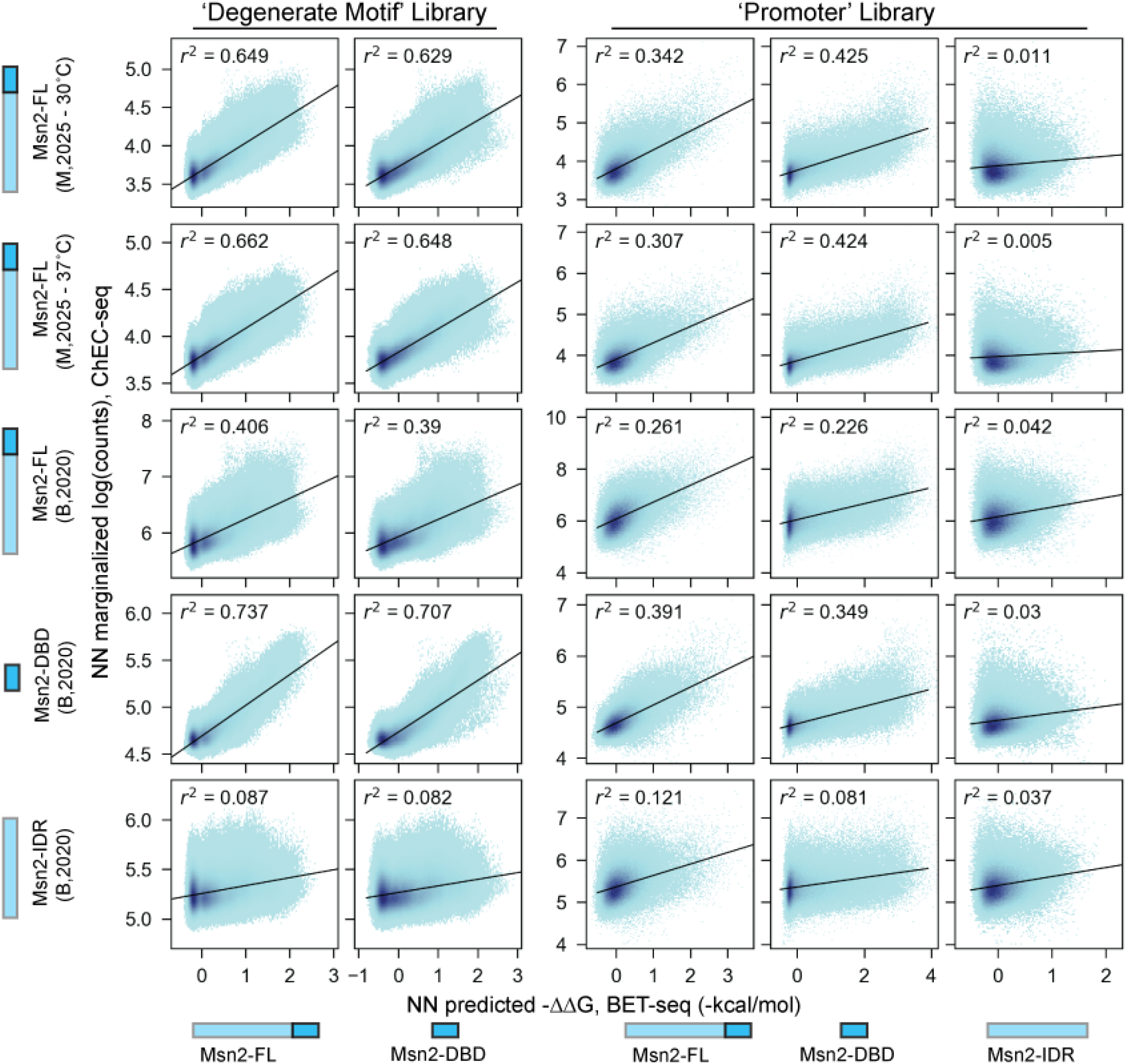
2D histograms of AffinityNet-predicted BET-seq -ΔΔGs versus marginalized BPNet-predicted log counts of ‘degenerate motif’ and ‘promoter’ library sequences for Msn2-FL and DBD constructs. Rows correspond to *in vivo* models and columns correspond to *in vitro* models; the left two columns compare predictions for the ‘degenerate motif’ library and the right three columns compare predictions for the ‘promoter’ library. Point color indicates data density per bin. Black lines indicate linear regression fit line.

**Supplementary Figure 59.**
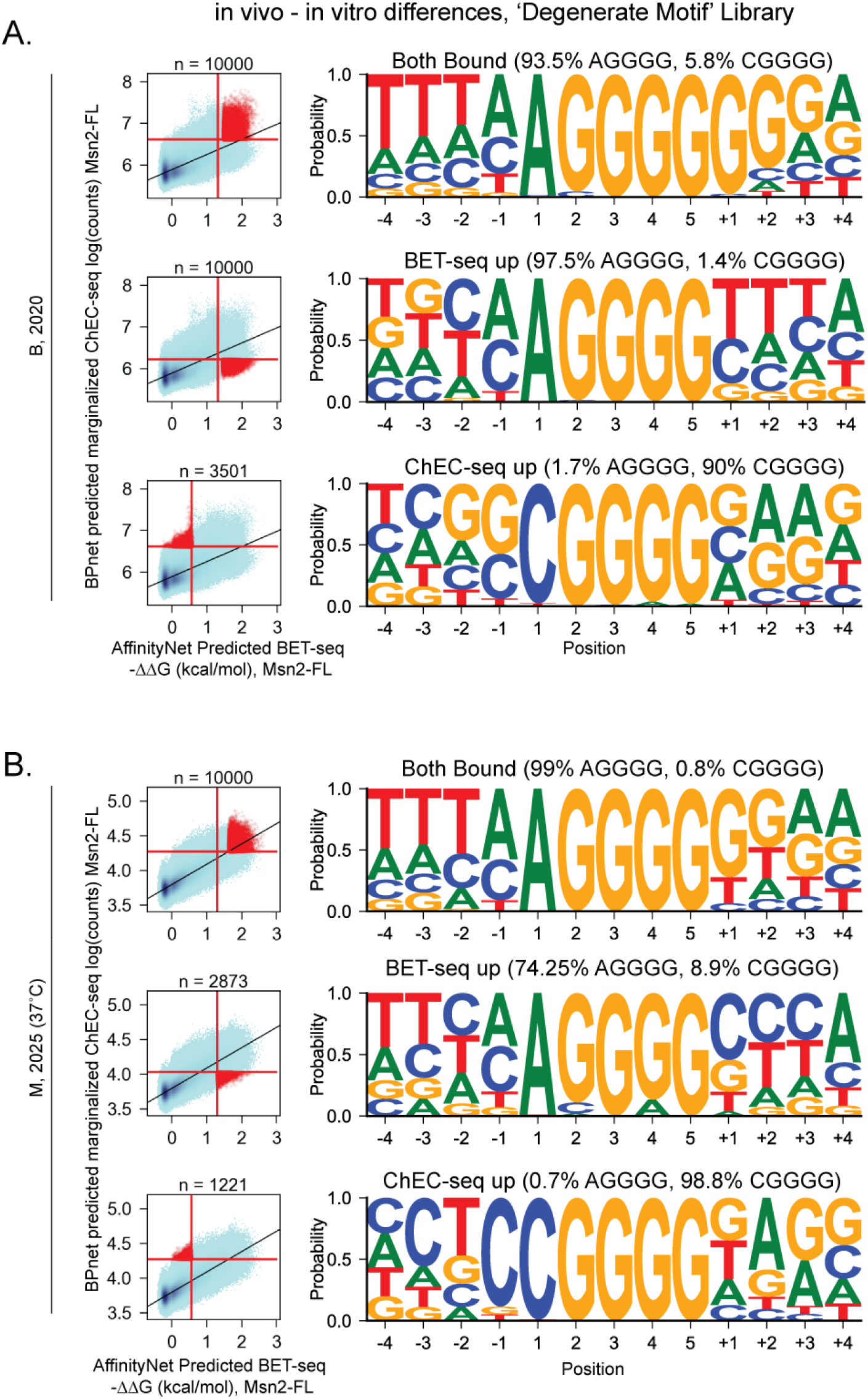
Analysis of differences between BET-seq and ChEC-seq predicted binding to ‘degenerate motif’ library. (**A**) (Left) 2D histograms of Msn2-FL AffinityNet predicted BET-seq -ΔΔGs versus marginalized BPNet predicted log-transformed counts of Msn2-FL (Brodsky, 2020) binding to the ‘degenerate motif’ library. Library sequences were binned by AffinityNet predicted -ΔΔG and BPNet predicted marginalized log-transformed counts, and with the shade of blue per pixel indicating density of sequences per bin. Vertical and horizontal red lines indicate the following thresholds using the standard deviations (σ) and means (µ) of fitted gaussian noise distributions to each sample used to define similarly and differentially bound sequence sets in each row: sequences with -ΔΔG and log(counts) > 2 σ from noise distribution µ values were bound in both BET-seq and ChEC-seq models (top); sequences with -ΔΔG > 2 σ and log(counts) < 1 σ from noise distribution µ values were bound only in BET-seq models (middle); sequences with -ΔΔG < 1 σ and log(counts) > 2 σ from noise distribution µ values were bound only in ChEC-seq models (bottom). Points colored in red were sequences that passed these thresholds, with a maximum of 10,000 being taken for further analysis. The total number of sequences used are indicated for each plot. (Right) Similarly and differentially bound sequences were used to generate Position Probability Matrices centered on matches to AGGGG (for the ‘Both Bound’ and ‘BET-seq up’ sets) or CGGGG (for the ‘ChEC-seq up’ set). For each set, the percentages of sequences with AGGGG and CGGGG matches are indicated. (**B**) shows the same analysis as in (A) comparing Msn2-FL AffinityNet BET-seq predictions to marginalized BPNet model predictions for a model trained on Mahendrawada, 2025 Msn2-FL (37°C) binding data.

**Supplementary Figure 60.**
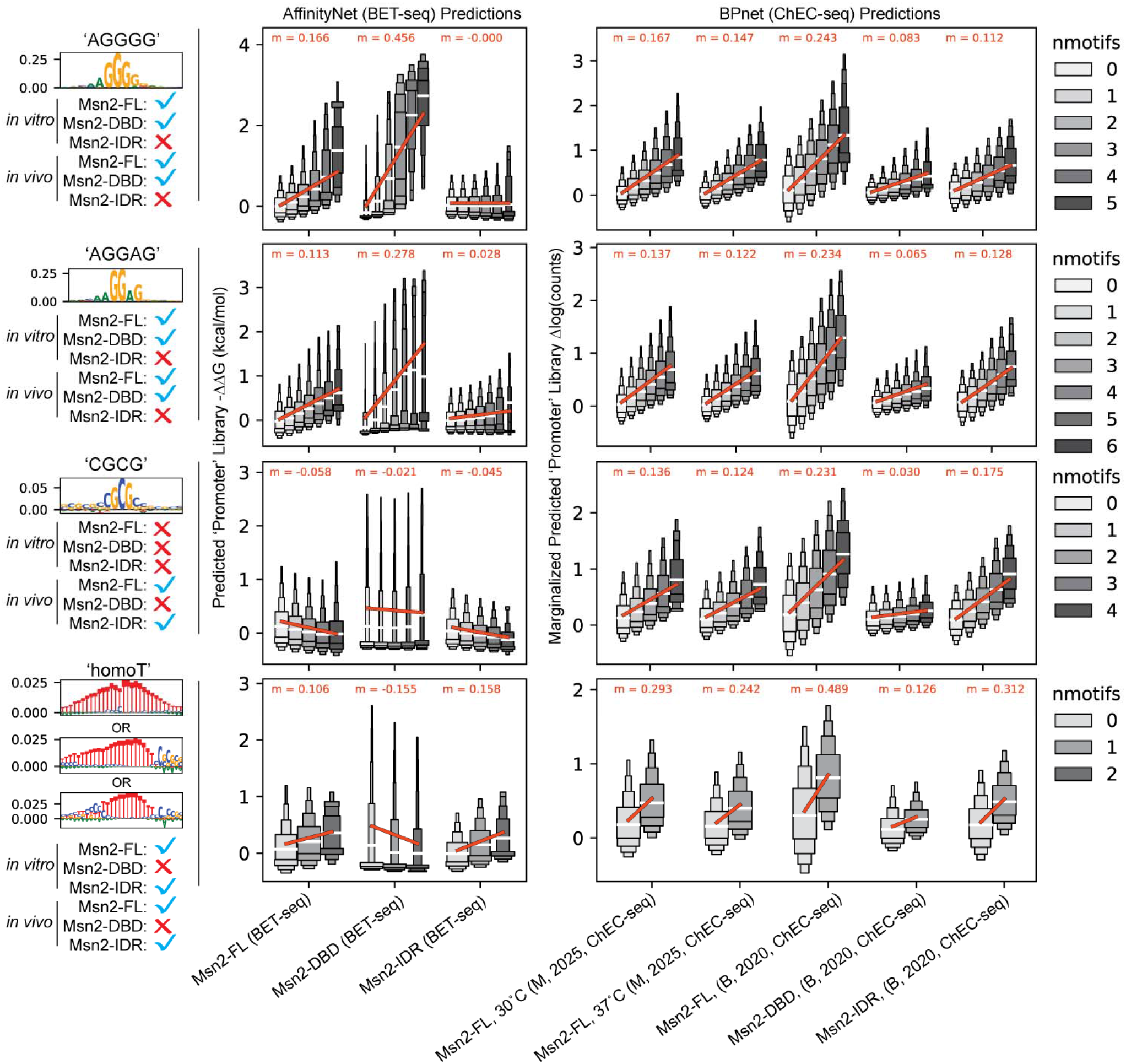
Predicted binding distributions of ‘promoter’ library tiles as a function of motif presence and number. ‘Boxenplots’ showing the distributions of min/max normalized (see Methods) predicted AffinityNet (-ΔΔG) and marginalized BPNet (log transformed counts) predicted binding to ‘promoter’ library sequences grouped by the number of FiNeMo motif matches to the shown CWMs. Rows indicate which *in vivo* model was compared. Boxes are centered around the medians (50^th^ percentile) of each distribution (denoted with a white line), and each subsequent layer represents percentile thresholds decreasing in size by powers of 2 (i.e. going out from the median, 25^th^ %, 12.5%, 6.25%, and 3.125%). Box widths represent the density of sequences in each percentile bin. Box colors indicate the number of motif matches to the CWM(s) shown to the left of each plot. Red lines indicate the linear fit line to each set of binding distributions as a function of the number of motifs with the fitted slopes annotated for each line. Sequences were binned by the max number of FiNeMo motif matches per sequence for a given CWM set and sample group. Sample groups are as follows: Msn2-FL, DBD, and IDR AffinityNet models; M, 2025 Msn2-FL 30°C and 37°C BPNet models; and B, 2020 Msn2-FL, DBD, and IDR BPNet models. The one exception to the above sequence binning was for the ‘CGCG’ motif binning with AffinityNet models, for which matches were counted using the M, 2025 Msn2-FL 30°C and 37°C model group since there were no ‘CGCG’ motif matches within the *in vitro* models. Sequence sets with a given number of motif matches were limited to those with at least 200 sequences. In order from top to bottom, CWMs for each row are from the following MoDISco results: ‘AGGGG’ - #1 CWM from the Msn2-DBD ‘promoter’ library AffinityNet model (Supplementary Fig. 34); ‘AGGAG’ - #2 CWM from the Msn2-DBD ‘promoter’ library AffinityNet model (Supplementary Fig. 34); ‘CGCG’ - #1 CWM from the B, 2020 Msn2-IDR BPNet model (Fig. 6D); ‘homoT’ - #9 CWM from the Msn2-FL ‘promoter’ library AffinityNet model (Supplementary Fig. 34), the #5 CWM from the M, 2025 (30°C) Msn2-FL BPNet model, and the #2 CWM from the B, 2020 Msn2-FL BPNet model. Below each motif blue check marks and red ‘X’s indicate whether the motif or a similar motif was found in each model or set of models.

**Supplementary Figure 61.**
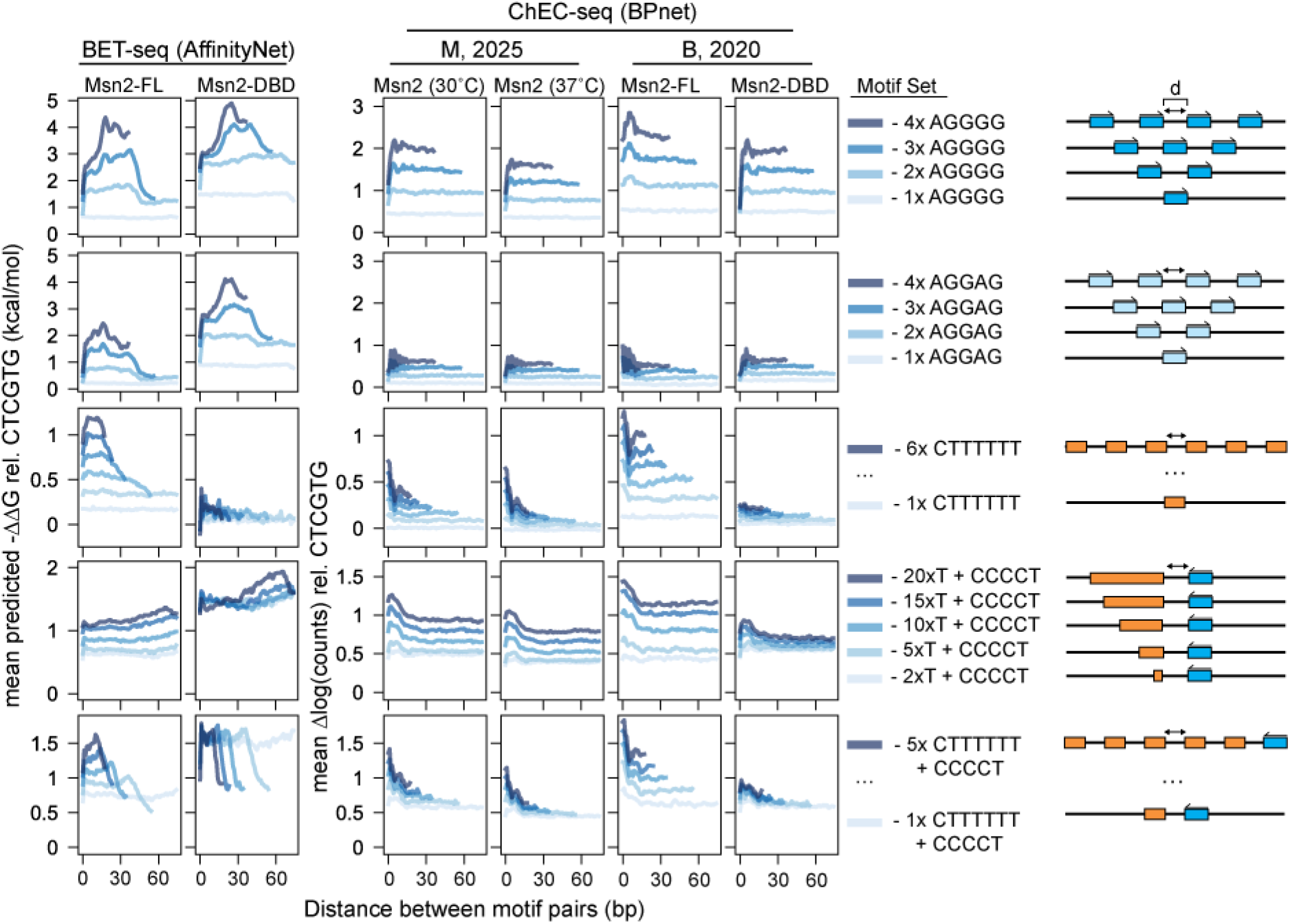
Simulated *in vitro* and *in vivo* binding to varied numbers of motifs and motif combinations. Lineplots of *in vitro* AffinityNet (left two columns) and *in vivo* BPNet (right two columns) mean simulated binding (-ΔΔG and Δ log(counts) respectively) of Msn2 FL and DBD constructs to sequences with varied numbers of AGGGG, and homoT motifs and motif variants as a function of distance between the motifs relative to background sequence with a non-binding E-box motif (CACGTG). Motif sets, indicated per row, were spaced equidistantly within 200 dinucleotide shuffled background sequences at a maximum distance of 75 bp between motifs.

## SUPPLEMENTARY METHODS FIGURES

**Supplementary Figure 62:**
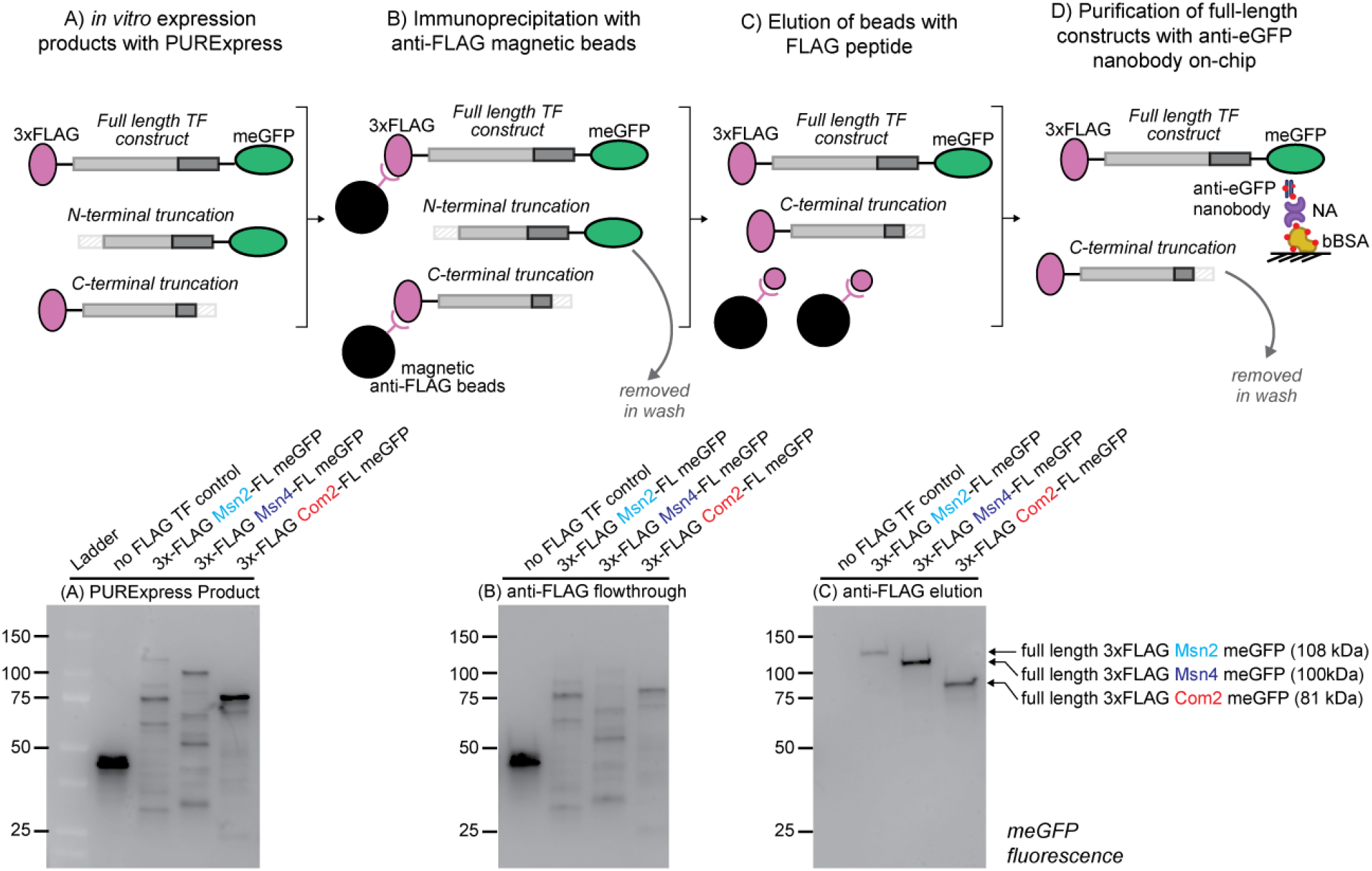
Schematic of double-immunoprecipitation purification strategy for isolating full-length TF constructs. (Top) *In vitro* expressed TF constructs are first pulled down using anti-FLAG magnetic beads that bind their N-terminal 3xFLAG tag and then eluted using purified FLAG peptide. Subsequently, TFs are purified by immunoprecipitating their C-terminal monomeric eGFP tag on MITOMI or BET-seq devices. (Bottom) Example semi-denaturing PAGEs with protein products from each of the above steps of FL Msn2, Msn4, and Com2 constructs with meGFP-tag fluorescence visualized using blue light excitation.

**Supplementary Figure 63.**
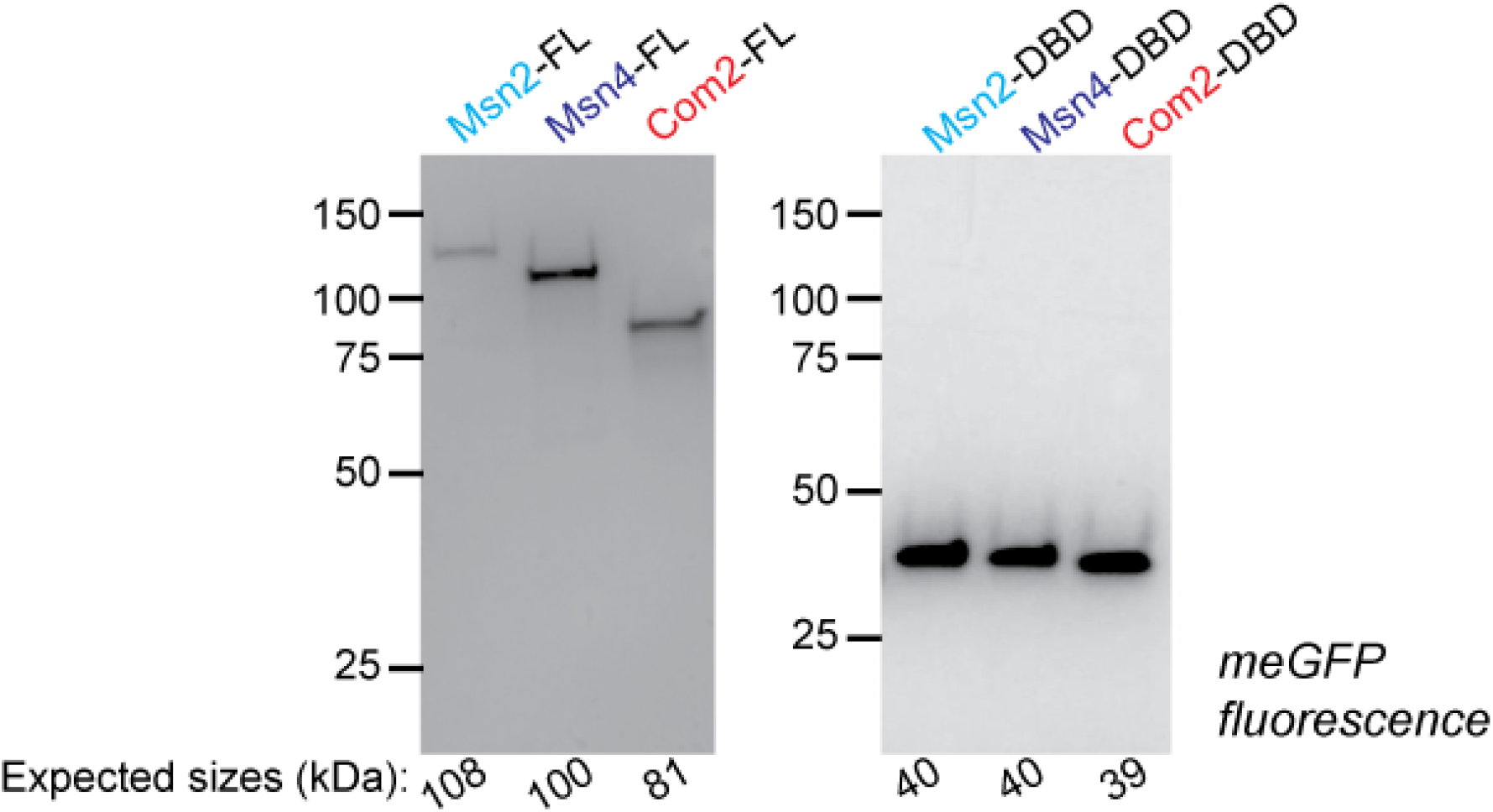
FL and DBD-only Msn2, Msn4, and Com2 constructs after elution from anti-FLAG beads. Gels were imaged using blue light excitation.

**Supplementary Figure 64.**
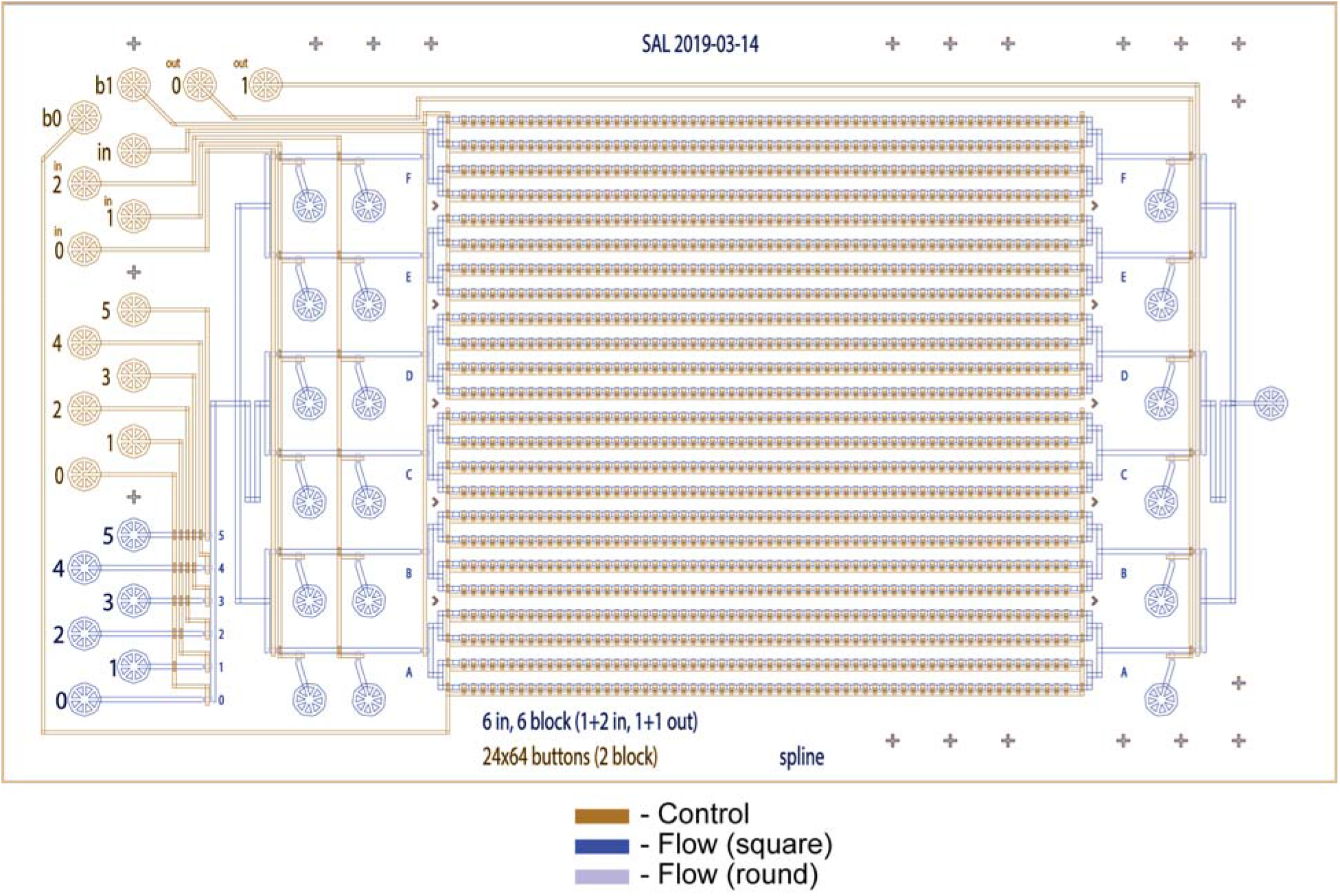
Design of the valved microfluidic device used for BET-seq in this study (‘spline 6-block’). The control layer (valves) is shown in red. The main, non-deformable flow layer (square) is shown in dark blue and the deformable flow layer (round) is shown in light purple.

**Supplementary Figure 65.**
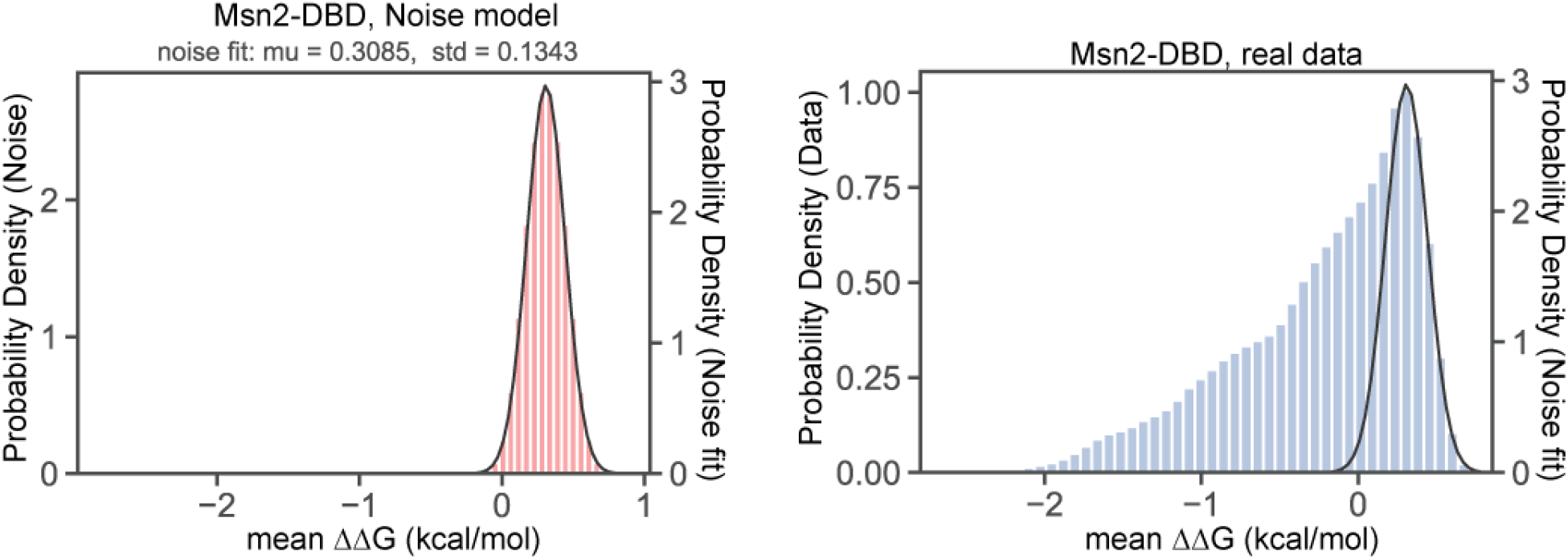
Example Gaussian fit to the ‘noise’ of BET-seq data. (Left) Synthetic ‘noise’ distribution constructed from Msn2-DBD ΔΔGs for the ‘promoter’ library (red bars) with the fitted Gaussian (black line; fitted mean and standard deviation annotated). (Right) Full distribution of Msn2-DBD ΔΔGs (blue bars) with the same Gaussian noise fit overlaid. The mean of the noise distribution is determined by finely binning the data by ΔΔG and selecting the bin center with the greatest number of data points. A symmetrical “noise” distribution is then created by reflecting the data with a ΔΔG greater than the noise mean across the mean and we fit a Gaussian curve to this noise distribution to determine the standard deviation of the noise.

**Supplementary Figure 66.**
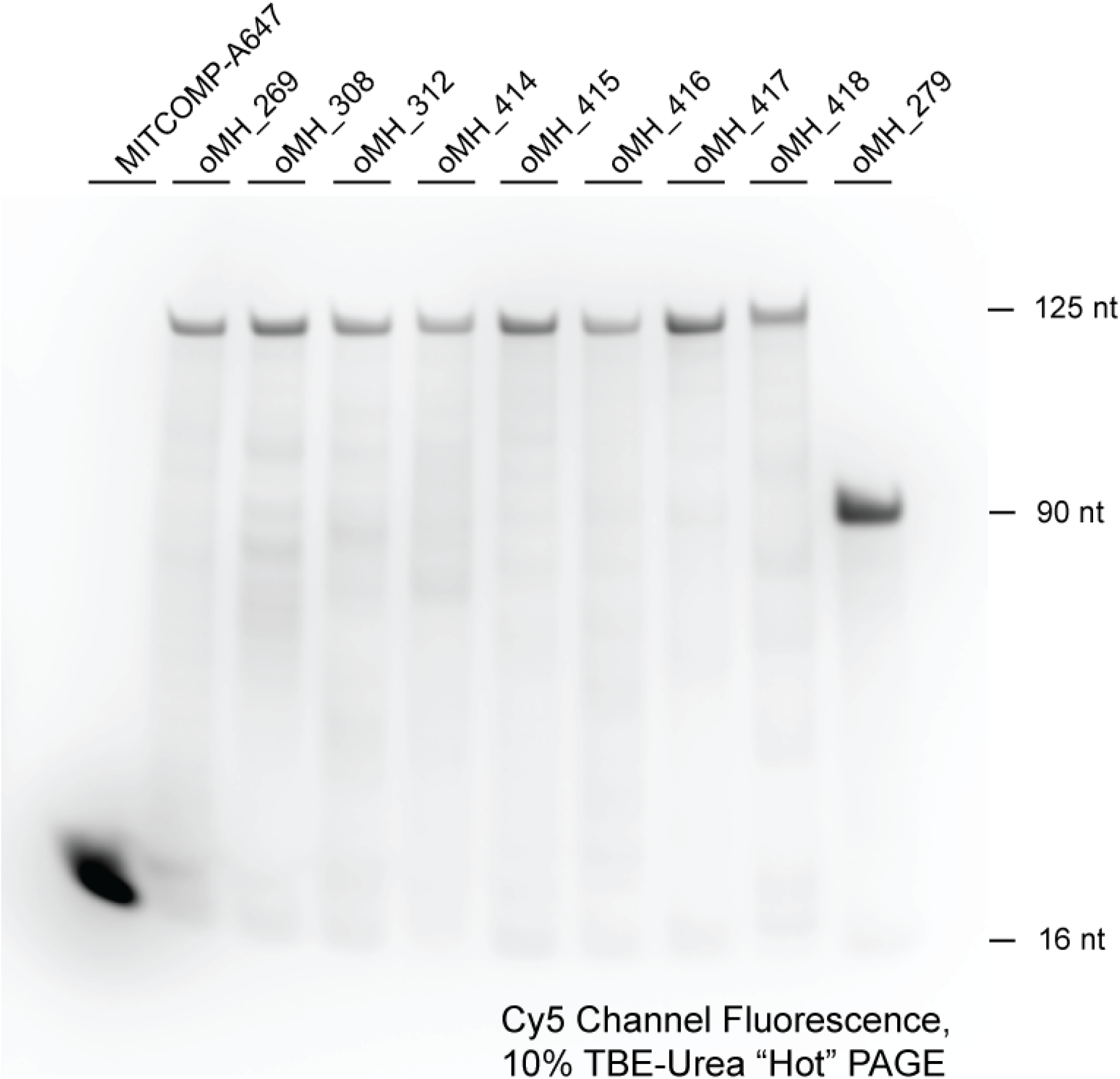
Representative denaturing 10% polyacrylamide TBE-Urea gel electrophoresis of AlexaFluor-647 labelled oligonucleotides. ssDNA gel showing proper extension of AlexaFluor647-labelled primer with various DNA sequences (from Fig. 5K, sequences available in **Supplementary File 6**) for subsequent binding measurements with MITOMI. Gel was imaged in the Cy5-channel with red light excitation.

**Supplementary Figure 67.**
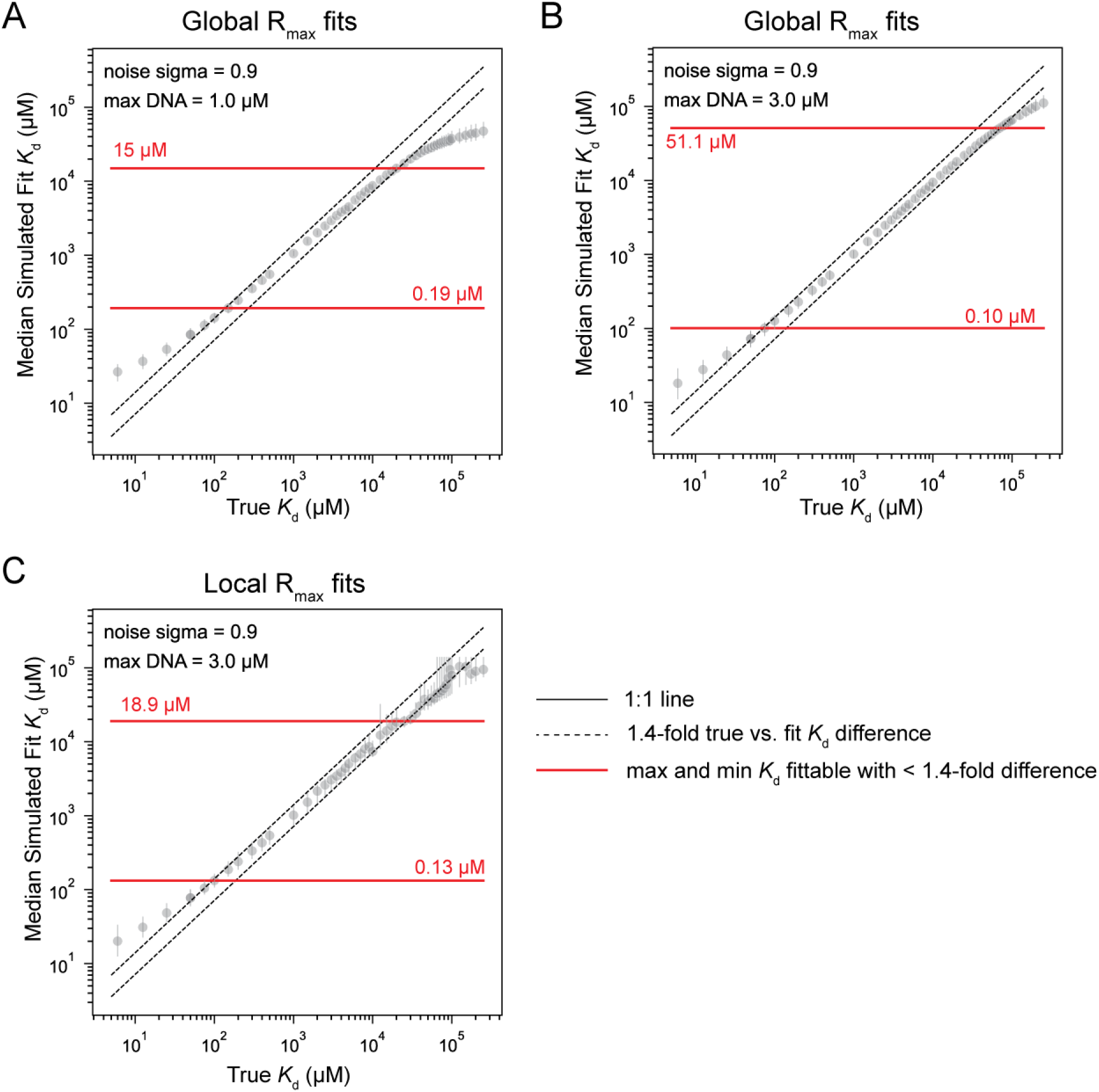
Simulated *K_d_* fits for interactions of various strengths to determine upper and lower limits on MITOMI *K_d_* fits. (**A**-**C**) Scatterplots of log-transformed versus median fit *K_d_* as a function of true *K_d_* for simulated binding curves with either global (**A**-**B**) or local (**C**) R_max_ fitting and with max total DNA concentrations of either 1 µM (**A**) or 3 µM (**B**-**C**) (**Methods**). Points represent individual binding interactions simulated; error bars represent boundaries of the upper and lower quartiles of the *K_d_*fit distributions; dashed black lines denote thresholds 1.4-fold difference between true and fitted *K_d_*; solid red lines denote the minimum and maximum fitted median *K_d_* values for which the difference between true and fitted *K_d_* is below 1.4-fold, or ∼ 0.2 kcal/mol with their values denoted in red.

**Supplementary Figure 68.**
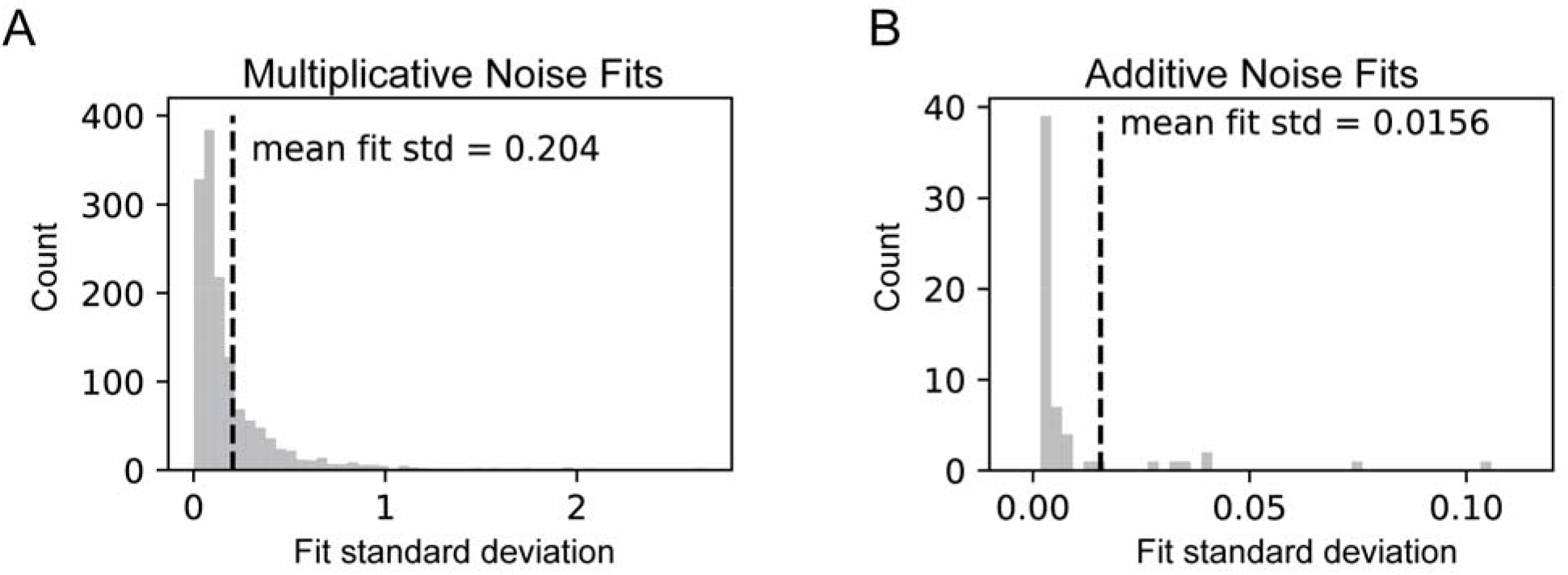
Parameterization of noise for binding simulations. (**A**) Histogram of fit standard deviation parameters for log normal distribution fits to distributions of alexafluor-647/GFP fluorescence ratios for TF-DNA-DNA dilution combinations across all MITOMI experiments with measurable binding (**Methods**). (**B**) Histogram of fit standard deviation parameters for normal distribution fits to alexafluor-647/GFP fluorescence ratios of “blank” chambers with GFP-tagged TF but no DNA printed. Dashed black lines indicate mean fit standard deviations with their values annotated.

**Supplementary Figure 69.**
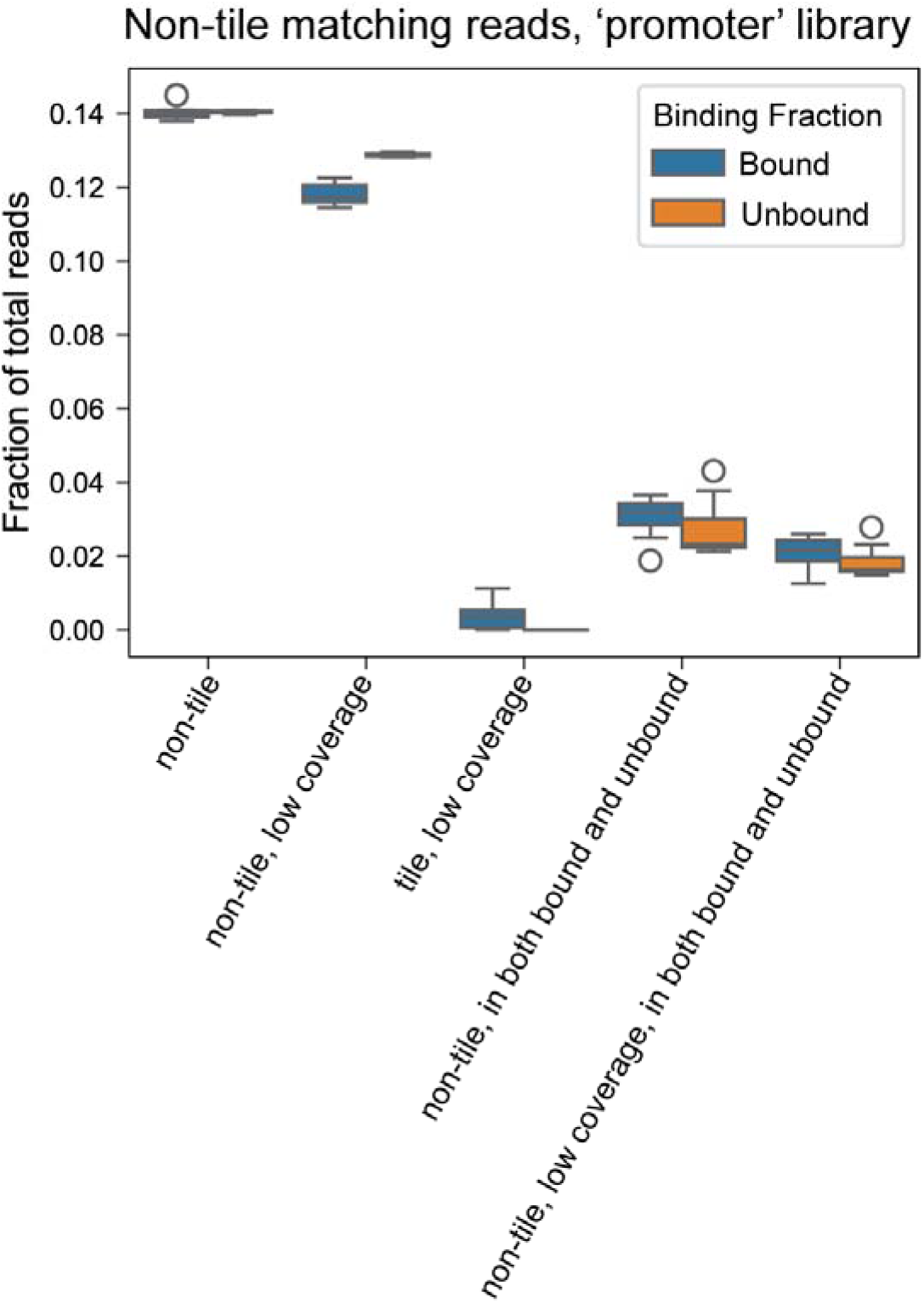
Analysis of reads not matching designed tiles in the ‘promoter’ library. Shown are distributions across a representative sequencing run (12 bound and 12 unbound libraries) of fractions of all unique reads comprised of (1) non-tile matching reads, (2) non-tile reads of sequences with low coverage (< 10 reads per sequence), (3) tile matching reads of sequences with low coverage, (4) non-tile matching reads to sequences that appear in both the bound and unbound libraries, and (5) non-tile matching reads to sequences that appear in both the bound and unbound libraries with low coverage.

