## Supplementary File 1 for "Intrinsically disordered regions within transcription factors impact DNA binding through both autoinhibition and independent DNA binding"

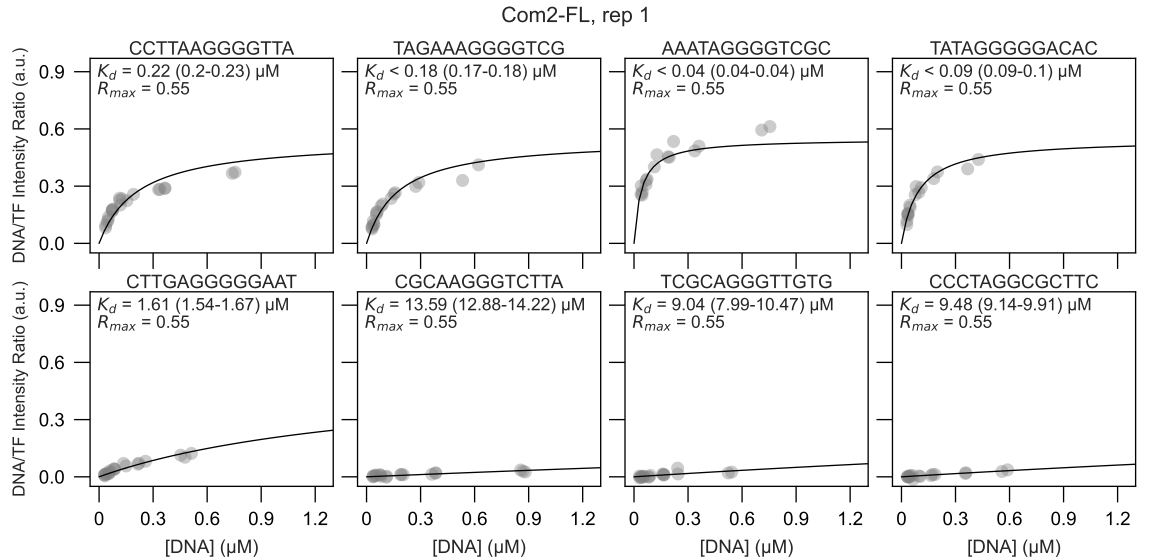

Com2-FL, rep 1
CCTTAAGGGGTTA
$K_d$ = 0.22 (0.2-0.23) μM
$R_{max}$ = 0.55
TAGAAAGGGGTCG
$K_d$ < 0.18 (0.17-0.18) μM
$R_{max}$ = 0.55
AAATAGGGGTCGC
$K_d$ < 0.04 (0.04-0.04) μM
$R_{max}$ = 0.55
TATAGGGGGACAC
$K_d$ < 0.09 (0.09-0.1) μM
$R_{max}$ = 0.55
CTTGAGGGGGAAT
$K_d$ = 1.61 (1.54-1.67) μM
$R_{max}$ = 0.55
CGCAAGGGTCTTA
$K_d$ = 13.59 (12.88-14.22) μM
$R_{max}$ = 0.55
TCGCAGGGTTGTG
$K_d$ = 9.04 (7.99-10.47) μM
$R_{max}$ = 0.55
CCCTAGGCGCTTC
$K_d$ = 9.48 (9.14-9.91) μM
$R_{max}$ = 0.55
DNA/TF Intensity Ratio (a.u.)
0.0
0.3
0.6
0.9
[DNA] (μM)
0
0.3
0.6
0.9
1.2

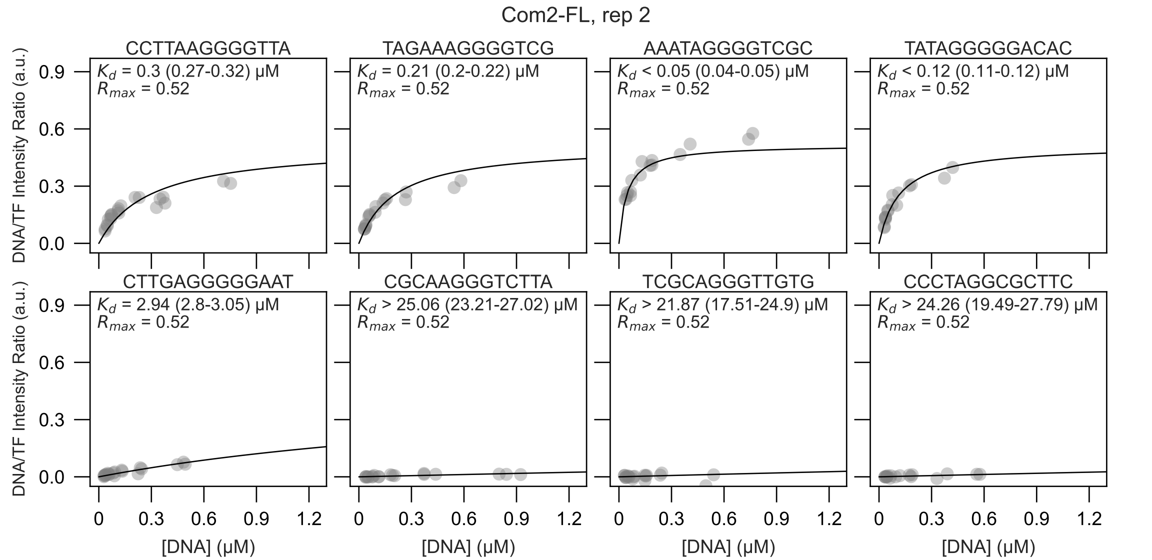
Com2-FL, rep 2
CCTTAAGGGGTTA
K_d = 0.3 (0.27-0.32) μM
R_max = 0.52
TAGAAAGGGGTCG
K_d = 0.21 (0.2-0.22) μM
R_max = 0.52
AAATAGGGGTCGC
K_d < 0.05 (0.04-0.05) μM
R_max = 0.52
TATAGGGGGACAC
K_d < 0.12 (0.11-0.12) μM
R_max = 0.52
CTTGAGGGGGAAT
K_d = 2.94 (2.8-3.05) μM
R_max = 0.52
CGCAAGGGTCTTA
K_d > 25.06 (23.21-27.02) μM
R_max = 0.52
TCGCAGGGTTGTG
K_d > 21.87 (17.51-24.9) μM
R_max = 0.52
CCCTAGGCGCTTC
K_d > 24.26 (19.49-27.79) μM
R_max = 0.52
DNA/TF Intensity Ratio (a.u.)
[DNA] (μM)

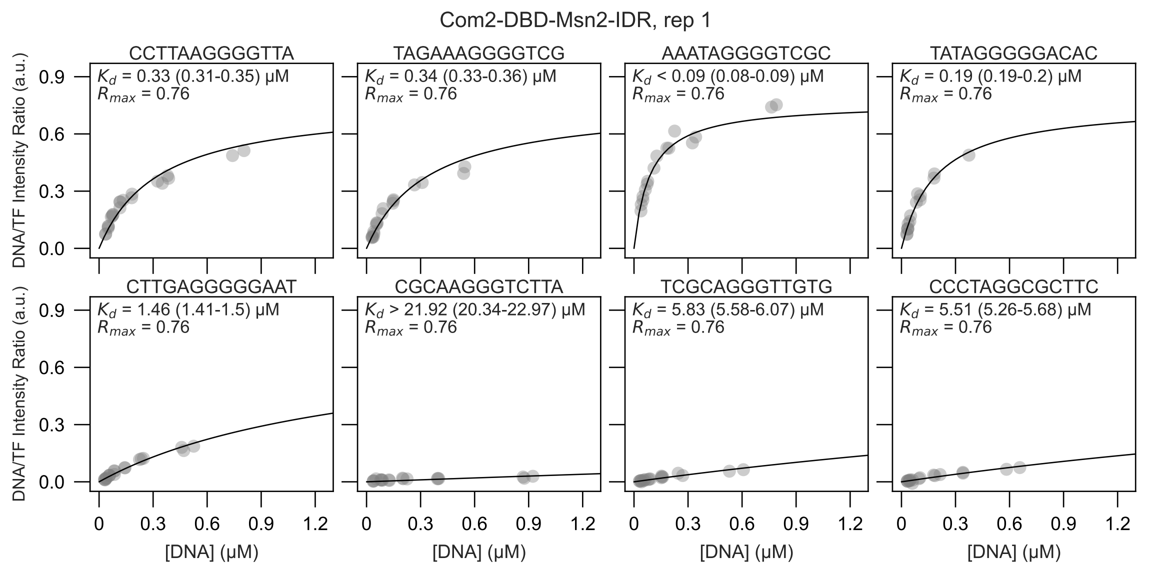

Com2-DBD-Msn2-IDR, rep 1
CCTTAAGGGGTTA
$K_d$ = 0.33 (0.31-0.35) μM
$R_{max}$ = 0.76
TAGAAAGGGGTCG
$K_d$ = 0.34 (0.33-0.36) μM
$R_{max}$ = 0.76
AAATAGGGGTCGC
$K_d$ < 0.09 (0.08-0.09) μM
$R_{max}$ = 0.76
TATAGGGGGACAC
$K_d$ = 0.19 (0.19-0.2) μM
$R_{max}$ = 0.76
CTTGAGGGGGAAT
$K_d$ = 1.46 (1.41-1.5) μM
$R_{max}$ = 0.76
CGCAAGGGTCTTA
$K_d$ > 21.92 (20.34-22.97) μM
$R_{max}$ = 0.76
TCGCAGGGTTGTG
$K_d$ = 5.83 (5.58-6.07) μM
$R_{max}$ = 0.76
CCCTAGGCGCTTC
$K_d$ = 5.51 (5.26-5.68) μM
$R_{max}$ = 0.76
DNA/TF Intensity Ratio (a.u.)
0.0
0.3
0.6
0.9
[DNA] (μM)
0
0.3
0.6
0.9
1.2

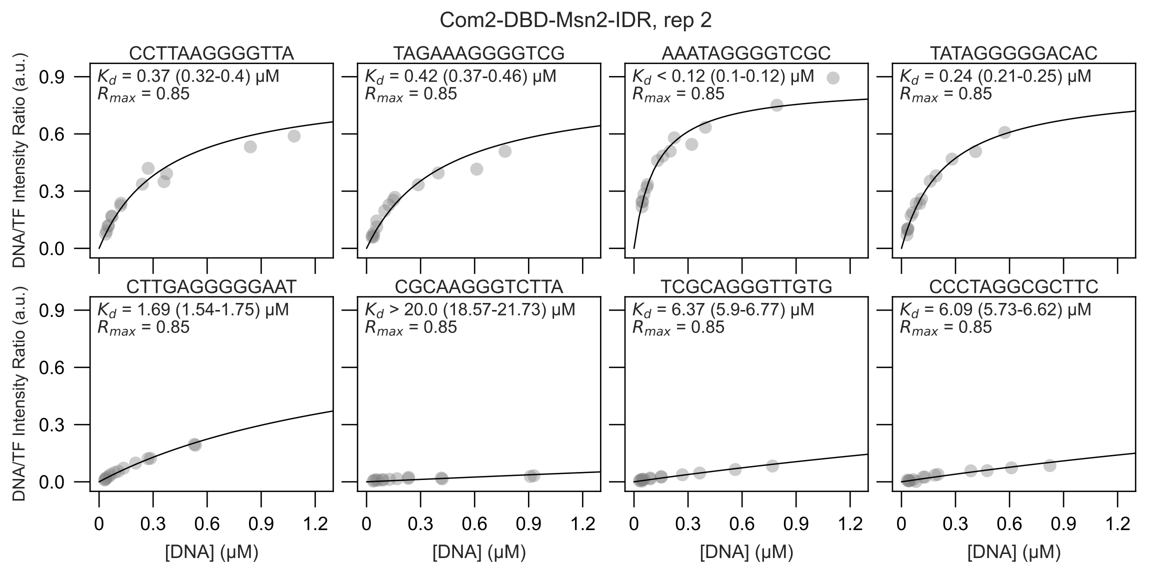
Com2-DBD-Msn2-IDR, rep 2
CCTTAAGGGGTTA
$K_d$ = 0.37 (0.32-0.4) μM
$R_{max}$ = 0.85
TAGAAAGGGGTCG
$K_d$ = 0.42 (0.37-0.46) μM
$R_{max}$ = 0.85
AAATAGGGGTCGC
$K_d$ < 0.12 (0.1-0.12) μM
$R_{max}$ = 0.85
TATAGGGGGACAC
$K_d$ = 0.24 (0.21-0.25) μM
$R_{max}$ = 0.85
CTTGAGGGGGAAT
$K_d$ = 1.69 (1.54-1.75) μM
$R_{max}$ = 0.85
CGCAAGGGTCTTA
$K_d$ > 20.0 (18.57-21.73) μM
$R_{max}$ = 0.85
TCGCAGGGTTGTG
$K_d$ = 6.37 (5.9-6.77) μM
$R_{max}$ = 0.85
CCCTAGGCGCTTC
$K_d$ = 6.09 (5.73-6.62) μM
$R_{max}$ = 0.85
DNA/TF Intensity Ratio (a.u.)
0.0
0.3
0.6
0.9
[DNA] (μM)
0
0.3
0.6
0.9
1.2

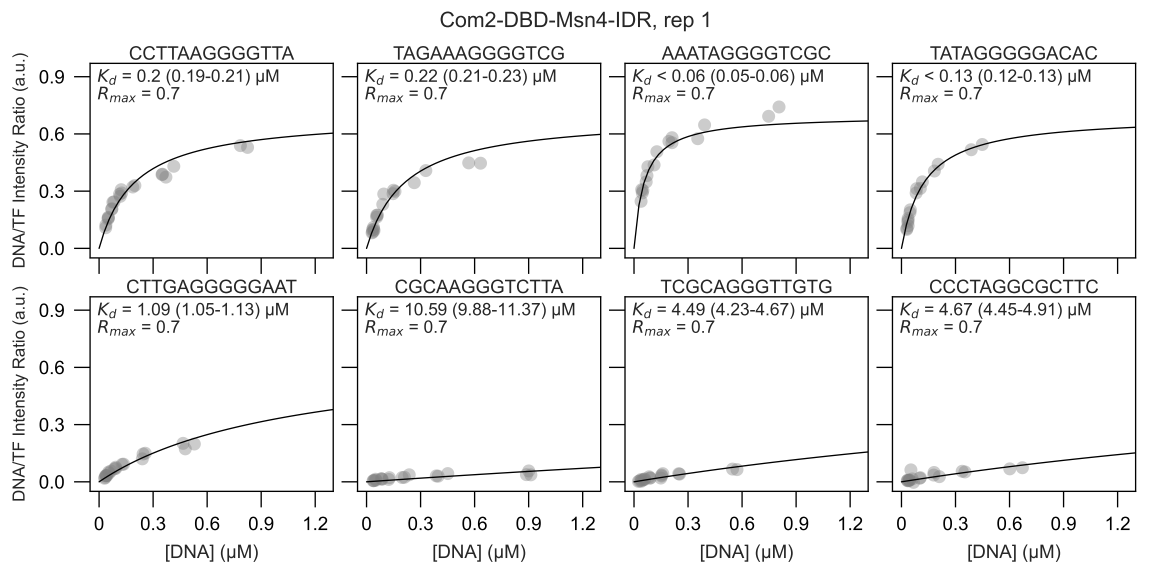

Com2-DBD-Msn4-IDR, rep 1
CCTTAAGGGGTTA
$K_d$ = 0.2 (0.19-0.21) μM
$R_{max}$ = 0.7
TAGAAAGGGGTCG
$K_d$ = 0.22 (0.21-0.23) μM
$R_{max}$ = 0.7
AAATAGGGGTCGC
$K_d$ < 0.06 (0.05-0.06) μM
$R_{max}$ = 0.7
TATAGGGGGACAC
$K_d$ < 0.13 (0.12-0.13) μM
$R_{max}$ = 0.7
CTTGAGGGGGAAT
$K_d$ = 1.09 (1.05-1.13) μM
$R_{max}$ = 0.7
CGCAAGGGTCTTA
$K_d$ = 10.59 (9.88-11.37) μM
$R_{max}$ = 0.7
TCGCAGGGTTGTG
$K_d$ = 4.49 (4.23-4.67) μM
$R_{max}$ = 0.7
CCCTAGGCGCTTC
$K_d$ = 4.67 (4.45-4.91) μM
$R_{max}$ = 0.7
DNA/TF Intensity Ratio (a.u.)
[DNA] (μM)
0.0
0.3
0.6
0.9
0
1.2

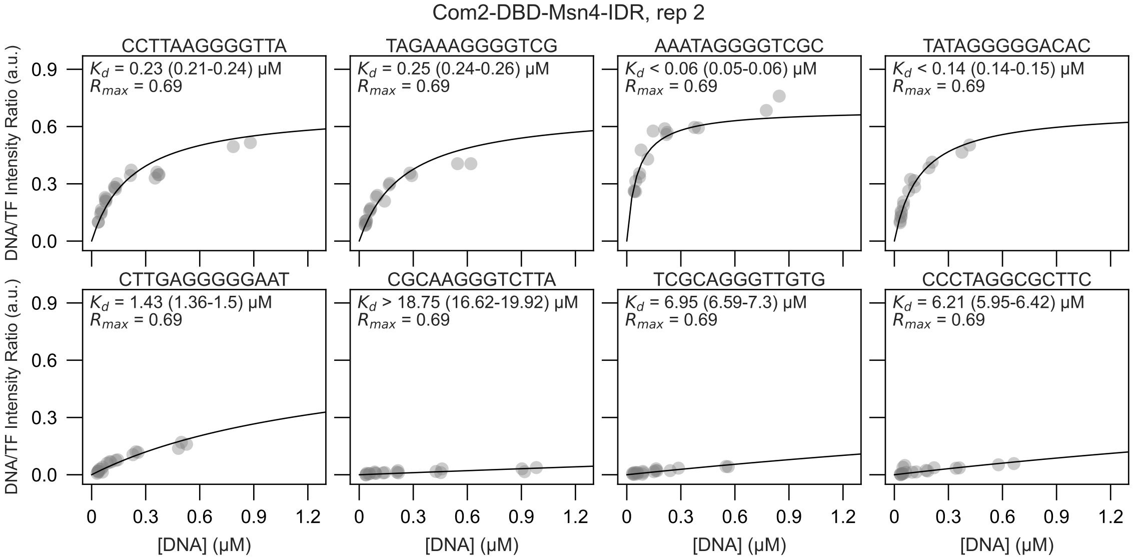
Com2-DBD-Msn4-IDR, rep 2
CCTTAAGGGGTTA
$K_d$ = 0.23 (0.21-0.24) μM
$R_{max}$ = 0.69
TAGAAAGGGGTCG
$K_d$ = 0.25 (0.24-0.26) μM
$R_{max}$ = 0.69
AAATAGGGGTCGC
$K_d$ < 0.06 (0.05-0.06) μM
$R_{max}$ = 0.69
TATAGGGGGACAC
$K_d$ < 0.14 (0.14-0.15) μM
$R_{max}$ = 0.69
CTTGAGGGGGAAT
$K_d$ = 1.43 (1.36-1.5) μM
$R_{max}$ = 0.69
CGCAAGGGTCTTA
$K_d$ > 18.75 (16.62-19.92) μM
$R_{max}$ = 0.69
TCGCAGGGTTGTG
$K_d$ = 6.95 (6.59-7.3) μM
$R_{max}$ = 0.69
CCCTAGGCGCTTC
$K_d$ = 6.21 (5.95-6.42) μM
$R_{max}$ = 0.69
DNA/TF Intensity Ratio (a.u.)
0.0
0.3
0.6
0.9
[DNA] (μM)
0
0.3
0.6
0.9
1.2

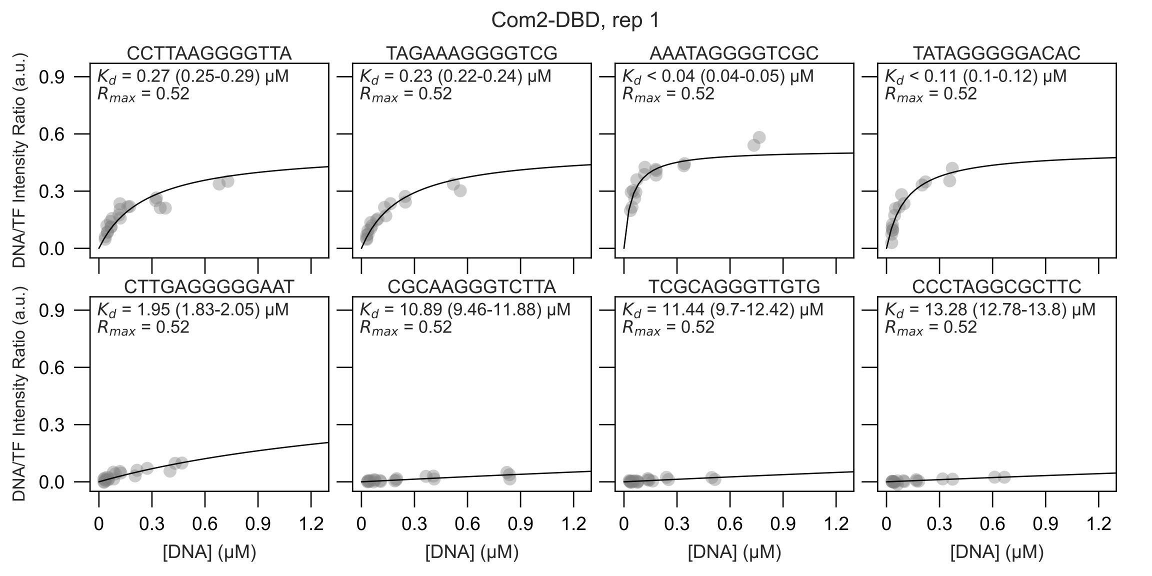

Com2-DBD, rep 1
CCTTAAGGGGTTA
$K_d$ = 0.27 (0.25-0.29) μM
$R_{max}$ = 0.52
TAGAAAGGGGTCG
$K_d$ = 0.23 (0.22-0.24) μM
$R_{max}$ = 0.52
AAATAGGGGTCGC
$K_d$ < 0.04 (0.04-0.05) μM
$R_{max}$ = 0.52
TATAGGGGGACAC
$K_d$ < 0.11 (0.1-0.12) μM
$R_{max}$ = 0.52
CTTGAGGGGGAAT
$K_d$ = 1.95 (1.83-2.05) μM
$R_{max}$ = 0.52
CGCAAGGGTCTTA
$K_d$ = 10.89 (9.46-11.88) μM
$R_{max}$ = 0.52
TCGCAGGGTTGTG
$K_d$ = 11.44 (9.7-12.42) μM
$R_{max}$ = 0.52
CCCTAGGCGCTTC
$K_d$ = 13.28 (12.78-13.8) μM
$R_{max}$ = 0.52
DNA/TF Intensity Ratio (a.u.)
0.0
0.3
0.6
0.9
[DNA] (μM)
0
0.3
0.6
0.9
1.2

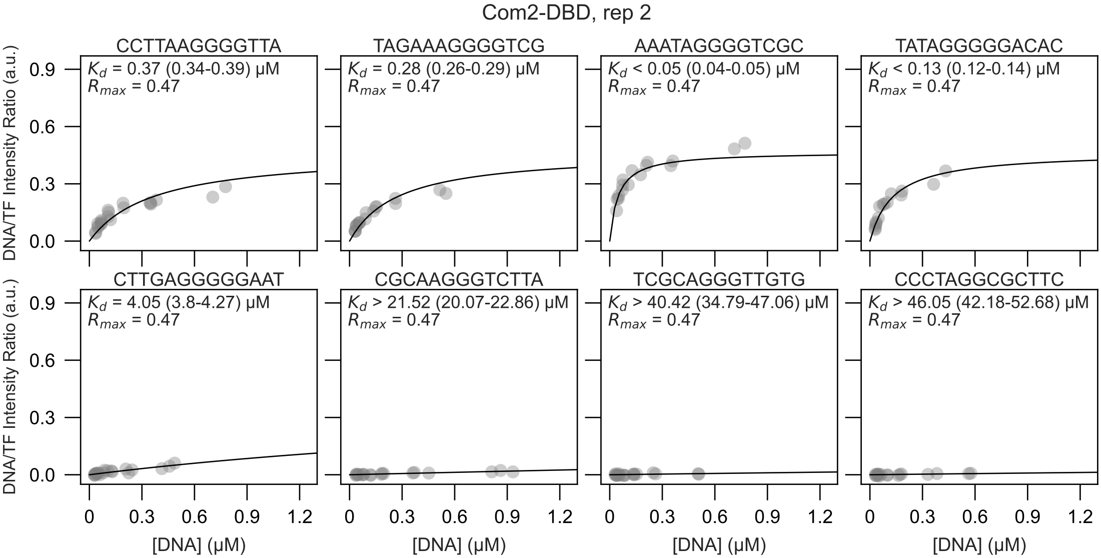

Com2-DBD, rep 2
CCTTAAGGGGTTA
$K_d$ = 0.37 (0.34-0.39) µM
$R_{max}$ = 0.47
TAGAAAGGGGTCG
$K_d$ = 0.28 (0.26-0.29) µM
$R_{max}$ = 0.47
AAATAGGGGTCGC
$K_d$ < 0.05 (0.04-0.05) µM
$R_{max}$ = 0.47
TATAGGGGGACAC
$K_d$ < 0.13 (0.12-0.14) µM
$R_{max}$ = 0.47
CTTGAGGGGGAAT
$K_d$ = 4.05 (3.8-4.27) µM
$R_{max}$ = 0.47
CGCAAGGGTCTTA
$K_d$ > 21.52 (20.07-22.86) µM
$R_{max}$ = 0.47
TCGCAGGGTTGTG
$K_d$ > 40.42 (34.79-47.06) µM
$R_{max}$ = 0.47
CCCTAGGCGCTTC
$K_d$ > 46.05 (42.18-52.68) µM
$R_{max}$ = 0.47
DNA/TF Intensity Ratio (a.u.)
0.0
0.3
0.6
0.9
[DNA] (µM)
0
0.3
0.6
0.9
1.2

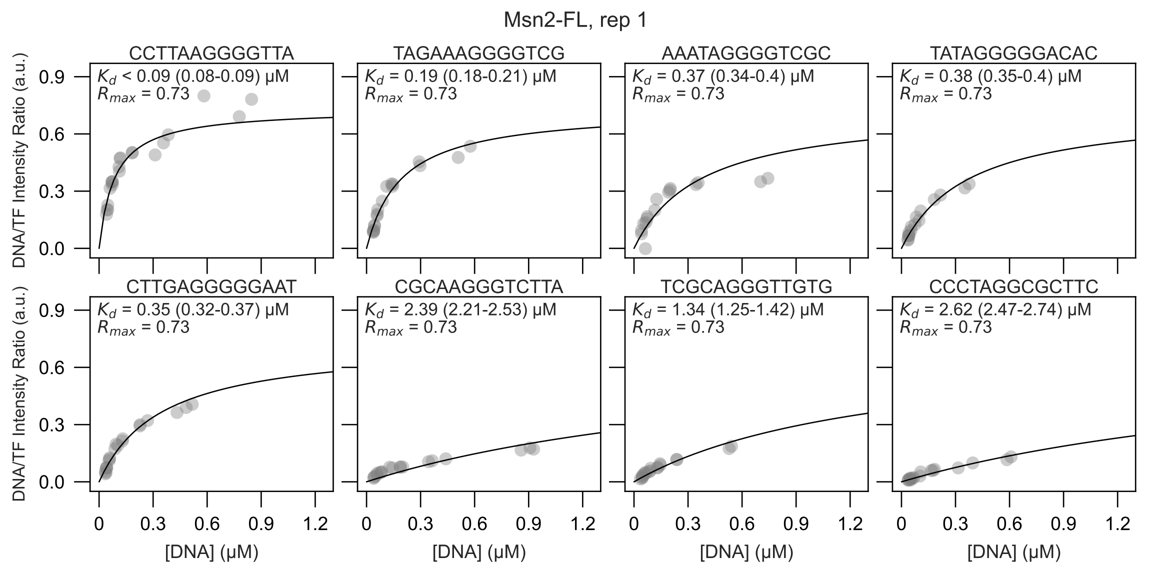

Msn2-FL, rep 1
CCTTAAGGGGTTA
Kd < 0.09 (0.08-0.09) μM
Rmax = 0.73
TAGAAAGGGGTCG
Kd = 0.19 (0.18-0.21) μM
Rmax = 0.73
AAATAGGGGTCGC
Kd = 0.37 (0.34-0.4) μM
Rmax = 0.73
TATAGGGGGACAC
Kd = 0.38 (0.35-0.4) μM
Rmax = 0.73
CTTGAGGGGGAAT
Kd = 0.35 (0.32-0.37) μM
Rmax = 0.73
CGCAAGGGTCTTA
Kd = 2.39 (2.21-2.53) μM
Rmax = 0.73
TCGCAGGGTTGTG
Kd = 1.34 (1.25-1.42) μM
Rmax = 0.73
CCCTAGGCGCTTC
Kd = 2.62 (2.47-2.74) μM
Rmax = 0.73
DNA/TF Intensity Ratio (a.u.)
[DNA] (μM)
0.0
0.3
0.6
0.9
0
0.3
0.6
0.9
1.2

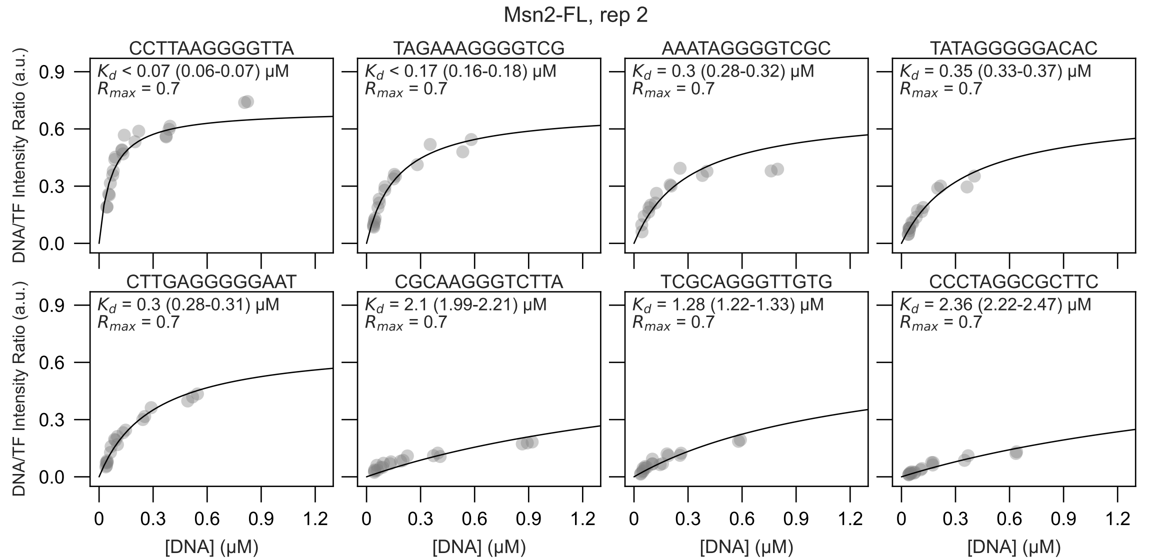

Msn2-FL, rep 2
CCTTAAGGGGTTA
$K_d$ < 0.07 (0.06-0.07) μM
$R_{max}$ = 0.7
TAGAAAGGGGTCG
$K_d$ < 0.17 (0.16-0.18) μM
$R_{max}$ = 0.7
AAATAGGGGTCGC
$K_d$ = 0.3 (0.28-0.32) μM
$R_{max}$ = 0.7
TATAGGGGGACAC
$K_d$ = 0.35 (0.33-0.37) μM
$R_{max}$ = 0.7
CTTGAGGGGGAAT
$K_d$ = 0.3 (0.28-0.31) μM
$R_{max}$ = 0.7
CGCAAGGGTCTTA
$K_d$ = 2.1 (1.99-2.21) μM
$R_{max}$ = 0.7
TCGCAGGGTTGTG
$K_d$ = 1.28 (1.22-1.33) μM
$R_{max}$ = 0.7
CCCTAGGCGCTTC
$K_d$ = 2.36 (2.22-2.47) μM
$R_{max}$ = 0.7
DNA/TF Intensity Ratio (a.u.)
[DNA] (μM)
0.0
0.3
0.6
0.9
0
0.3
0.6
0.9
1.2

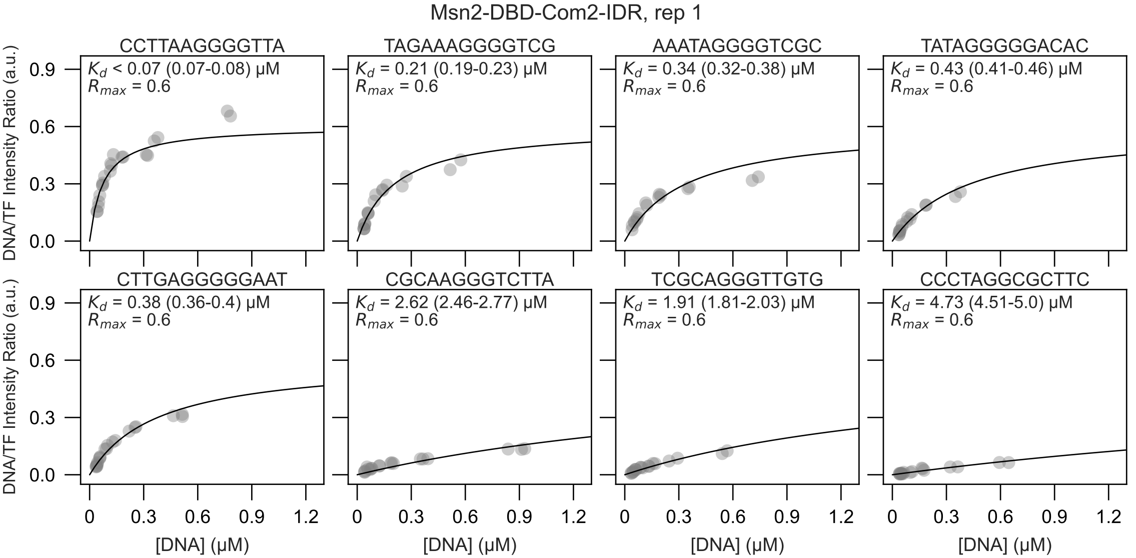

Msn2-DBD-Com2-IDR, rep 1
CCTTAAGGGGTTA
$K_d$ < 0.07 (0.07-0.08) μM
$R_{max}$ = 0.6
TAGAAAGGGGTCG
$K_d$ = 0.21 (0.19-0.23) μM
$R_{max}$ = 0.6
AAATAGGGGTCGC
$K_d$ = 0.34 (0.32-0.38) μM
$R_{max}$ = 0.6
TATAGGGGGACAC
$K_d$ = 0.43 (0.41-0.46) μM
$R_{max}$ = 0.6
CTTGAGGGGGAAT
$K_d$ = 0.38 (0.36-0.4) μM
$R_{max}$ = 0.6
CGCAAGGGTCTTA
$K_d$ = 2.62 (2.46-2.77) μM
$R_{max}$ = 0.6
TCGCAGGGTTGTG
$K_d$ = 1.91 (1.81-2.03) μM
$R_{max}$ = 0.6
CCCTAGGCGCTTC
$K_d$ = 4.73 (4.51-5.0) μM
$R_{max}$ = 0.6
DNA/TF Intensity Ratio (a.u.)
[DNA] (μM)
0.0
0.3
0.6
0.9
0
1.2

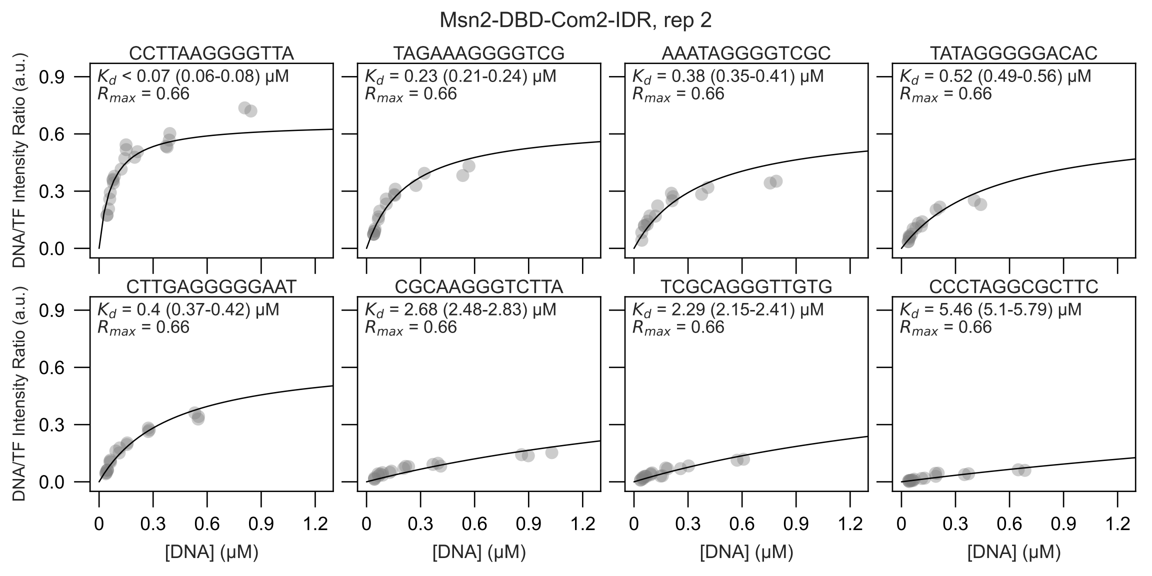

Msn2-DBD-Com2-IDR, rep 2
CCTTAAGGGGTTA
$K_d$ < 0.07 (0.06-0.08) μM
$R_{max}$ = 0.66
TAGAAAGGGGTCG
$K_d$ = 0.23 (0.21-0.24) μM
$R_{max}$ = 0.66
AAATAGGGGTCGC
$K_d$ = 0.38 (0.35-0.41) μM
$R_{max}$ = 0.66
TATAGGGGGACAC
$K_d$ = 0.52 (0.49-0.56) μM
$R_{max}$ = 0.66
CTTGAGGGGGAAT
$K_d$ = 0.4 (0.37-0.42) μM
$R_{max}$ = 0.66
CGCAAGGGTCTTA
$K_d$ = 2.68 (2.48-2.83) μM
$R_{max}$ = 0.66
TCGCAGGGTTGTG
$K_d$ = 2.29 (2.15-2.41) μM
$R_{max}$ = 0.66
CCCTAGGCGCTTC
$K_d$ = 5.46 (5.1-5.79) μM
$R_{max}$ = 0.66
DNA/TF Intensity Ratio (a.u.)
0.0
0.3
0.6
0.9
[DNA] (μM)
0
0.3
0.6
0.9
1.2

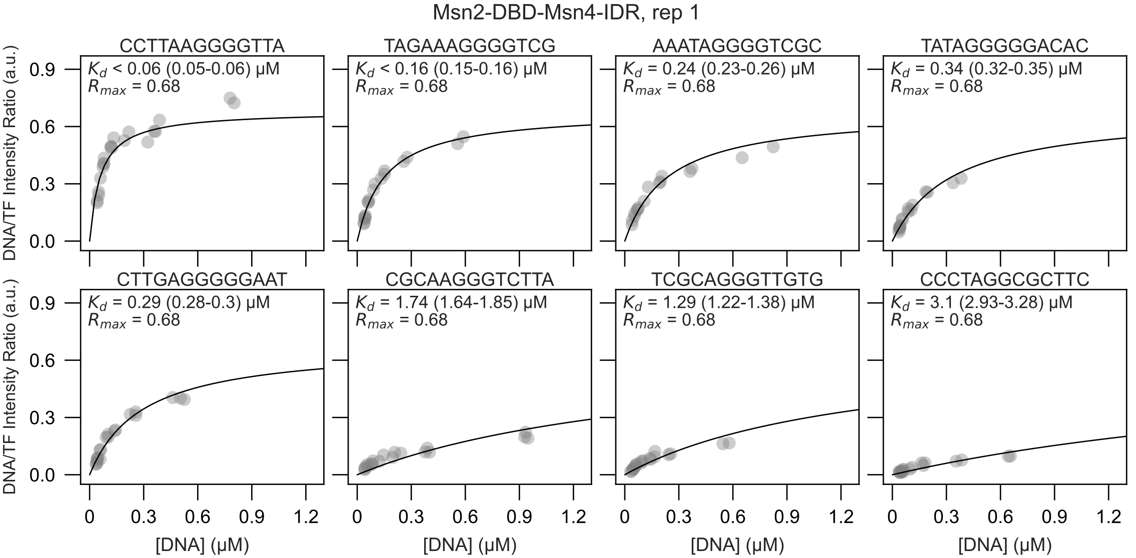

Msn2-DBD-Msn4-IDR, rep 1
CCTTAAGGGGTTA
$K_d$ < 0.06 (0.05-0.06) μM
$R_{max}$ = 0.68
TAGAAAGGGGTCG
$K_d$ < 0.16 (0.15-0.16) μM
$R_{max}$ = 0.68
AAATAGGGGTCGC
$K_d$ = 0.24 (0.23-0.26) μM
$R_{max}$ = 0.68
TATAGGGGGACAC
$K_d$ = 0.34 (0.32-0.35) μM
$R_{max}$ = 0.68
CTTGAGGGGGAAT
$K_d$ = 0.29 (0.28-0.3) μM
$R_{max}$ = 0.68
CGCAAGGGTCTTA
$K_d$ = 1.74 (1.64-1.85) μM
$R_{max}$ = 0.68
TCGCAGGGTTGTG
$K_d$ = 1.29 (1.22-1.38) μM
$R_{max}$ = 0.68
CCCTAGGCGCTTC
$K_d$ = 3.1 (2.93-3.28) μM
$R_{max}$ = 0.68
DNA/TF Intensity Ratio (a.u.)
[DNA] (μM)
0.0
0.3
0.6
0.9
0
0.3
0.6
0.9
1.2

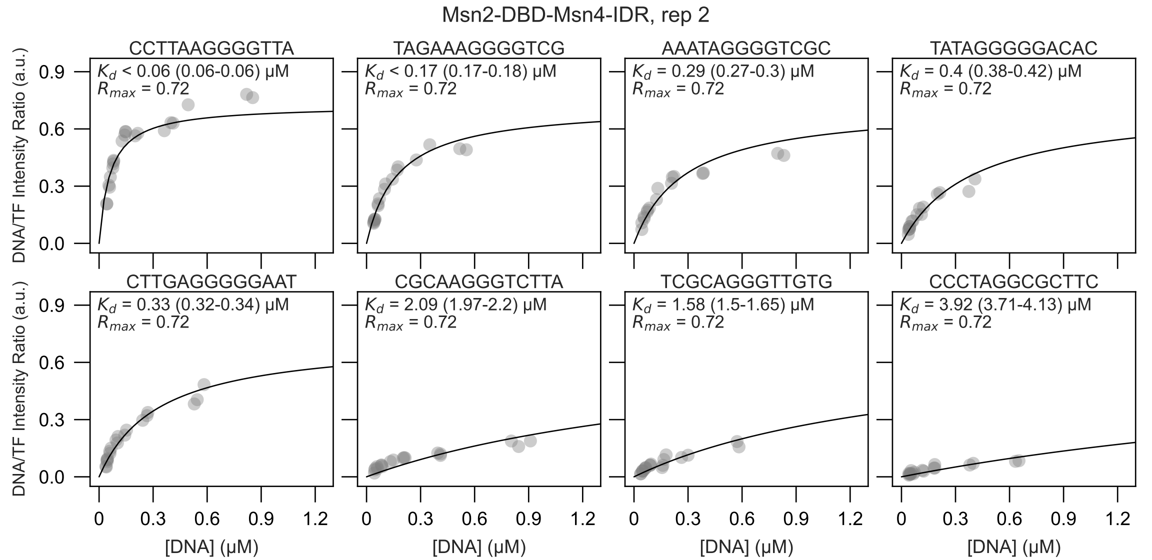

Msn2-DBD-Msn4-IDR, rep 2
CCTTAAGGGGTTA
$K_d$ < 0.06 (0.06-0.06) µM
$R_{max}$ = 0.72
TAGAAAGGGGTCG
$K_d$ < 0.17 (0.17-0.18) µM
$R_{max}$ = 0.72
AAATAGGGGTCGC
$K_d$ = 0.29 (0.27-0.3) µM
$R_{max}$ = 0.72
TATAGGGGGACAC
$K_d$ = 0.4 (0.38-0.42) µM
$R_{max}$ = 0.72
CTTGAGGGGGAAT
$K_d$ = 0.33 (0.32-0.34) µM
$R_{max}$ = 0.72
CGCAAGGGTCTTA
$K_d$ = 2.09 (1.97-2.2) µM
$R_{max}$ = 0.72
TCGCAGGGTTGTG
$K_d$ = 1.58 (1.5-1.65) µM
$R_{max}$ = 0.72
CCCTAGGCGCTTC
$K_d$ = 3.92 (3.71-4.13) µM
$R_{max}$ = 0.72
DNA/TF Intensity Ratio (a.u.)
0.0
0.3
0.6
0.9
[DNA] (µM)
0
0.3
0.6
0.9
1.2

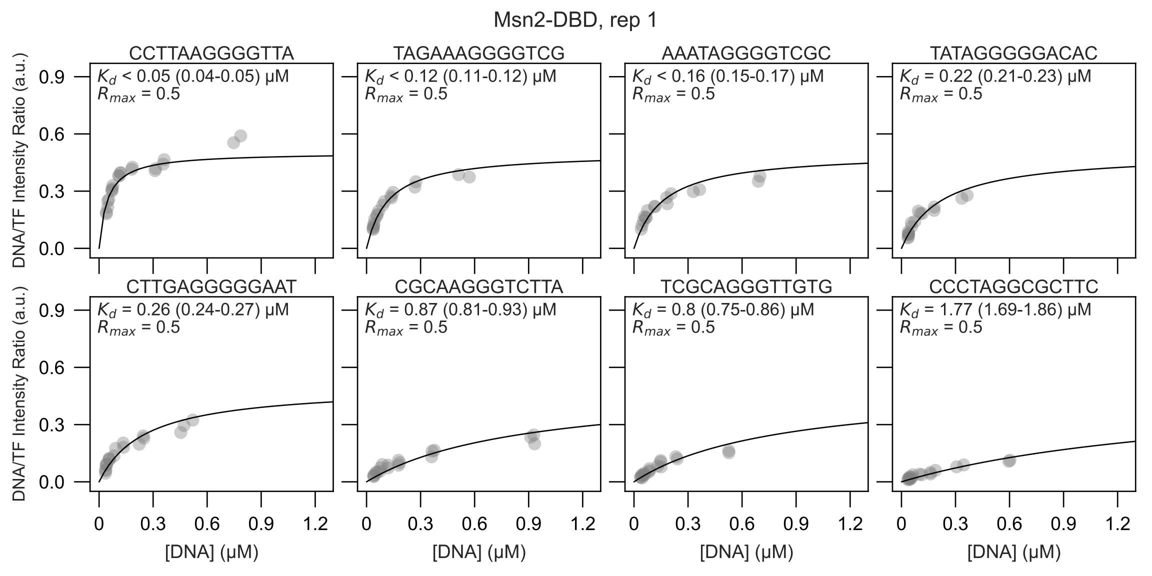

Msn2-DBD, rep 1
CCTTAAGGGGTTA
Kd < 0.05 (0.04-0.05) μM
Rmax = 0.5
TAGAAAGGGGTCG
Kd < 0.12 (0.11-0.12) μM
Rmax = 0.5
AAATAGGGGTCGC
Kd < 0.16 (0.15-0.17) μM
Rmax = 0.5
TATAGGGGGACAC
Kd = 0.22 (0.21-0.23) μM
Rmax = 0.5
CTTGAGGGGGAAT
Kd = 0.26 (0.24-0.27) μM
Rmax = 0.5
CGCAAGGGTCTTA
Kd = 0.87 (0.81-0.93) μM
Rmax = 0.5
TCGCAGGGTTGTG
Kd = 0.8 (0.75-0.86) μM
Rmax = 0.5
CCCTAGGCGCTTC
Kd = 1.77 (1.69-1.86) μM
Rmax = 0.5
DNA/TF Intensity Ratio (a.u.)
[DNA] (μM)
0.0
0.3
0.6
0.9
0
1.2

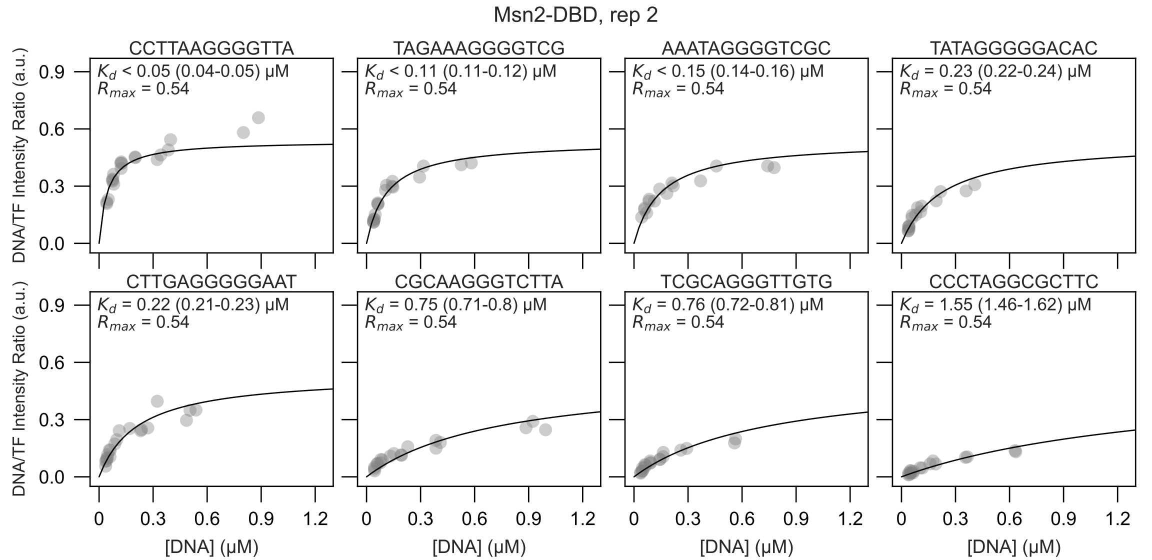

Msn2-DBD, rep 2
CCTTAAGGGGTTA
$K_d$ < 0.05 (0.04-0.05) μM
$R_{max}$ = 0.54
TAGAAAGGGGTCG
$K_d$ < 0.11 (0.11-0.12) μM
$R_{max}$ = 0.54
AAATAGGGGTCGC
$K_d$ < 0.15 (0.14-0.16) μM
$R_{max}$ = 0.54
TATAGGGGGACAC
$K_d$ = 0.23 (0.22-0.24) μM
$R_{max}$ = 0.54
CTTGAGGGGGAAT
$K_d$ = 0.22 (0.21-0.23) μM
$R_{max}$ = 0.54
CGCAAGGGTCTTA
$K_d$ = 0.75 (0.71-0.8) μM
$R_{max}$ = 0.54
TCGCAGGGTTGTG
$K_d$ = 0.76 (0.72-0.81) μM
$R_{max}$ = 0.54
CCCTAGGCGCTTC
$K_d$ = 1.55 (1.46-1.62) μM
$R_{max}$ = 0.54
DNA/TF Intensity Ratio (a.u.)
0.0
0.3
0.6
0.9
[DNA] (μM)
0
0.3
0.6
0.9
1.2

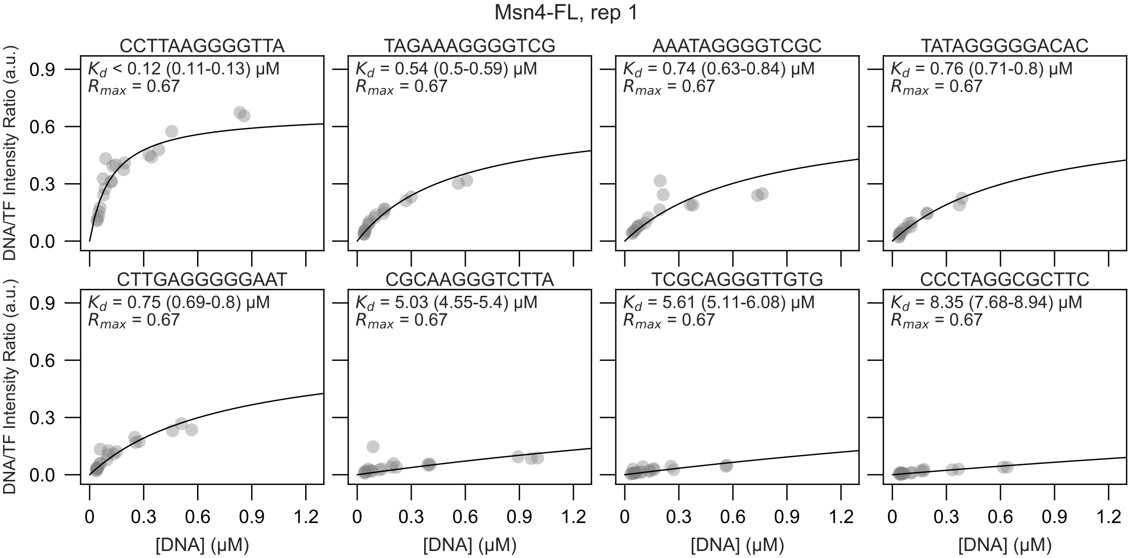

Msn4-FL, rep 1
CCTTAAGGGGTTA
$K_d$ < 0.12 (0.11-0.13) μM
$R_{max}$ = 0.67
TAGAAAGGGGTCG
$K_d$ = 0.54 (0.5-0.59) μM
$R_{max}$ = 0.67
AAATAGGGGTCGC
$K_d$ = 0.74 (0.63-0.84) μM
$R_{max}$ = 0.67
TATAGGGGGACAC
$K_d$ = 0.76 (0.71-0.8) μM
$R_{max}$ = 0.67
CTTGAGGGGGAAT
$K_d$ = 0.75 (0.69-0.8) μM
$R_{max}$ = 0.67
CGCAAGGGTCTTA
$K_d$ = 5.03 (4.55-5.4) μM
$R_{max}$ = 0.67
TCGCAGGGTTGTG
$K_d$ = 5.61 (5.11-6.08) μM
$R_{max}$ = 0.67
CCCTAGGCGCTTC
$K_d$ = 8.35 (7.68-8.94) μM
$R_{max}$ = 0.67
DNA/TF Intensity Ratio (a.u.)
0.0
0.3
0.6
0.9
[DNA] (μM)
0
0.3
0.6
0.9
1.2

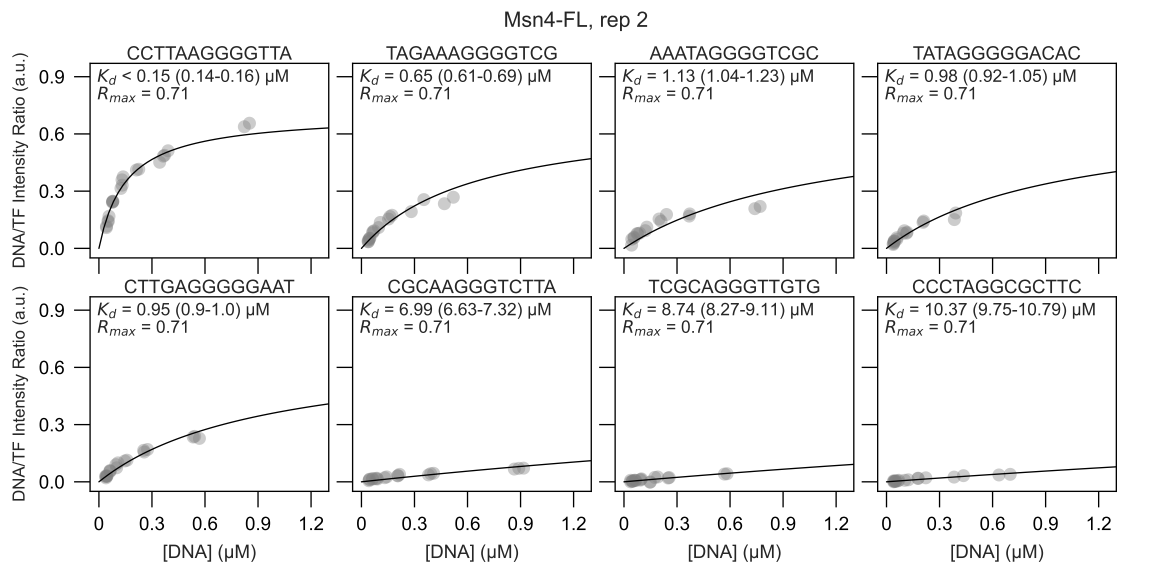

Msn4-FL, rep 2
CCTTAAGGGGTTA
$K_d$ < 0.15 (0.14-0.16) µM
$R_{max}$ = 0.71
TAGAAAGGGGTCG
$K_d$ = 0.65 (0.61-0.69) µM
$R_{max}$ = 0.71
AAATAGGGGTCGC
$K_d$ = 1.13 (1.04-1.23) µM
$R_{max}$ = 0.71
TATAGGGGGACAC
$K_d$ = 0.98 (0.92-1.05) µM
$R_{max}$ = 0.71
CTTGAGGGGGAAT
$K_d$ = 0.95 (0.9-1.0) µM
$R_{max}$ = 0.71
CGCAAGGGTCTTA
$K_d$ = 6.99 (6.63-7.32) µM
$R_{max}$ = 0.71
TCGCAGGGTTGTG
$K_d$ = 8.74 (8.27-9.11) µM
$R_{max}$ = 0.71
CCCTAGGCGCTTC
$K_d$ = 10.37 (9.75-10.79) µM
$R_{max}$ = 0.71
DNA/TF Intensity Ratio (a.u.)
0.0
0.3
0.6
0.9
[DNA] (µM)
0
0.3
0.6
0.9
1.2

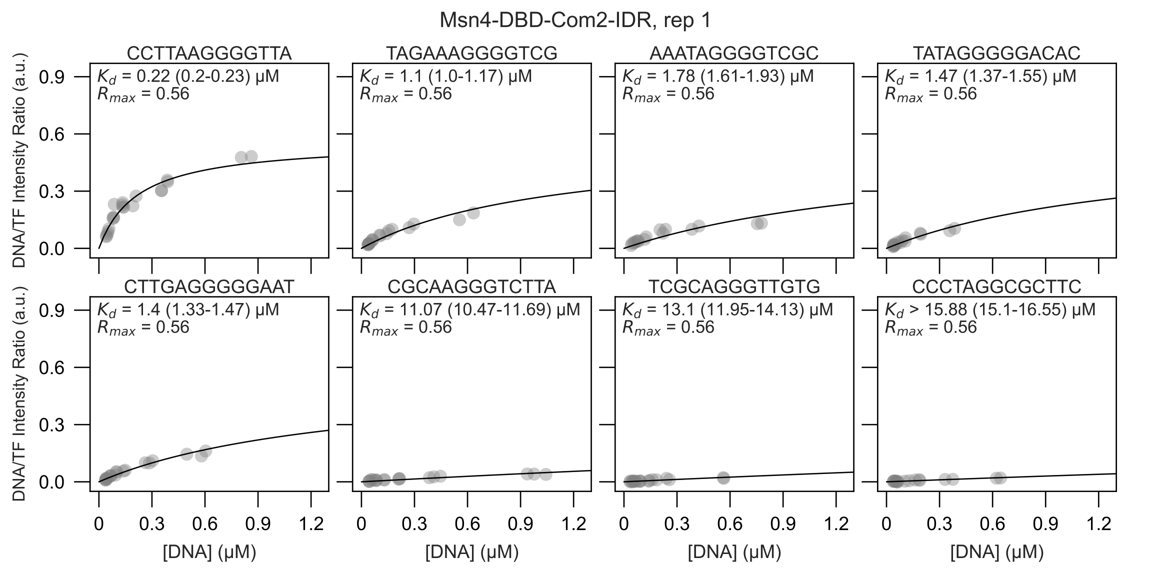

Msn4-DBD-Com2-IDR, rep 1
CCTTAAGGGGTTA
$K_d$ = 0.22 (0.2-0.23) μM
$R_{max}$ = 0.56
TAGAAAGGGGTCG
$K_d$ = 1.1 (1.0-1.17) μM
$R_{max}$ = 0.56
AAATAGGGGTCGC
$K_d$ = 1.78 (1.61-1.93) μM
$R_{max}$ = 0.56
TATAGGGGGACAC
$K_d$ = 1.47 (1.37-1.55) μM
$R_{max}$ = 0.56
CTTGAGGGGGAAT
$K_d$ = 1.4 (1.33-1.47) μM
$R_{max}$ = 0.56
CGCAAGGGTCTTA
$K_d$ = 11.07 (10.47-11.69) μM
$R_{max}$ = 0.56
TCGCAGGGTTGTG
$K_d$ = 13.1 (11.95-14.13) μM
$R_{max}$ = 0.56
CCCTAGGCGCTTC
$K_d$ > 15.88 (15.1-16.55) μM
$R_{max}$ = 0.56
DNA/TF Intensity Ratio (a.u.)
[DNA] (μM)
0.0
0.3
0.6
0.9
0
1.2

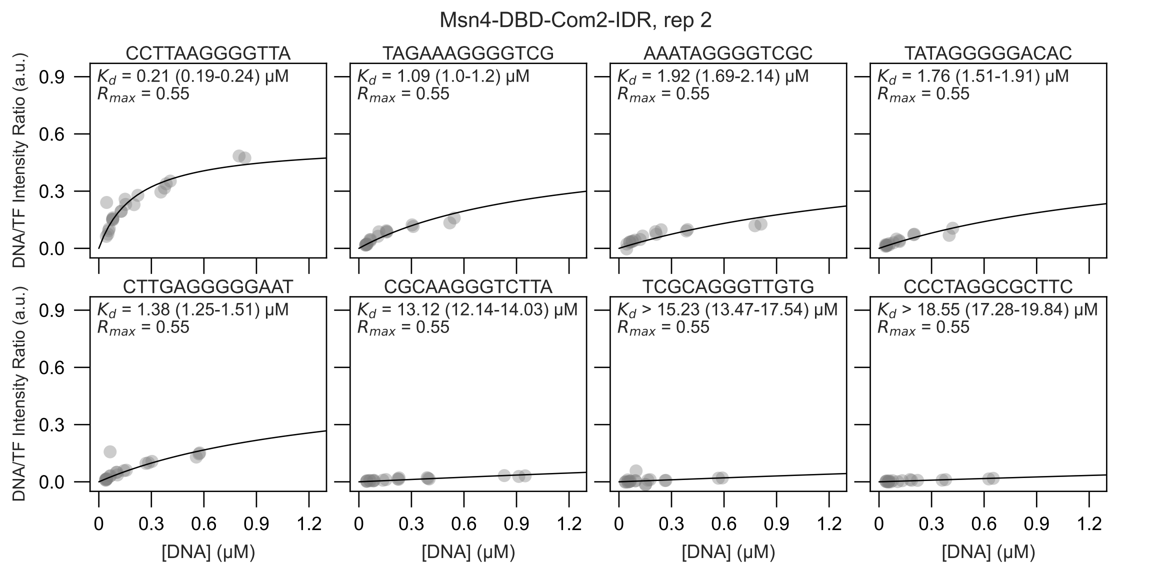

Msn4-DBD-Com2-IDR, rep 2
CCTTAAGGGGTTA
$K_d$ = 0.21 (0.19-0.24) µM
$R_{max}$ = 0.55
TAGAAAGGGGTCG
$K_d$ = 1.09 (1.0-1.2) µM
$R_{max}$ = 0.55
AAATAGGGGTCGC
$K_d$ = 1.92 (1.69-2.14) µM
$R_{max}$ = 0.55
TATAGGGGGACAC
$K_d$ = 1.76 (1.51-1.91) µM
$R_{max}$ = 0.55
CTTGAGGGGGAAT
$K_d$ = 1.38 (1.25-1.51) µM
$R_{max}$ = 0.55
CGCAAGGGTCTTA
$K_d$ = 13.12 (12.14-14.03) µM
$R_{max}$ = 0.55
TCGCAGGGTTGTG
$K_d$ > 15.23 (13.47-17.54) µM
$R_{max}$ = 0.55
CCCTAGGCGCTTC
$K_d$ > 18.55 (17.28-19.84) µM
$R_{max}$ = 0.55
DNA/TF Intensity Ratio (a.u.)
0.0
0.3
0.6
0.9
[DNA] (µM)
0
0.3
0.6
0.9
1.2

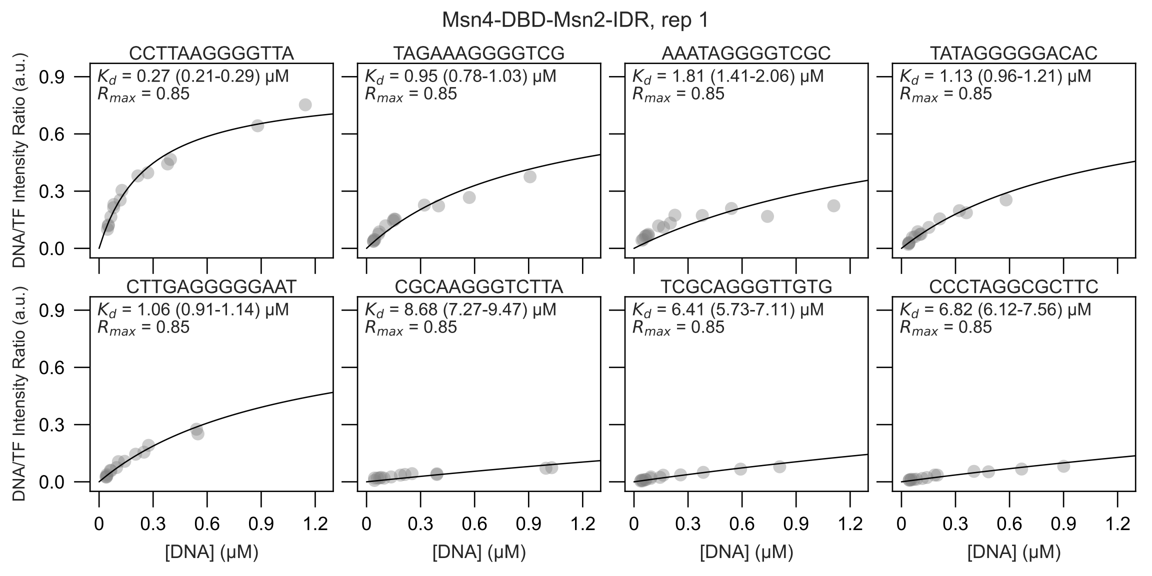

Msn4-DBD-Msn2-IDR, rep 1
CCTTAAGGGGTTA
$K_d$ = 0.27 (0.21-0.29) μM
$R_{max}$ = 0.85
TAGAAAGGGGTCG
$K_d$ = 0.95 (0.78-1.03) μM
$R_{max}$ = 0.85
AAATAGGGGTCGC
$K_d$ = 1.81 (1.41-2.06) μM
$R_{max}$ = 0.85
TATAGGGGGACAC
$K_d$ = 1.13 (0.96-1.21) μM
$R_{max}$ = 0.85
CTTGAGGGGGAAT
$K_d$ = 1.06 (0.91-1.14) μM
$R_{max}$ = 0.85
CGCAAGGGTCTTA
$K_d$ = 8.68 (7.27-9.47) μM
$R_{max}$ = 0.85
TCGCAGGGTTGTG
$K_d$ = 6.41 (5.73-7.11) μM
$R_{max}$ = 0.85
CCCTAGGCGCTTC
$K_d$ = 6.82 (6.12-7.56) μM
$R_{max}$ = 0.85
DNA/TF Intensity Ratio (a.u.)
0.0
0.3
0.6
0.9
[DNA] (μM)
0
0.3
0.6
0.9
1.2

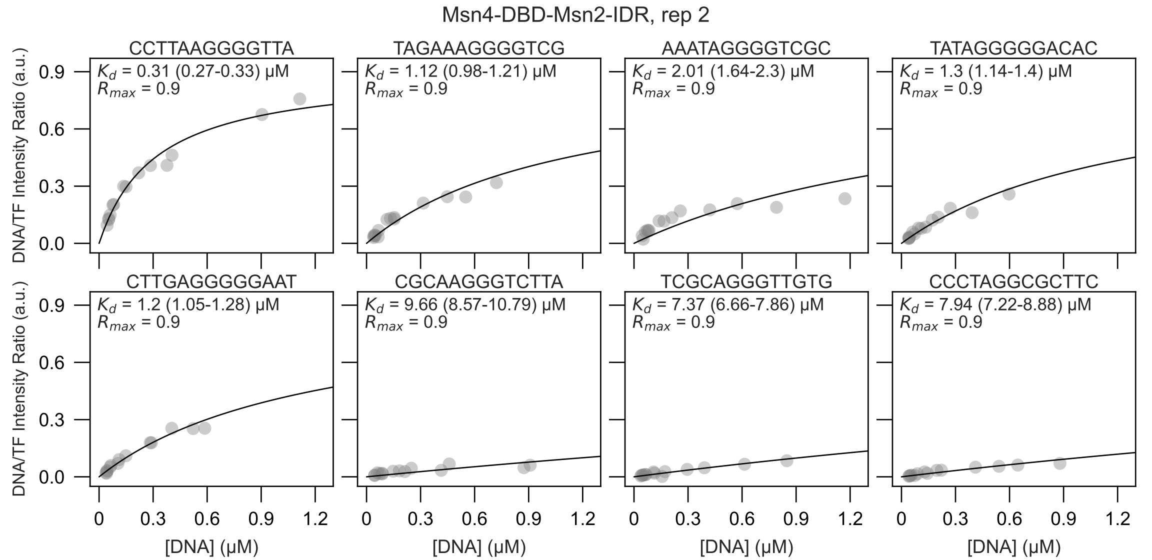

Msn4-DBD-Msn2-IDR, rep 2
CCTTAAGGGGTTA
$K_d$ = 0.31 (0.27-0.33) μM
$R_{max}$ = 0.9
TAGAAAGGGGTCG
$K_d$ = 1.12 (0.98-1.21) μM
$R_{max}$ = 0.9
AAATAGGGGTCGC
$K_d$ = 2.01 (1.64-2.3) μM
$R_{max}$ = 0.9
TATAGGGGGACAC
$K_d$ = 1.3 (1.14-1.4) μM
$R_{max}$ = 0.9
CTTGAGGGGGAAT
$K_d$ = 1.2 (1.05-1.28) μM
$R_{max}$ = 0.9
CGCAAGGGTCTTA
$K_d$ = 9.66 (8.57-10.79) μM
$R_{max}$ = 0.9
TCGCAGGGTTGTG
$K_d$ = 7.37 (6.66-7.86) μM
$R_{max}$ = 0.9
CCCTAGGCGCTTC
$K_d$ = 7.94 (7.22-8.88) μM
$R_{max}$ = 0.9
DNA/TF Intensity Ratio (a.u.)
0.0
0.3
0.6
0.9
[DNA] (μM)
0
0.3
0.6
0.9
1.2

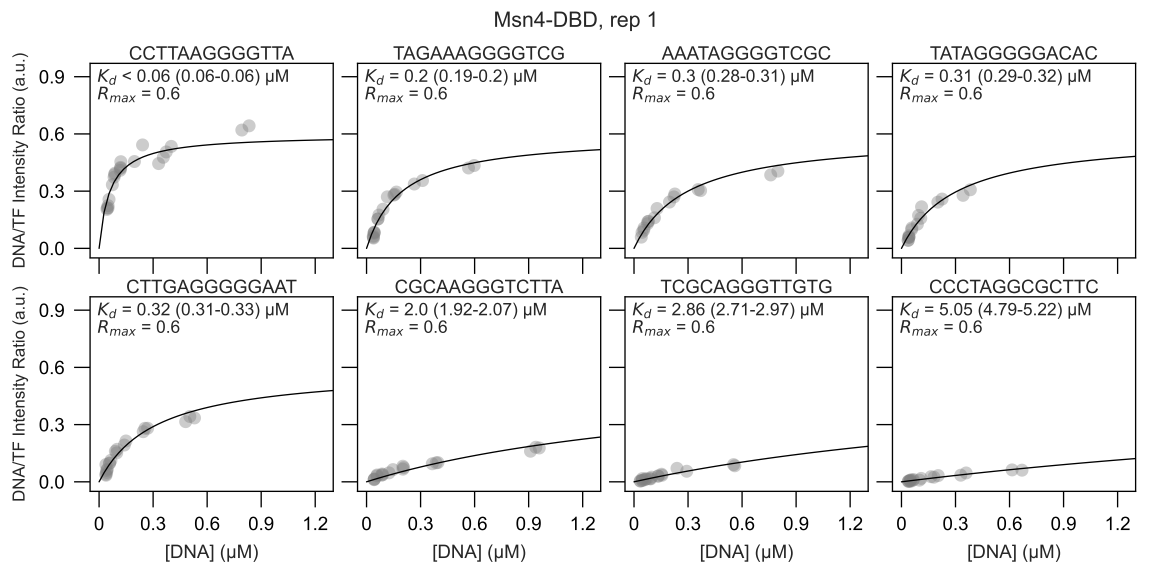

Msn4-DBD, rep 1
CCTTAAGGGGTTA
Kd < 0.06 (0.06-0.06) μM
Rmax = 0.6
TAGAAAGGGGTCG
Kd = 0.2 (0.19-0.2) μM
Rmax = 0.6
AAATAGGGGTCGC
Kd = 0.3 (0.28-0.31) μM
Rmax = 0.6
TATAGGGGGACAC
Kd = 0.31 (0.29-0.32) μM
Rmax = 0.6
CTTGAGGGGGAAT
Kd = 0.32 (0.31-0.33) μM
Rmax = 0.6
CGCAAGGGTCTTA
Kd = 2.0 (1.92-2.07) μM
Rmax = 0.6
TCGCAGGGTTGTG
Kd = 2.86 (2.71-2.97) μM
Rmax = 0.6
CCCTAGGCGCTTC
Kd = 5.05 (4.79-5.22) μM
Rmax = 0.6
DNA/TF Intensity Ratio (a.u.)
0.0
0.3
0.6
0.9
[DNA] (μM)
0
0.3
0.6
0.9
1.2

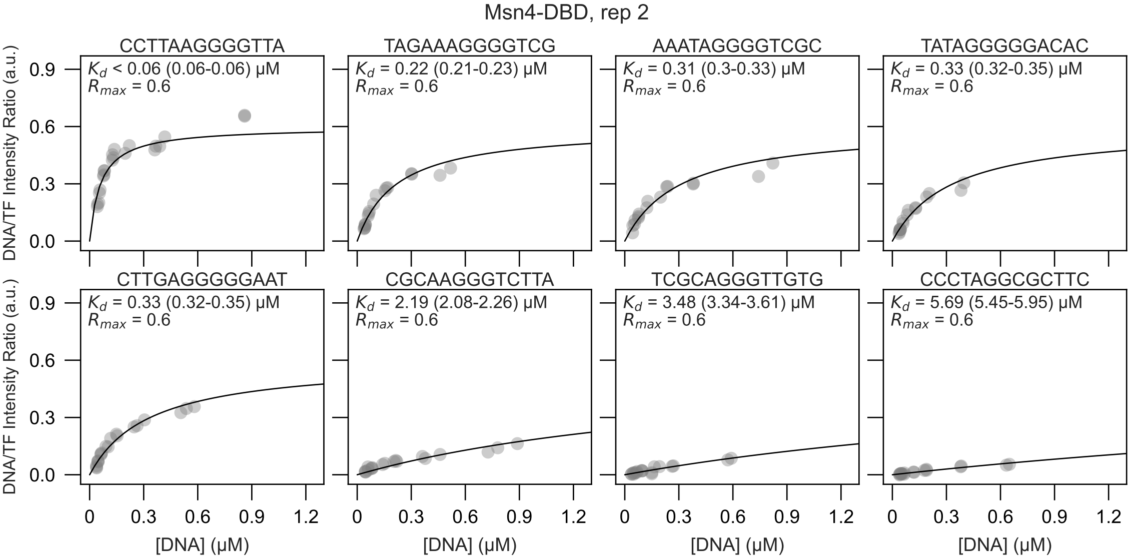
Msn4-DBD, rep 2
CCTTAAGGGGTTA
$K_d$ < 0.06 (0.06-0.06) μM
$R_{max}$ = 0.6
TAGAAAGGGGTCG
$K_d$ = 0.22 (0.21-0.23) μM
$R_{max}$ = 0.6
AAATAGGGGTCGC
$K_d$ = 0.31 (0.3-0.33) μM
$R_{max}$ = 0.6
TATAGGGGGACAC
$K_d$ = 0.33 (0.32-0.35) μM
$R_{max}$ = 0.6
CTTGAGGGGGAAT
$K_d$ = 0.33 (0.32-0.35) μM
$R_{max}$ = 0.6
CGCAAGGGTCTTA
$K_d$ = 2.19 (2.08-2.26) μM
$R_{max}$ = 0.6
TCGCAGGGTTGTG
$K_d$ = 3.48 (3.34-3.61) μM
$R_{max}$ = 0.6
CCCTAGGCGCTTC
$K_d$ = 5.69 (5.45-5.95) μM
$R_{max}$ = 0.6
DNA/TF Intensity Ratio (a.u.)
0.0
0.3
0.6
0.9
[DNA] (μM)
0
0.3
0.6
0.9
1.2
