## Supplementary File 2 for "Intrinsically disordered regions within transcription factors impact DNA binding through both autoinhibition and independent DNA binding"

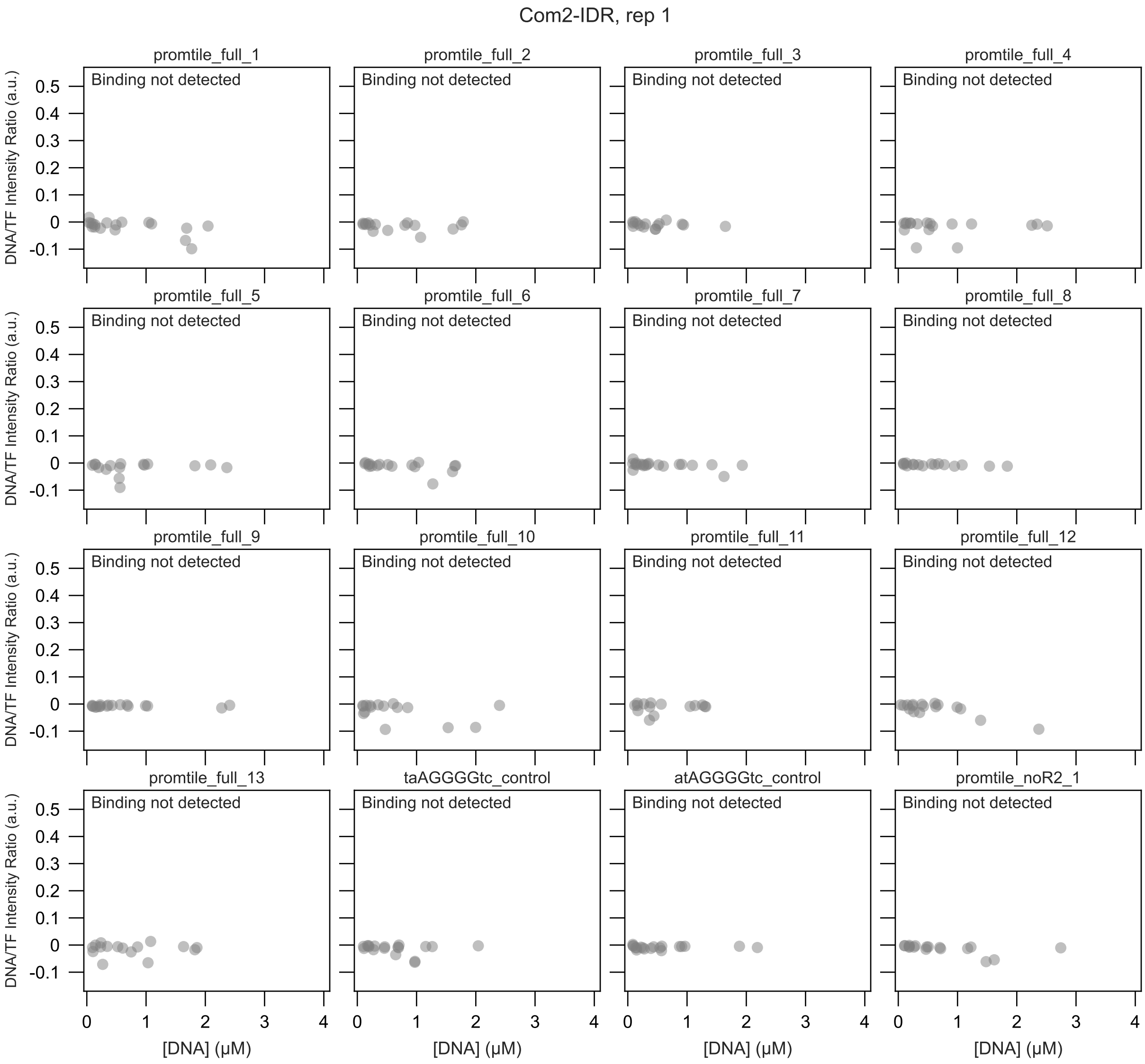
Com2-IDR, rep 1
promtile_full_1
promtile_full_2
promtile_full_3
promtile_full_4
promtile_full_5
promtile_full_6
promtile_full_7
promtile_full_8
promtile_full_9
promtile_full_10
promtile_full_11
promtile_full_12
promtile_full_13
taAGGGGtc_control
atAGGGGtc_control
promtile_noR2_1
Binding not detected
DNA/TF Intensity Ratio (a.u.)
[DNA] (μM)
0.5
0.4
0.3
0.2
0.1
0
-0.1
0
1
2
3
4

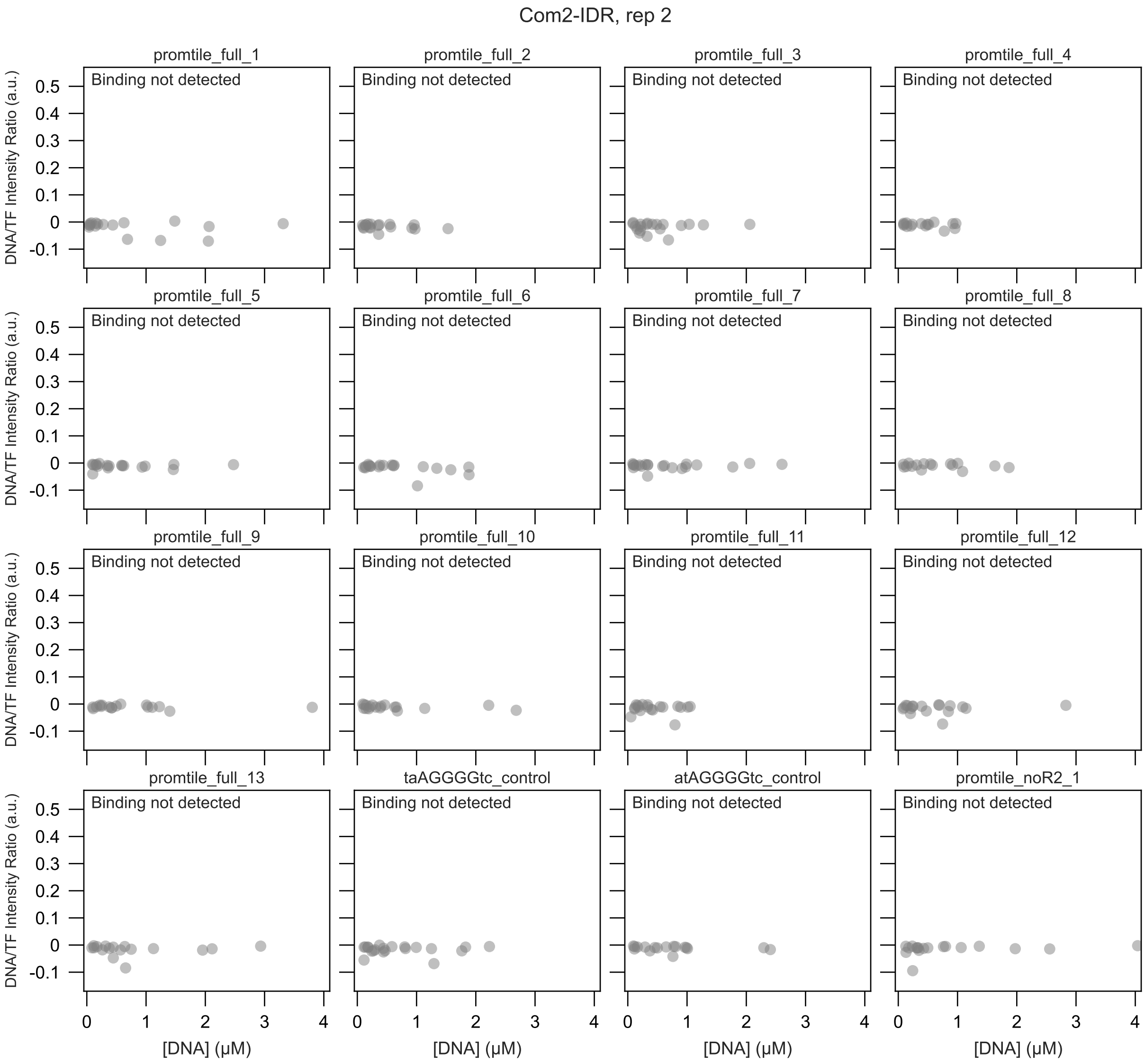

Com2-IDR, rep 2
promtile_full_1
promtile_full_2
promtile_full_3
promtile_full_4
promtile_full_5
promtile_full_6
promtile_full_7
promtile_full_8
promtile_full_9
promtile_full_10
promtile_full_11
promtile_full_12
promtile_full_13
taAGGGGtc_control
atAGGGGtc_control
promtile_noR2_1
Binding not detected
DNA/TF Intensity Ratio (a.u.)
[DNA] (μM)

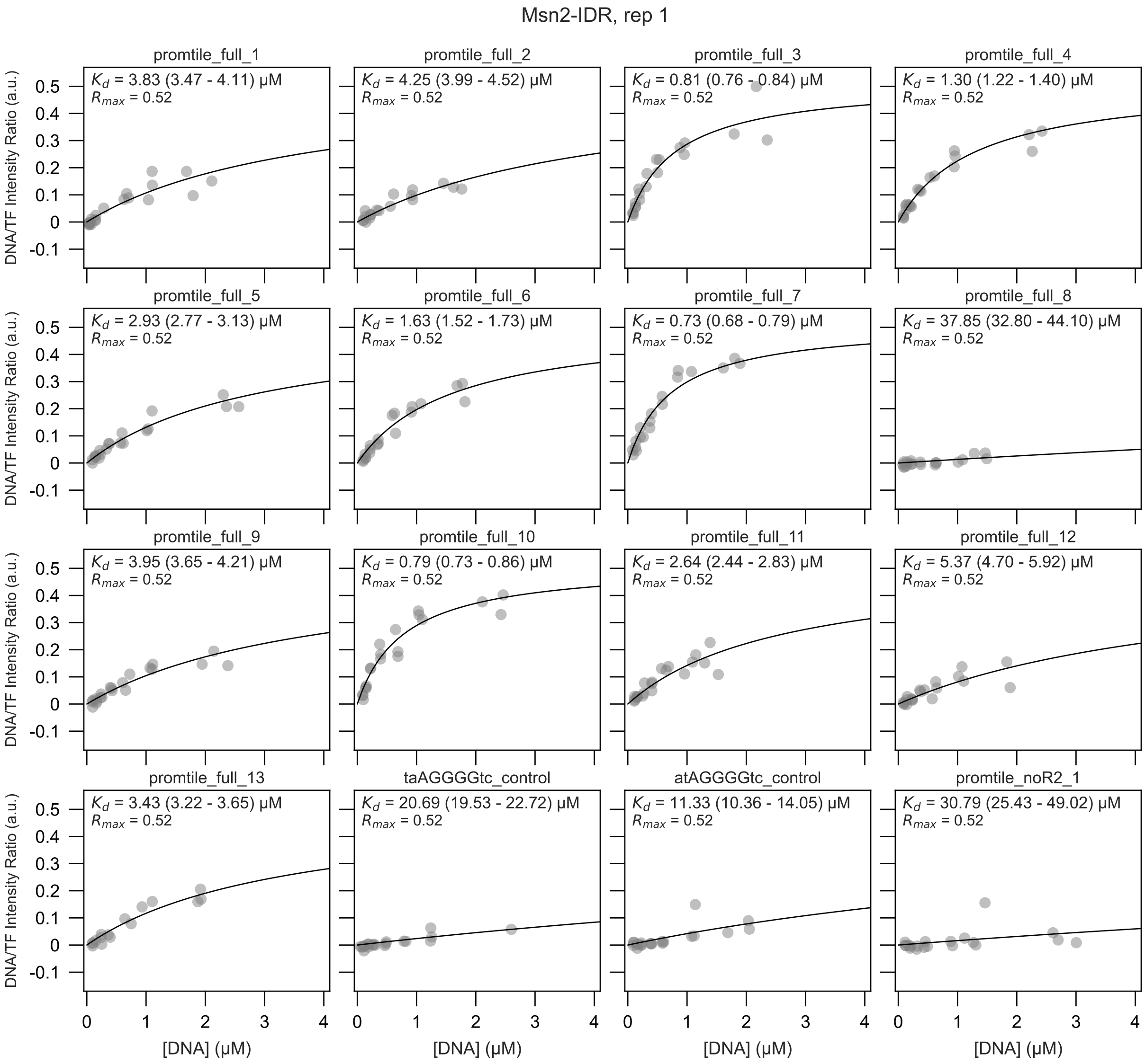

Msn2-IDR, rep 1
promtile_full_1
$K_d$ = 3.83 (3.47 - 4.11) μM
$R_{max}$ = 0.52
promtile_full_2
$K_d$ = 4.25 (3.99 - 4.52) μM
$R_{max}$ = 0.52
promtile_full_3
$K_d$ = 0.81 (0.76 - 0.84) μM
$R_{max}$ = 0.52
promtile_full_4
$K_d$ = 1.30 (1.22 - 1.40) μM
$R_{max}$ = 0.52
promtile_full_5
$K_d$ = 2.93 (2.77 - 3.13) μM
$R_{max}$ = 0.52
promtile_full_6
$K_d$ = 1.63 (1.52 - 1.73) μM
$R_{max}$ = 0.52
promtile_full_7
$K_d$ = 0.73 (0.68 - 0.79) μM
$R_{max}$ = 0.52
promtile_full_8
$K_d$ = 37.85 (32.80 - 44.10) μM
$R_{max}$ = 0.52
promtile_full_9
$K_d$ = 3.95 (3.65 - 4.21) μM
$R_{max}$ = 0.52
promtile_full_10
$K_d$ = 0.79 (0.73 - 0.86) μM
$R_{max}$ = 0.52
promtile_full_11
$K_d$ = 2.64 (2.44 - 2.83) μM
$R_{max}$ = 0.52
promtile_full_12
$K_d$ = 5.37 (4.70 - 5.92) μM
$R_{max}$ = 0.52
promtile_full_13
$K_d$ = 3.43 (3.22 - 3.65) μM
$R_{max}$ = 0.52
taAGGGGtc_control
$K_d$ = 20.69 (19.53 - 22.72) μM
$R_{max}$ = 0.52
atAGGGGtc_control
$K_d$ = 11.33 (10.36 - 14.05) μM
$R_{max}$ = 0.52
promtile_noR2_1
$K_d$ = 30.79 (25.43 - 49.02) μM
$R_{max}$ = 0.52
DNA/TF Intensity Ratio (a.u.)
0.5
0.4
0.3
0.2
0.1
0
-0.1
[DNA] (μM)
0
1
2
3
4

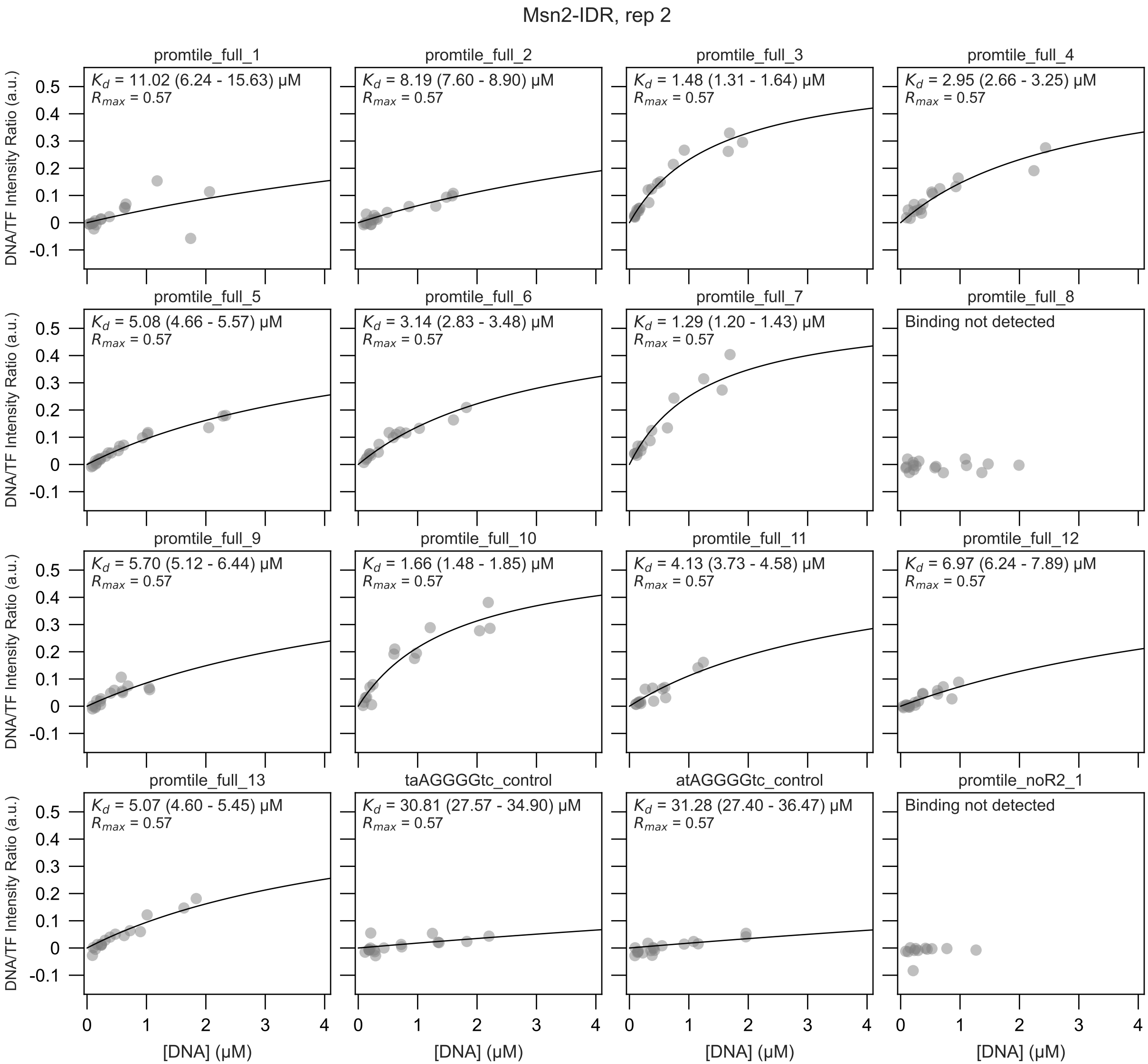
Msn2-IDR, rep 2
promtile_full_1
$K_d$ = 11.02 (6.24 - 15.63) μM
$R_{max}$ = 0.57
promtile_full_2
$K_d$ = 8.19 (7.60 - 8.90) μM
$R_{max}$ = 0.57
promtile_full_3
$K_d$ = 1.48 (1.31 - 1.64) μM
$R_{max}$ = 0.57
promtile_full_4
$K_d$ = 2.95 (2.66 - 3.25) μM
$R_{max}$ = 0.57
promtile_full_5
$K_d$ = 5.08 (4.66 - 5.57) μM
$R_{max}$ = 0.57
promtile_full_6
$K_d$ = 3.14 (2.83 - 3.48) μM
$R_{max}$ = 0.57
promtile_full_7
$K_d$ = 1.29 (1.20 - 1.43) μM
$R_{max}$ = 0.57
promtile_full_8
Binding not detected
promtile_full_9
$K_d$ = 5.70 (5.12 - 6.44) μM
$R_{max}$ = 0.57
promtile_full_10
$K_d$ = 1.66 (1.48 - 1.85) μM
$R_{max}$ = 0.57
promtile_full_11
$K_d$ = 4.13 (3.73 - 4.58) μM
$R_{max}$ = 0.57
promtile_full_12
$K_d$ = 6.97 (6.24 - 7.89) μM
$R_{max}$ = 0.57
promtile_full_13
$K_d$ = 5.07 (4.60 - 5.45) μM
$R_{max}$ = 0.57
taAGGGGtc_control
$K_d$ = 30.81 (27.57 - 34.90) μM
$R_{max}$ = 0.57
atAGGGGtc_control
$K_d$ = 31.28 (27.40 - 36.47) μM
$R_{max}$ = 0.57
promtile_noR2_1
Binding not detected
DNA/TF Intensity Ratio (a.u.)
[DNA] (μM)
0.5
0.4
0.3
0.2
0.1
0
-0.1
0
1
2
3
4

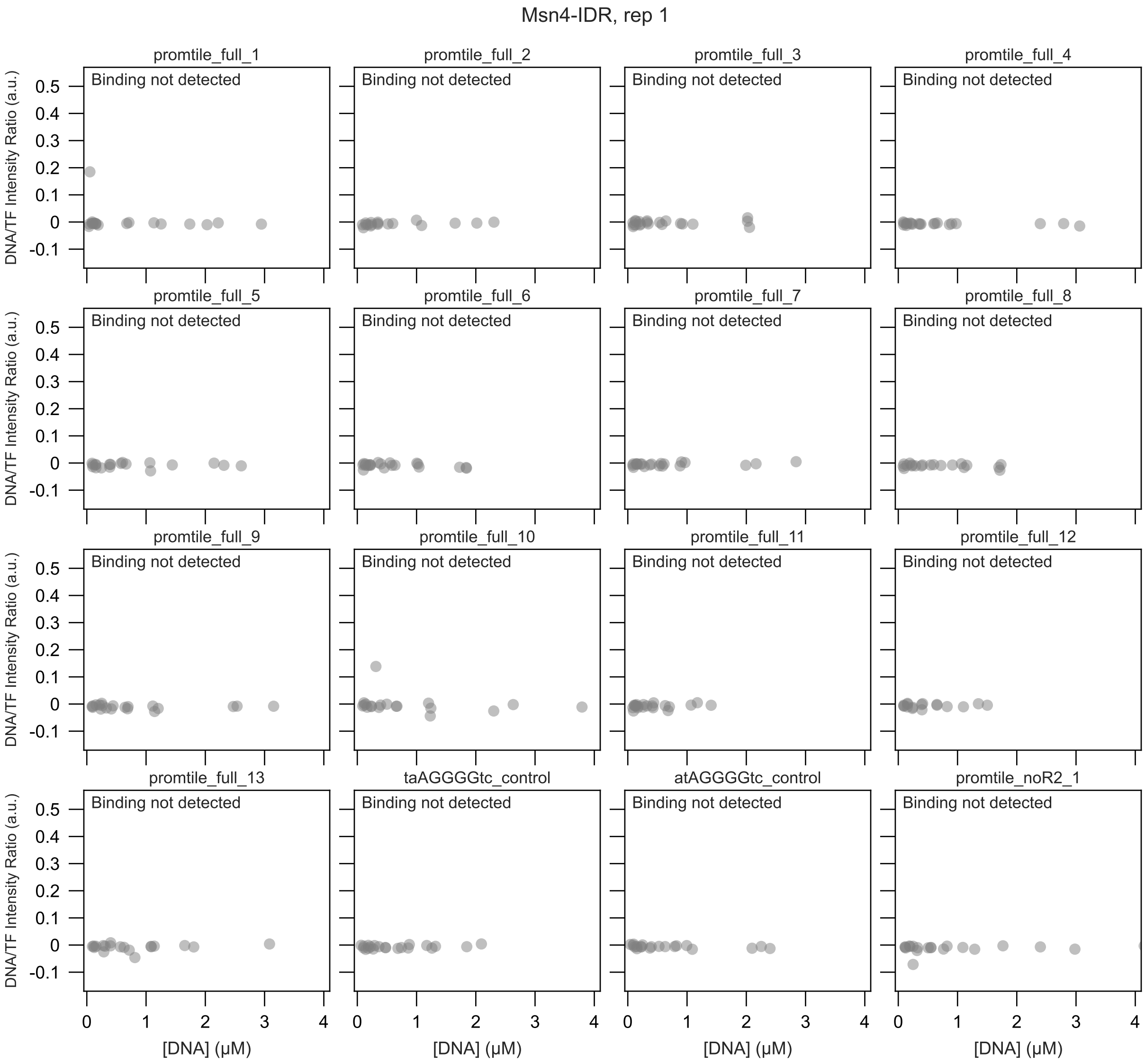

Msn4-IDR, rep 1
promtile_full_1
promtile_full_2
promtile_full_3
promtile_full_4
promtile_full_5
promtile_full_6
promtile_full_7
promtile_full_8
promtile_full_9
promtile_full_10
promtile_full_11
promtile_full_12
promtile_full_13
taAGGGGtc_control
atAGGGGtc_control
promtile_noR2_1
Binding not detected
DNA/TF Intensity Ratio (a.u.)
[DNA] (μM)
0.5
0.4
0.3
0.2
0.1
0
-0.1
0
1
2
3
4

Msn4-IDR, rep 2

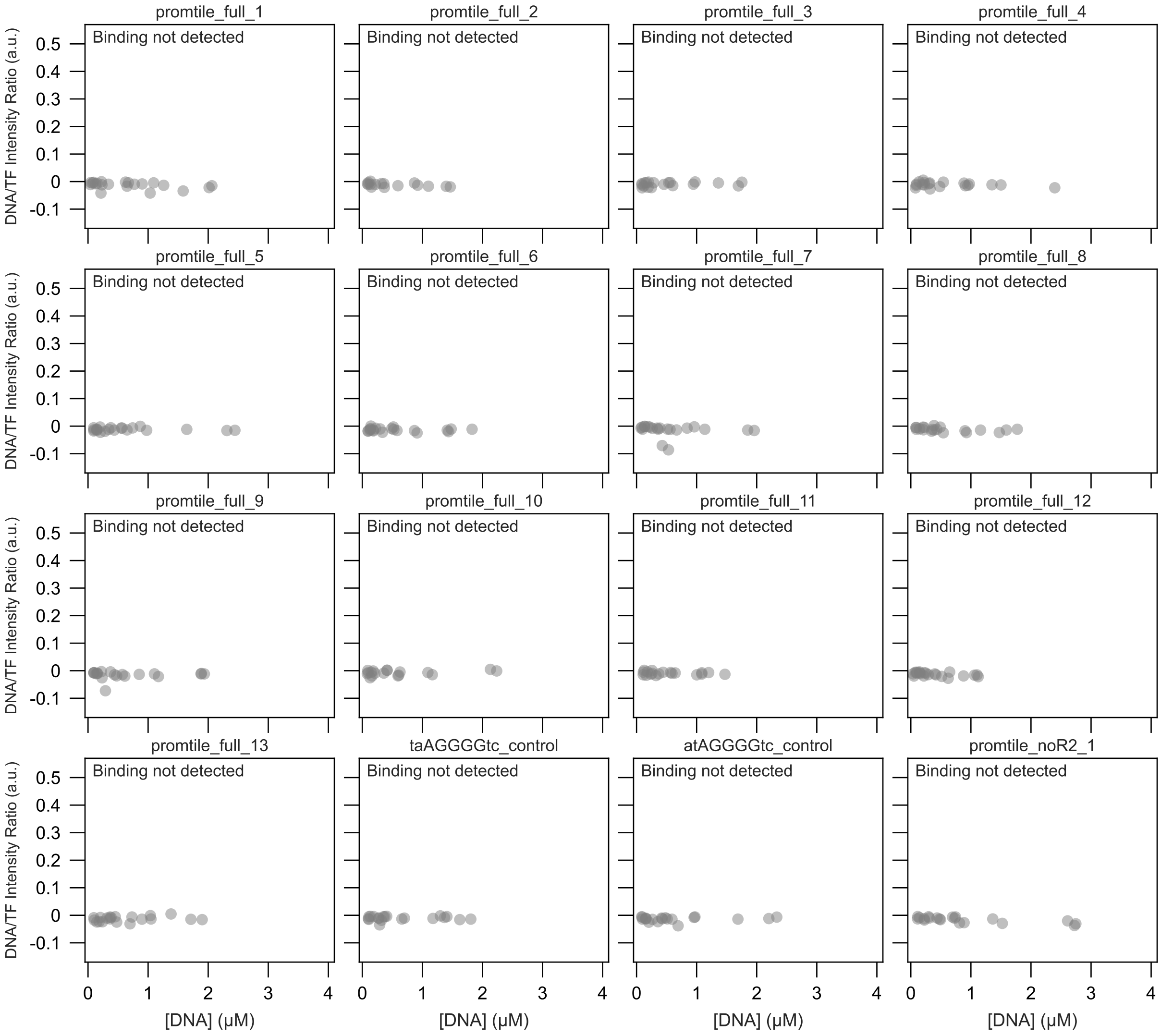
