## Supplementary File 3 for "Intrinsically disordered regions within transcription factors impact DNA binding through both autoinhibition and independent DNA binding"

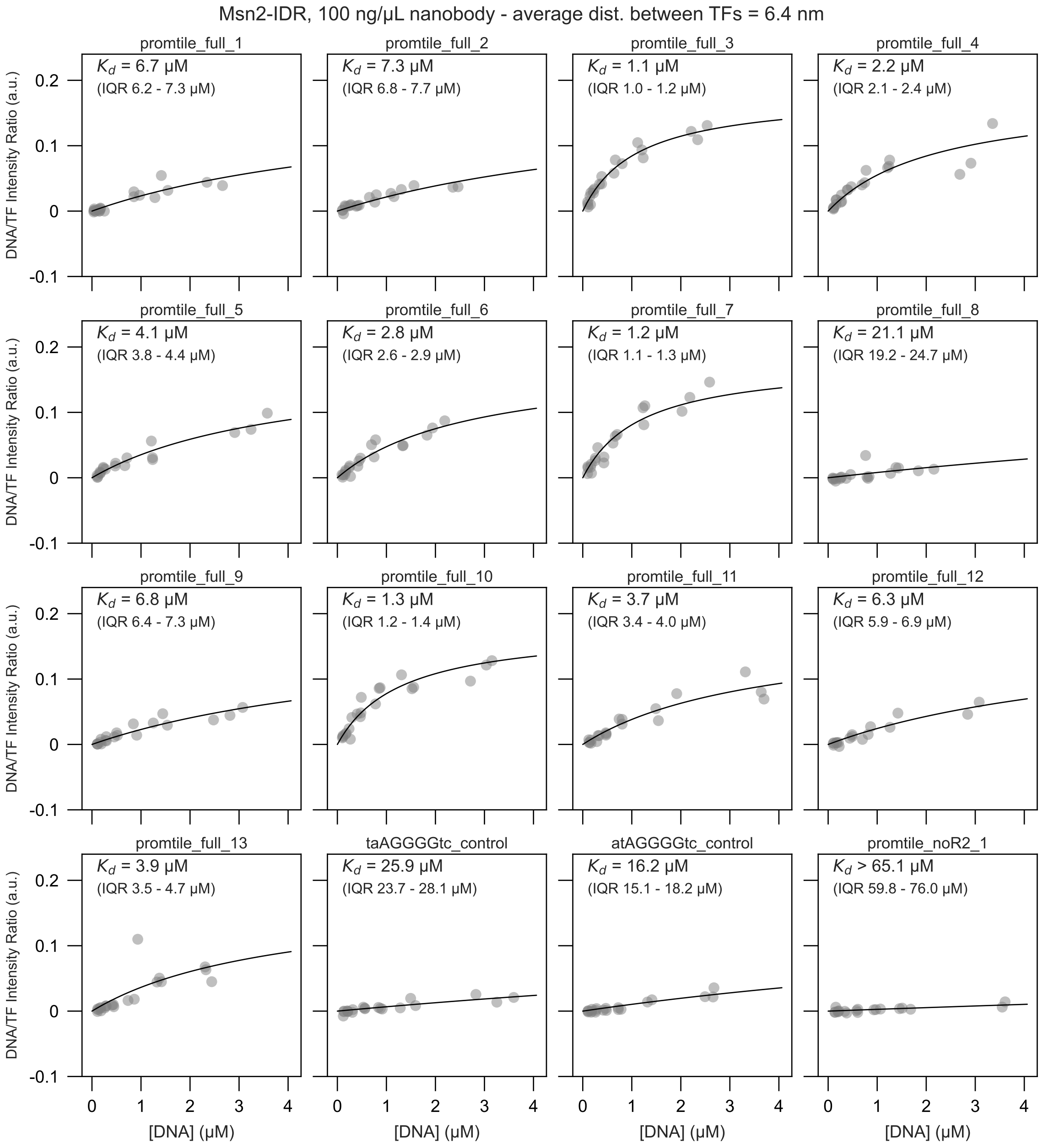

Msn2-IDR, 100 ng/µL nanobody - average dist. between TFs = 6.4 nm
promtile_full_1
$K_d$ = 6.7 µM
(IQR 6.2 - 7.3 µM)
promtile_full_2
$K_d$ = 7.3 µM
(IQR 6.8 - 7.7 µM)
promtile_full_3
$K_d$ = 1.1 µM
(IQR 1.0 - 1.2 µM)
promtile_full_4
$K_d$ = 2.2 µM
(IQR 2.1 - 2.4 µM)
promtile_full_5
$K_d$ = 4.1 µM
(IQR 3.8 - 4.4 µM)
promtile_full_6
$K_d$ = 2.8 µM
(IQR 2.6 - 2.9 µM)
promtile_full_7
$K_d$ = 1.2 µM
(IQR 1.1 - 1.3 µM)
promtile_full_8
$K_d$ = 21.1 µM
(IQR 19.2 - 24.7 µM)
promtile_full_9
$K_d$ = 6.8 µM
(IQR 6.4 - 7.3 µM)
promtile_full_10
$K_d$ = 1.3 µM
(IQR 1.2 - 1.4 µM)
promtile_full_11
$K_d$ = 3.7 µM
(IQR 3.4 - 4.0 µM)
promtile_full_12
$K_d$ = 6.3 µM
(IQR 5.9 - 6.9 µM)
promtile_full_13
$K_d$ = 3.9 µM
(IQR 3.5 - 4.7 µM)
taAGGGGtc_control
$K_d$ = 25.9 µM
(IQR 23.7 - 28.1 µM)
atAGGGGtc_control
$K_d$ = 16.2 µM
(IQR 15.1 - 18.2 µM)
promtile_noR2_1
$K_d$ > 65.1 µM
(IQR 59.8 - 76.0 µM)
DNA/TF Intensity Ratio (a.u.)
[DNA] (µM)
0.2
0.1
0
-0.1
0
1
2
3
4

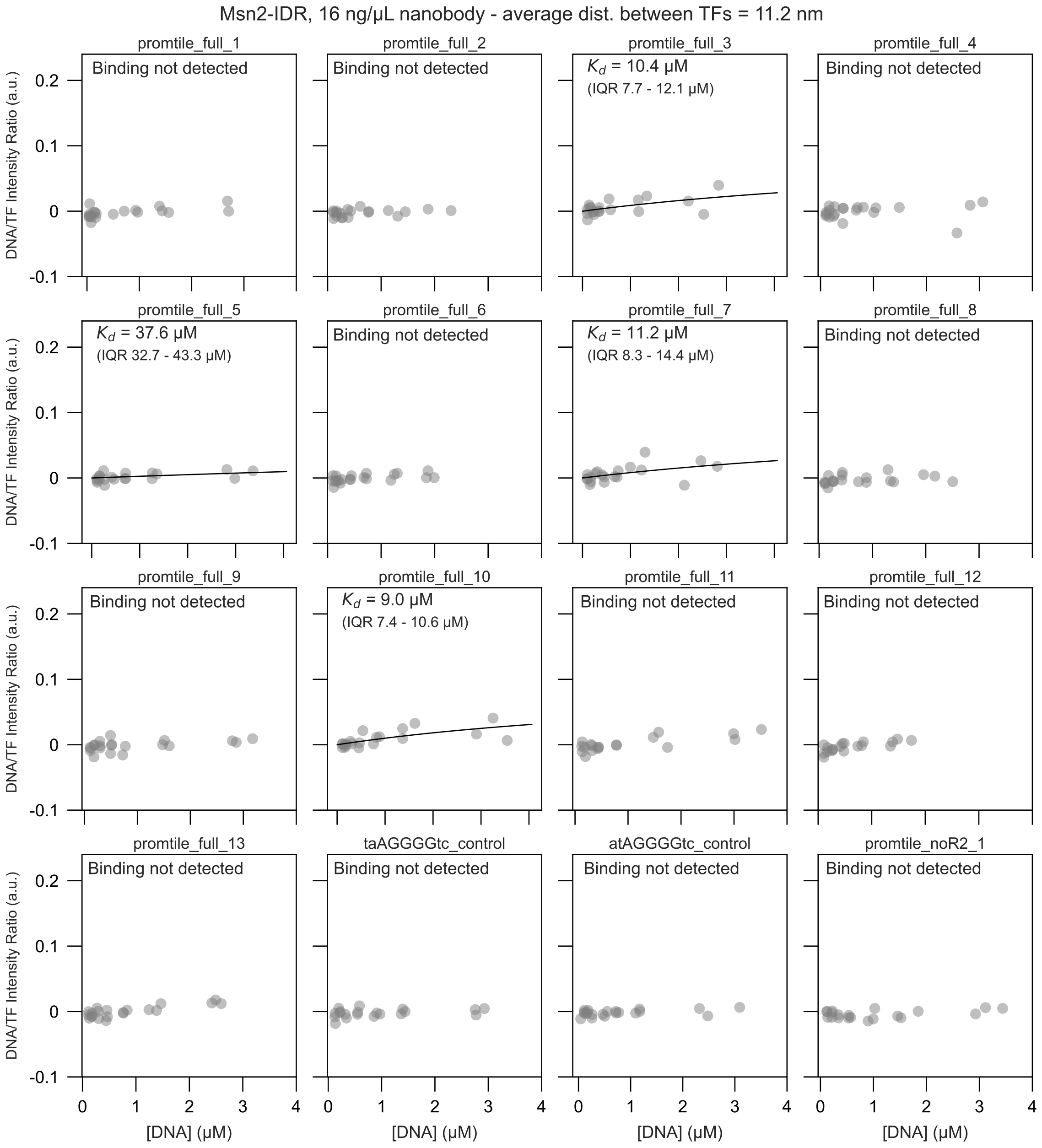
Msn2-IDR, 16 ng/μL nanobody - average dist. between TFs = 11.2 nm
promtile_full_1
Binding not detected
promtile_full_2
Binding not detected
promtile_full_3
$K_d$ = 10.4 μM
(IQR 7.7 - 12.1 μM)
promtile_full_4
Binding not detected
promtile_full_5
$K_d$ = 37.6 μM
(IQR 32.7 - 43.3 μM)
promtile_full_6
Binding not detected
promtile_full_7
$K_d$ = 11.2 μM
(IQR 8.3 - 14.4 μM)
promtile_full_8
Binding not detected
promtile_full_9
Binding not detected
promtile_full_10
$K_d$ = 9.0 μM
(IQR 7.4 - 10.6 μM)
promtile_full_11
Binding not detected
promtile_full_12
Binding not detected
promtile_full_13
Binding not detected
taAGGGGtc_control
Binding not detected
atAGGGGtc_control
Binding not detected
promtile_noR2_1
Binding not detected
DNA/TF Intensity Ratio (a.u.)
0.2
0.1
0
-0.1
0
1
2
3
4
[DNA] (μM)

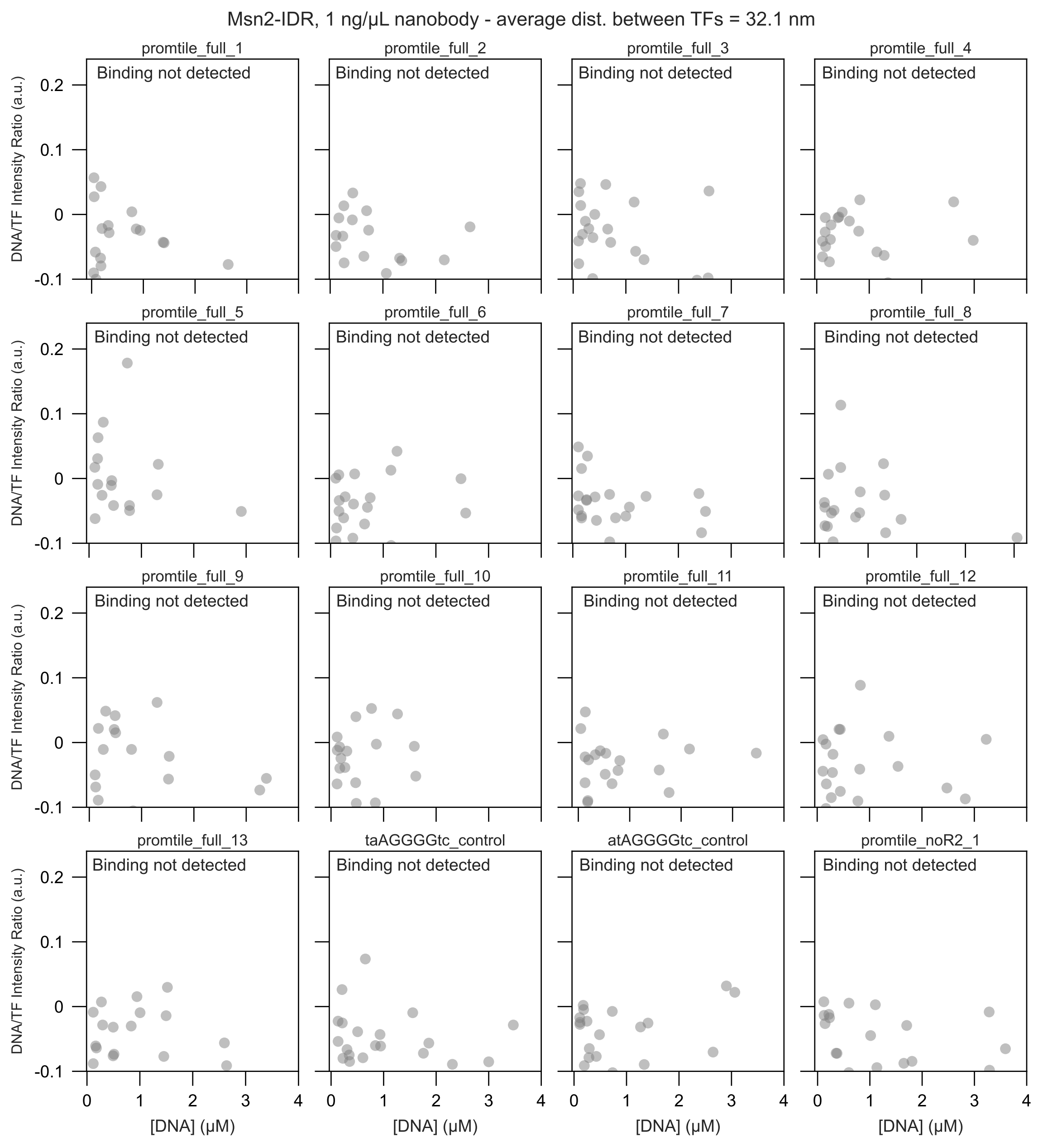
Msn2-IDR, 1 ng/μL nanobody - average dist. between TFs = 32.1 nm
promtile_full_1
promtile_full_2
promtile_full_3
promtile_full_4
promtile_full_5
promtile_full_6
promtile_full_7
promtile_full_8
promtile_full_9
promtile_full_10
promtile_full_11
promtile_full_12
promtile_full_13
taAGGGGtc_control
atAGGGGtc_control
promtile_noR2_1
Binding not detected
DNA/TF Intensity Ratio (a.u.)
[DNA] (μM)
0.2
0.1
0
-0.1
0
1
2
3
4
