## Supplementary File 4 for "Intrinsically disordered regions within transcription factors impact DNA binding through both autoinhibition and independent DNA binding"

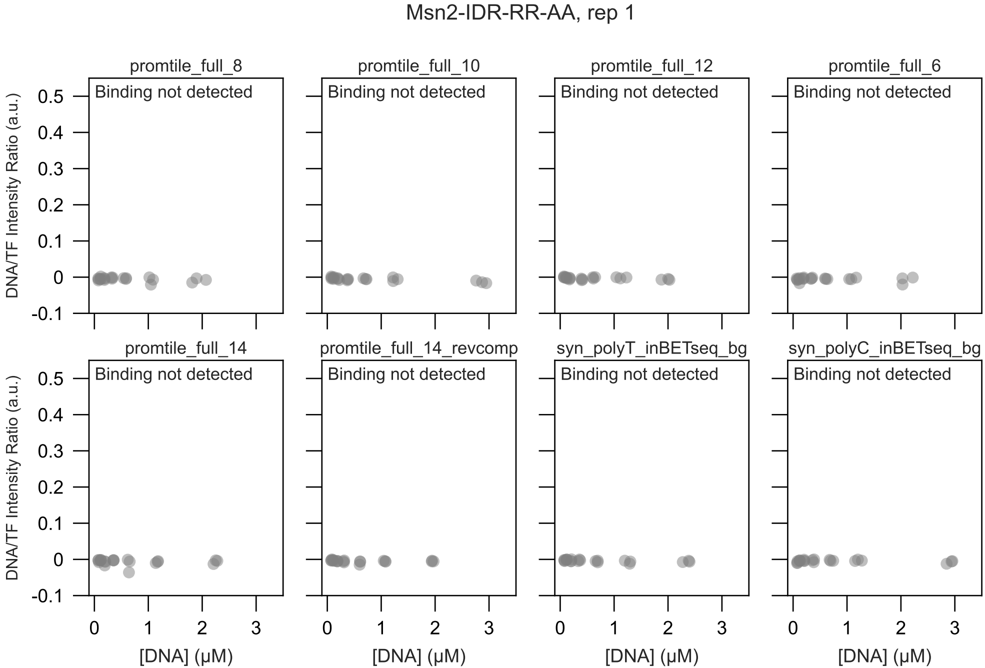
Msn2-IDR-RR-AA, rep 1
promtile_full_8
promtile_full_10
promtile_full_12
promtile_full_6
promtile_full_14
promtile_full_14_revcomp
syn_polyT_inBETseq_bg
syn_polyC_inBETseq_bg
Binding not detected
DNA/TF Intensity Ratio (a.u.)
0.5
0.4
0.3
0.2
0.1
0
-0.1
0
1
2
3
[DNA] (μM)

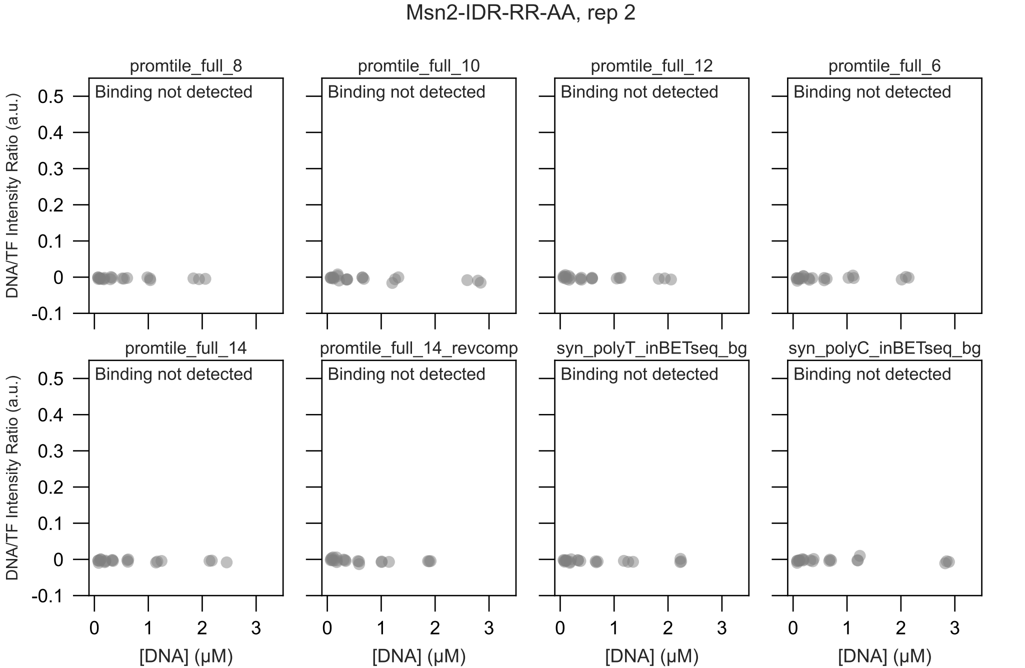

Msn2-IDR-RR-AA, rep 2
promtile_full_8
promtile_full_10
promtile_full_12
promtile_full_6
promtile_full_14
promtile_full_14_revcomp
syn_polyT_inBETseq_bg
syn_polyC_inBETseq_bg
Binding not detected
DNA/TF Intensity Ratio (a.u.)
[DNA] (μM)
0.5
0.4
0.3
0.2
0.1
0
-0.1
0
1
2
3

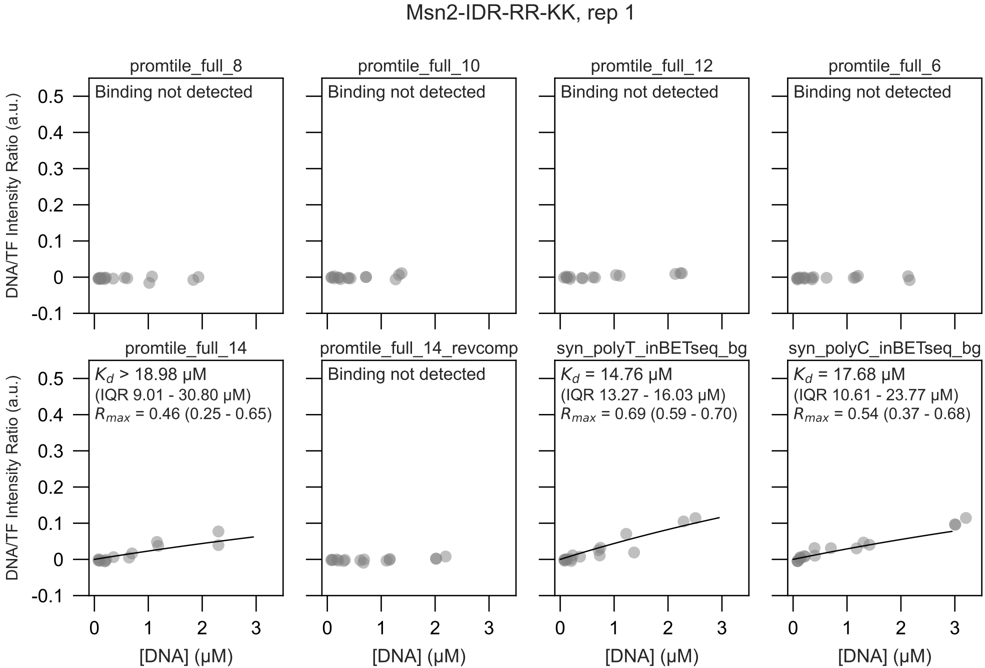
Msn2-IDR-RR-KK, rep 1
promtile_full_8
Binding not detected
promtile_full_10
Binding not detected
promtile_full_12
Binding not detected
promtile_full_6
Binding not detected
promtile_full_14
$K_d$ > 18.98 μM
(IQR 9.01 - 30.80 μM)
$R_{max}$ = 0.46 (0.25 - 0.65)
promtile_full_14_revcomp
Binding not detected
syn_polyT_inBETseq_bg
$K_d$ = 14.76 μM
(IQR 13.27 - 16.03 μM)
$R_{max}$ = 0.69 (0.59 - 0.70)
syn_polyC_inBETseq_bg
$K_d$ = 17.68 μM
(IQR 10.61 - 23.77 μM)
$R_{max}$ = 0.54 (0.37 - 0.68)
DNA/TF Intensity Ratio (a.u.)
[DNA] (μM)

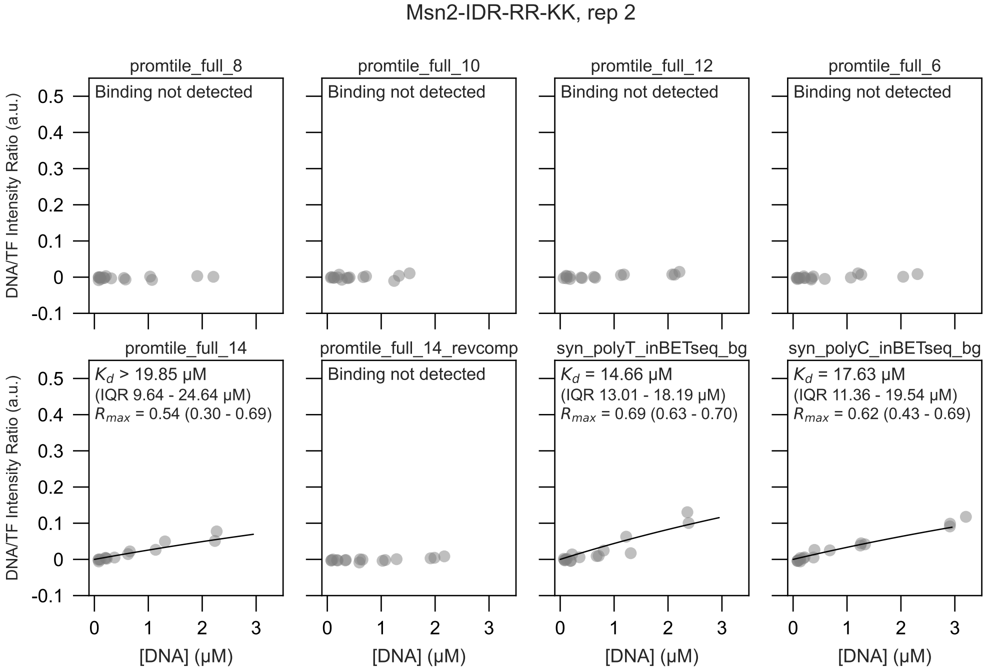

Msn2-IDR-RR-KK, rep 2
promtile_full_8
Binding not detected
promtile_full_10
Binding not detected
promtile_full_12
Binding not detected
promtile_full_6
Binding not detected
promtile_full_14
$K_d$ > 19.85 μM
(IQR 9.64 - 24.64 μM)
$R_{max}$ = 0.54 (0.30 - 0.69)
promtile_full_14_revcomp
Binding not detected
syn_polyT_inBETseq_bg
$K_d$ = 14.66 μM
(IQR 13.01 - 18.19 μM)
$R_{max}$ = 0.69 (0.63 - 0.70)
syn_polyC_inBETseq_bg
$K_d$ = 17.63 μM
(IQR 11.36 - 19.54 μM)
$R_{max}$ = 0.62 (0.43 - 0.69)
DNA/TF Intensity Ratio (a.u.)
[DNA] (μM)
0.5
0.4
0.3
0.2
0.1
0
-0.1
0
1
2
3

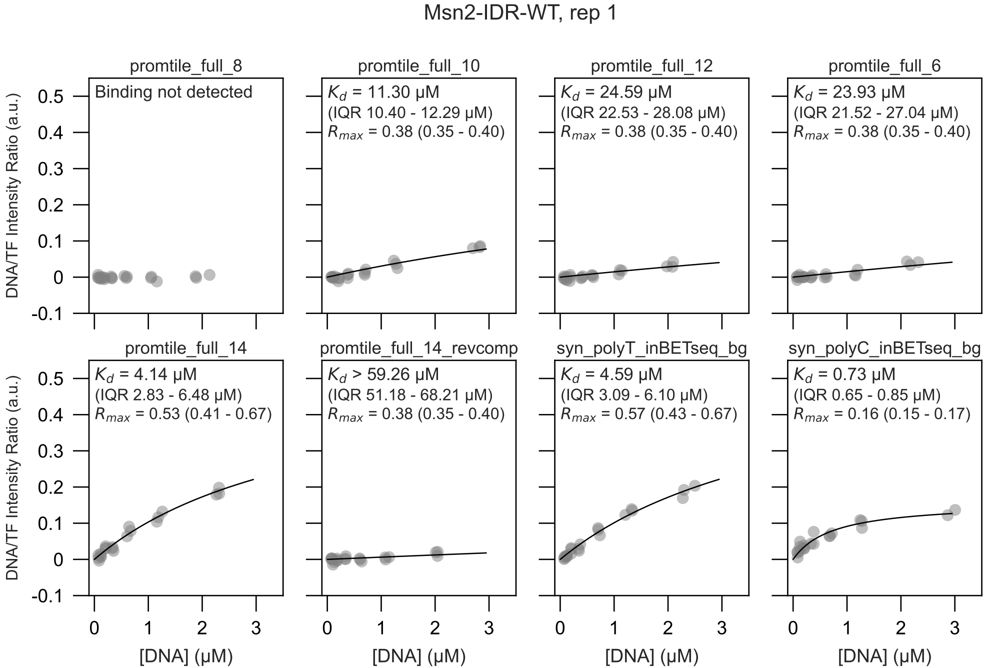
Msn2-IDR-WT, rep 1
promtile_full_8
Binding not detected
promtile_full_10
$K_d$ = 11.30 μM
(IQR 10.40 - 12.29 μM)
$R_{max}$ = 0.38 (0.35 - 0.40)
promtile_full_12
$K_d$ = 24.59 μM
(IQR 22.53 - 28.08 μM)
$R_{max}$ = 0.38 (0.35 - 0.40)
promtile_full_6
$K_d$ = 23.93 μM
(IQR 21.52 - 27.04 μM)
$R_{max}$ = 0.38 (0.35 - 0.40)
promtile_full_14
$K_d$ = 4.14 μM
(IQR 2.83 - 6.48 μM)
$R_{max}$ = 0.53 (0.41 - 0.67)
promtile_full_14_revcomp
$K_d$ > 59.26 μM
(IQR 51.18 - 68.21 μM)
$R_{max}$ = 0.38 (0.35 - 0.40)
syn_polyT_inBETseq_bg
$K_d$ = 4.59 μM
(IQR 3.09 - 6.10 μM)
$R_{max}$ = 0.57 (0.43 - 0.67)
syn_polyC_inBETseq_bg
$K_d$ = 0.73 μM
(IQR 0.65 - 0.85 μM)
$R_{max}$ = 0.16 (0.15 - 0.17)
DNA/TF Intensity Ratio (a.u.)
[DNA] (μM)
0.5
0.4
0.3
0.2
0.1
0
-0.1
0
1
2
3

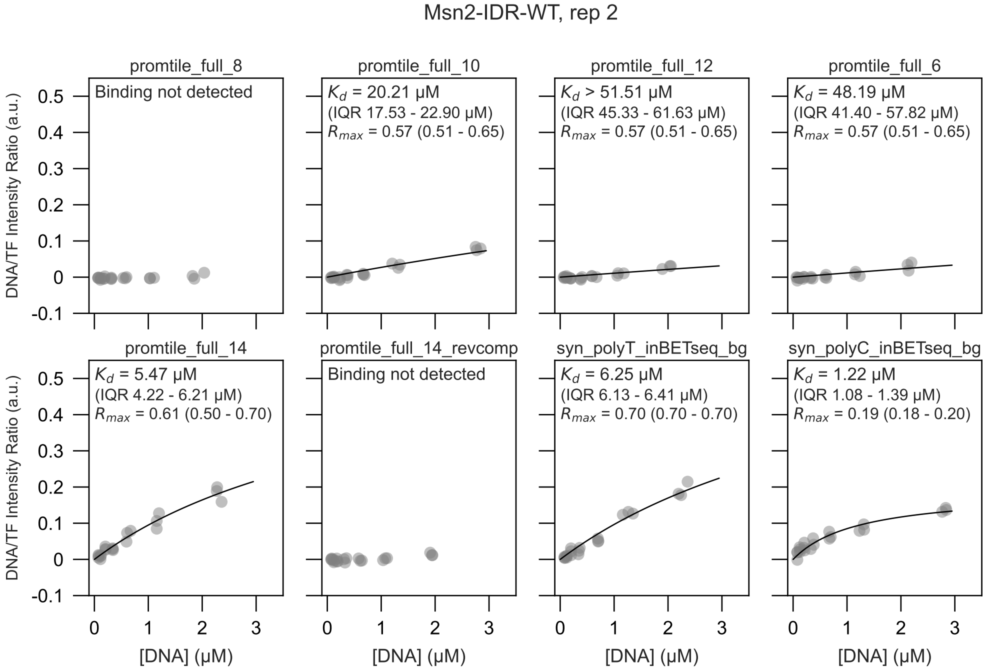

Msn2-IDR-WT, rep 2
promtile_full_8
Binding not detected
promtile_full_10
$K_d$ = 20.21 μM
(IQR 17.53 - 22.90 μM)
$R_{max}$ = 0.57 (0.51 - 0.65)
promtile_full_12
$K_d$ > 51.51 μM
(IQR 45.33 - 61.63 μM)
$R_{max}$ = 0.57 (0.51 - 0.65)
promtile_full_6
$K_d$ = 48.19 μM
(IQR 41.40 - 57.82 μM)
$R_{max}$ = 0.57 (0.51 - 0.65)
promtile_full_14
$K_d$ = 5.47 μM
(IQR 4.22 - 6.21 μM)
$R_{max}$ = 0.61 (0.50 - 0.70)
promtile_full_14_revcomp
Binding not detected
syn_polyT_inBETseq_bg
$K_d$ = 6.25 μM
(IQR 6.13 - 6.41 μM)
$R_{max}$ = 0.70 (0.70 - 0.70)
syn_polyC_inBETseq_bg
$K_d$ = 1.22 μM
(IQR 1.08 - 1.39 μM)
$R_{max}$ = 0.19 (0.18 - 0.20)
DNA/TF Intensity Ratio (a.u.)
[DNA] (μM)
0.5
0.4
0.3
0.2
0.1
0
-0.1
0
1
2
3

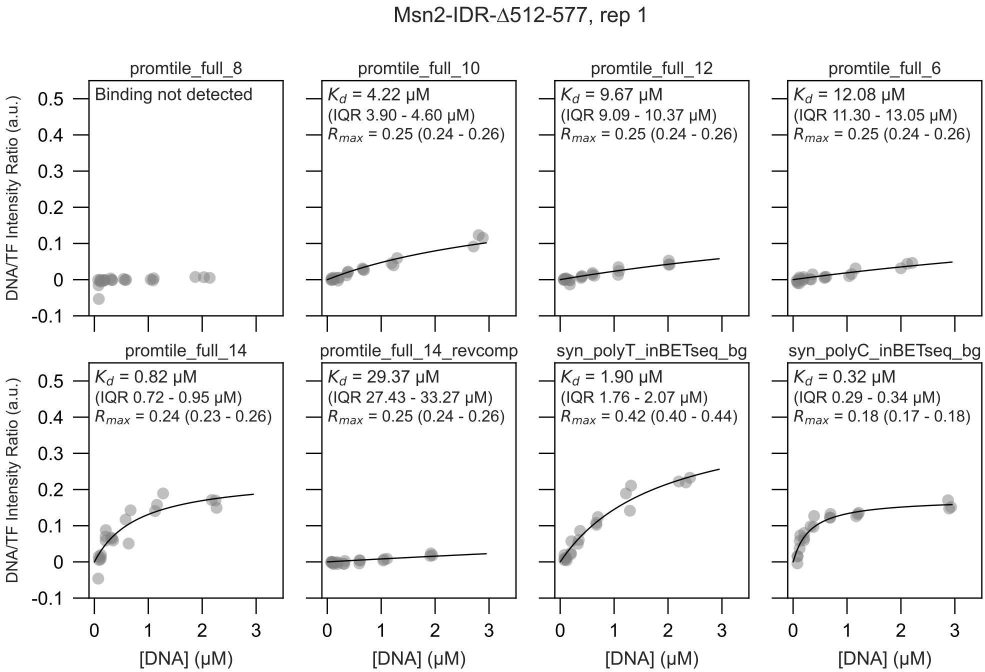
Msn2-IDR-Δ512-577, rep 1
promtile_full_8
Binding not detected
promtile_full_10
$K_d$ = 4.22 μM
(IQR 3.90 - 4.60 μM)
$R_{max}$ = 0.25 (0.24 - 0.26)
promtile_full_12
$K_d$ = 9.67 μM
(IQR 9.09 - 10.37 μM)
$R_{max}$ = 0.25 (0.24 - 0.26)
promtile_full_6
$K_d$ = 12.08 μM
(IQR 11.30 - 13.05 μM)
$R_{max}$ = 0.25 (0.24 - 0.26)
promtile_full_14
$K_d$ = 0.82 μM
(IQR 0.72 - 0.95 μM)
$R_{max}$ = 0.24 (0.23 - 0.26)
promtile_full_14_revcomp
$K_d$ = 29.37 μM
(IQR 27.43 - 33.27 μM)
$R_{max}$ = 0.25 (0.24 - 0.26)
syn_polyT_inBETseq_bg
$K_d$ = 1.90 μM
(IQR 1.76 - 2.07 μM)
$R_{max}$ = 0.42 (0.40 - 0.44)
syn_polyC_inBETseq_bg
$K_d$ = 0.32 μM
(IQR 0.29 - 0.34 μM)
$R_{max}$ = 0.18 (0.17 - 0.18)
DNA/TF Intensity Ratio (a.u.)
[DNA] (μM)
0.5
0.4
0.3
0.2
0.1
0
-0.1
0
1
2
3

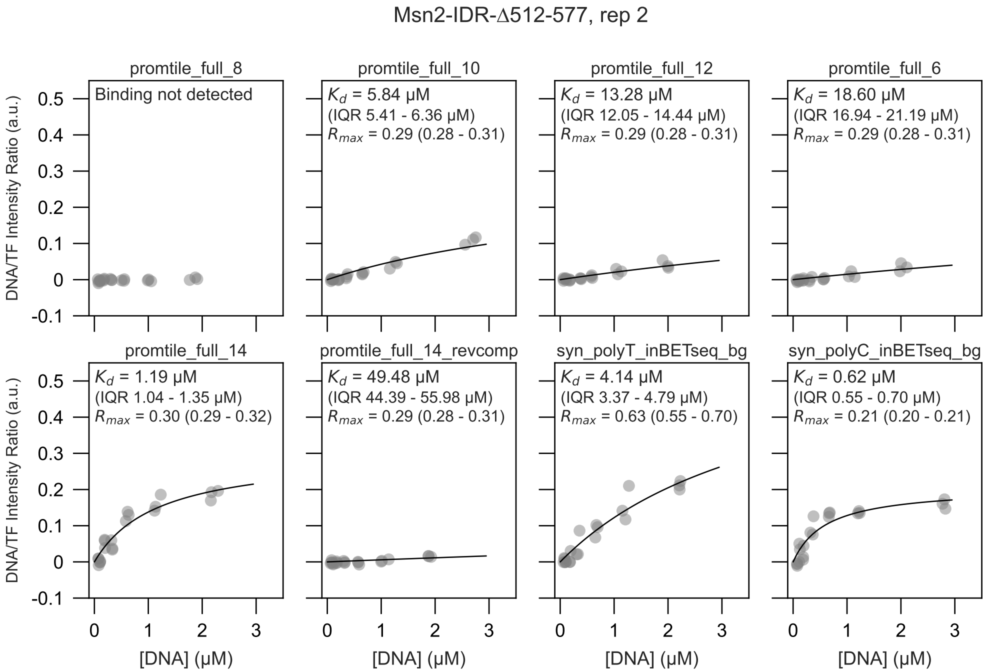

Msn2-IDR-Δ512-577, rep 2
promtile_full_8
Binding not detected
promtile_full_10
$K_d$ = 5.84 μM
(IQR 5.41 - 6.36 μM)
$R_{max}$ = 0.29 (0.28 - 0.31)
promtile_full_12
$K_d$ = 13.28 μM
(IQR 12.05 - 14.44 μM)
$R_{max}$ = 0.29 (0.28 - 0.31)
promtile_full_6
$K_d$ = 18.60 μM
(IQR 16.94 - 21.19 μM)
$R_{max}$ = 0.29 (0.28 - 0.31)
promtile_full_14
$K_d$ = 1.19 μM
(IQR 1.04 - 1.35 μM)
$R_{max}$ = 0.30 (0.29 - 0.32)
promtile_full_14_revcomp
$K_d$ = 49.48 μM
(IQR 44.39 - 55.98 μM)
$R_{max}$ = 0.29 (0.28 - 0.31)
syn_polyT_inBETseq_bg
$K_d$ = 4.14 μM
(IQR 3.37 - 4.79 μM)
$R_{max}$ = 0.63 (0.55 - 0.70)
syn_polyC_inBETseq_bg
$K_d$ = 0.62 μM
(IQR 0.55 - 0.70 μM)
$R_{max}$ = 0.21 (0.20 - 0.21)
DNA/TF Intensity Ratio (a.u.)
[DNA] (μM)

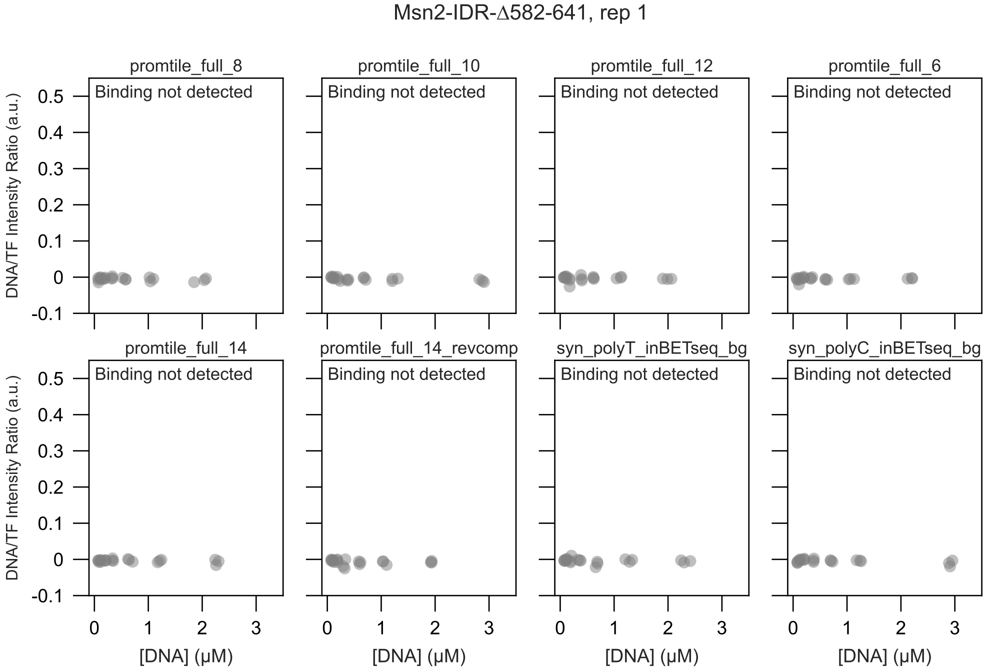

Msn2-IDR-Δ582-641, rep 1
promtile_full_8
promtile_full_10
promtile_full_12
promtile_full_6
promtile_full_14
promtile_full_14_revcomp
syn_polyT_inBETseq_bg
syn_polyC_inBETseq_bg
Binding not detected
DNA/TF Intensity Ratio (a.u.)
[DNA] (μM)
0.5
0.4
0.3
0.2
0.1
0
-0.1
0
1
2
3

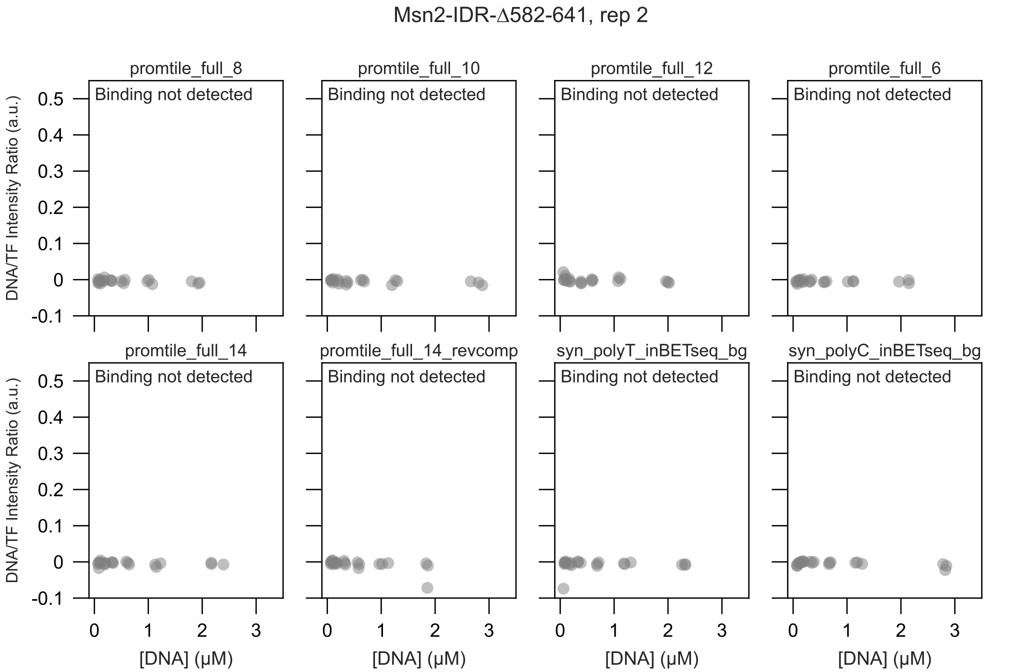

Msn2-IDR-Δ582-641, rep 2
promtile_full_8
promtile_full_10
promtile_full_12
promtile_full_6
promtile_full_14
promtile_full_14_revcomp
syn_polyT_inBETseq_bg
syn_polyC_inBETseq_bg
Binding not detected
DNA/TF Intensity Ratio (a.u.)
0.5
0.4
0.3
0.2
0.1
0
-0.1
0
1
2
3
[DNA] (μM)

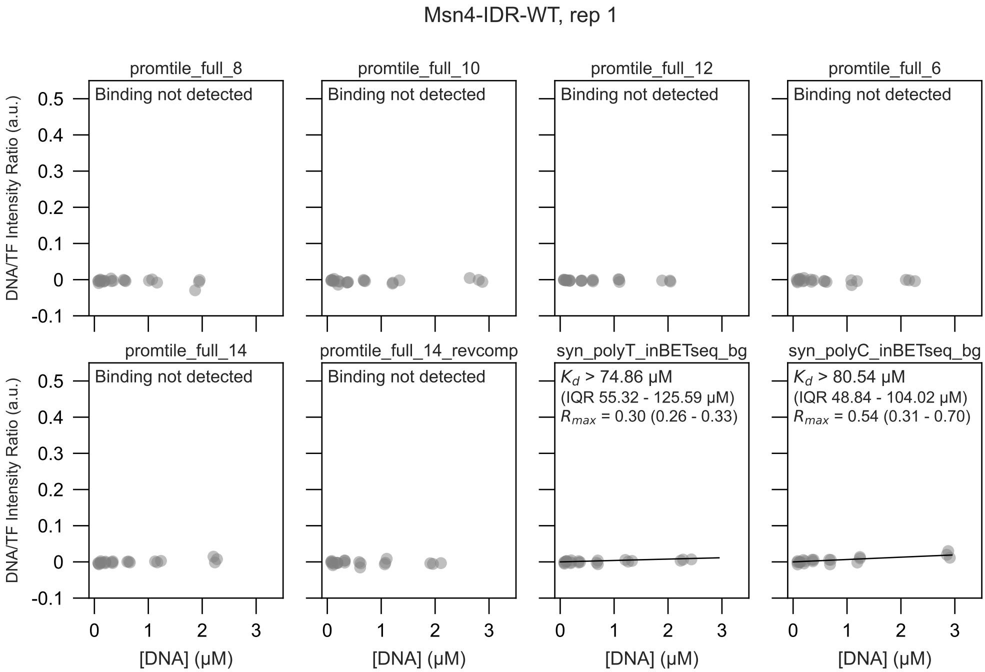

Msn4-IDR-WT, rep 1
promtile_full_8
Binding not detected
promtile_full_10
Binding not detected
promtile_full_12
Binding not detected
promtile_full_6
Binding not detected
promtile_full_14
Binding not detected
promtile_full_14_revcomp
Binding not detected
syn_polyT_inBETseq_bg
$K_d$ > 74.86 µM
(IQR 55.32 - 125.59 µM)
$R_{max}$ = 0.30 (0.26 - 0.33)
syn_polyC_inBETseq_bg
$K_d$ > 80.54 µM
(IQR 48.84 - 104.02 µM)
$R_{max}$ = 0.54 (0.31 - 0.70)
DNA/TF Intensity Ratio (a.u.)
0.5
0.4
0.3
0.2
0.1
0
-0.1
0
1
2
3
[DNA] (µM)

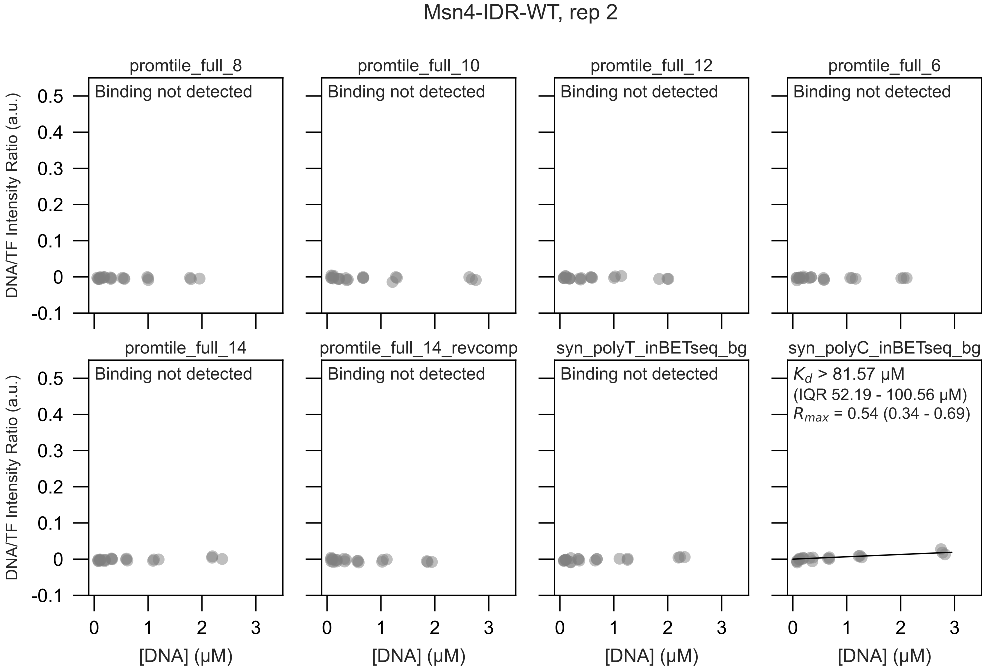

Msn4-IDR-WT, rep 2
promtile_full_8
Binding not detected
promtile_full_10
Binding not detected
promtile_full_12
Binding not detected
promtile_full_6
Binding not detected
promtile_full_14
Binding not detected
promtile_full_14_revcomp
Binding not detected
syn_polyT_inBETseq_bg
Binding not detected
syn_polyC_inBETseq_bg
$K_d$ > 81.57 μM
(IQR 52.19 - 100.56 μM)
$R_{max}$ = 0.54 (0.34 - 0.69)
DNA/TF Intensity Ratio (a.u.)
0.5
0.4
0.3
0.2
0.1
0
-0.1
0
1
2
3
[DNA] (μM)

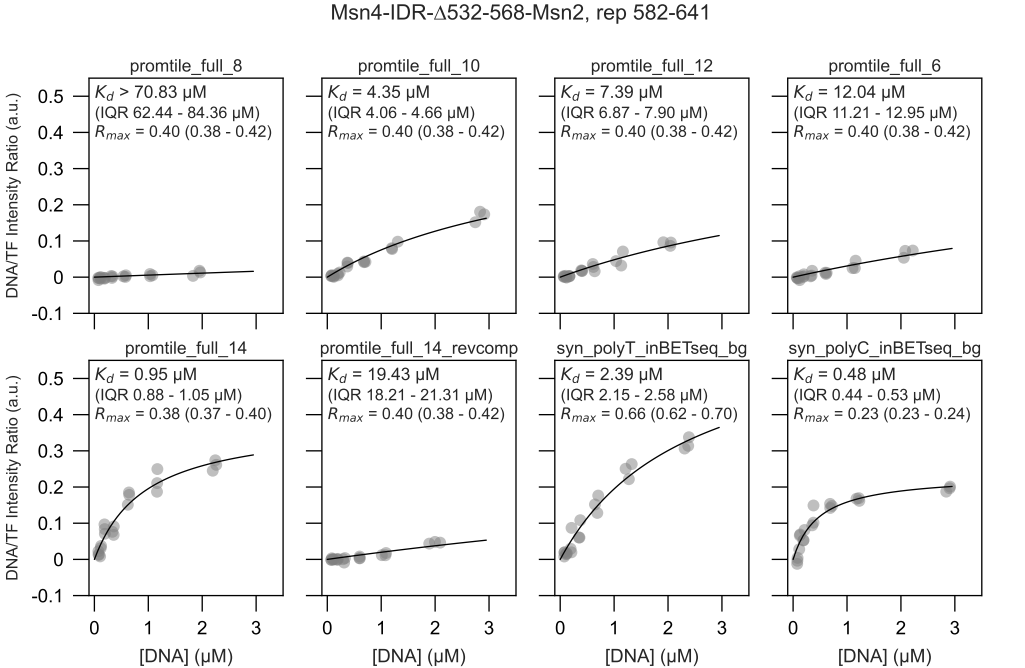
Msn4-IDR-Δ532-568-Msn2, rep 582-641
promtile_full_8
$K_d$ > 70.83 μM
(IQR 62.44 - 84.36 μM)
$R_{max}$ = 0.40 (0.38 - 0.42)
promtile_full_10
$K_d$ = 4.35 μM
(IQR 4.06 - 4.66 μM)
$R_{max}$ = 0.40 (0.38 - 0.42)
promtile_full_12
$K_d$ = 7.39 μM
(IQR 6.87 - 7.90 μM)
$R_{max}$ = 0.40 (0.38 - 0.42)
promtile_full_6
$K_d$ = 12.04 μM
(IQR 11.21 - 12.95 μM)
$R_{max}$ = 0.40 (0.38 - 0.42)
promtile_full_14
$K_d$ = 0.95 μM
(IQR 0.88 - 1.05 μM)
$R_{max}$ = 0.38 (0.37 - 0.40)
promtile_full_14_revcomp
$K_d$ = 19.43 μM
(IQR 18.21 - 21.31 μM)
$R_{max}$ = 0.40 (0.38 - 0.42)
syn_polyT_inBETseq_bg
$K_d$ = 2.39 μM
(IQR 2.15 - 2.58 μM)
$R_{max}$ = 0.66 (0.62 - 0.70)
syn_polyC_inBETseq_bg
$K_d$ = 0.48 μM
(IQR 0.44 - 0.53 μM)
$R_{max}$ = 0.23 (0.23 - 0.24)
DNA/TF Intensity Ratio (a.u.)
[DNA] (μM)
0.5
0.4
0.3
0.2
0.1
0
-0.1
0
1
2
3

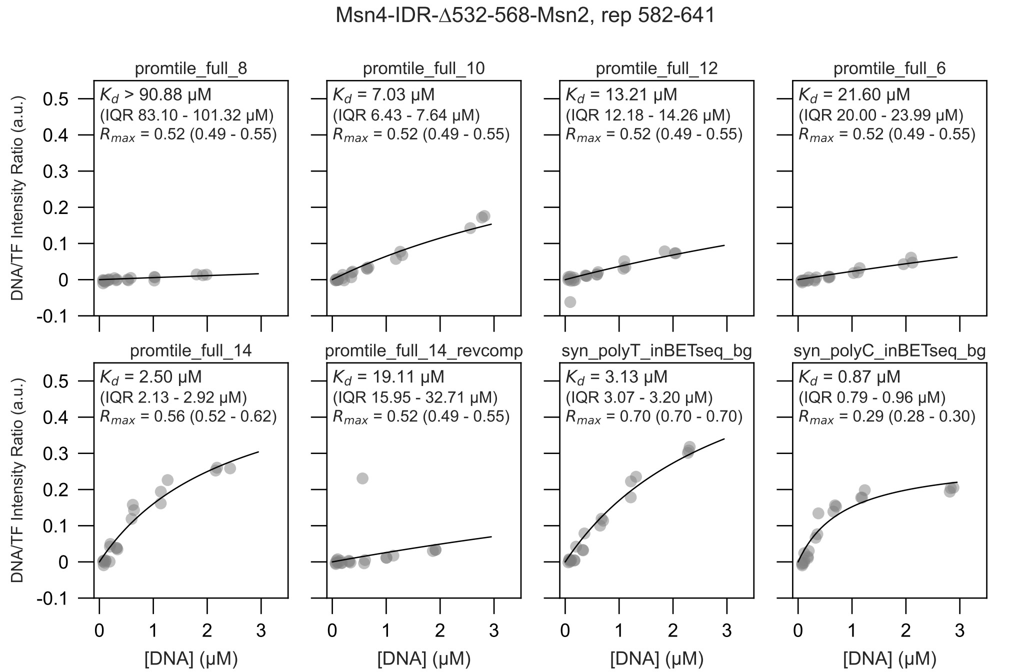

Msn4-IDR-Δ532-568-Msn2, rep 582-641
promtile_full_8
$K_d$ > 90.88 μM
(IQR 83.10 - 101.32 μM)
$R_{max}$ = 0.52 (0.49 - 0.55)
promtile_full_10
$K_d$ = 7.03 μM
(IQR 6.43 - 7.64 μM)
$R_{max}$ = 0.52 (0.49 - 0.55)
promtile_full_12
$K_d$ = 13.21 μM
(IQR 12.18 - 14.26 μM)
$R_{max}$ = 0.52 (0.49 - 0.55)
promtile_full_6
$K_d$ = 21.60 μM
(IQR 20.00 - 23.99 μM)
$R_{max}$ = 0.52 (0.49 - 0.55)
promtile_full_14
$K_d$ = 2.50 μM
(IQR 2.13 - 2.92 μM)
$R_{max}$ = 0.56 (0.52 - 0.62)
promtile_full_14_revcomp
$K_d$ = 19.11 μM
(IQR 15.95 - 32.71 μM)
$R_{max}$ = 0.52 (0.49 - 0.55)
syn_polyT_inBETseq_bg
$K_d$ = 3.13 μM
(IQR 3.07 - 3.20 μM)
$R_{max}$ = 0.70 (0.70 - 0.70)
syn_polyC_inBETseq_bg
$K_d$ = 0.87 μM
(IQR 0.79 - 0.96 μM)
$R_{max}$ = 0.29 (0.28 - 0.30)
DNA/TF Intensity Ratio (a.u.)
[DNA] (μM)

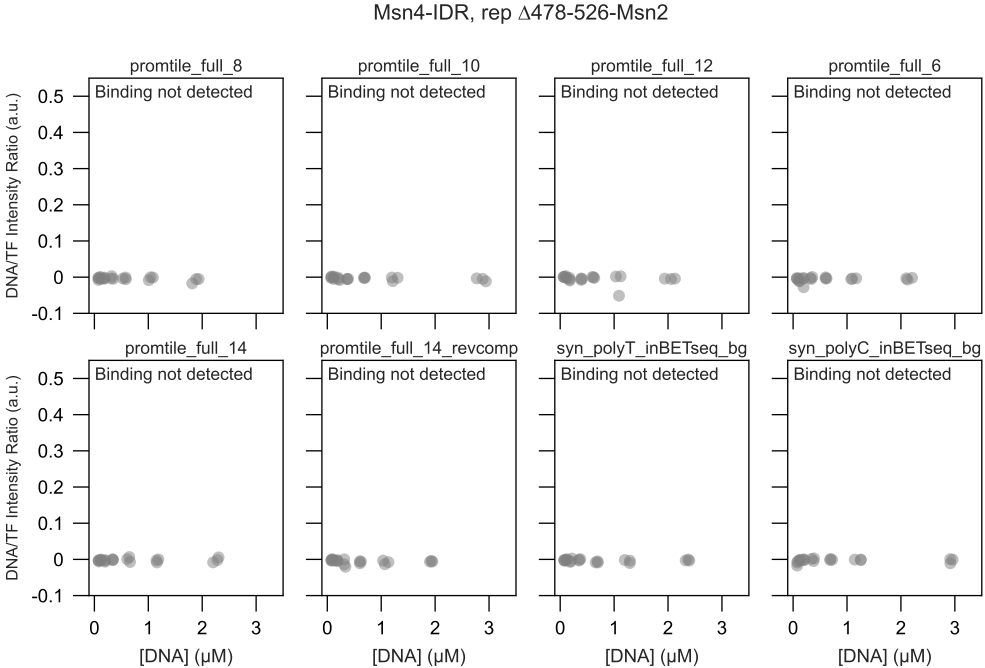
Msn4-IDR, rep Δ478-526-Msn2
promtile_full_8
promtile_full_10
promtile_full_12
promtile_full_6
promtile_full_14
promtile_full_14_revcomp
syn_polyT_inBETseq_bg
syn_polyC_inBETseq_bg
Binding not detected
DNA/TF Intensity Ratio (a.u.)
[DNA] (μM)
0.5
0.4
0.3
0.2
0.1
0
-0.1
0
1
2
3

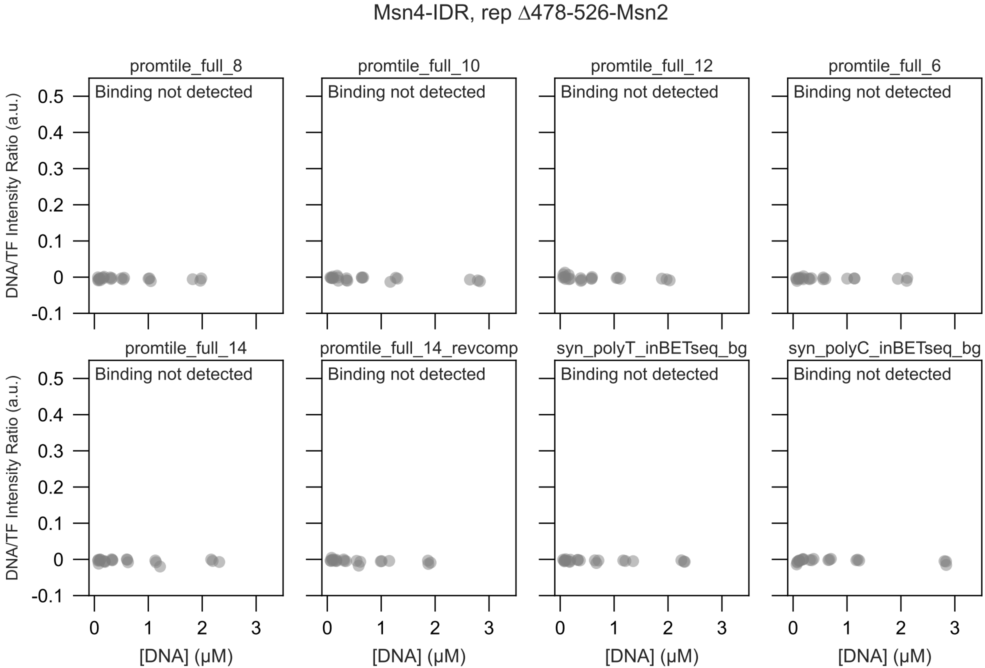
Msn4-IDR, rep Δ478-526-Msn2
promtile_full_8
promtile_full_10
promtile_full_12
promtile_full_6
promtile_full_14
promtile_full_14_revcomp
syn_polyT_inBETseq_bg
syn_polyC_inBETseq_bg
Binding not detected
DNA/TF Intensity Ratio (a.u.)
[DNA] (μM)
0.5
0.4
0.3
0.2
0.1
0
-0.1
0
1
2
3
