## Supplementary Note 1 for "Intrinsically disordered regions within transcription factors impact DNA binding through both autoinhibition and independent DNA binding"

### Derivation of the effects of autoinhibition and avidity on observed equilibrium binding constants

November 19, 2026

#### 1 Binding of TF to DNA with 2 binding sites, with no avidity

Here we consider how the observed $K_d$ is affected by the presence 2 identical binding sites on a single DNA molecule when binding to a TF in a setup with sparsely immobilized TF molecules, such that two TFs cannot bind both binding sites of a single DNA molecule simultaneously.

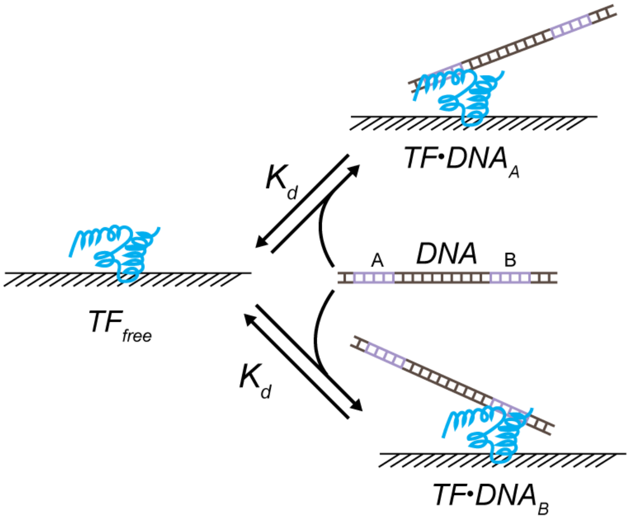


Figure 1: Model of binding of a single TF to DNA with two binding sites

To derive the effects of two DNA binding sites on our observed $K_d$ we use the fact that the observed $K_d$ is the concentration of DNA at which fractional binding, $\theta = \frac{[TF_{bound}]}{[TF_{total}]} = 0.5$. Because the two bound states are equivalent, we can define $[TF_{total}]$ and $[TF_{bound}]$ as follows:

$$[TF_{total}] = [TF_{free}] + [TF \cdot DNA_A] + [TF \cdot DNA_B] = [TF_{free}] + 2[TF \cdot DNA_A]$$

and

$$[TF_{bound}] = [TF \cdot DNA_A] + [TF \cdot DNA_B] = 2[TF \cdot DNA_A]$$

Since $K_d = \frac{[TF_{free}][DNA]}{[TF \cdot DNA_A]}$, we can solve for $[TF \cdot DNA_A]$ and find that $[TF \cdot DNA_A] = \frac{[DNA][TF_{free}]}{K_d}$. We can then plug this expression into our equations for $[TF_{total}]$ and $[TF_{bound}]$ and divide by $TF_{free}$ to get the following equations:

$$\frac{[TF_{total}]}{[TF_{free}]} = 1 + 2\frac{[DNA]}{K_d}$$

and

$$\frac{[TF_{bound}]}{[TF_{free}]} = 2\frac{[DNA]}{K_d}$$

We can then plug these values of $[TF_{total}]$ and $[TF_{bound}]$ into equation for $\theta$, set it equal to 0.5:

$$\theta = 0.5 = \frac{2\frac{[DNA]}{K_d}}{1+2\frac{[DNA]}{K_d}}$$

We then solve for $[DNA]$ to get $K_{d,obs}$:

$$[DNA] = K_{d,obs} = \frac{K_d}{2}$$

Thus, having two identical binding sites on a molecule of DNA that cannot be simultaneously bound by two immobilized TFs simply decreases the observed $K_d$ by a factor of 2.

### 2 Avid binding with no autoinhibition

In our experimental setup, we immobilize TFs on the device surface. When measuring binding to the promoter tiles, we are exposing these immobilized TFs to DNA molecules with multiple potential binding sites. Thus, we must consider the possibility of simultaneous binding of multiple adjacent binding sites on the same DNA molecule by adjacent TFs - in other words, avidity.

#### 2.1 Avidity with two binding sites

We begin by considering the simplest case, in which we have two adjacent, immobilized TFs and a DNA molecule with two binding sites of equal affinity.

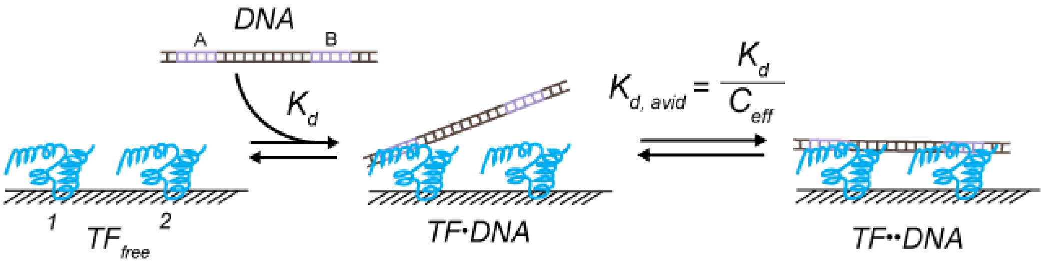


Figure 2: Model of avid binding with 2 sites

In this simplified model, we make two assumptions. (1) Both the TFs and the two binding sites on the DNA are identical, rendering binding of $TF_1$ to binding site A degenerate with binding of $TF_2$ to binding site B. (2) Binding orientations that ostensibly cannot lead to the avid-bound state (e.g., binding of $TF_1$ to binding site B) in reality are geometries that can lead to avid binding since, in our experiment, each TF is surrounded by other TFs and has freedom to rotate on the surface. With these two assumptions, we can operate under the assumption that all single-bound states are equivalent and degenerate. Therefore, we consider three states that the TFs can occupy: the unbound state ($TF_{free}$, which is the sum of unbound $TF_1$ and $TF_2$), the single-bound state ($TF \cdot DNA$), and the avid, double-bound state ($TF \cdot \cdot DNA$, in which $TF_1$ and $TF_2$ are simultaneously bound).
We can model the first binding step as a normal binding interaction where:

$$K_d = \frac{[TF_{free}][DNA]}{[TF{\cdot}DNA]}$$

The subsequent binding can be considered as an intramolecular binding reaction, similar to how we think of autoinhibition, in which the equilibrium constant is an empirical constant (as in a $K_{auto}$) independent of concentration. However, based on first principles, we can break this constant into component parts, namely (1) an equilibrium association constant ($K_a$) between the second DNA binding site and the TF and (2) the effective concentration of the second binding sites relative to each other ($C_{eff}$), where

$K = K_a \cdot C_{eff}$[1]. Written as an equilibrium dissociation constant:

$$K_{d,avid} = \frac{[TF \cdot DNA]}{[TF \cdot\cdot DNA]} = \frac{K_d}{C_{eff}}$$

To derive how these two binding steps affect the observed $K_d$(or $K_{d,obs}$), we consider the equation describing sum of the states the TFs can populate:

$$TF_{total} = [TF_{free}] + 2[TF \cdot DNA] + [TF \cdot \cdot DNA]$$

Given our defined equilibrium constants, we can solve for $[TF \cdot DNA]$ and $[TF \cdot \cdot DNA]$ in terms of $K_d$, $[DNA]$, $[TF_{free}]$, and $C_{eff}$, finding that $[TF \cdot DNA] = \frac{[TF_{free}][DNA]}{K_d}$ and $[TF \cdot \cdot DNA] = \frac{[TF_{free}][DNA]C_{eff}}{(K_d)^2}$. We then substitute these into the equation above, and normalize all terms to $[TF_{free}]$:

$$\frac{TF_{total}}{[TF_{free}]} = 1 + 2\frac{[DNA]}{K_d} + \frac{[DNA]C_{eff}}{(K_d)^2}$$

We then define $\theta$ as the fraction of bound TF relative to the total TF:

$$\theta = \frac{2[TF \cdot DNA] + [TF \cdot\cdot DNA]}{TF_{total}}$$

We can again normalize the numerator and denominator of the above equation to $[TF_{free}]$, arriving at:

$$\theta = \frac{2\frac{[DNA]}{K_d} + \frac{[DNA]C_{eff}}{(K_d)^2}}{1 + 2\frac{[DNA]}{K_d} + \frac{[DNA]C_{eff}}{(K_d)^2}}$$

Our $K_{d,obs}$ can be defined as the concentration of DNA at which $\theta = 0.5$, so we can set our above equation for $\theta$ to 0.5, and then solve for $[DNA]$, arriving at:

$$K_{d,obs} = \frac{(K_d)^2}{2K_d + C_{eff}}$$

Under our experimental conditions, we estimate $C_{eff}$ to be in the range of hundreds of µM - mM (see Supplemental Methods), and therefore can often make the assumption that $C_{eff} >> 2K_d$. Under this assumption, $2K_d + C_{eff} \rightarrow C_{eff}$, and thus:

$$\mathbf{K_{d,obs} = \frac{(K_d)^2}{C_{eff}}}$$

#### 2.2 Avidity with n binding sites

To describe the case where we have an arbitrary number of additional sites on the DNA and an arbitrary number of adjacent, immobilized TFs, we make the following assumptions: (1) The effective concentrations of the DNA binding site to their respective TFs are equal, and (2), as in section 1.1, any binding that would not lead to all n sites being bound in our cartoon models (for example, $TF_2$ binding to DNA site 1 in Figure 3) can be assumed to create a situation in which binding to all n sites is indeed possible with additional adjacent TFs present on the device surface. Thus, any "off-path" binding can be considered degenerate. In the example of $TF_2$ binding to DNA site 1, there would be another TF molecule adjacent to $TF_n$, such that this $TF_2$ would really be equivalent to $TF_1$ if we shift our frame over by 1 TF. With these assumptions, we can create the following table of states and weights, where the weights represent the concentrations of each state normalized to the concentration of unbound TF.

| # Bound | 0 | 1 | 2 | k | n |
|---|---|---|---|---|---|
| Weight (norm to unbound) | 1 | x | xy | $xy^{k-1}$ | Where $xy^{n-1}$ |
| Multiplicity | n choose 0 = 1 | n choose 1 = n | n choose 2 | n choose k | n choose n = 1 |
| States | | | | | |

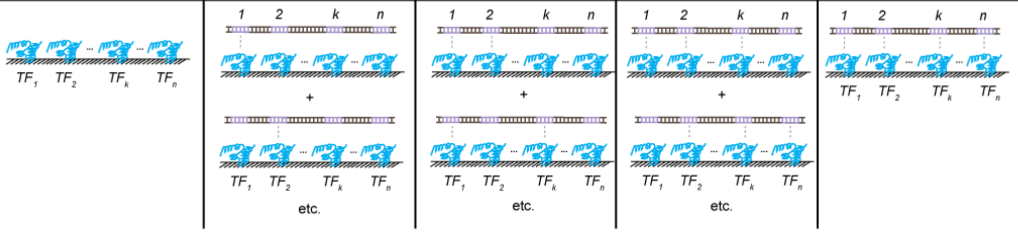


Where $x = \frac{[DNA]}{K_d}$ and $y = \frac{C_{eff}}{K_d}$

Figure 3: States and weights for avid binding with arbitrary sites n

The weights above have been simplified to terms of $x$ and $y$, where (similarly to the setup of the case with avid binding with 2 sites) $x = \frac{[DNA]}{K_d}$ and $y = \frac{C_{eff}}{K_d}$. Intuitively, for each additional bound site after 1, the weight is multiplied by an additional $y$ term, since, using the definition of $K_{d,avid}$ we can express the concentration of each avid bound state with $k$ bound sites as

$$[TF \cdot DNA_{k,bound}] = [TF \cdot DNA_{k-1,bound}]\frac{C_{eff}}{K_d} = [TF \cdot DNA_{k-1,bound}]y$$

If $k = 2$, then $[TF \cdot DNA_{2,bound}] = [TF_{free}]xy$; if $k = 3$, then $[TF \cdot DNA_{3,bound}] = [TF_{free}]xy^2$; and so on.

Describing the multiplicity, or the number of degenerate and equivalent states, for each  of bound DNA sites in Figure 3 is a classical problem of calculating the number of combinations in which one can choose $k$ elements from $n$ total elements, since the order of sites chosen does not matter. For example, we know that there are $n$ ways to bind 1 out of $n$ total TFs (or n choose 1, $C_1^n$) for the $TF \cdot DNA_{1,bound}$ state. Similarly, in the case of the $TF \cdot DNA_{2,bound}$ state, the multiplicity can in turn be described as n choose 2 ($C_2^n$).

Given these weights and multiplicities, we can express the total TF concentration normalized to $TF_{free}$ as follows.

$$\frac{[TF_{total}]}{[TF_{free}]} = 1 + C_1^n x + C_2^n xy + ... + C_k^n xy^{k-1} + C_n^n xy^{n-1} = 1 + \sum_{k=1}^{n} C_k^n xy^{k-1}$$
$$\frac{[TF_{total}]}{[TF_{free}]} = 1 + \frac{x}{y}\sum_{k=1}^{n} C_k^n y^k$$

Using the binomial theorem, which states that $\sum_{k=0}^{n} C_k^n y^k = (1+y)^k$, and the fact that when $k = 0$, $C_0^n y^0 = 1$ we again simplify:

$$\frac{[TF_{total}]}{[TF_{free}]} = 1 + \frac{x}{y}((1+y)^k - 1)$$

We can then solve for $K_{d,obs}$ in a similar fashion as before, by setting the fraction bound, $\theta = \frac{TF_{bound}}{TF_{total}} = \frac{\frac{TF_{bound}}{[TF_{free}]}}{\frac{TF_{total}}{[TF_{free}]}}$, to 0.5, and solving for $[DNA]$ in terms of $K_d$ and $C_{eff}$. We begin by substituting our weights from before into this equation:

$$\frac{1}{2} = \frac{\frac{x}{y}((1+y)^k - 1)}{1 + \frac{x}{y}((1+y)^k - 1)}$$

We can then solve for $x$, which contains our $[DNA]$ term:

$$x = \frac{y}{(1+y)^n - 1}$$

Substituting our values for x and y from above into this equation and solving for $[DNA]$ we arrive at the following.

$$[DNA] = K_{d,obs} = \frac{C_{eff}}{(1+\frac{Ceff}{K_d})^n - 1}$$

If we make the same assumption as before that $C_{eff} >> K_d$, then $(1 + \frac{Ceff}{K_d})^n - 1 \approx (\frac{Ceff}{K_d})^n$. Substituting this into the equation above and simplifying gives our final result that, for avid binding to n binding sites under conditions in which $C_{eff} >> K_d$:

$$\mathbf{K_{d,obs}} = \frac{\mathbf{(K_d)^n}}{\mathbf{(C_{eff})^{n-1}}}$$

### 3 Derivation for Binding with Autoinhibition

Below we derive equations that describe the effects of TF autoinhibition on observed binding to DNA with a single binding site.

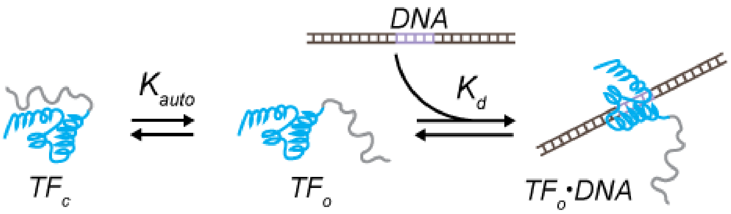


Figure 4: Model of simple autoinhibition and single-site binding

For a simple 3 state autoinhibition model, we define the three states as (1) $TF_c$, (2) $TF_o$, and (3) $TF \cdot DNA$, where $TF_c$ signifies the closed state of the TF and $TF_o$ signifies the open, binding-competent state. The equilibrium constants can be defined as follows:

$$K_{auto} = \frac{[TF_c]}{[TF_o]}$$

$$K_d = \frac{[TF_o][DNA]}{[TF \cdot DNA]}$$

We define $f_{active}$ as the fraction of uninhibited, unbound TF at equilibrium:

$$f_{active} = \frac{[TF_o]}{[TF_o]+[TF_c]}$$

If we multiply this by $\frac{\frac{1}{[TF_o]}}{\frac{1}{[TF_o]}}$ and substitute $K_{auto}$ for $\frac{[TF_c]}{[TF_o]}$, we arrive at:

$$f_{active} = \frac{1}{1+K_{auto}}$$

While $K_d = \frac{[TF_o][DNA]}{[TF \cdot DNA]}$, we can only observe the total DNA-bound TF and non-DNA-bound TF in an experiment. Therefore, within an experiment $K_{d,obs} = \frac{([TF_o]+[TF_c])[DNA]}{[TF \cdot DNA]}$

By definition, $[TF_o] + [TF_c] = \frac{[TF_o]}{f_{active}}$. Therefore, $K_{d,obs} = \frac{\frac{[TF_o]}{f_{active}}[DNA]}{[TF \cdot DNA]}$

Since $\frac{1}{f_{active}} = 1 + K_{auto}$, we can get our observed equilibrium binding constant in terms of $K_{auto}$:

$$K_{d,obs} = \frac{[TF_o][DNA]}{[TF \cdot DNA]}(1 + K_{auto})$$

Therefore, $\mathbf{K_{d,obs} = K_d(1 + K_{auto})}$

### 4 Derivation for avid binding with autoinhibition

#### 4.1 Avidity with two binding sites and autoinhibition

We have observed the presence of an autoinhibited, binding incompetent state in our TFs, so we now consider the case in which we have two adjacent, immobilized TFs that have a closed, autoinhibited state and a DNA molecule with two binding sites of equal affinity.

In this simplified model we make the same assumptions as in section 1.1, with the additional assumption that the autoinhibition of each TF is independent and identical, meaning that the two possible unbound states with one TF in the open state and the other in the closed state ($TF_{c,o}$ and $TF_{o,c}$)

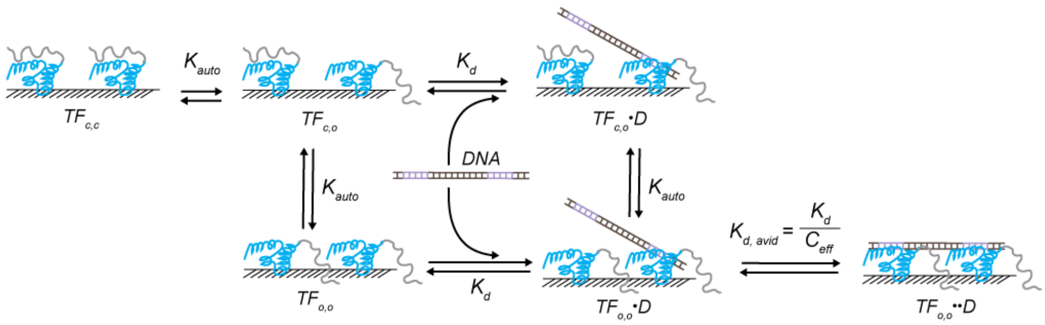


Figure 5: Simplified model of avid binding with 2 sites and autoinhibition

are degenerate. Similarly, the two possible single-DNA-bound states with one TF in the open state and the other in the closed state ($TF_{c,o} \cdot DNA$ and $TF_{o,c} \cdot DNA$) are equivalent and degenerate. Shortening our DNA term down to the symbol $D$, we can enumerate the sum of possible states as follows:

$$TF_{total} = [TF_{c,c}] + 2[TF_{c,o}] + [TF_{o,o}] + 2[TF_{c,o} \cdot D] + 2[TF_{o,o} \cdot D] + [TF_{o,o} \cdot \cdot D]$$

Given the simplifying assumptions above, we can also define our equilibrium constants as follows:

$$K_{auto} = \frac{[TF_{c,c}]}{[TF_{c,o}]} = \frac{[TF_{c,o}]}{[TF_{o,o}]} = \frac{[TF_{c,o} \cdot DNA]}{[TF_{o,o} \cdot DNA]}$$
$$K_d = \frac{[TF_{c,o}][D]}{[TF_{c,o} \cdot D]} = \frac{[TF_{o,o}][D]}{[TF_{o,o} \cdot D]}$$
$$K_{d,avid} = \frac{K_d}{C_{eff}} = \frac{[TF_{o,o} \cdot D]}{[TF_{o,o} \cdot \cdot D]}$$

Again using the fact that $K_{d,obs}$ is equal to the DNA concentration at which half the total TF is bound, we can define the fraction bound, $\theta$, as follows:

$$\theta = \frac{2[TF_{c,o} \cdot DNA] + 2[TF_{o,o} \cdot DNA], + [TF_{,o} \cdot \cdot DNA]}{TF_{total}}$$

We can then solve each term in this equation in terms of $TF_{c,c}$, $K_{auto}$, $K_d$, and $C_{eff}$, multiply the numerator and denominator of this fraction by $\frac{1}{TF_{c,c}}$, and set $\theta$ to 0.5 resulting in the following equation:

$$0.5 = \frac{2\frac{[D]}{K_{auto}K_d} + 2\frac{[D]}{(K_{auto})^2 K_d} + \frac{[D]C_{eff}}{(K_{auto})^2(K_d)^2}}{1 + \frac{2}{K_{auto}} + \frac{1}{(K_{auto})^2} + 2\frac{[D]}{K_{auto}K_d} + 2\frac{[D]}{(K_{auto})^2 K_d} + \frac{[D]C_{eff}}{(K_{auto})^2(K_d)^2}}$$

Solving for $[D]$ in this equation yields the following:

$$K_{d,obs} = [D] = \frac{(K_d)^2(1+K_{auto})^2}{2K_d(1+K_{auto})+C_{eff}}$$

Finally, if we make a similar assumption as in prior derivations that $C_{eff} >> 2K_d(1+K_{auto})$, then the above equation simplifies to the following:

$$\mathbf{K_{d,obs}} = \frac{\mathbf{(K_d)^2(1+K_{auto})^2}}{\mathbf{C_{eff}}}$$

#### 4.2 Avidity with n binding sites and autoinhibition

We again make the same initital assumptions as in section 1.2. In order to derive a generalized formula for the observed avid $K_d$ with n binding sites and autoinhibition we can build on our derivation used in section 1.2, aiming to describe the observed $K_d$ as the DNA concentration at which the fraction bound, $\theta = \frac{[TF_{bound}]}{[TF_{total}]} = 0.5$, in terms of $K_d$, $K_{auto}$, and $C_{eff}$. To begin we enumerate all possible states, their multiplicities, and their weights in terms of $K_d$, $K_{auto}$, $C_{eff}$, and $[TF_{c_n}]$ (the state with all n TFs in closed conformation) in Figure 5:

### of sites bound by DNA with n total sites

| STATES (# of open (uninhibited) TFs) | 0 | 1 | 2 | j | n |
|---|---|---|---|---|---|
| 0 | $TF_{c(n)}$ | X | X | X | X |
| 1 | $TF_{c(n-1), o}$ | $TF_{c(n-1), o}{\cdot}DNA_1$ | X | X | X |
| 2 | $TF_{c(n-2), o(2)}$ | $TF_{c(n-2), o(2)}{\cdot}DNA_1$ | $TF_{c(n-2), o(2)}{\cdot}DNA_2$ | X | X |
| k | $TF_{c(n-k), o(k)}$ | $TF_{c(n-k), o(k)}{\cdot}DNA_1$ | $TF_{c(n-k), o(k)}{\cdot}DNA_2$ | $TF_{c(n-k), o(k)}{\cdot}DNA_j$ | X |
| n | $TF_{o(n)}$ | $TF_{o(n)}{\cdot}DNA_1$ | $TF_{o(n)}{\cdot}DNA_2$ | $TF_{o(n)}{\cdot}DNA_j$ | $TF_{o(n)}{\cdot}DNA_n$ |

| MULTIPLICITY (# of open (uninhibited) TFs) | 0 | 1 | 2 | j | n |
|---|---|---|---|---|---|
| 0 | $C_0^n \bullet C_0^0$ | X | X | X | X |
| 1 | $C_1^n \bullet C_0^1$ | $C_1^n \bullet C_1^1$ | X | X | X |
| 2 | $C_2^n \bullet C_0^2$ | $C_2^n \bullet C_1^2$ | $C_2^n \bullet C_2^2$ | X | X |
| k | $C_k^n \bullet C_0^k$ | $C_k^n \bullet C_1^k$ | $C_k^n \bullet C_2^k$ | $C_k^n \bullet C_j^k$ | X |
| n | $C_n^n \bullet C_0^n$ | $C_n^n \bullet C_1^n$ | $C_n^n \bullet C_2^n$ | $C_n^n \bullet C_j^n$ | $C_n^n \bullet C_n^n$ |

| WEIGHTS (# of open (uninhibited) TFs) | 0 | 1 | 2 | j | n |
|---|---|---|---|---|---|
| 0 | $[TF_{c(n)}]$ | X | X | X | X |
| 1 | $\frac{[TF_{c(n)}]}{K_{auto}}$ | $\frac{[TF_{c(n)}]}{K_{auto}} \frac{[D]}{K_d}$ | X | X | X |
| 2 | $\frac{[TF_{c(n)}]}{K_{auto}{}^2}$ | $\frac{[TF_{c(n)}]}{(K_{auto})^2} \frac{[D]}{K_d}$ | $\frac{[TF_{c(n)}][D]\,C_{eff}}{(K_{auto})^2 (K_d)^2}$ | X | X |
| k | $\frac{[TF_{c(n)}]}{K_{auto}{}^k}$ | $\frac{[TF_{c(n)}][D]}{(K_{auto})^k (K_d)}$ | $\frac{[TF_{c(n)}][D]\,C_{eff}}{(K_{auto})^k (K_d)^2}$ | $\frac{[TF_{c(n)}][D](C_{eff})^{j-1}}{(K_{auto})^k (K_d)^j}$ | X |
| n | $\frac{[TF_{c(n)}]}{K_{auto}{}^n}$ | $\frac{[TF_{c(n)}][D]}{(K_{auto})^n (K_d)}$ | $\frac{[TF_{c(n)}][D]\,C_{eff}}{(K_{auto})^n (K_d)^2}$ | $\frac{[TF_{c(n)}][D](C_{eff})^{j-1}}{(K_{auto})^n (K_d)^j}$ | $\frac{[TF_{c(n)}][D](C_{eff})^{n-1}}{(K_{auto})^n (K_d)^n}$ |

Figure 6: Possible states, multiplicities, and weights when considering $n$ TFs, within each of which autoinhibition is possible, binding to a DNA molecule with $n$ binding sites

To further illustrate why the multiplicity of each state is a product of combinations, we can consider the example of $n = 3$ TFs total, $k = 2$ open TFs, and $j = 1$ DNA site bound in Figure 6. Here, there are $C_2^3 = 3$ possible ways to arrange two open TFs among the three. Within each of those possibilities, there are $C_1^2 = 2$ ways to bind 1 out of 3 possible binding sites on the DNA, resulting in $3 \cdot 2 = 6$ possible total degenerate microstates that account for this example.

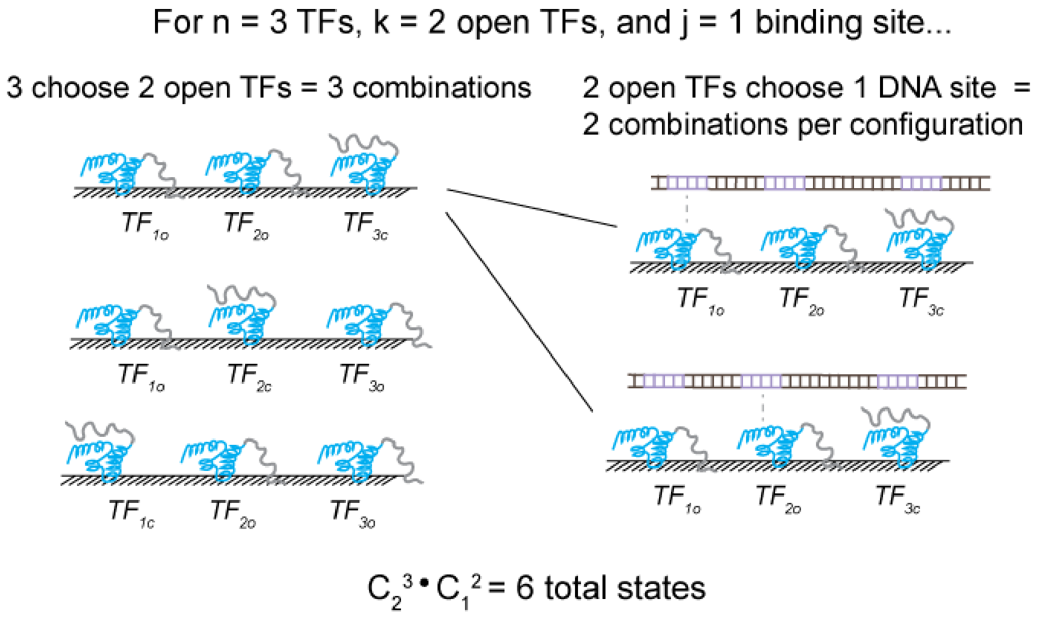


Figure 7: Accounting for all microstates for the example with $n = 3$ TFs, $k = 2$ open TFs, and $j = 1$ DNA site bound

We can then describe $[TF_{total}]$ as the sum of $[TF_{bound}]$ and $[TF_{unbound}]$, each of which can in turn be described as sums of the weights and multiplicities for each state. $[TF_{unbound}]$ is the sum of column 0 of Figure 5, and $[TF_{bound}]$ is the sum of columns 1 through $n$. We will also normalize all of these weights to the concentration of the state in which all $n$ TFs are in closed conformation, $[TF_{c_n}]$, since this is a constant factor in all weights (Figure 5). Using the binomial theorem as in section 1.2, $[TF_{unbound}]$ can thus be described as the following:

$$\frac{[TF_{unbound}]}{[TF_{cn}]} = \sum_{k=0}^{n} C_k^n (\frac{1}{K_{auto}})^k = (1 + \frac{1}{K_{auto}})^n$$

To describe $[TF_{bound}]$, we will use our shorthand from section 1.2 again and set $x = \frac{[D]}{K_d}$ and $y = \frac{C_{eff}}{K_d}$. In this case, $[TF_{bound}]$ is the sum of all possible conformational states (1) given $k$ open TFs out of $n$ total TFs, where $k$ goes from $k = 1$ to $n$, and (2) given $j$ TFs bound to DNA out of $k$ open TFs total, where $j$ goes from $j = 1$ to $k$. In this way, we sum across the rows and columns of our tables in Figure 5.

$$\frac{[TF_{bound}]}{[TF_{cn}]} = x \sum_{k=1}^{n} C_k^n (\frac{1}{K_{auto}})^k [\sum_{j=1}^{k} C_j^k (y)^{j-1}]$$

Since $\sum_{j=1}^{k} C_j^k (y)^{j-1} = \frac{1}{y}[\sum_{j=0}^{k} C_j^k (y)^j - 1]$, we can use the binomial theorem again as in section 1.2 to simplify the equation above to:

$$\frac{[TF_{bound}]}{[TF_{cn}]} = \frac{x}{y} \sum_{k=1}^{n} C_k^n (\frac{1}{K_{auto}})^k [(1+y)^k - 1]$$

We can then split this expression into two sums ($\frac{x}{y}[\sum_{k=1}^{n} C_k^n (\frac{1+y}{K_{auto}})^k - \sum_{k=1}^{n} C_k^n (\frac{1}{K_{auto}})^k]$), use the fact that $\sum_{k=1}^{n} C_k^n (z)^k = \sum_{k=0}^{n} C_k^n (z)^k - 1$ (with $z$ here representing any variable), and finally use the binomial theorem to simplify this equation again to get the following:

$$\frac{[TF_{bound}]}{[TF_{cn}]} = \frac{x}{y}((1 + \frac{1+y}{K_{auto}})^n - (1 + \frac{1}{K_{auto}})^n)$$

We then can set the fraction bound $\theta = \frac{\frac{[TF_{bound}]}{[TF_{cn}]}}{\frac{[TF_{unbound}]}{[TF_{cn}]} + \frac{[TF_{bound}]}{[TF_{cn}]}} = 0.5$ and plug in our expressions thus far:

$$\theta = \frac{1}{2} = \frac{\frac{x}{y}((1+\frac{1+y}{K_{auto}})^n - (1+\frac{1}{K_{auto}})^n)}{(1+\frac{1}{K_{auto}})^n + \frac{x}{y}((1+\frac{1+y}{K_{auto}})^n - (1+\frac{1}{K_{auto}})^n)}$$

Solving this equation for x, which contains our $[D]$ term, substituting in our values for $x$ and $y$, and then subsequently solving for $[D]$ yields the following:

$$K_{d,obs} = [D] = \frac{C_{eff}(1+\frac{1}{K_{auto}})^n}{(1+\frac{1+\frac{C_{eff}}{K_d}}{K_{auto}})^n - (1+\frac{1}{K_{auto}})^n}$$

We can further simplify this and factor out $K_{auto}$ terms in the denominators since $(1 + \frac{1}{K_{auto}})^n = (\frac{K_{auto}+1}{K_{auto}})^n$, resulting in the following:

$$\mathbf{K_{d,obs}} = \frac{\mathbf{C_{eff}}}{(\mathbf{1}+\frac{\mathbf{C_{eff}}}{\mathbf{K_d(1+K_{auto})}})^{\mathbf{n}}-\mathbf{1}}$$

To further our intuitions, if we assume that $C_{eff} >> K_d(1 + K_{auto})$ similarly as before, then $(1 + \frac{C_{eff}}{K_d(1+K_{auto})})^n - 1 \approx (\frac{C_{eff}}{K_d(1+K_{auto})})^n$ and our equation simplifies to the following:

$$\mathbf{K_{d,obs}} = \frac{(\mathbf{K_d})^{\mathbf{n}}(\mathbf{1}+\mathbf{K_{auto}})^{\mathbf{n}}}{(\mathbf{C_{eff}})^{\mathbf{n-1}}}$$
